# RELAXING SELECTIVE PRESSURES ON DEVELOPMENTALLY COMPLEX INTEGUMENTARY STRUCTURES: FEATHER VANE SYMMETRY EVOLVES IN ADDITION TO BODY MASS AND WING LENGTH AFTER FLIGHT LOSS IN RECENT BIRDS

**DOI:** 10.1101/2023.10.24.563691

**Authors:** Evan T. Saitta, Lilja Carden, Jonathan S. Mitchell, Peter J. Makovicky

## Abstract

Feathers are complex integumentary structures with high diversity across species and within plumage and have varied functions (e.g., thermoregulation, flight). Flight is lost in many crown lineages, and frequently occurs in island ‘founding’ or semiaquatic context. Different extant lineages lost flight across at least three orders of magnitude of time (∼79.58 Ma–15 Ka). Flight loss’s effect on sensory capacity, brain size, and skeletomusculature have been studied, but less work exists on relations between flightlessness and feathers. To understand how flight loss affects feather anatomy, we measured 11 feather metrics (e.g., barb length, barb angle) from primaries, tertials, rectrices, and contour feathers on skins of 30 flightless taxa and their phylogenetically closest volant taxa, supplemented with broader sampling of primaries across all orders of volant crown birds. Our sample includes 27 independent losses of flight; the sample contains nearly half the extant flightless species count and matches its ∼3:2 terrestrial:semiaquatic ratio. Vane symmetry increases in flightless lineages, and these patterns are strongest in flight feathers and weakest in coverts. Greatest changes in feathers are in the oldest flightless lineages like penguins, which show robust filaments (rachis, barbs, and barbules) on small feathers, and ratites, which show high interspecific diversity with plumulaceous filaments and/or filament loss. Phylogenetic comparative methods show that some of these microscopic feather traits, such as barb/barbule length and rachis width, are not as dramatically modified upon flight loss as are body mass increase and relative wing and tail fan reduction, whereas the effect on vane symmetry is more easily detected. Upon relaxing selection for flight, feathers do not soon significantly modify many of their flight adaptations, although increased vane symmetry is likely the most detectable shift. Feathers of recently flightless lineages are in many ways like those of their volant relatives. Feather microstructure evolution is often subtle in flightless taxa, except when flight loss is ancient, perhaps because developmental constraints act upon feathers and/or selection for novel feather morphologies is not strong. Changes in skeletomusculature of the flight apparatus are likely more evident in recently flightless taxa and may be a more reliable way to detect flight loss in fossils, with increased vane symmetry as potentially a microscopic signal. Finally, we see an intriguing, reversed pattern in feather evolution after flight loss from the pattern proposed in popular developmental models of feathers, with the later stages of feather development (asymmetric displacement of barb loci) being lost more readily, while early stages of development (e.g., differentiated barb ridges on follicle collar) are only lost after many millions of years of flightlessness.

## INTRODUCTION

With an estimated diversity of 11,000 to 18,000 species (Barrowclough *et al*. 2016), a global distribution under a wide variety of habitats and ecological niches (Mitchell & Makovicky 2014), and an evolutionary history stretching over 150 million years (von Meyer 1861), birds are one of the most successful vertebrate groups ever. While many adaptations might help explain their evolutionary longevity, diversity, and disparity, such as their elevated metabolism (Legendre & Davesne 2020), a key adaptation certainly are feathers. While often associated with their role in flight, feathers serve a diverse array of functions including thermoregulation (McFarland & Budgell 1970), water repellence (Rijke & Jesser 2011), camouflage (Tökölyi *et al*. 2008), and sexual display (Aparicio *et al*. 2003). Feathers come in a variety of morphologies across species and sexes of a single species, but they also vary within a single individual across the plumage (e.g., flight versus contour feathers) as well as through time across seasons or ontogeny (e.g., neonatal down versus adult plumage) (Lucas & Stettenheim 1972). Feathers are also impressive from a mathematical biology perspective, representing complex fractal structures with several orders of filament (i.e., rachis, barbs, and barbules) branching under the control of precise developmental mechanisms (Ng & Li 2018). Despite their broad utility and complexity, as metabolically inactive (i.e., ‘dead’) extracellular integumentary structures consisting largely of the common ‘keratin’ proteins (Fraser *et al*. 1971) (i.e., α-keratins and corneous β-proteins [Holthaus *et al*. 2018]) along with pigments [Roy *et al*. 2020], secreted lipid waxes [Jacob 1978], and calcium phosphates [Blakey *et al*. 1963], they are presumably not highly energetically demanding on the individual as evidenced by their continuous and regular replacement throughout life (Palmer 1972). These attributes make bird feathers an intriguing evolutionary system to examine.

Another wrinkle in the evolution of feathers relates to their most remarkable function, flight. Flight has been independently reversed in many different lineages of birds throughout their deep evolutionary history (Roots 2006). Even if only examining extant lineages, occurrences of flight loss span at least three orders of magnitude of time, with ostrich (*Struthio camelus*) ancestors having lost flight about 79.58 Ma (Yonezawa *et al*. 2017), while Fuegian steamer ducks (*Tachyeres pteneres*) lost flight about 15 Ka (Fulton *et al*. 2012). Some poor-flying species even appear to be in the process of losing flight and might be described as ‘incipiently flightless’, such as the extant saddleback of New Zealand (*Philesturnus*) (Taylor *et al*. 2007), or the recently extinct, semiaquatic New Zealand merganser (*Mergus australis*) (Livezey 1989). In some cases, there is uncertainty as to whether a species is truly flightless, such as the brown mesite (*Mesitornis unicolor*), which has no reliable record of ever being observed to fly (Roots 2006). While some domestic species are flightless or poor flyers as a result of artificial selection to yield unique, fluffy feathers, such as ‘silkie’ chickens and pigeons (Miller 1956; Feng *et al*. 2014), we do not focus on these instances because they are not only a result of human artificial selection, but also represent flight loss after breeding for novel feather morphologies, rather than flight loss occurring due to ecological selection that then impact subsequent feather evolution. Flight can also be lost during ontogeny – the flightless adult giant coot (*Fulica gigantea*) retains volant juveniles (Fjeldså 1981; Livezey 2003) (coded in this study as a flightless taxon). In contrast, only the heaviest males of the flying steamer duck (*Tachyeres patachonicus*) are flightless (Humphrey & Livezey 1982; Livezey 2003) (coded here as a volant taxon). Furthermore, some taxa are seasonally flightless for certain periods of time due to simultaneous/synchronous molting (Dial & Heers 2021).

Flight loss tends to occur in one of two ecological contexts. Terrestrial flight loss seems to occur when ‘founder’ individuals fly to new regions – often islands without predators, including nest predators (Clout & Craig 1995; Wright *et al*. 2016). Alternatively, many bird lineages lost flight upon a transition to semiaquatic ecologies in fresh or salt water (Elliott *et al*. 2013; Fish 2016). Both scenarios are possibly related to some of the most heavily discussed phenomena in evolutionary biology – island evolutionary dynamics (e.g., proposed examples of island dwarfism or gigantism [Lomolino *et al*. 2013]) and secondarily aquatic convergent evolution (Kelley & Pyenson 2015), respectively.

The effect of flight loss on brain size (Ksepka *et al*. 2020), sensory capacity (Torres & Clarke 2018), and skeletomusculature (e.g., Cubo & Arthur 2001; Watanabe *et al*. 2021) of birds has been previously studied, but less work seems to have been done on feathers themselves, especially at the microscopic level (McGowan 1989). This study is a preliminary examination of feather variation in extant and recently extinct birds to characterize the changes that occur to feathers when flight is lost in these lineages. While the results may have implications for recognizing potential flight capacity in fossils of exceptional preservation (Feduccia & Tordoff 1979), they bear directly on evolutionary questions related to flightlessness in crown group or recent/extant lineages: selection (e.g., relating to predation, locomotion, thermoregulation) and constraints (e.g., developmental of complex fractals). The ecological context of flight loss is also intriguing; for example, if terrestrial flightless bird feathers evolve via predation reduction, while semiaquatic flightless bird feathers evolve via locomotory selection in a novel, dense medium. At what rate do feathers evolve after flight loss, especially in comparison to other morphological changes, such as in the skeletomusculature? Under one hypothesis, feathers, which are highly morphologically plastic over macroevolutionary timeframes, might evolve rapidly when selection for flight and predation pressure are released. Under an alternate hypothesis, metabolically cheap feathers might evolve less rapidly than metabolically expensive bone and muscle of the flight apparatus due to weaker selection on feathers and/or because of development constraints that are inherent in the development of these complex fractal structures.

## MATERIALS AND METHODS

### Sample choice

There are estimated to be roughly 64 species of extant flightless birds, of which ∼59% are terrestrial and ∼41% are semiaquatic; furthermore, ∼28% are penguins (Sphenisciformes) (Roots 2006). Our sample contained 30 taxa known to be fully flightless from nine different orders (Palaeognathae, Gruiformes, Psittaciformes, Charadriiformes, Anseriformes, Podicipediformes, Suliformes, Sphenisciformes, and Passeriformes), nearly half of which are rare or threatened species (IUCN 2022). Specimens were obtained from the bird skin collections of the Field Museum of Natural History (FMNH) and the American Museum of Natural History (AMNH). Our sample is ∼57% terrestrial and ∼43% semiaquatic, similar to the ecological breakdown observed among all extant flightless birds.

We reduce redundancy in our sample (supplemental material) by avoiding the overweighting of single losses of flight among lineages with multiple flightless species that subsequently diversified, such that our sample is only 10% penguins, while broadly covering three of the major penguin lineages (sampled taxa: *Spheniscus*, *Eudyptes*, *Aptenodytes* [Ksepka *et al*. 2006]). Additionally, only one species of each of the five genera of ratites (i.e., flightless Palaeognathae) is included for what would otherwise be 13 species that represent only four independent losses of flight.

Six sampled flightless taxa are recently extinct (*Porzana sandwichensis*, *Porzana palmeri*, *Gallinula nesiotis*, *Xenicus lyalli*, *Podilymbus gigas*, and *Pinguinus impennis*), as well as one recently extinct, poor-flying/‘incipiently flightless’ species (*Mergus australis*).

For each flightless (and ‘incipiently’ flightless) taxon, we measured feather traits in its phylogenetically closest volant taxon. For penguins, we included three volant species of tubenoses (Procellariiformes), covering three of the major lineages within that clade (*Oceanites oceanicus*, *Pelecanoides urinatrix*, and *Diomedea immutabilis*), given that penguins represent a deeply diverging flightless lineage most closely related to a diverse tubenose clade with disparate locomotory habits and ecologies.

Finally, we included a broader sampling of crown birds to obtain at least one species from each commonly recognized order across the entire phylogeny (Reddy *et al*. 2017) (the inclusion of two owl species can be used as a way justify sufficient measurement precision, assuming both species have similar feather morphometrics to each other compared to other taxa). All these broadly sampled species are volant, except for *Mesitornis unicolor*, whose flight capabilities are unknown.

This sampling yielded a total of 83 taxa (307 feathers, as discussed below). Based on the above considerations, we consider our sample to be useful for informing us about flight loss in birds under a phylogenetic framework.

### Data acquisition

A wide range of macro- and microscopic measurements were made on the skins, covering the size/proportions, densities, and positions of various skeletal, scale, and feather traits (see supplemental material for a full listing), and were supplemented with values from the literature (e.g., species body mass was commonly taken from Dunning [2008]).

Measurement tools included calipers, digital calipers, and measuring tape as required for macroscopic traits. As for macroscopic traits, we were particularly interested in body mass, wing length as measured from the wrist to the apical end of the longest primary, and tail fan length as measured from the posterior end of the bony pygostyle to the apical tip of the longest rectrix.

For microscopic observations of feathers, a Dino-Lite Edge AM73915MZTL (10X∼140X; 5.0MP; USB 3.0) Digital Microscope with a calibrated scalebar feature (supplemented with a physical scale bar placed alongside the feather when necessary) was used with a maneuverable Dino-Lite MS36A-C2 adjustable precision mount with clamp. Measurements were then taken from the micrographs using Adobe Photoshop (version 20.0.6).

For each specimen within clades containing flightless extant taxa, five feathers across the body were measured: the longest primary remex, the dorsal-most tertial remex, the dorsal-most rectrix, and dorsal and ventral contours from roughly the middle of the torso. For orders of birds that do not contain known extant flightless taxa, we only examined the longest primary remex for logistical purposes, operating under the assumption that the patterns of flight loss are strongest in these feathers (a hypothesis that, as seen below, was supported by our results). While flight feathers can easily be assigned leading and trailing veins, for internal consistency within our dataset, we refer to the ‘trailing’ vanes of contour feathers as those more medial and the ‘leading’ vanes as those more lateral, even though we understand that these terms are not truly applicable to feathers that do not generate lift.

The feather of interest was carefully manipulated by hand for maximum exposure from the plumage and a piece of paper was placed underneath it to provide contrast for photography. Feather barbs were manipulated as required with a small metal pick and sewing pins to expose them for observation.

For each feather, observations were made at the apical, middle, and basal regions of the exposed extent of the rachis (see supplemental material for complete data table). We focused our analyses on the middle region due to our observation that the apex is often damaged in museum specimens and can have high amounts of individual asymmetry (even between comparable feather positions on the left and right sides of a single individual) and the basal region would sometimes, but not always, include plumulaceous regions of feathers depending on how far down its length the feather could be manipulated and exposed from neighboring plumage.

Specific feather metrics of greatest interest are shown in Fig. 1A. Also note the semi- alternating barb pattern and partial barbule loss along the barb farthest from the rachis observed in the kiwi feather, which could lead to variation in the metrics depending on the specific barb pair measured [Fig. 1B]; for the kiwi feathers studied, a barb pair with at least one well-developed barb was always chosen for measurement. For logistical reasons, only the traits of the barbules and mid- barb width on the trailing vane barb were measured, since the trailing vane has a greater area than the leading vane on each flight feather as it relates to the overall lift surface of the wing. To control for absolute size of the feather and the position along the length of rachis, linear microscopic feather metrics (rachis width, mid-barb width on trailing barb, leading barb length, trailing barb length, and distal barbule length on trailing barb) were divided by the length from the apical most barb divergence/node to the specific barbs that were measured. Proximal barbule length was not examined due to difficulties in manually unfurling the tightly overlapping/imbricating barbules observed in many taxa in a manner that would have been consistent and safe across specimens.

**Figure 1.**
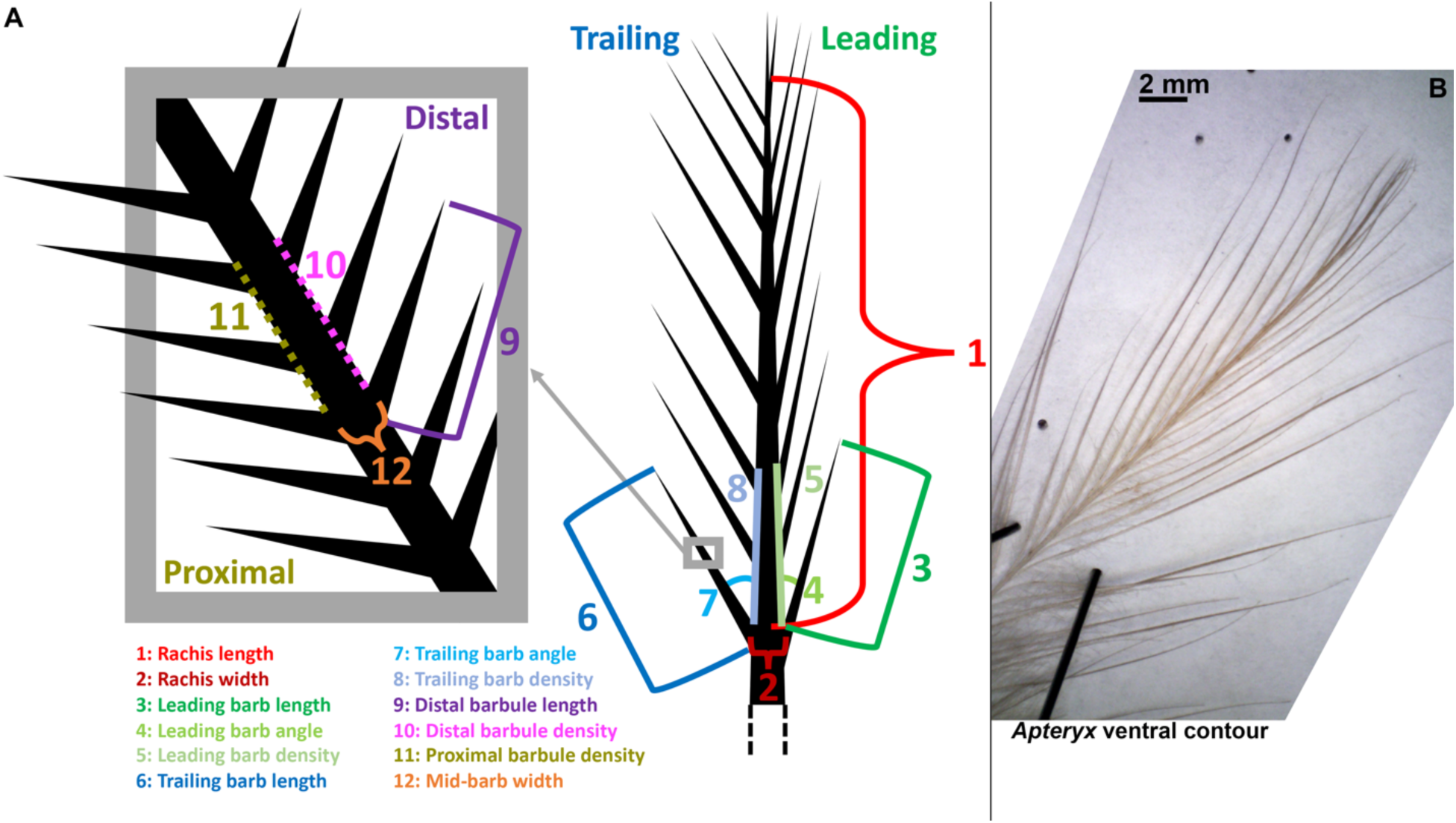
Feather morphology examined in this study. A, Diagram of key microscopic feather metrics measured in this study. Note that we focus on the midpoint of the exposed rachis on the museum skin (i.e., the full extent of the feather exposed from the plumage [dashed lines] would roughly double the length of value #1). B, *Apteryx australis* ventral contour showing the semi-alternating pattern of barb development and partial barbule loss along the length of the barb farthest from the rachis.

To account for changes in barb orientation as a result of manipulation as well as curvature at the base of barbs, barb angle was examined as a measure of vane asymmetry at a given barb pair along the rachis in order to control for any distortion at that position along the rachis or variation in the manner in which angles were measured by an observer on our team from one barb pair to another. Vane asymmetry was calculated by subtracting the angle that the leading barb branches off from the rachis by the angle of its associated trailing barb off from the rachis at that barb pair’s rachis position – consistent with the developmental model of feather asymmetry that involves increased barb angles (Feo & Prum 2014).

Filament densities were measured in absolute values of the number of filaments per unit distance (i.e., the number of barbs or barbules per mm along the rachis or barb, respectively). While this does not consider issues of scaling (e.g., increased isometric scaling of a feather would decrease these density values), it is likely more appropriate based on feather development, in which the feather erupts from the follicle such that the apical regions are the first to form (Prum 1999). Therefore, our approach essentially assumes that larger feathers develop not so much in a geometrically isometric way, but rather by allometry (consistent with models suggested by Maderson & Alibardi 2000; Prum & Brush 2002; Sawyer *et al*. 2003; Prum 2005; Lin *et al*. 2006; Chernova & Kiladze 2019). The absolute density values of filaments thereby reflect the rate at which new branching is initiated in development. Despite the logic of these absolute metrics in a development sense, bear in mind that they can be influenced by scaling, such that similarly shaped feathers can vary in density due to size alone.

See supplemental material for further details and considerations of the data acquisition.

Data, scripts, archiving, etc. is also available at http://github.com/paleomitchelljs/FeatherEvo.

### Analysis

To account for any noise in the data, we employ an analytical approach that embraces triangulation (Munafò & Smith 2018), whereby simple to more sophisticated statistical analyses (e.g., with increasing phylogenetic control), rerun under varying conditions (supplemental material), are expected to converge upon similar patterns so as to detect the most prevalent signals in the dataset. We furthermore emphasize comparing the strength of signals relative each other (i.e., their relative effect sizes), rather than only relying on arbitrary thresholds of statistical significance that can lead to falsely dichotomous declarations that effects are either present of absent (Amrhein *et al*. 2019). Finally, we attempt as much as possible given sample sizes to make broad comparisons expected to yield the most prominent patterns (e.g., volant versus flightless, recent versus ancient divergences, inter-order differences, terrestrial versus semiaquatic). With respect to semiaquatic behavior, we can code a score of 0 to 5 based largely on Fish (2016): 0 as not semiaquatic, 1 as adaptive surface hindfeet paddling (beyond wading or ability to swim), 2 as surface wing paddling/‘steaming’ with surface hindfeet paddling, 3 as underwater hindfeet rowing, 4 as asymmetrical underwater flight, and 5 as symmetrical underwater flight. However, such a fine parsing often leads to very small sample sizes in each category, and so the categories of 1–5 can alternatively be lumped into a single, broader semiaquatic ecological category.

Basic data analyses and statistics were performed using R coding language (RStudio version 1.3.1056). We start by using sister taxon comparison between the phylogenetically closest volant taxa and its associated flightless relative. The difference examined involves subtracting the metric for the volant taxa (e.g., a density or angle symmetry measure) from that of their phylogenetically closest flightless taxa: Δ = Metric_Flightless_ – Metric_Volant_. For penguins, the volant metric is based on the average values of the three tubenoses examined (*Oceanites oceanicus*, *Pelecanoides urinatrix*, and *Diomedea immutabilis*). For principal component analysis (PCA), the prcomp() command was set to scale=TRUE (i.e., data was normalized).

Data was not log-transformed when non-transformable data were being considered, such as when feathers without barbules or barbs have nonapplicable values or values of 0 prior to attempted transformation that would yield undefined values.

### Phylogenetic comparative methods

Finally, we employed a series of phylogenetically controlled comparisons to evaluate how traits changed with the loss of flight. Using the phylogeny from Jetz *et al*. (2012) combined with the detailed Gruiformes phylogeny from Garcia-R. & Matzke (2021), we added recently extinct taxa and their dates to their appropriate branches at a stochastically determined time and we repeated this procedure 100 times to create a set of trees of 82 living (Cassowary dropped because 0 values of various primary metrics cannot be log-transformed) and recently extinct taxa. We then employed stochastic mapping (Revell 2012) to generate timings for the loss of flight along the tree. The transition rates were set to be unidirectional (flight could be lost but not re-evolved) and the ancestral state was fixed as flightless, as the relatively small size of the tree (82 taxa) with an over representation of flightless taxa otherwise led to unrealistically high rates of change.

We then compared a suite of macroscopic traits (body mass, wing length to body mass ratio, tail length to body mass ratio, and tarsus length to body mass ratio) and microscopic traits at the middle region of the exposed primary remex (rachis width, rachis length, barb length, barb density, barb angle for the leading & trailing vanes, and distal barbule density for the trailing vane). We compared the traits between flightless and volant taxa directly. Note that the incipiently flightless *Mergus australis* was coded as volant, while the ‘unknown’ volant or flightless *Mesitornis unicolor* was coded as flightless. We compared the differences between 14 volant and flightless sister pairs in a strict sense (i.e., immediate sister lineages, unlike clades with multiple flightless taxa more closely related to each other than to their closest volant relative [e.g., ratites and penguins]), and we used a set of phylogenetically informed Bayesian regressions to evaluate changes in traits with the onset of flightlessness (Kruschke 2011; de Villemereuil *et al*. 2012; Bürkner 2017, 2018, 2021; Fuentes-G *et al*. 2020).

Our Bayesian regressions took two forms. We performed logistic regressions with phylogenetically structured errors to evaluate how well trait differences predict flightlessness. This hierarchical approach is particularly useful for these data, since there are a large number of predictor variables (i.e., traits), and so by using a Bayesian hierarchical approach we were able to put restrictive priors that shrank most regression estimates towards zero to avoid multiple comparison issues. The only traits with non-zero priors on the marginal effect size were body mass and wing length, since these are a part of the skeletomusculature, which has been established through many prior studies (e.g., Cubo & Arthur 2001; Watanabe *et al*. 2021) to change predictably with the loss of flight.

We analyzed variation in rate in two ways. First, we compared the average Akaike weights of single-rate BM versus two-rate BM from 100 stochastically mapped trees using OUwie (Beaulieu *et al*. 2012). The volant and flightless rates and AICc were retained for each model, and the Akaike weights were calculated for each trait-tree combination. We considered Akaike weights of >80% as strong evidence for one model over another.

Our other approach to rates was fitting a two-slope, two-rate phylogenetic regression using RJAGS following Fuentes-G *et al*. (2020). We performed separate two-slope, two-rate regression for the relationship between wing length and body mass, as well as the relationships between trailing and leading vane barb angles, lengths, and densities.

Each analysis built on the previous sets. Given the exploratory nature of this project, we used the results from each progressively less naïve, but more assumption-laden, analysis to inform our next model. Therefore, our basic comparisons informed which traits we analyzed with immediate sister species comparisons, and the traits that stood out in immediate sister-clade comparisons were then used to build the logistic regression models. The parameters with the strongest signal from the logistic regressions were then used to construct the hierarchical multi-slope model (Fuentes-G et al., 2020). Posterior predictive checks were performed along with convergence diagnostics to evaluate the reliability of the regressions.

## RESULTS

### PCA of linear feather metrics across the entire plumage without phylogenetic control

For PCA, linear data on the rachis, barbs, and barbules was not log-transformed beforehand to allow for the inclusion of taxa that lack barbs or barbules, whose lengths and widths of zero would be undefined under log-transformation (i.e., some ratites [Table 1]), although the data was normalized during PCA in R coding language. Therefore, the PCA here should be used to examine relative positions of the taxa to each other on the morphospace rather than absolute distances of PC values between taxa.

**Table 1.** Barb and distal barbule (at approximately mid-length along the trailing vane’s barb) expression across the observed middle and basal feather regions in ratites with estimated divergence ages from Yonezawa *et al*. (2017) shown. X = presence. O = absence. *Note that *Apteryx* often has appreciable partial barbule loss along the barb farthest from the rachis, even when they are present at mid-length, as well as semi-alternating barb developmental patterns.

|  |  | Middle region |  | Basal region |  |
| --- | --- | --- | --- | --- | --- |
|  |  | Barbs | Barbules<br>at mid-barb<br>(trailing vane) | Barbs | Barbules<br>at mid-barb<br>(trailing vane) |
| ***Struthio***<br>79.68 Ma | Primary | X | X | X | X |
|  | Tertial | X | X | X | X |
|  | Rectrix | X | X | X | X |
|  | Dorsal contour | X | X | X | X |
|  | Ventral contour | X | X | X | X |
| ***Rhea***<br>70.62 Ma | Primary | X | O | X | X |
|  | Tertial | X | X | X | X |
|  | Rectrix | X | X | X | X |
|  | Dorsal contour | X | X | X | X |
|  | Ventral contour | X | X | X | X |
| ***Casuarius***<br>66.69 Ma | Primary | O | O | O | O |
|  | Tertial | X | O | X | X |
|  | Rectrix | X | O | X | O |
|  | Dorsal contour | X | X | X | X |
|  | Ventral contour | X | O | X | O |
| ***Dromaius***<br>66.69 Ma | Primary | X | O | X | X |
|  | Tertial | X | X | X | X |
|  | Rectrix | X | O | X | O |
|  | Dorsal contour | X | X | X | X |
|  | Ventral contour | X | X | X | X |
| ***Apteryx*\***<br>62.01 Ma | Primary | X | X | X | X |
|  | Tertial | X | X | X | X |
|  | Rectrix | X | O | X | X |
|  | Dorsal contour | X | X | X | X |
|  | Ventral contour | X | X | X | X |

PCA results for all the feathers in the sample, including those from clades that do not include flightless taxa (Fig. 2), without any phylogenetic control (i.e., not ΔPC values), one sees that flightless taxa overall have more diverse feather morphologies than do volant taxa in the morphospace (Fig. 2A), including many with shifts seemingly toward long barbs and barbules. The poor flying *Mergus australis* does not show particularly extreme feather morphologies (i.e., PC values near the origin and close to most of the other feathers in the sample). Penguins tend to have relatively wider filaments, especially the rachis, while grebes and rails have some feathers with very long filaments (Fig. 2B). One also observes a shift from relatively shorter/wider filaments to longer/thinner filaments according to feather position in the following order: from primary, rectrix, tertial, dorsal contour, to ventral contour (Fig. 2C). Finally, semiaquatic taxa potentially show slightly wider filaments in remiges and rectrices compared to terrestrial taxa that occupy the bottom edge (i.e., lowest PC2 values in the region of high PC1 values) of the morphospace (Fig. 2D). The results are generally consistent when repeating the PCA under varying initial conditions (i.e., with and without ‘long branch’ clades where flightlessness occurred 10’s of millions of years ago, as well as with and without clades that do not include flightless taxa [supplemental material]).

**Figure 2.**
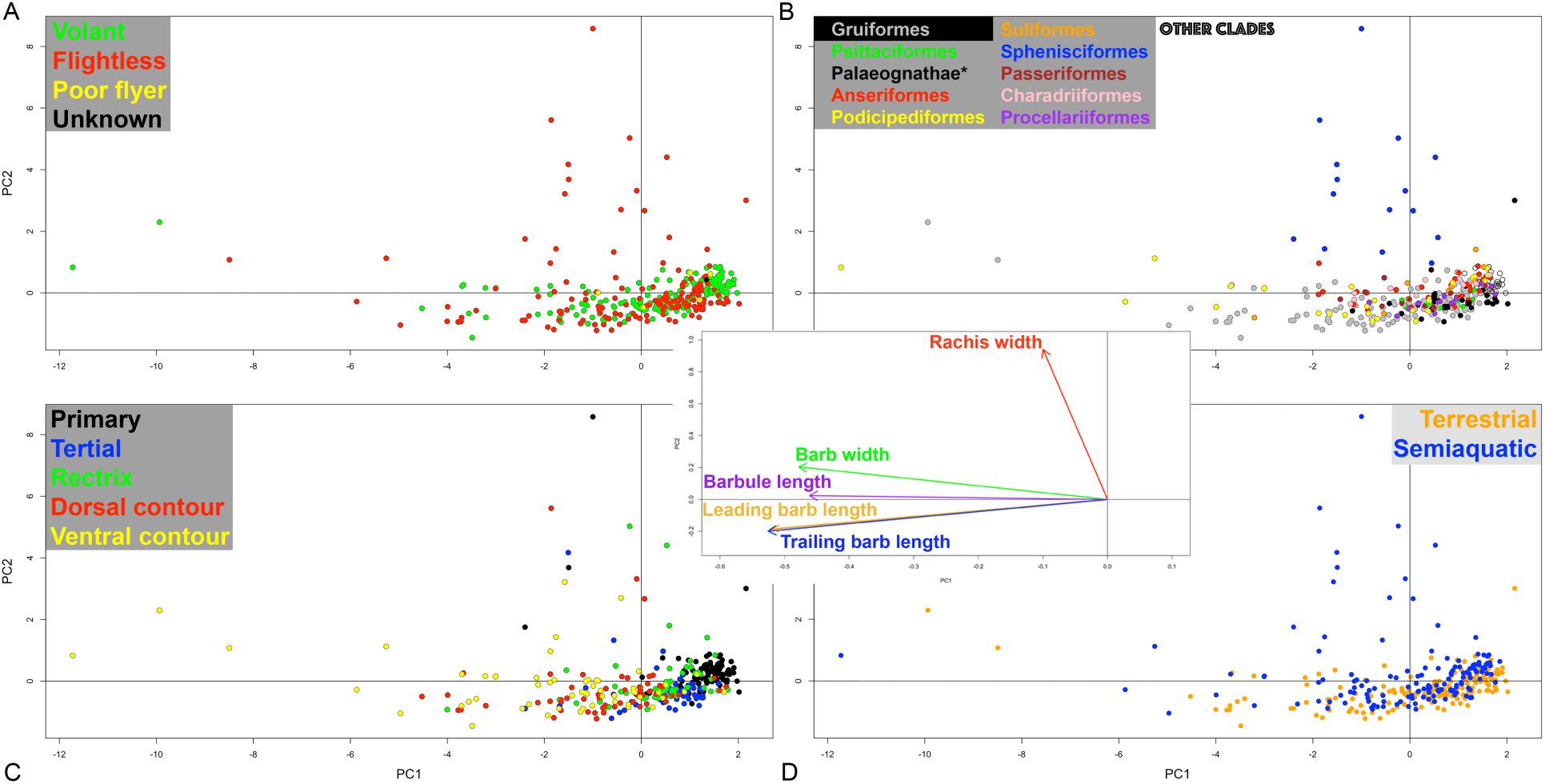
PCA of all feathers studied, including clades without flightless taxa, without phylogenetic control. PC1 and PC2 explain ∼63.63% and ∼21.06% of the variation, respectively. Feathers categorized by, A, flight capability, B, clade (clades without flightless taxa are in open circles; *kiwi are black squares to indicate that their position could vary depending on the barb pair selected due to the semi-alternating barb developmental pattern), C, feather position on the body, and, D, ecology. Center inset shows the loading plot for the five linear metrics on PC1 and PC2. Each of the five metrics is divided by the length of the rachis from the apical-most node to the middle barb pair measured here. N = 307 feathers, 83 taxa.

### Observations of feathers in ancient diverging flightless lineages

Ratites show diverse shifts from their volant tinamou relative in their primaries, with some shifting to shorter filaments or loss of filaments rather than the longer filaments of ostrich barbs and barbules and emu barbs – as exemplified by the thick, quill-like cassowary primaries that lack both barbules and barbs. Other instances of filament loss (Table 1) in the middle regions of ratite primaries include the loss of barbules at approximately mid-length along the barb in rhea and emu. The most recently diverging ratite, the kiwi, can show partial barbule loss along the barb farthest from the rachis as well as a semi-alternating pattern of barbs that are small and weakly developed between more developed barbs (Fig. 1B). The oldest diverging flightless ratites (i.e., ostrich and rhea) are not more likely to have widespread filament loss than the more recent divergences. The more recently diverging emu shows an interesting pattern of barbule loss alongside lengthening barbs.

Penguins tend to shift toward relatively wider filaments from their volant relatives compared to other flightless taxa, yielding small, stiff, scale-like feathers with a very wide rachis.

### Vane asymmetry in sister taxon comparisons

When examining the difference in vane asymmetry, calculated as the asymmetry of the flightless taxa minus the asymmetry of their closest volant relative(s), the strongest shifts related to flightlessness occur in the primaries, whereby we see a decrease in asymmetry with flight loss (Fig. 3C-G). The trend is most prominent in the primaries of ratites, penguins, and potentially auk (although the sample size is limited) (Fig 5.A-B). The pattern seems to be less apparent in ducks, cormorants, and grebes (again, barring limited samples sizes), which are not underwater flyers.

**Figure 3.**
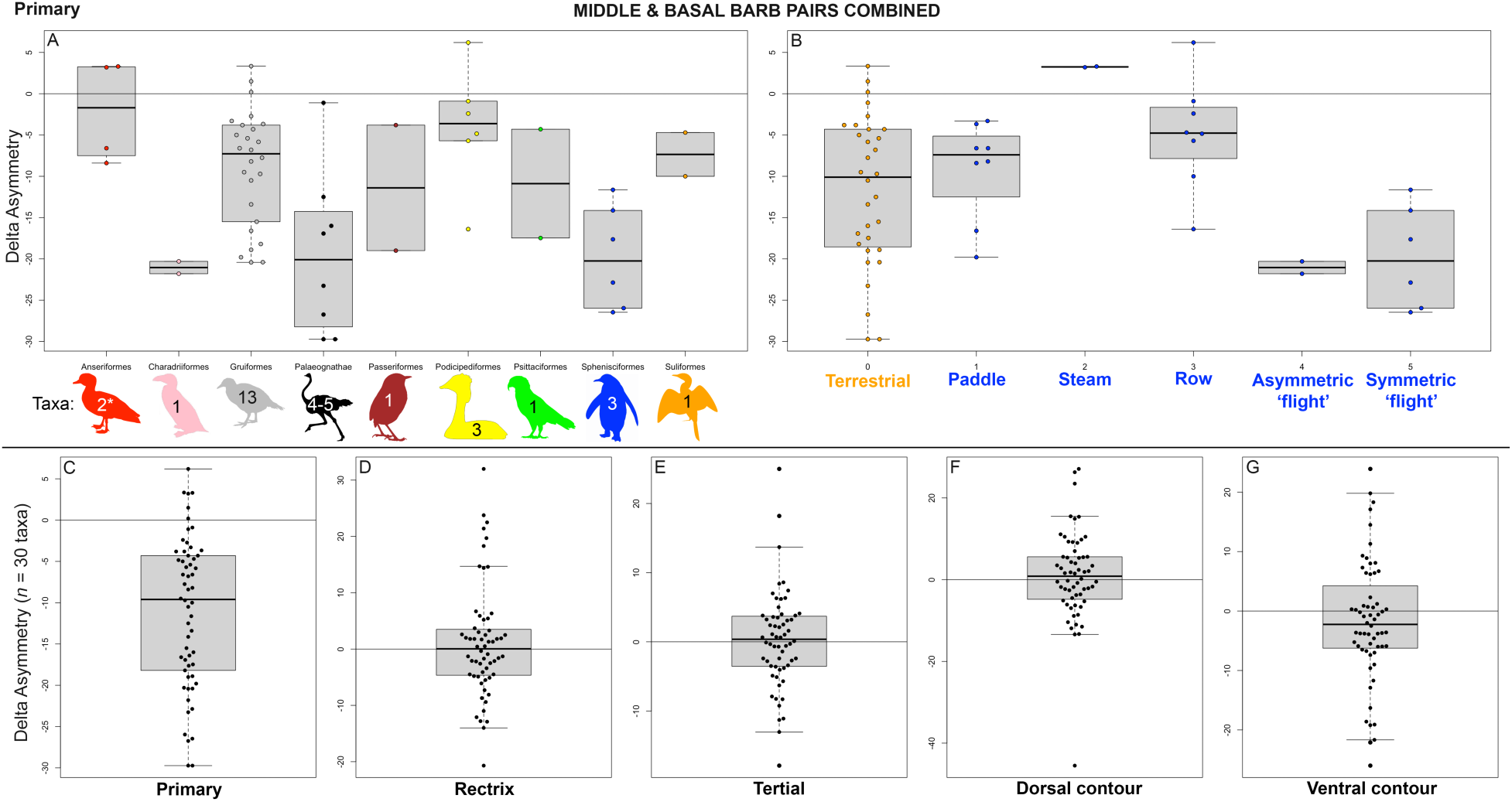
Shifts in vane asymmetry of flightless taxa from their closest volant relative(s) using combined data from the middle and basal barb pairs for the, A, primaries according to clade, B, primaries according to ecology and swim style, as well as whole feather distributions for, C, primaries, D, rectrices, E, tertials, F, dorsal contours, and, G, ventral contours. In flightless semi-aquatic taxa, paddle refers to surface hindlimb paddling, steam refers to surface ‘steaming’ seen in steamer ducks that on occasion use a combination of forelimb and hindlimb surface paddling, row refers to underwater hindlimb rowing, and ‘flight’ refers to underwater forelimb strokes. *The poor-flying/‘incipiently flightless’ *Mergus australis* was not included in Anseriformes. The number of taxa for Palaeognathae varies from 4 to 5 since Cassowary primaries lack barbs and therefore do not have barb angle or vane asymmetry values. N = 60 barb pairs from 30 taxa for all except the primary, which has N = 58 barb pairs from 29 taxa due to the lack of barbs in Cassowary primaries.

**Figure 5.**
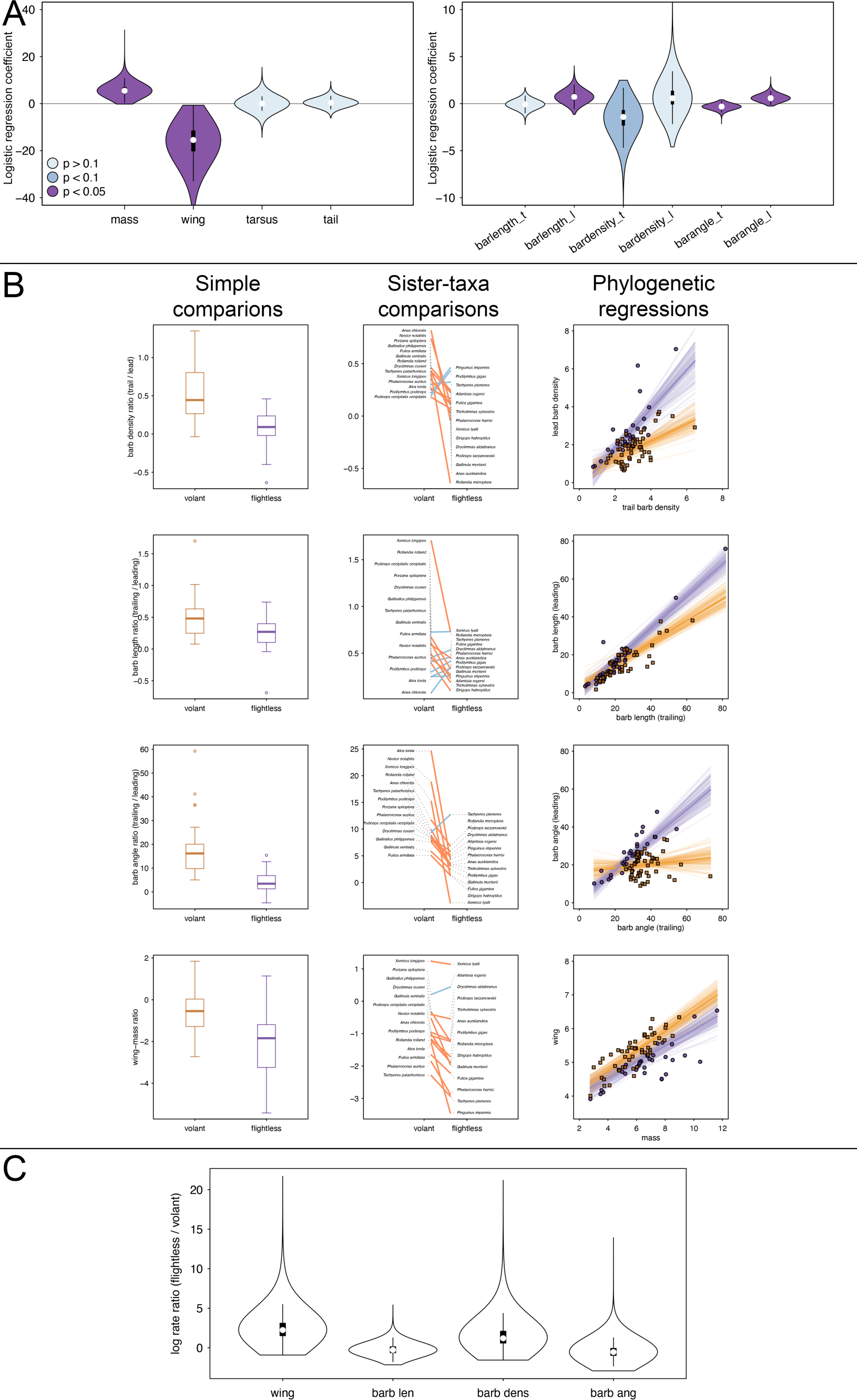
Phylogenetic comparative methods key results summary. A, Logistic regression coefficients. These values reflect the strength and directionality (positive or negative) of the association between macroscopic traits (left: body mass, wing length relative to body mass, tail fan length relative to body mass, tarsus length relative to body mass) or microscopic traits (right: leading/trailing barb density ratio, leading/trailing barb length ratio, rachis width/length ratio) with flightlessness. The spindles indicate the distribution of the posterior estimates for each coefficient from the phylogenetic logistic regression model. Color coding indicates the one-tailed p-value calculated by the proportion of post-burnin samples. B, Tri-plots of barb density ratios (leading/trailing), barb length ratios (leading/trialing), barb angle, and wing length ratios (wing length/body mass), presented as log_10_-transformed values. Left column reports simple comparisons of these measures between volant and flightless taxa as boxplots with outliers plotted. Middle column reports immediate sister-taxa comparisons. Right column reports the multi-slope, multi- intercept phylogenetic regressions between volant (orange) and flightless (purple) taxa, with the shading representing the 1,000 randomly drawn slopes from the posterior. C, Comparison of rates between flightless and volant taxa for wing length to body mass, and trailing/leading barb length, density, and angle. These rates were computed from the multi-slope, multi-rate model of Fuentes- G *et al*. (2020) using rJAGS, and the spindles represent the posterior distribution of the log rate ratio. The poor-flying/‘incipiently flightless’ *Mergus australis* was coded here as volant and the unknown flyer *Mesitornis unicolor* was coded here as flightless. N = 82 taxa for all but the immediate sister-taxa comparisons, which are 14 pairs of volant and flightless comparisons.

*Filament densities in sister taxon comparisons.* When looking broadly at the distributions of shifts in filament densities observed in flightless taxa from their phylogenetically closest volant taxa (i.e., flightless taxon density minus phylogenetically closest volant taxa/taxon’s density), one can see that the densities of both barbs and barbules tend to decrease in terms of their median values (Fig. 4E-R), apart from the leading vane barbs on primaries (Fig. 4A,D). Focusing on the primaries (i.e., the most crucial feather position in our sample as it relates to flight), ratites show the most prominent decreases in barb and barbule densities, while penguins appear to show the most positive shifts in barb densities (Fig. 4B-C).

**Figure 4.**
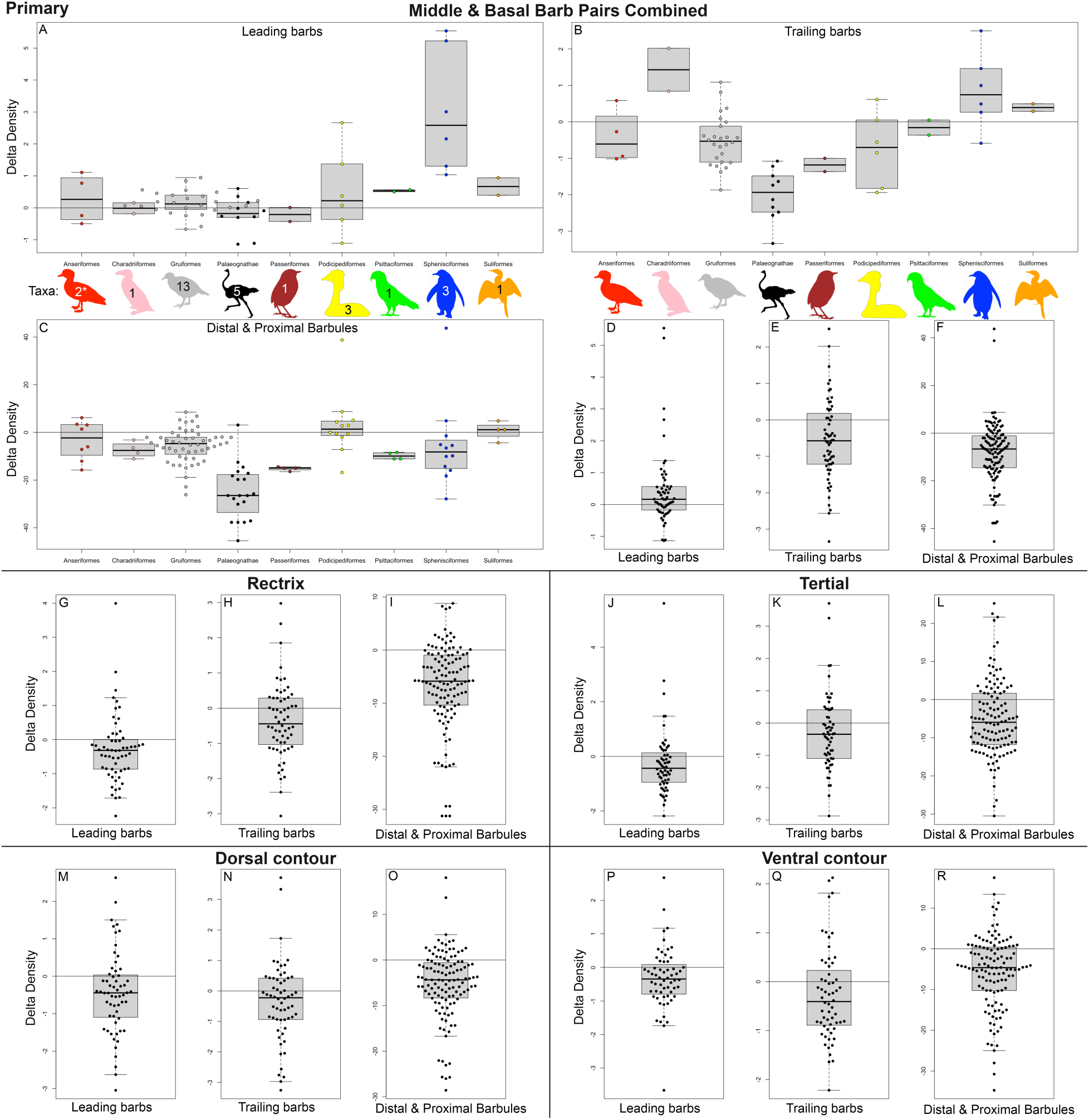
Shifts in filament densities of flightless taxa from their closest volant relative(s) using combined data from the middle and basal barb pairs for, A, primary leading barbs, B, primary trailing barbs, and, C, primary distal and proximal barbules on trailing barbs, as well as the total distributions for the leading barbs, trailing barbs, and distal and proximal barbules on the trailing barbs, respectively, for the, D-F, primaries, G-I, rectrices, J-L, tertials, M-O, dorsal contours, and, P-R, ventral contours. *The poor-flying/‘incipiently flightless’ *Mergus australis* was not included in Anseriformes. N = 60 leading or trailing barb density measures from 30 taxa or 120 distal and proximal barbule density measures on 60 trailing barbs from 30 taxa.

### Phylogenetic comparative methods

Hypothesized macroscopic and microscopic morphological changes linked to the loss of flight from regressions of sister taxon comparisons (supplemental material) were more rigorously tested using more sophisticated phylogenetic comparative methods. Note that the small number of taxa for each of the diverse semi-aquatic locomotory categories limited our ability to analyze flight loss in semi-aquatic taxa using these methods.

### Macroscopic traits

Both logistic regression and immediate sister clade comparison support the hypotheses that body mass increases upon flight loss and wing length relative to body mass decreases upon flight loss (Fig. 5).

The hypothesis (supplemental material) that tail fan length relative to body mass decreases upon flight loss is not supported by logistic regression, with the same being true for changes in tarsus length relative to body mass, but it is supported by immediate sister taxa comparison.

These patterns described in the above macroscopic traits appear to be consistent even for recently diverging flightless taxa (i.e., when dropping ratites and penguins from regression). This consistency in recently diverging flightless taxa is further supported by immediate sister clade analyses for body mass and wing length (at least relative to body mass).

However, these patterns described in the above macroscopic traits cannot as confidently be said to occur more rapidly after flight loss in semi-aquatic taxa compared to terrestrial taxa. The problem arises from the fact that there are not enough taxa relative to the fine parsing of aquatic locomotion categories that might be of relevance.

### Microscopic traits

Microscopic trait measures were examined under phylogenetic comparative methods at the middle region of exposed primary remiges. The full table of logistic regression coefficients are shown in Table 2. The hypothesis (supplemental material) that rachis width generally decreases upon flight loss is not well supported by logistic regression; the effect is functionally zero but is consistent with a decrease (-0.23 +/- 1.37). However, the hypothesis is partially supported by immediate sister taxa comparison; the middle region of the primary’s rachis width-to-length ratio generally decreases for flightless species, indicating a narrower rachis for the same length up the feather towards it’s calamus.

**Table 2.** Regression coefficient estimates, error, 95% confidence intervals, and chain convergence (R-hat) for the phylogenetic logistic regression. The estimate for the phylogenetic group effect standard deviation was 0.01 +/- 0.02.

|  | Estimate | Est. Error | l-95% CI | u-95% CI | R-hat |
| --- | --- | --- | --- | --- | --- |
| Intercept | -3.59 | 1.46 | -6.95 | -1.25 | 1 |
| mass | 5.72 | 2.02 | 2.54 | 10.36 | 1 |
| wing | -16.34 | 6.55 | -31.5 | -6.4 | 1 |
| tail | 0.48 | 1.16 | -1.6 | 3.09 | 1 |
| tarsus | 0.07 | 1.39 | -2.7 | 2.98 | 1 |
| Rachis width | -0.23 | 1.37 | -3.25 | 2.33 | 1 |
| Rachis length | -0.16 | 0.12 | -0.4 | 0.06 | 1 |
| Barb density (lead) | 0.73 | 1.21 | -1.21 | 3.59 | 1 |
| barb density (trail) | -1.61 | 1.28 | -4.64 | 0.37 | 1 |
| Barb length (lead) | 0.75 | 0.49 | -0.11 | 1.81 | 1 |
| Barb length (trail) | -0.08 | 0.34 | -0.78 | 0.58 | 1 |
| Barb angle (trail) | -0.33 | 0.22 | -0.82 | 0.04 | 1 |
| Barb angle (lead) | 0.62 | 0.28 | 0.17 | 1.26 | 1 |
| Barbule density (distal) | -0.2 | 0.14 | -0.5 | 0.05 | 1 |

The hypothesis that barb length generally increases upon flight loss (supplemental material) is more complicated, as leading barb length shows a significant increase while trailing barb length is estimated to change little (Table 2; Figure 5). Barb density shows a similar leading versus trailing pattern, although the increase in density on the leading margin is not significant (Table 2, Figure 5). Barb angle showed the strongest contrasting response, with trailing angles decreasing while leading angles increased with the loss of flight (Table 2; Figure 5).

### Rates of macroscopic versus microscopic trait evolution

We hypothesized that, upon flight loss, the macroscopic traits typically shift earlier and with greater effect size (i.e., more prominently) than microscopic feather traits (supplemental material). The average Akaike weight of the two-rate BM model was above 90% for all traits analyzed except tarsus length (87%) and tail fan length (78%). Logistic regressions show significant change for wing length and body mass with the evolution of flightlessness (shorter wings and greater mass in flightless birds [Table 2; Figure 5A]). The relationship between wing length and body mass showed the greatest change in rate (Figure 5B).

Based on our two-slope, two-rate regressions, wing length relative to body size showed the strongest rate increase, with >99% of post-burnin samples showing an increased rate (average rate of 0.002 for flightless, 0.0004 for volant; log ratio of 2.5). Barb density between trailing and leading vanes showed a comparable degree of rate change, with an increased flightless rate in >93% of post-burnin samples and a rate change of 0.003 for volant and 0.008 for flightless taxa (log ratio of 1.6). On the other hand, barb length (flightless rate 0.29, volant 0.37; log ratio -0.22) and barb angle (flightless 0.44, volant 0.78; log ratio -0.57) of the trailing and leading vanes showed slower rates in flightless species, although the results were less clear (decreased rate in 68% and 78% of samples for length and angle, respectively).

## DISCUSSION

### Flightless taxa have more diverse feather morphologies, and flight feathers are quantifiably more robust than contours

Initial examination of the data from PCA of linear feather metrics across the entire plumage without phylogenetic controls might lead one to hypothesize that the greater overall diversity of feather morphologies in flightless taxa indicates a release of selective pressure from flight requirements as well as the exaptation of feathers previously used primarily in flight to alternative primary functions, such as thermoregulation or swimming. The pattern from primary, rectrix, tertial, dorsal contour, to ventral contour in terms of shorter/wider filaments to relatively longer/thinner filaments indicates that the metrics observed here do indeed capture functional signals, with more robust feathers (i.e., short filaments relative to rachis width) presumably playing a greater role in locomotion (whether that be flight or swimming) and less robust feathers possibly playing a greater role in thermoregulation. This pattern is consistent with the observation that the least robust remiges and rectrices (i.e., the edge of the data cluster with the lowest PC2 values in the morphospace, but with high PC1 values) are from terrestrial taxa rather than semiaquatic taxa, the latter of which often use their feathers for locomotion in the dense medium of water even when they are not flying.

### Extreme feather modification in ancient diverging flightless lineages

Ratites show diverse shifts in primary feather morphology with some shifting towards long filaments, while others reduce their filaments, sometimes to the point of filament loss (e.g., cassowary primaries lack barbules and barbs). The diversity of morphologies in ratite feathers likely indicates a release from selective pressure, since their forelimbs and pectoral gridles are often greatly reduced and are minimally used (Cubo & Arthur 2001; Nudds *et al*. 2004; Habib & Ruff 2008; Novas *et al*. 2020; Serrano *et al*. 2020; Lowi-Merri *et al*. 2021), while enough evolutionary time has passed to allow for any developmental constraints on feather growth to be overcome. The loss of filaments (i.e., the loss of fractal branching) may indicate that the feathers are beginning to function more analogously to the morphologically simpler hair of mammals, playing a role solely in thermoregulation, camouflage, abrasion resistance, or water repellency rather than locomotion.

The wide filaments, and especially a wide rachis, in penguins might be interpreted as a result of the dense medium of water as well as a shift towards small, more scale-like feathers as the wing is adapted into more of a hydrodynamic flipper.

### Barb angles support an increase in flight feather vane symmetry upon flight loss

Our results when examining vane asymmetry in sister taxon comparison are consistent with previous reports that vane asymmetry is an adaptation for flight (e.g., Feo *et al*. 2015), since the primaries, but not the other feather positions less crucial for flight, of flightless taxa are appreciably more symmetric than their closest volant relative(s). The fact that the most prominent shifts toward greater vane symmetry seem to occur in ratites and penguins is unsurprising given that they represent end- member examples of highly adapted terrestrial taxa and underwater flyers with long evolutionary histories of flightlessness. Ratites long lost the locomotor function of the forelimb, while underwater ‘flyers’, such as penguins, ultimately shifted their wing into more of a flipper (i.e., a relatively fleshier ‘waterfoil’, rather than a purely keratinous airfoil).

### Filament densities decrease in absolute measure upon flight loss, but scaling effects confound, while the signal of increased vane symmetry persists

When examining filament densities in sister taxon comparison, a general tendency to decrease filament densities upon losing flight is possibly consistent with the release of selection pressure for lift and thrust on a tightly packed and well- zipped feather vane that resists against low density air during flight. All feather positions show density decreases, and in terrestrial flightless taxa, this might reflect a release from selection pressure for aerodynamic plumage across the body. The fact that semiaquatic flightless taxa also appear to share this trend towards decreased filament densities might be reflective of the higher density of water than air as well as the cohesive ability of water molecules, and therefore feathers used in a hydrofoil require less tight barb and barbule packing than those used in an airfoil. Alternatively, wider (and stiffer) barbs that are less densely packed might be selected for in semiaquatic flightless taxa against the dense water medium.

When it comes to the pattern of strongly negative shifts in filament densities in ratites and relatively most positive shifts in penguin barb densities, again, bear in mind that some of this signal is in part driven by scaling effects (i.e., ratites other than kiwis can have very large feathers, while penguins have very small feathers). Given that overall body mass also tends to increase with flightlessness, this caveat of feather size cannot be readily dismissed. Interestingly, penguin feathers have been shown to be not just hydrophobically water-shedding, but also ice-shedding due to tight interweaving of filaments, down to the hooklets/barbicels of the barbules, that lead to anti-icing through interfacial cracking and therefore low adhesion of ice (Wood *et al*. 2022). This is consistent with their shift to high filament densities (in absolute measures).

At the very least, this filament density data provides even further evidence for the impact of flight loss on vane symmetry. The pattern of increased barb densities in the leading vanes of primaries is unsurprising given that, as we saw above, feather asymmetry decreases upon flight loss, and higher branching angles allow for easier packing of barbs along the rachis.

### Tight phylogenetic controls shows that body mass, wing length, and feather symmetry are key traits impacted by flight loss

The more sophisticated phylogenetic comparative methods generally support the results from the analyses above, but especially help to identify the most prominent evolutionary changes upon the loss of flight. Body mass increase and wing length (relative to body mass) decrease are most strongly associated with flight loss on the macroscopic level, whereas indicators of an increase in vane symmetry observed in the ratios of leading/trailing barb lengths, angles, and densities are most strongly associated with flight loss on the microscopic level (e.g., more so than an increase in barb or barbule length). In fact, the correlation between these indicators of increased vane symmetry and flight loss, as well as increased body mass and decreased wing length (relative to body mass) and flight loss, appears repeatedly across our series of analyses. Notably, the pattern is complicated by which side of the vane is analyzed, as the relationship between trailing and leading barb density and angle changes with the evolution of flightlessness.

The rates of evolution for the most strongly correlated traits with flightlessness observed here (i.e., body mass, wing length to body mass ratio, tail fan length to body mass ratio, tarsus length to body mass ratio, rachis width to length ratio, and ratios of leading/trailing barb length and density) indicate that the loss of flight does indeed induce for novel evolution in these traits, likely as a result relaxed selection pressures relating to flight and a shift towards primary functions other than flight (e.g., swimming, thermoregulation).

### Summary of key results

This is the first time that morphometric feather data of this nature has been collected from a diverse sample of volant and flightless extant and recently extinct birds. Overall, our different analyses all converge upon the conclusion that feathers can adapt radically upon flightlessness (e.g., the quill-like cassowary primary), but that the rates of feather evolution upon flight loss are not necessarily fast.

### Feather developmental stages and flight loss

Our results have interesting implications on how evolutionary changes in feathers after flight loss might relate to underlying developmental processes. If we assume a model of feather development along the lines of Prum & Brush (2002) and Prum (2005), then we find that the changes in feather morphology after flight loss appear to occur in reverse order of this ‘evo-devo’ model. Stage Va (asymmetric flight feathers) is the most readily lost stage after a lineage loses flight, reverting to symmetric Stage IV. This would imply a loss of lateral displacement of new barb loci in the feather follicle. After lineages have been flightless for many millions of years, as seen in the feathers of various ratites, vanes can change from closed to open (reverting from Stage IV to IIIa+b with the loss of differentiated distal and proximal barbules) and barbules can be lost (reverting from Stage IIIa+b to Stage IIIa with a loss of periphal barbule plates). Interestingly, the primary feathers of cassowaries, which show a loss of barbs to result in a quill-like, bare rachis show a further reversion from Stage IIIA all the way back to Stage I, with a loss of barb ridges completely. It would appear, at least based upon this model, that the later stages of feather development (e.g., lateral displacement of barb loci in the follicle) are more liable to loss than are early stages (e.g., barb ridge differentiation in the follicle collar).

## CONCLUSION

Upon flight loss we clearly observe an increase in body mass, while the length of the wings decrease relative to body mass. Without selection for flight, the upper limit on body mass can be easily raised, further compounded by the supportive buoyancy of water on semiaquatic flightless taxa. In a classic scenario for terrestrial flightlessness, namely a newly colonized island without predators, limited island resources and land area likely select against the maintenance of costly flight-related skeletomusculature and limb development shifts away from the forelimbs. In semiaquatic contexts of flight loss, wings possibly shorten to become stout paddles, requiring less surface area to generate lift and thrust in the dense water medium and increased buoyancy, or alternatively, locomotion is instead shifted towards the hindlimbs for paddling/rowing taxa.

In contrast to these macroscopic traits, many feather microstructural traits, with the exception of vane symmetry, appear to be less susceptible to predictable or prominent change, at least early on in evolution. This is perhaps unsurprising given that, as extracellular integumentary ‘keratin’, feathers are likely not as metabolically costly as flight skeletomusculature, as evidenced by their continual shedding and regeneration, and so selection is not as intense to alter feathers in flightless taxa. Alternative to, or perhaps in conjunction with, a selection hypothesis is a developmental constraint hypothesis, where the complex developmental mechanisms by which these fractal integumentary structures grow are not susceptible to rapid evolution. With respect to locomotory selective pressures, semiaquatic flightless taxa, unlike terrestrial flightless taxa, experience a combination of a release from selection for flight and new selection for swimming.

Ultimately, the most prominent microscopic changes after the loss of flight seem to occur in the flight feathers, especially the primaries, where vane asymmetry decreases. While we might hypothesize that, early in the evolution of flightless taxa, primary remex filaments tend to increase in length (supplemental material) and, in semiaquatic flightless taxa, that there is an increase in rachis width (perhaps to better maintain feather shape in dense water media for forelimb-driven steaming/swimming), these patterns, if present, are clearly not as prominent or detachable as changes in vane asymmetry.

Later in the evolution of flightless taxa (i.e., as seen in taxa that diverged from their volant ancestor the least recently), we observe the most prominent shift to symmetric vanes. Penguins evolve very robust, scale-like feathers with wide filaments that are small in absolute size and have high filament density in absolute measure as a result. Their hydrofoil for lift and thrust during swimming consists relatively more of flesh than of feather. Ratites show diverse feather morphotypes – some with large feathers in absolute size (kiwis being an exception) and a shaggy overall plumage, alongside an overall trend for low filament density in absolute measure as a result. Ostrich have very plumulaceous primary remiges, while cassowary feathers tend to lose many barbules and even barbs in their primaries. Shaggy plumage and the simplification of feathers in ratites (i.e., other than the primaries of ostrich, for example, the other feathers of ratites across the body often tend toward filament reduction) suggest that terrestrial flightless bird feathers might begin to function more akin to non-branching mammalian hair given sufficient evolutionary time to overcome any developmental constraints.

Implications of these results include the possibility that some signatures of secondary flightlessness are consistent across independent losses of flight, but there may be a difference based on ecological context (i.e., terrestrial versus semiaquatic). Furthermore, it indicates that for extinct taxa or recently deceased specimens, skeletal anatomy rather than feather anatomy (perhaps with the exception of vane symmetry) is probably the best indicator of flight loss, at least if flight loss was recent. This result is consistent with prior studies, for example, showing a relationship in birds between coracoid strength and wing-beat propulsion (Akeda & Fujiwara 2022). However, we hypothesize that somewhat predictable changes in feathers do occur, especially given sufficient evolutionary time. Interestingly, feather evolution after flight loss appears to occur in the reverse order of commonly invoked developmental models for feathers.

## Acknowledgements

Research was supported by the Bass postdoctoral research fund at the Field Museum. We thank the following at the FMNH for access to the bird collections and helpful discussion: Ben Marks, John Bates, and Shannon Hackett. Furthermore, we thank the following at the AMNH for access to their bird collections and helpful discussion: Bentley Bird and Paul Sweet. We also thank Andrew Romanelli for assistance in data collection. Finally, we thank the Jeff Metcalf Internship Program (University of Chicago) for supporting LC.

Silhouettes in main text and supplement are from phylopic.org: *Pinguinus impennis* modified from John James Audubon (Public Domain Mark 1.0), *Xenicus* by Wynston Cooper (photo) and Albert Chen (a.k.a. “Albertonykus”) (silhouette) (CC BY-SA 3.0), *Aptenodytes* by Steven Traver (Public Domain Mark 1.0), *Phalacrocorax* by L. Shyamal (CC BY-SA 3.0), *Podiceps* by Doug Backlund (photo), John E. McCormack, Michael G. Harvey, Brant C. Faircloth, Nicholas G. Crawford, Travis C. Glenn, Robb T. Brumfield & T. Michael Keesey (CC BY-SA 3.0), Anatidae by Rebecca Groom (CC BY-SA 3.0), *Apteryx australis* by Steven Traver (Public Domain Mark 1.0), *Casuarius* by Ferran Sayol (Public Domain Mark 1.0), *Dromaius novaehollandiae* by Darren Naish (vectorized by T. Michael Keesey) (CC BY-SA 3.0), *Struthio camelus* by Ferran Sayol (Public Domain Mark 1.0), Rhea americana by Ferran Sayol (Public Domain Mark 1.0), *Nestor notabilis* by Ferran Sayol (Public Domain Mark 1.0), Gallirallus australis by T. Michael Keesey (vectorization) and HuttyMcphoo (photography) (CC BY-SA 3.0).

## LITERATURE CITED

Akeda, T. and Fujiwara, S.I., 2022. Coracoid strength as an indicator of wing-beat propulsion in birds. Journal of Anatomy.

Aparicio, J.M., Bonal, R. and Cordero, P.J., 2003. Evolution of the structure of tail feathers: implications for the theory of sexual selection. Evolution, 57(2), pp.397–405.

Amrhein V, Greenland S, McShane B. 2019. Scientists rise up against statistical significance. Nature 567: 305–307.

Barrowclough, G.F., Cracraft, J., Klicka, J. and Zink, R.M., 2016. How many kinds of birds are there and why does it matter?. PLoS One, 11(11), p.e0166307.

Beaulieu, J.M., Jhwueng, D.C., Boettiger, C. and O’Meara, B.C., 2012. Modeling stabilizing selection: expanding the Ornstein–Uhlenbeck model of adaptive evolution. Evolution, 66(8), pp.2369–2383.

Blakey, P.R., Earland, C. and Stell, J.G.P., 1963. Calcification of keratin. Nature, 198(4879), pp.481–481.

Bürkner, P.C., 2017. brms: An R package for Bayesian multilevel models using Stan. Journal of statistical software, 80, pp.1–28.

Bürkner P (2018). “Advanced Bayesian Multilevel Modeling with the R Package brms.” The R Journal, 10(1), 395–411.

Bürkner P (2021). “Bayesian Item Response Modeling in R with brms and Stan.” Journal of Statistical Software, 100(5), 1–54.

Chernova, O.F. and Kiladze, A.B., 2019. Heterochrony as the basis for inter-and intraspecific diversity of skin in vertebrates. Biology Bulletin Reviews, 9, pp.174–189.

Clout, M.N. and Craig, J.L., 1995. The conservation of critically endangered flightless birds in New Zealand. Ibis, 137, pp.S181–S190.

Cubo, J. and Arthur, W., 2001. Patterns of correlated character evolution in flightless birds: a phylogenetic approach. Evolutionary Ecology, 14(8), pp.693–702.

de Villemereuil, P.D., Wells, J.A., Edwards, R.D. and Blomberg, S.P., 2012. Bayesian models for comparative analysis integrating phylogenetic uncertainty. BMC Evolutionary Biology, 12, pp.1–16.

Dial, K.P. and Heers, A.M., 2021. Waxing and Waning of Wings. Trends in Ecology & Evolution. Dunning, J.J.B. 2008. CRC Handbook of Avian Body Masses, 2nd ed.; CRC Press: Boca Raton, FL, USA.

Elliott, K.H., Ricklefs, R.E., Gaston, A.J., Hatch, S.A., Speakman, J.R. and Davoren, G.K., 2013. High flight costs, but low dive costs, in auks support the biomechanical hypothesis for flightlessness in penguins. Proceedings of the National Academy of Sciences, 110(23), pp.9380–9384.

Feduccia, A. and Tordoff, H.B., 1979. Feathers of *Archaeopteryx*: asymmetric vanes indicate aerodynamic function. Science, 203(4384), pp.1021–1022.

Feng, C., Gao, Y., Dorshorst, B., Song, C., Gu, X., Li, Q., Li, J., Liu, T., Rubin, C.J., Zhao, Y. and Wang, Y., 2014. A cis-regulatory mutation of PDSS2 causes silky-feather in chickens. PLoS genetics, 10(8), p.e1004576.

Feo, T.J. and Prum, R.O., 2014. Theoretical morphology and development of flight feather vane asymmetry with experimental tests in parrots. Journal of Experimental Zoology Part B: Molecular and Developmental Evolution, 322(4), pp.240–255.

Feo, T.J., Field, D.J. and Prum, R.O., 2015. Barb geometry of asymmetrical feathers reveals a transitional morphology in the evolution of avian flight. Proceedings of the Royal Society B: Biological Sciences, 282(1803), p.20142864.

Fish, F.E., 2016. Secondary evolution of aquatic propulsion in higher vertebrates: validation and prospect. Integrative and comparative biology, 56(6), pp.1285–1297.

Fjeldsa, J., 1981. Biological notes on the giant coot *Fulica gigantea*. Ibis, 123(4), pp.423–437.

Fraser, R.D.B., MacRae, T.P., Parry, D.A.D. and Suzuki, E., 1971. The structure of feather keratin. Polymer, 12(1), pp.35–56.

Fuentes-G, J.A., Polly, P.D. and Martins, E.P., 2020. A Bayesian extension of phylogenetic generalized least squares: Incorporating uncertainty in the comparative study of trait relationships and evolutionary rates. Evolution, 74(2), pp.311–325.

Fulton, T.L., Letts, B. and Shapiro, B., 2012. Multiple losses of flight and recent speciation in steamer ducks. Proceedings of the Royal Society B: Biological Sciences, 279(1737), pp.2339–2346.

Garcia-R, J.C. and Matzke, N.J., 2021. Trait-dependent dispersal in rails (Aves: Rallidae): Historical biogeography of a cosmopolitan bird clade. Molecular Phylogenetics and Evolution, 159, p.107106.

Habib, M.B. and Ruff, C.B., 2008. The effects of locomotion on the structural characteristics of avian limb bones. Zoological Journal of the Linnean Society, 153(3), pp.601–624.

Holthaus, K.B., Eckhart, L., Dalla Valle, L. and Alibardi, L., 2018. Evolution and diversification of corneous beta-proteins, the characteristic epidermal proteins of reptiles and birds. Journal of Experimental Zoology Part B: Molecular and Developmental Evolution, 330(8), pp.438–453.

Humphrey, P.S. and Livezey, B.C., 1982. Flightlessness in flying steamer-ducks. The Auk, 99(2), pp.368–372.

IUCN. 2022. The IUCN Red List of Threatened Species. Version 2022–2. https://www.iucnredlist.org

Jacob, J., 1978. Uropygial gland secretions and feather waxes. Chemical Zoology (AH Brush, ed.), 10, pp.165–211.

Jetz, W., Thomas, G.H., Joy, J.B., Hartmann, K. and Mooers, A.O., 2012. The global diversity of birds in space and time. Nature, 491(7424), pp.444–448.

Kelley, N.P. and Pyenson, N.D., 2015. Evolutionary innovation and ecology in marine tetrapods from the Triassic to the Anthropocene. Science, 348(6232).

Kruschke, J. K. 2011. Doing Bayesian data analysis. A tutorial with R and BUGS. Academic Press/Elsevier, Burlington, MA

Ksepka, D.T., Bertelli, S. and Giannini, N.P., 2006. The phylogeny of the living and fossil Sphenisciformes (penguins). Cladistics, 22(5), pp.412–441.

Ksepka, D.T., Balanoff, A.M., Smith, N.A., Bever, G.S., Bhullar, B.A.S., Bourdon, E., Braun, E.L., Burleigh, J.G., Clarke, J.A., Colbert, M.W. and Corfield, J.R., 2020. Tempo and pattern of avian brain size evolution. Current Biology, 30(11), pp.2026–2036.

Legendre, L.J. and Davesne, D., 2020. The evolution of mechanisms involved in vertebrate endothermy. Philosophical Transactions of the Royal Society B, 375(1793), p.20190136.

Lin, C.M., Jiang, T.X., Widelitz, R.B. and Chuong, C.M., 2006. Molecular signaling in feather morphogenesis. Current opinion in cell biology, 18(6), pp.730–741.

Livezey, B.C., 1989. Phylogenetic relationships and incipient flightlessness of the extinct Auckland Islands Merganser. The Wilson Bulletin, pp.410–435.

Livezey, B.C., 2003. *Evolution of flightlessness in rails (Gruiformes, Rallidae)*. American Ornithologists’ Union.

Lomolino, M.V., van der Geer, A.A., Lyras, G.A., Palombo, M.R., Sax, D.F. and Rozzi, R., 2013. Of mice and mammoths: generality and antiquity of the island rule. Journal of Biogeography, 40(8), pp.1427–1439.

Lowi-Merri, T.M., Benson, R.B., Claramunt, S. and Evans, D.C., 2021. The relationship between sternum variation and mode of locomotion in birds. BMC biology, 19(1), pp.1–23.

Lucas, A.M. and Stettenheim, P.R. 1972. Avian anatomy integument. Part 1. US Government Printing Office, Washington, DC.

Maderson, P.F. and Alibardi, L., 2000. The development of the sauropsid integument: a contribution to the problem of the origin and evolution of feathers. American Zoologist, 40(4), pp.513–529.

McFarland, D. and Budgell, P., 1970. The thermoregulatory role of feather movements in the barbary dove (Streptopelia risoria). Physiology & behavior, 5(7), pp.763–771.

McGowan, C., 1989. Feather structure in flightless birds and its bearing on the question of the origin of feathers. Journal of Zoology, 218(4), pp.537–547.

Miller, W.J., 1956. Silky plumage in the ring neck dove. Journal of Heredity, 47(1), pp.37–40.

Mitchell, J.S. and Makovicky, P.J., 2014. Low ecological disparity in Early Cretaceous birds. Proceedings of the Royal Society B: Biological Sciences, 281(1787), p.20140608.

Munafò, M.R. and Smith, G.D. 2018. Robust research needs many lines of evidence. Nature, 553:339–401.

Ng, C.S. and Li, W.H., 2018. Genetic and molecular basis of feather diversity in birds. Genome biology and evolution, 10(10), pp.2572–2586.

Novas, F.E., Agnolin, F., Brissón Egli, F. and Lo Coco, G.E., 2020. Pectoral girdle morphology in early-diverging paravians and living ratites: implications for the origin of flight.

Nudds, R.L., Dyke, G.J. and Rayner, J.M.V., 2004. Forelimb proportions and the evolutionary radiation of Neornithes. Proceedings of the Royal Society of London. Series B: Biological Sciences, 271(suppl_5), pp.S324-S327.

Palmer, R.S., 1972. Patterns of molting. Avian biology, 2, pp.65–101.

Prum, R.O., 1999. Development and evolutionary origin of feathers. Journal of Experimental Zoology, 285(4), pp.291–306.

Prum, R.O., 2005. Evolution of the morphological innovations of feathers. Journal of Experimental Zoology Part B: Molecular and Developmental Evolution, 304(6), pp.570–579.

Prum, R.O. and Brush, A.H., 2002. The evolutionary origin and diversification of feathers. The Quarterly review of biology, 77(3), pp.261–295.

Reddy, S., Kimball, R.T., Pandey, A., Hosner, P.A., Braun, M.J., Hackett, S.J., Han, K.L., Harshman, J., Huddleston, C.J., Kingston, S., Marks, B.D., Miglia, K.J., Moore, W.S., Sheldon, F.H., Witt, C.C., Yuri, T., and Braun, E.L., 2017. Why do phylogenomic data sets yield conflicting trees? Data type influences the avian tree of life more than taxon sampling. Systematic Biology, 66(5), pp.857–879.

Revell, L.J., 2012. phytools: an R package for phylogenetic comparative biology (and other things). Methods in ecology and evolution, (2), pp.217-223.

Rijke, A.M. and Jesser, W.A., 2011. The water penetration and repellency of feathers revisited. The Condor, 113(2), pp.245–254.

Roots, C., 2006. Flightless birds. Greenwood Publishing Group.

Roy, A., Pittman, M., Saitta, E.T., Kaye, T.G. and Xu, X., 2020. Recent advances in amniote palaeocolour reconstruction and a framework for future research. Biological Reviews, 95(1), pp.22–50.

Sawyer, R.H., Washington, L.D., Salvatore, B.A., Glenn, T.C. and Knapp, L.W., 2003. Origin of archosaurian integumentary appendages: The bristles of the wild turkey beard express feather-type β keratins. Journal of Experimental Zoology Part B: Molecular and Developmental Evolution, 297(1), pp.27–34.

Serrano, F.J., Costa-Pérez, M., Navalón, G. and Martín-Serra, A., 2020. Morphological Disparity of the Humerus in Modern Birds. Diversity, 12(5), p.173.

Taylor, S.S., Jamieson, I.G. and Wallis, G.P., 2007. Historic and contemporary levels of genetic variation in two New Zealand passerines with different histories of decline. Journal of Evolutionary Biology, 20(5), pp.2035–2047.

Tökölyi, J., Bokony, V. and Barta, Z., 2008. Seasonal colour change by moult or by the abrasion of feather tips: a comparative study. Biological Journal of the Linnean Society, 94(4), pp.711–721.

Torres, C.R. and Clarke, J.A., 2018. Nocturnal giants: evolution of the sensory ecology in elephant birds and other palaeognaths inferred from digital brain reconstructions. Proceedings of the Royal Society B, 285(1890), p.20181540.

von Meyer, H., 1861. *Archaeopteryx lithographica* (vogel-feder) und *Pterodactylus* von Solnhofen. Neues Jahrbuch für Mineralogie, Geognosie, Geologie und Petrefakten- Kunde, 1861, pp.678–679.

Watanabe, J., Field, D.J. and Matsuoka, H., 2021. Wing musculature reconstruction in extinct flightless auks (Pinguinus and Mancalla) reveals incomplete convergence with penguins (Spheniscidae) due to differing ancestral states. Integrative Organismal Biology, 3(1), p.obaa040.

Wood, M.J., Brock, G., Debray, J., Servio, P. and Kietzig, A.M., 2022. Robust Anti-Icing Surfaces Based on Dual Functionality─ Microstructurally-Induced Ice Shedding with Superimposed Nanostructurally-Enhanced Water Shedding. ACS Applied Materials & Interfaces, 14(41), pp.47310–47321.

Wright, N.A., Steadman, D.W. and Witt, C.C., 2016. Predictable evolution toward flightlessness in volant island birds. Proceedings of the National Academy of Sciences, 113(17), pp.4765–4770.

Yonezawa, T., Segawa, T., Mori, H., Campos, P.F., Hongoh, Y., Endo, H., Akiyoshi, A., Kohno, N., Nishida, S., Wu, J. and Jin, H., 2017. Phylogenomics and morphology of extinct paleognaths reveal the origin and evolution of the ratites. Current Biology, 27(1), pp.68–77.

