## Appendix for "RELAXING SELECTIVE PRESSURES ON DEVELOPMENTALLY COMPLEX INTEGUMENTARY STRUCTURES: FEATHER VANE SYMMETRY EVOLVES IN ADDITION TO BODY MASS AND WING LENGTH AFTER FLIGHT LOSS IN RECENT BIRDS"

### APPENDIX: SAITTA *ET AL.*

#### 1. Pairwise phylogenetic controls for the simple morphometric comparisons

| Flightless taxon | Closest volant relative(s) available for study |
|---|---|
| *Anas aucklandica* | *Anas chlorotis* |
| *Aptenodytes forsteri* | *Pelecanoides urinatrix*, *Oceanites oceanicus*, & *Diomedea immutabilis* |
| *Apteryx australis* | *Tinamus major saturatus* |
| *Aramidopsis plateni* | *Gallirallus striatus* |
| *Atlantisia rogersi* | *Porzana spiloptera* |
| *Casuarius unappendiculatus* | *Tinamus major saturatus* |
| *Dromaius novaehollandiae* | *Tinamus major saturatus* |
| *Dryolimnas cuvieri aldabranus* | *Dryolimnas cuvieri cuvieri* |
| *Eudyptes chrysocome crestatus* | *Pelecanoides urinatrix*, *Oceanites oceanicus*, & *Diomedea immutabilis* |
| *Fulica gigantea* | *Fulica armillata* |
| *Gallinula nesiotis* | *Gallinula chloropus chloropus* |
| *Gallirallus (=Tricholimnas) sylvestris* | *Ralls (=Gallirallus) philippensis philippensis* |
| *Gallirallus australis* | *Ralls (=Gallirallus) philippensis philippensis* |
| *Habroptila wallacii* | *Ralls (=Gallirallus) philippensis philippensis* |
| *Megacrex inepta* | *Himantornis haematopus* |
| *Mergus australis** | *Mergus merganser americanus* |
| *Nesoclopeus woodfordi* | *Ralls (=Gallirallus) philippensis philippensis* |
| *Phalacrocorax harrisi* | *Phalacrocorax auritus* |
| *Pinguinus impennis* | *Alca torda* |
| *Podiceps taczanowskii* | *Podiceps occipitalis occipitalis* |
| *Podilymbus gigas* | *Podilymbus podiceps* |
| *Porzana palmeri* | *Porzana pusilla pusilla* |
| *Porzana sandwichensis* | *Porzana tabuensis tabuensis* |
| *Rhea americana intermedia* | *Tinamus major saturatus* |
| *Rollandia microptera* | *Rollandia rolland chilensis* |
| *Spheniscus demersus* | *Pelecanoides urinatrix*, *Oceanites oceanicus*, & *Diomedea immutabilis* |
| *Strigops habroptilus* | *Nestor notabilis* |
| *Struthio camelus* | *Tinamus major saturatus* |
| *Tachyeres pteneres* | *Tachyeres patachonicus* |
| *Tribonyx mortierii* | *Tribonyx ventralis whitei* |
| *Xenicus lyalli* | *Xenicus longipes* |

Table S1. Pairwise phylogenetic comparisons for simple statistical analyses of morphometric shifts from volant to flightless or poor flighted taxa as sister taxon comparisons. Values of the three tubesnoses were averaged together. * indicates poor flying/'incipiently flightless'.

#### 2. *Apteryx australis* semi-alternating barb development pattern

##### 2.1. *Apteryx australis* primary

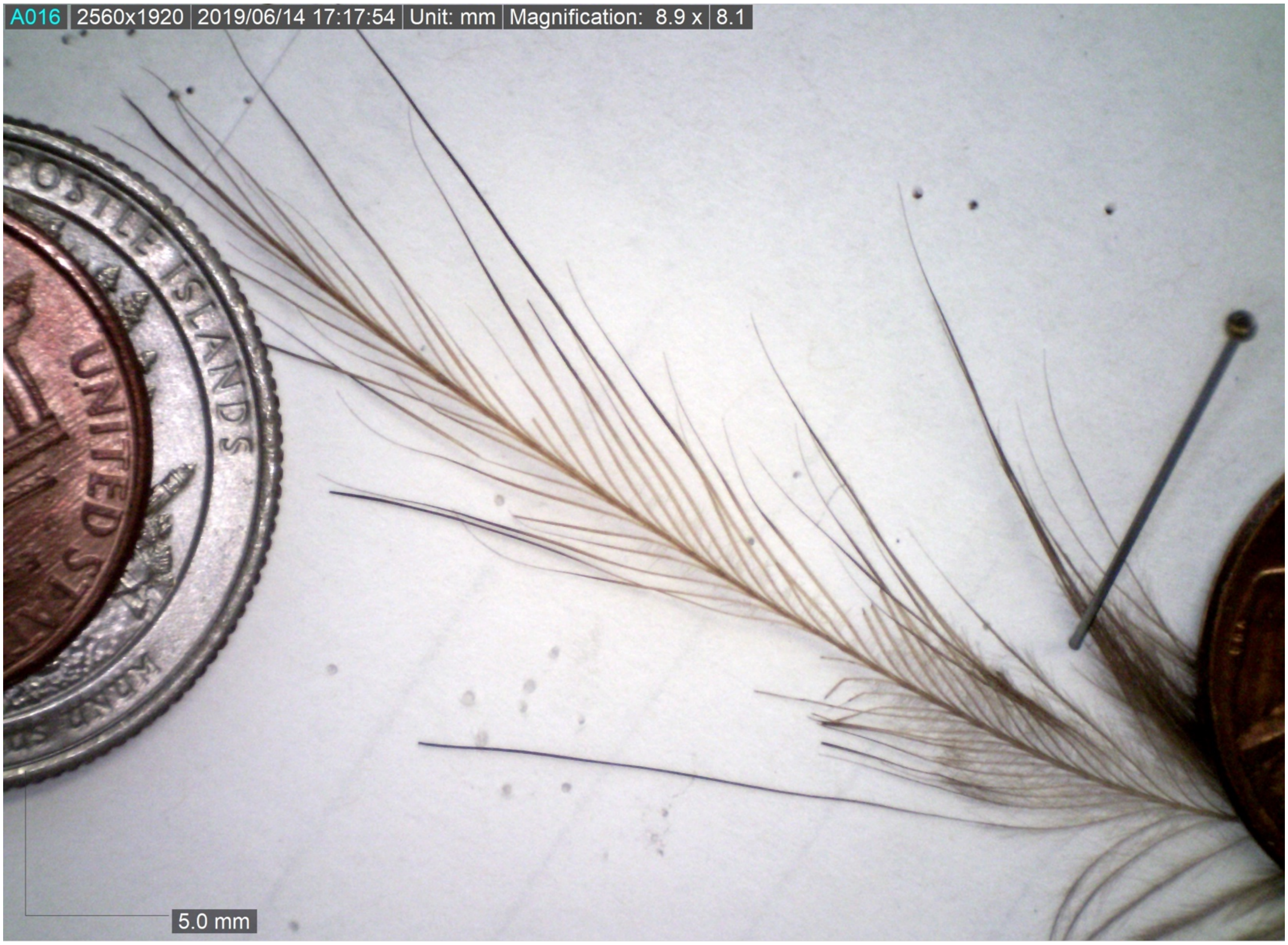


Figure S1. *Apteryx australis* primary remex showing semi-alternating barb development pattern.

### 2.2. *Apteryx australis* tertial

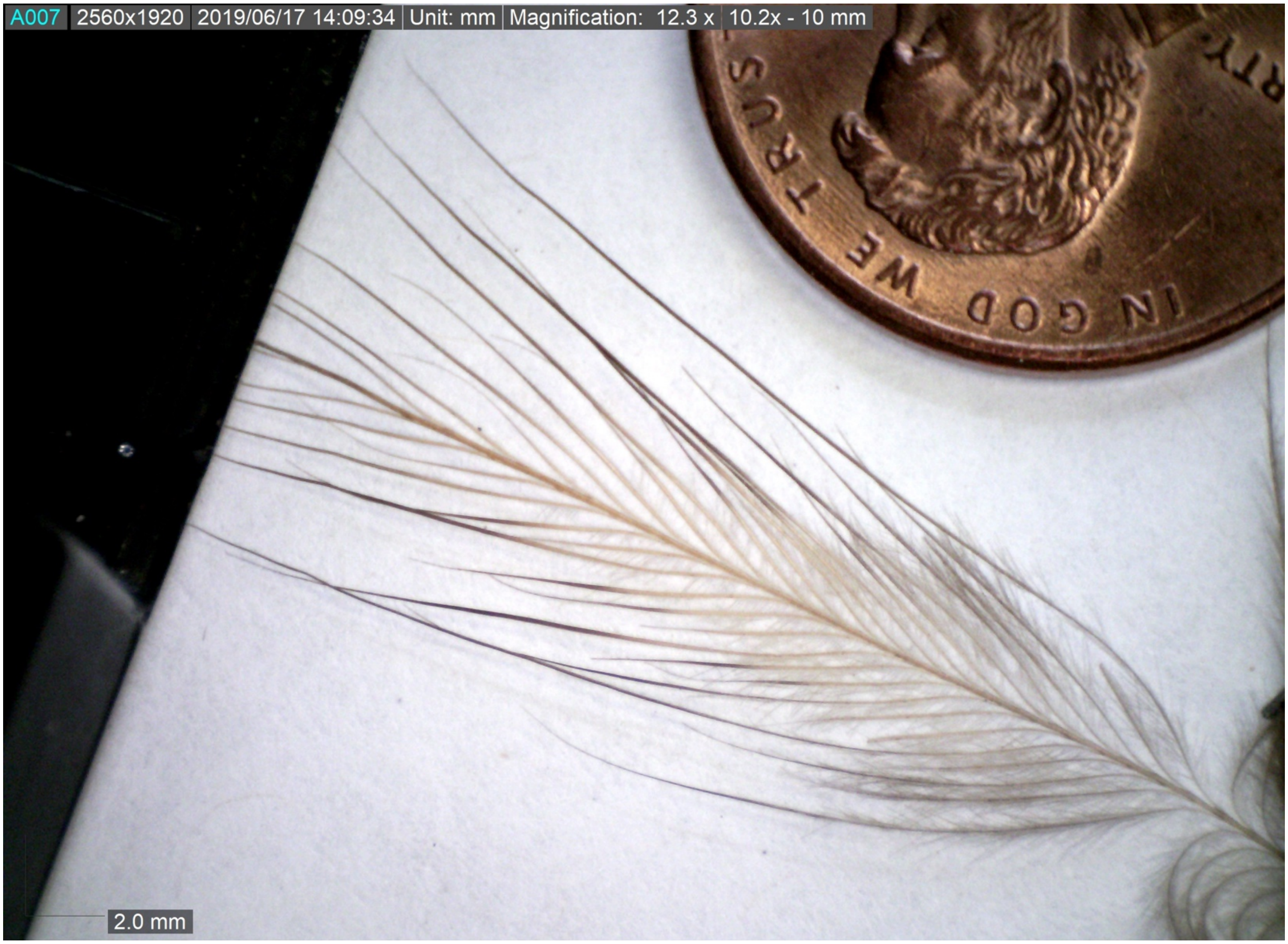

Figure S2. *Apteryx australis* tertial showing semi-alternating barb development pattern.

### 2.3. *Apteryx australis* rectrix

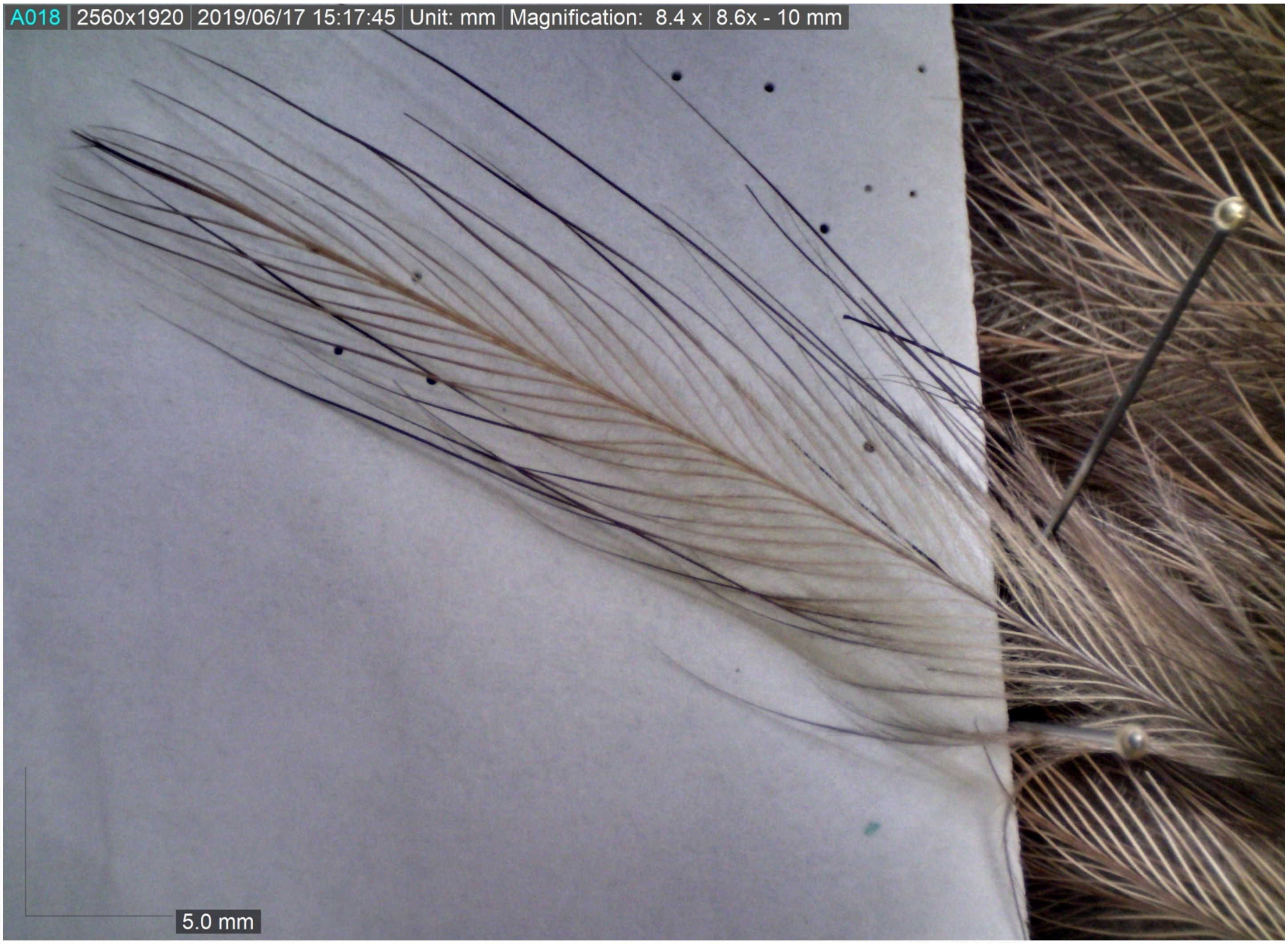


Figure S3. *Apteryx australis* rectrix showing semi-alternating barb development pattern.

### 2.4. *Apteryx australis* dorsal contour

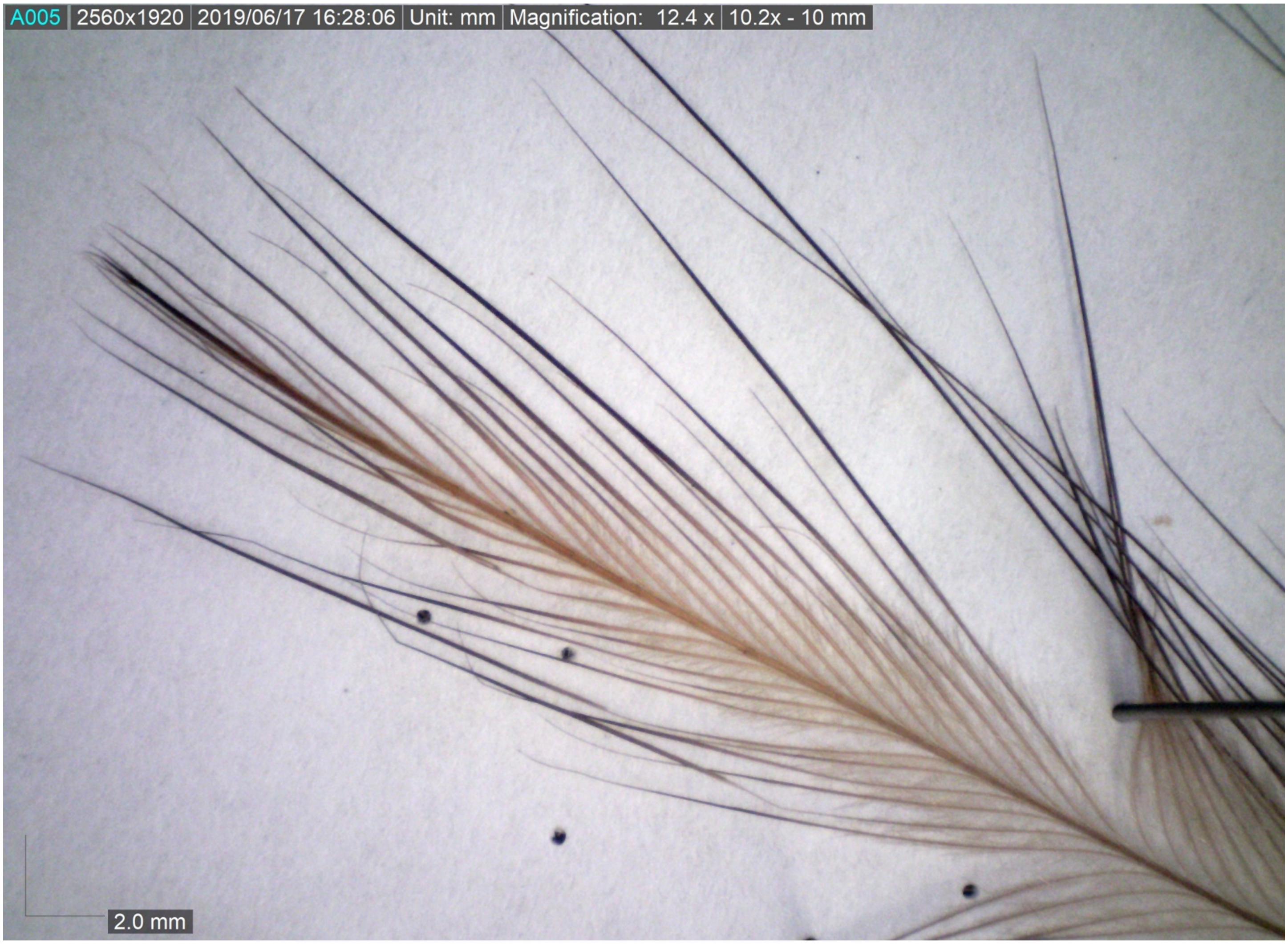


Figure S4. *Apteryx australis* dorsal contour showing semi-alternating barb development pattern.

### 3. Considerations of data acquisition

We attempted to account for challenges arising from the use of museum bird skins, which were often collected many years ago (especially true for extinct taxa) and therefore worn and degraded to some degree, as well as sewn together such that wings are almost always positioned tightly folded up against the body and rectrices are folded medially (hence why we examined the longest primary remex, the dorsal-most tertial remex, and the dorsal-most rectrix). This leads to problems of feather manipulation while obtaining data, especially when observing microscopic structures, and difficulty in controlling for homology between and within feathers.

As for homology, feather development occurring from the apex downward as the feather extends from the follicle (Ng & Li 2018) might suggest that homologous barbs are best described as those equally distant from the apex in terms of the number of barbs. However, differences in barb angle and density along the rachis can vary between the leading and trailing vanes, complicating this approach (e.g., the 10$^{th}$ barb from the apex on the leading vane with lower barb angle/density might be much further down the feather than the 10$^{th}$ barb from the apex on the trailing vane with higher barb angle/density). Furthermore, different feathers will not have the same length or total number of barbs (e.g., the 40$^{th}$ barb on remex of one species might not be in the same functional region of the feather as the 40$^{th}$ barb on the remex of another species). Additionally, counting barbs in such a manner is extremely time intensive, so an alternative view of homology might examine the entire feather from apex to calamus and describe the extent down the rachis as homologous (e.g., the barbs midway down the rachis are homologous between feathers). However, without plucking the feathers from rare, strictly conserved museum skin specimens, this cannot be done with precision since the feather bases are inaccessible on the skins. These considerations of homology at the microscopic level do not even cover considerations between follicles across the body/plumage (e.g., differences in the number of primary remiges between species). Therefore, we chose to focus on the middle position along an exposed feather as a rough approximation of what might be considered a comparable intra-feather region/possible module across taxa, even if not precisely developmentally homologous.

As fractal structures, the largest and smallest order branches of feather filaments range from the macro- to microscopic, leading one to carefully consider measurement tools, especially the use of scaled microscopic images that are at risk of parallax errors when feathers are observed on three dimensional skins with extreme topography and challenges positioning the specimen into the perpendicular focal plane of the microscope. All these factors lead to noise in the data that we attempted to minimize through the use of the maneuverable microscope mounted stand, allowing for three dimensional skins to be observed with minimal parallax error, as well as our considerations of the subsequent statistical analyses (described in main text). Furthermore, different authors independently measured the same subset of the feathers in the dataset, finding reasonably consistent measurement values between researchers.

### 4. Repeated PCA under different initial conditions

| | Only clades with flightless taxa | | All clades | |
|---|---|---|---|---|
| | Long branches | No long branches | Long branches | No long branches |
| Whole plumage | 84.17 | 85.98 | **84.69** | 86.53 |
| Primaries | **84.26** | 77.91 | 84.41 | 77.33 |
| Tertials | **86.44** | 86.62 | | |
| Rectrices | **81.27** | 85.2 | | |
| Dorsal contours | **78.75** | 79.47 | | |
| Ventral contours | **82.37** | 84.37 | | |

Table S2. Total proportion of the variation in the data explained by both PC1 and PC2. Bold values are those associated with PCA presented in the figures of the main text.

### 4.1. Whole plumage

#### 4.1.1. Whole plumage: long branches, PCA of only clades with flightless taxa

PC1 and PC2 explain ~63.10% and ~21.07% of the variation, respectively.

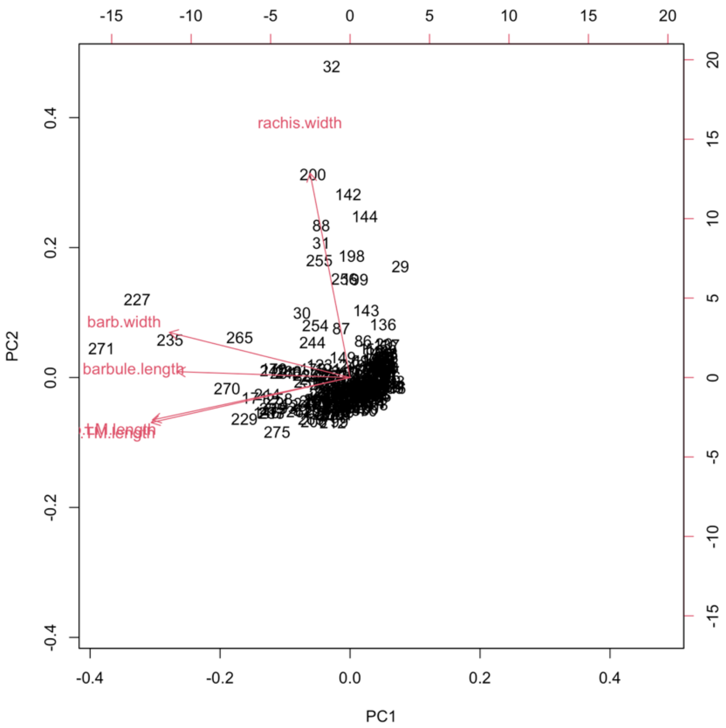


Figure S5. PCA biplot of feathers from the whole plumage of only clades containing flightless taxa, with penguins/tubenoses and ratites/tinamous included. Red arrows indicate variable loadings. Leading and trailing barb lengths show similar loadings. N = 280 feathers, 56 taxa.

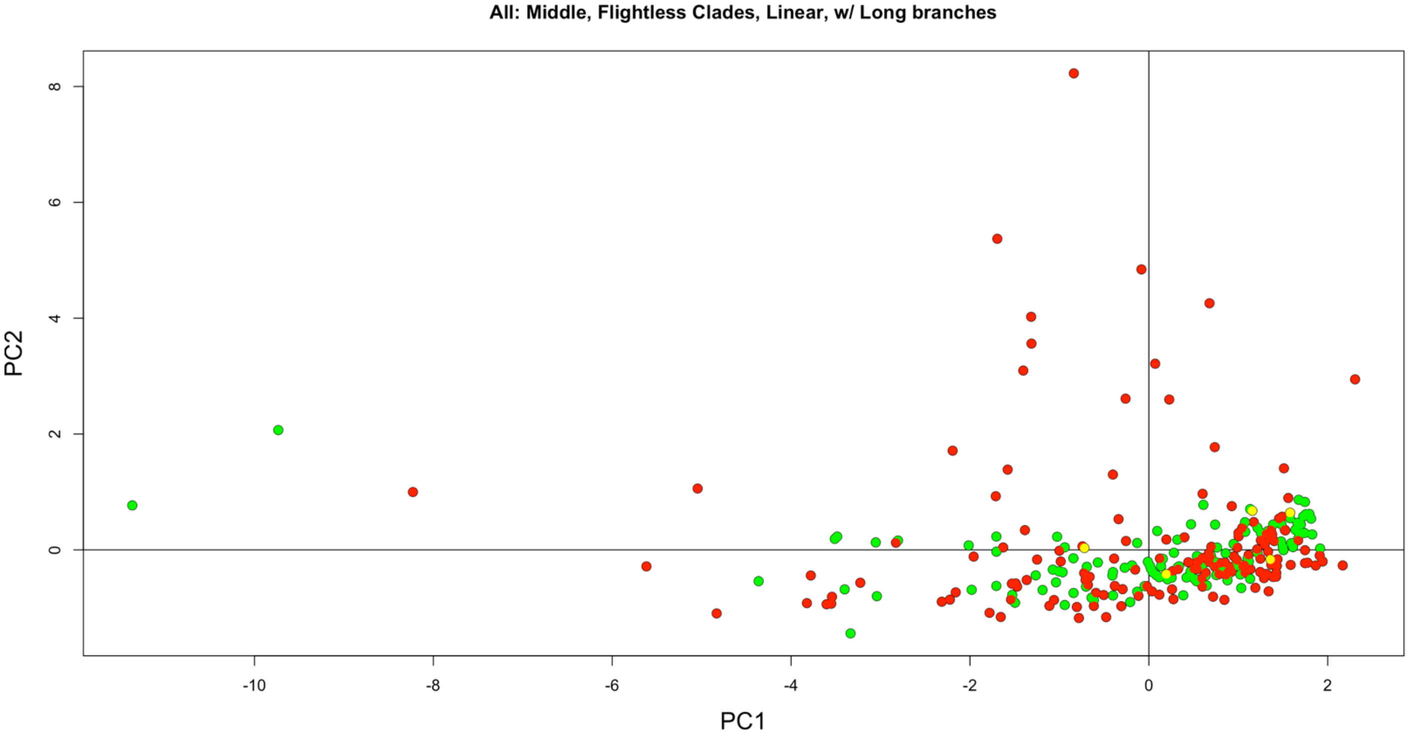

Figure S6. PCA of feathers from the whole plumage of only clades containing flightless taxa, with penguins/tubenoses and ratites/tinamous included. Green are volant taxa, red are flightless taxa, and yellow are poor flying/'incipiently flightless' taxa. N = 280 feathers, 56 taxa.

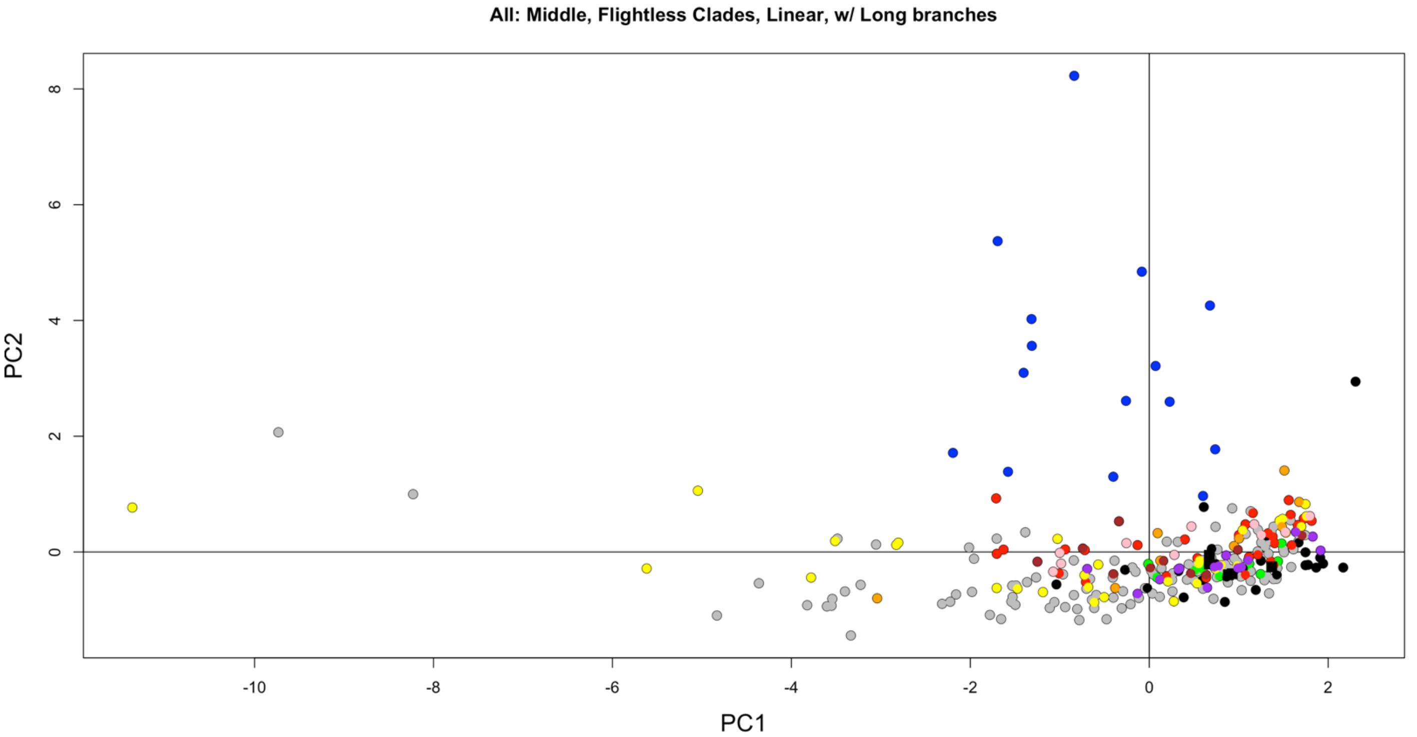


Figure S7. PCA of feathers from the whole plumage of only clades containing flightless taxa, with penguins/tubenoses and ratites/tinamous included. Color and shape indicate taxonomy as in Figure 2B. N = 280 feathers, 56 taxa.

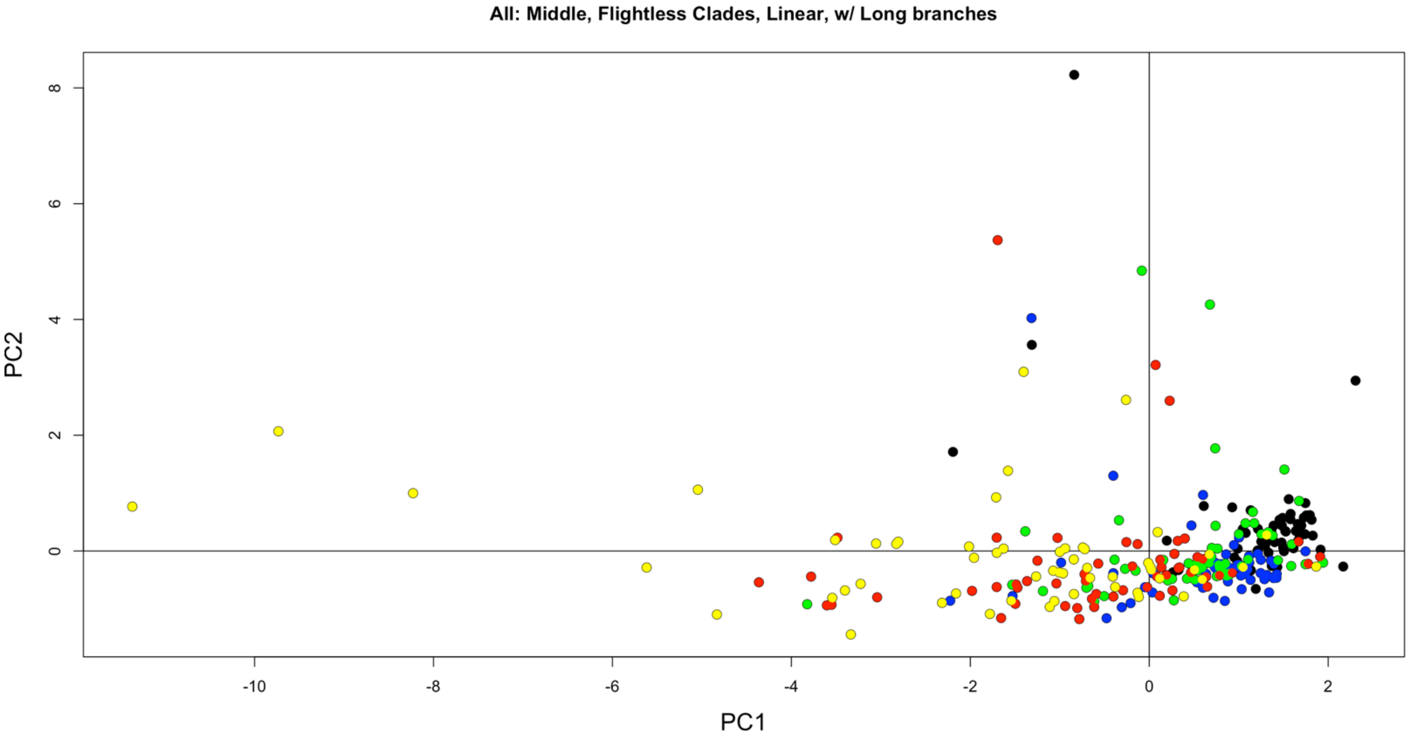


Figure S8. PCA of feathers from the whole plumage of only clades containing flightless taxa, with penguins/tubenoses and ratites/tinamous included. Black are primaries, blue are tertials, green are rectrices, red are dorsal contours, and yellow are ventral contours. N = 280 feathers, 56 taxa.

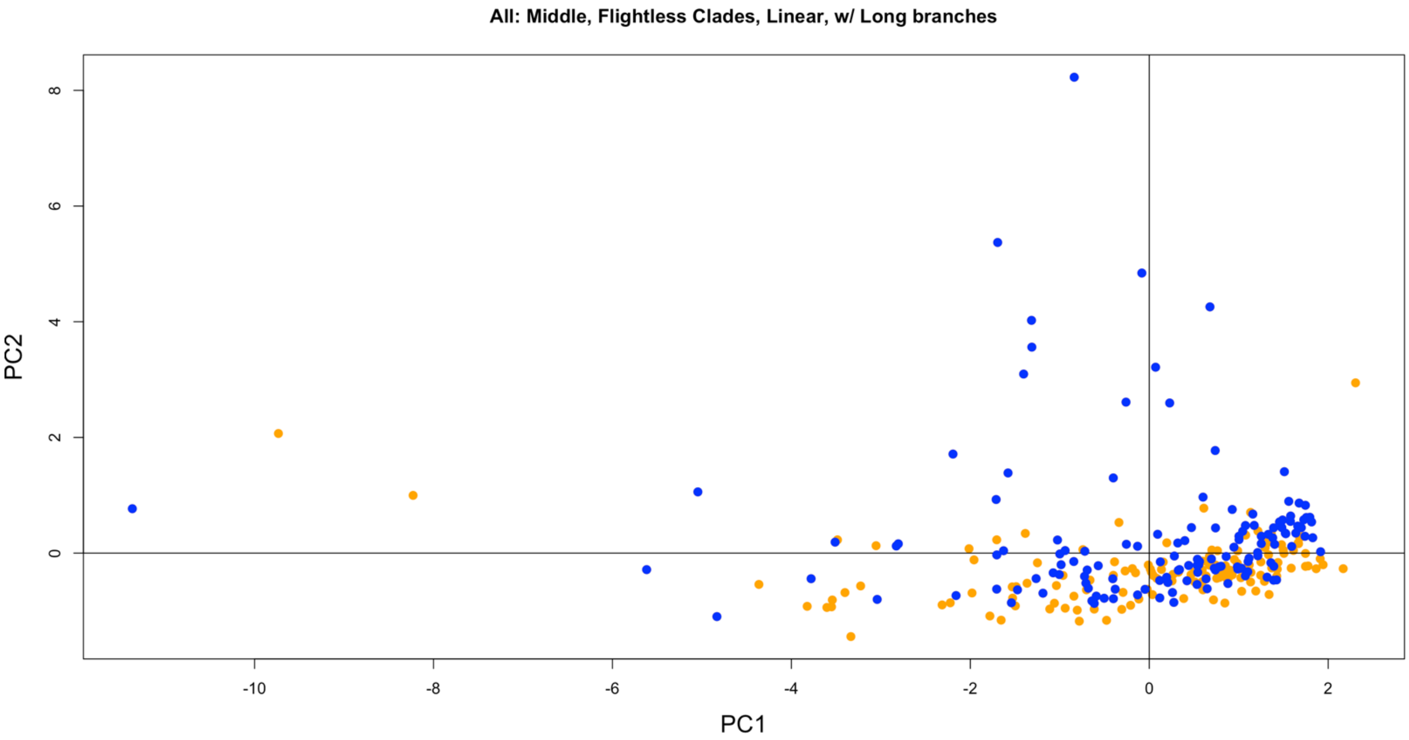


Figure S9. PCA of feathers from the whole plumage of only clades containing flightless taxa, with penguins/tubenoses and ratites/tinamous included. Blue are semiaquatic, and orange are terrestrial. N = 280 feathers, 56 taxa.

#### 4.1.2. Whole plumage: long branches, PCA of all clades (see Fig. 2)

PC1 and PC2 explain ~63.63% and ~21.06% of the variation, respectively.

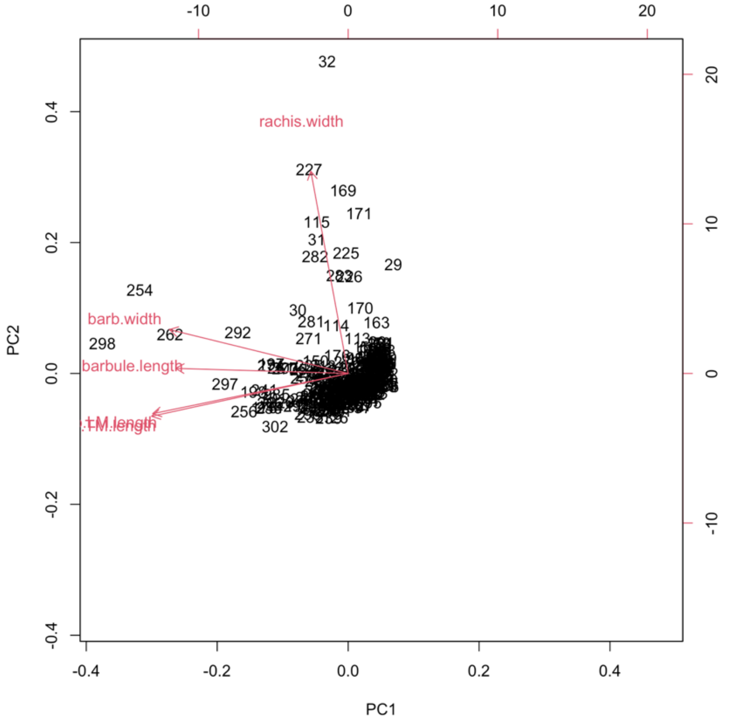


Figure S10. PCA biplot of feathers from the whole plumage of all clades, with penguins/tubenoses and ratites/tinamous included. Red arrows indicate variable loadings. Leading and trailing barb lengths show similar loadings. N = 307 feathers, 83 taxa.

#### 4.1.3. Whole plumage: no long branches (i.e., without Palaeognathae, Sphenisciformes, and Procellariiformes), PCA of only clades with flightless taxa

PC1 and PC2 explain ~66.53% and ~19.45% of the variation, respectively.

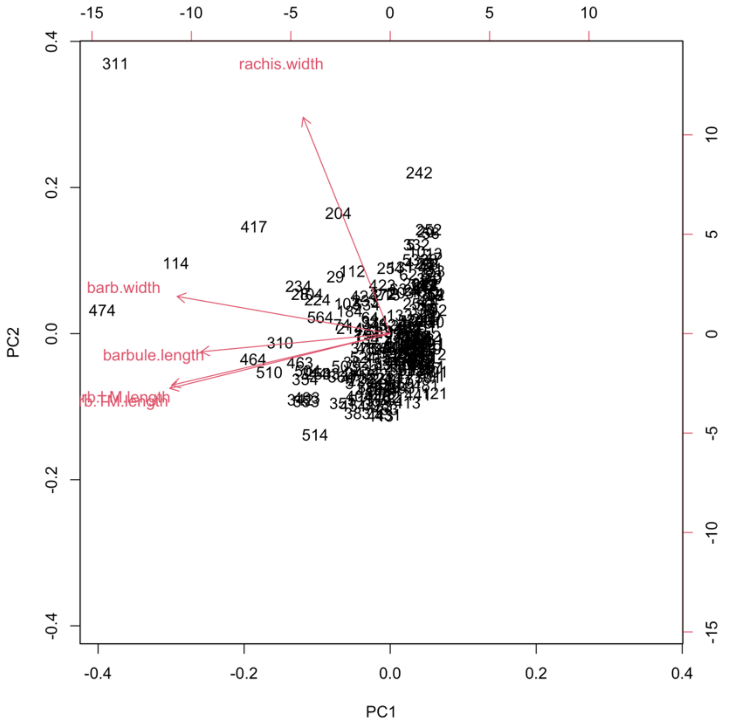


Figure S11. PCA biplot of feathers from the whole plumage of only clades containing flightless taxa, except penguins/tubenoses and ratites/tinamous. Red arrows indicate variable loadings. Leading and trailing barb lengths show similar loadings. N = 220 feathers, 44 taxa.

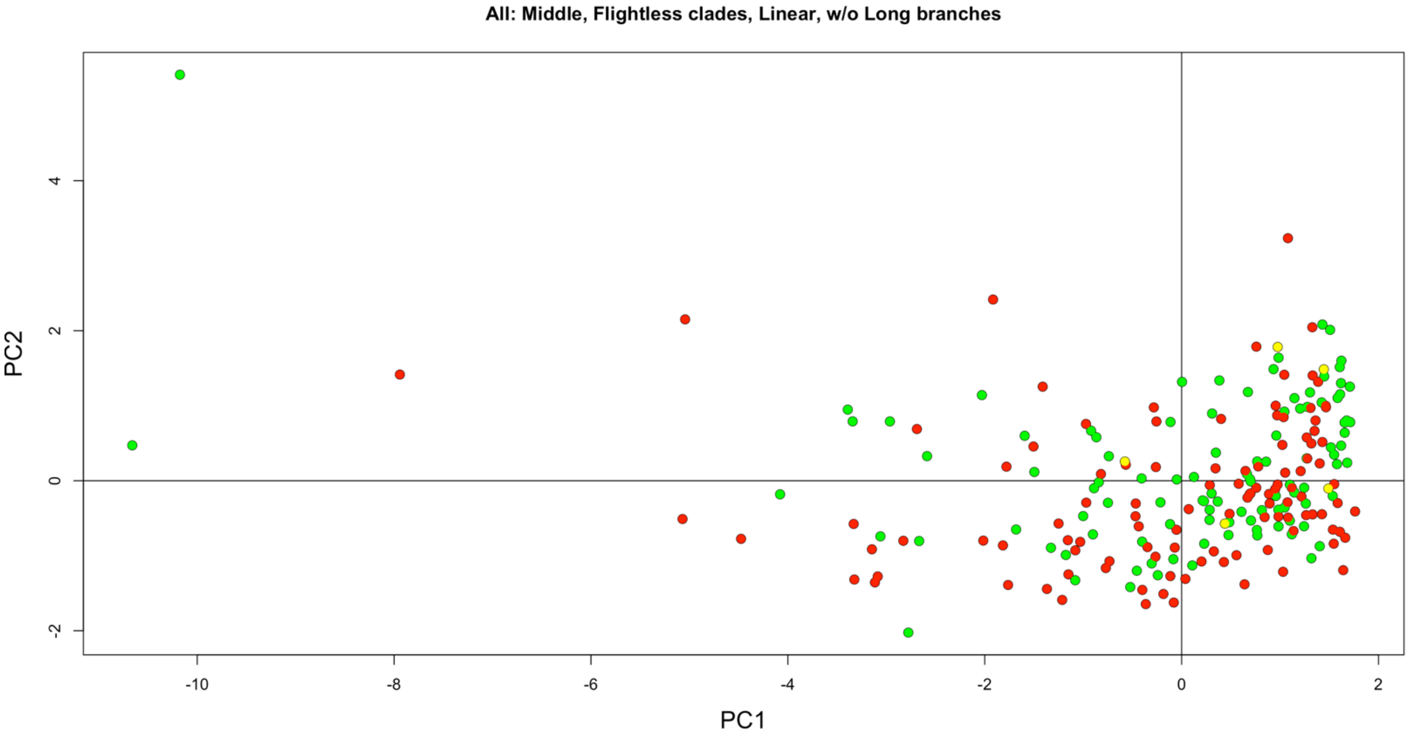

Figure S12. PCA of feathers from the whole plumage of only clades containing flightless taxa, except penguins/tubenoses and ratites/tinamous. Green are volant taxa, red are flightless taxa, and yellow are poor flying/‘incipiently flightless’ taxa. N = 220 feathers, 44 taxa.

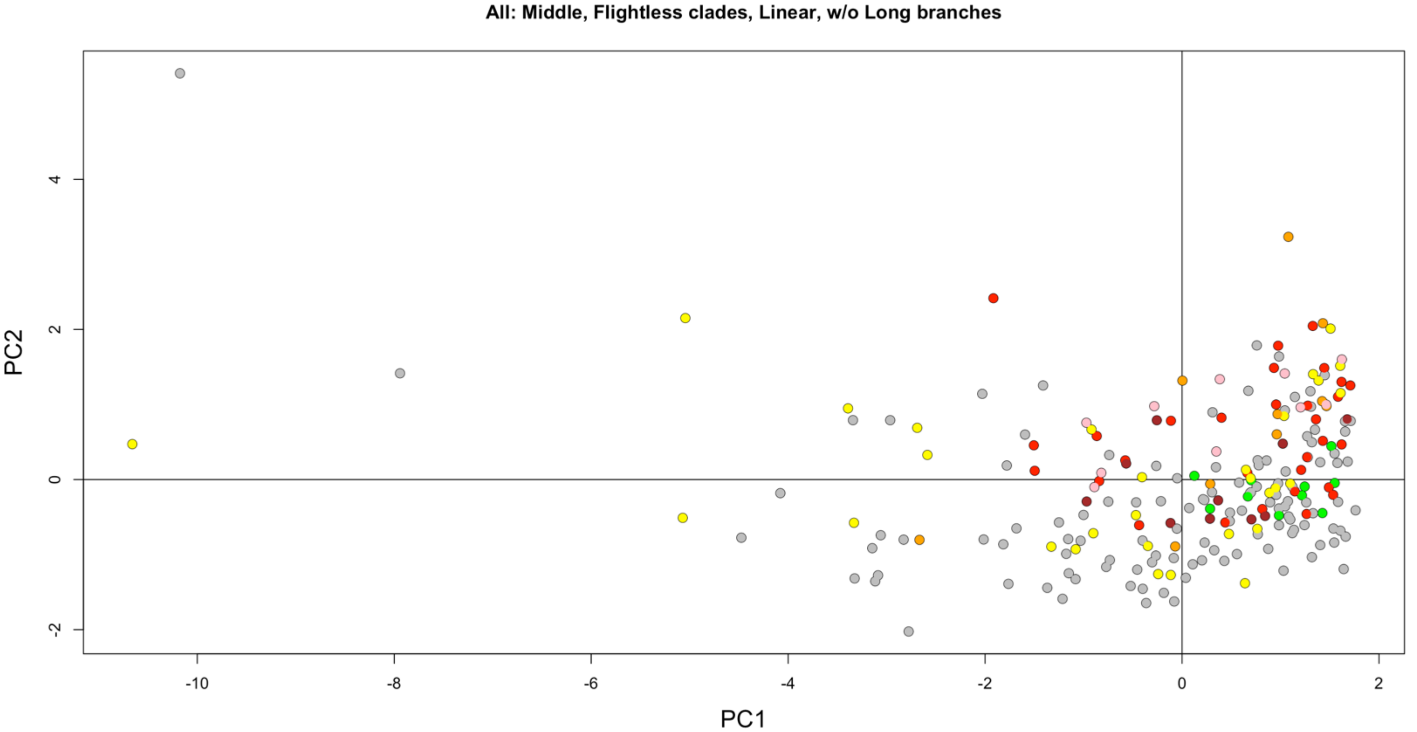


Figure S13. PCA of feathers from the whole plumage of only clades containing flightless taxa, except penguins/tubenoses and ratites/tinamous. Color indicates clade as in Figure 2B. N = 220 feathers, 44 taxa.

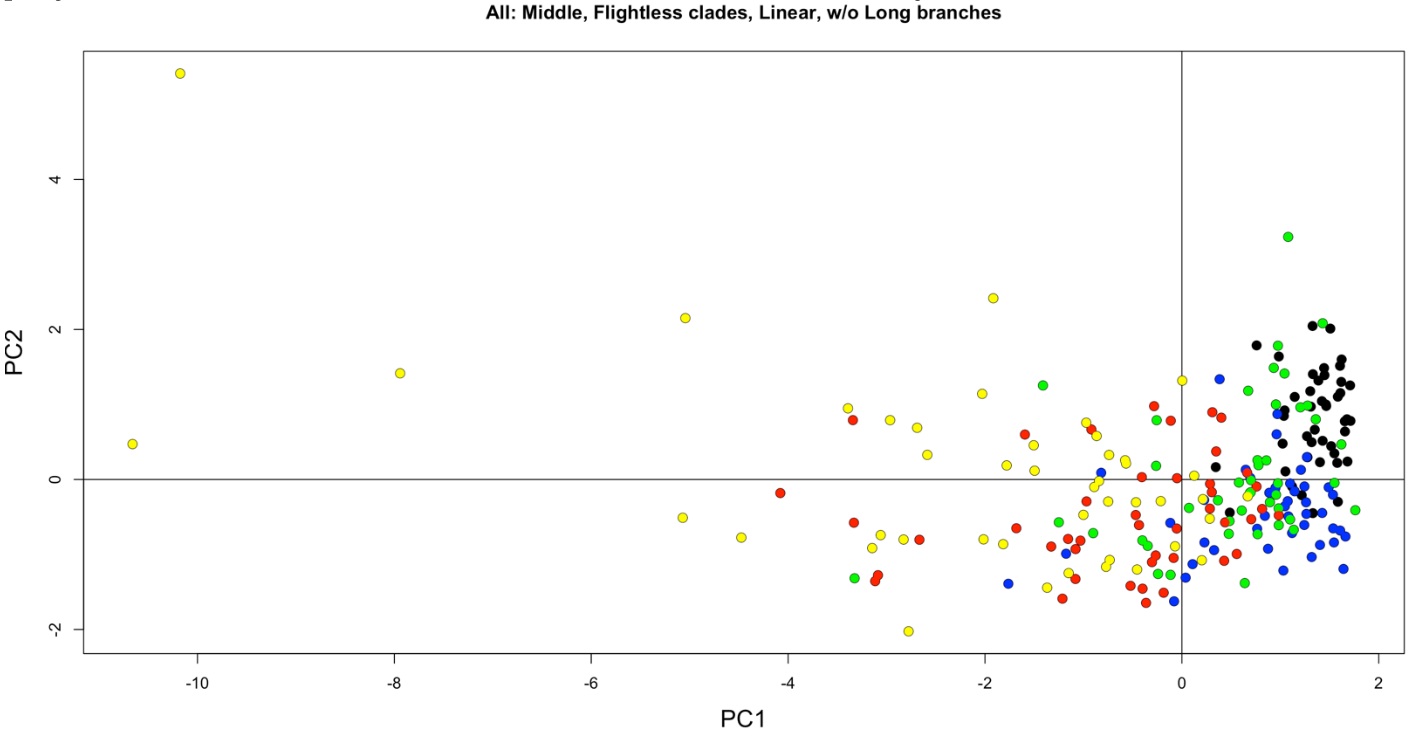


Figure S14. PCA of feathers from the whole plumage of only clades containing flightless taxa, except penguins/tubenoses and ratites/tinamous. Black are primaries, blue are tertials, green are rectrices, red are dorsal contours, and yellow are ventral contours. N = 220 feathers, 44 taxa.

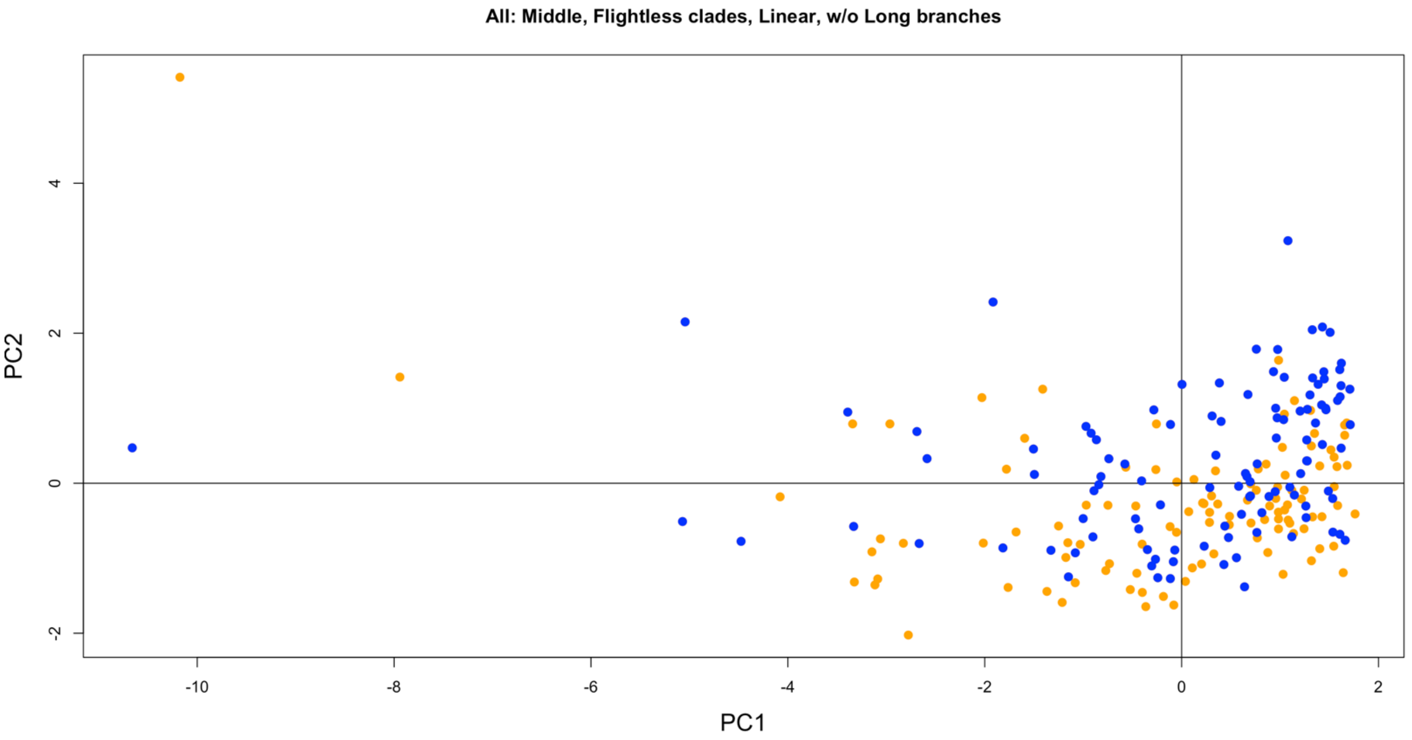


Figure S15. PCA of feathers from the whole plumage of only clades containing flightless taxa, except penguins/tubenoses and ratites/tinamous. Blue are semiaquatic, and orange are terrestrial. N = 220 feathers, 44 taxa.

#### 4.1.4. Whole plumage: no long branches (i.e., without Palaeognathae, Sphenisciformes, and Procellariiformes), PCA of all clades

PC1 and PC2 explain ~65.96% and ~20.57% of the variation, respectively.

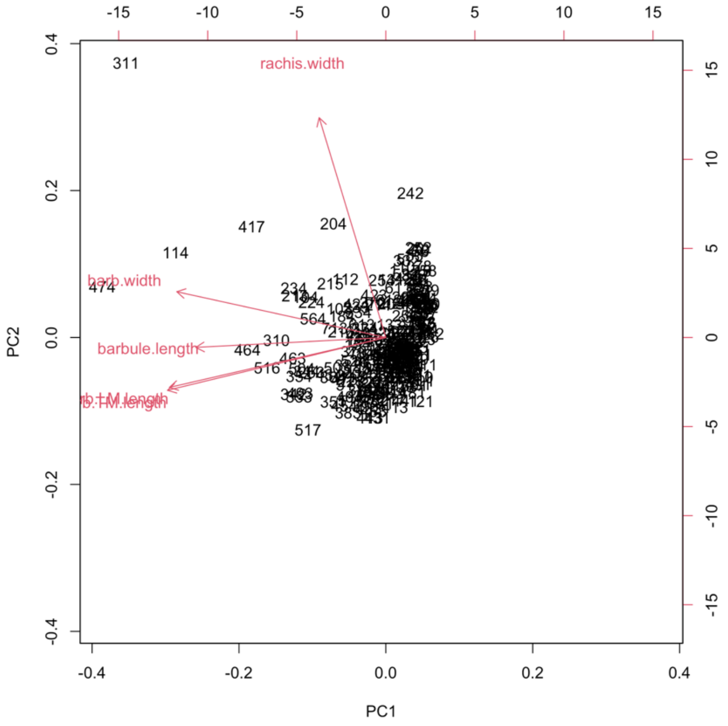

Figure S16. PCA biplot of feathers from the whole plumage of all clades, except penguins/tubenoses and ratites/tinamous. Red arrows indicate variable loadings. Leading and trailing barb lengths show similar loadings. N = 267 feathers, 75 taxa.

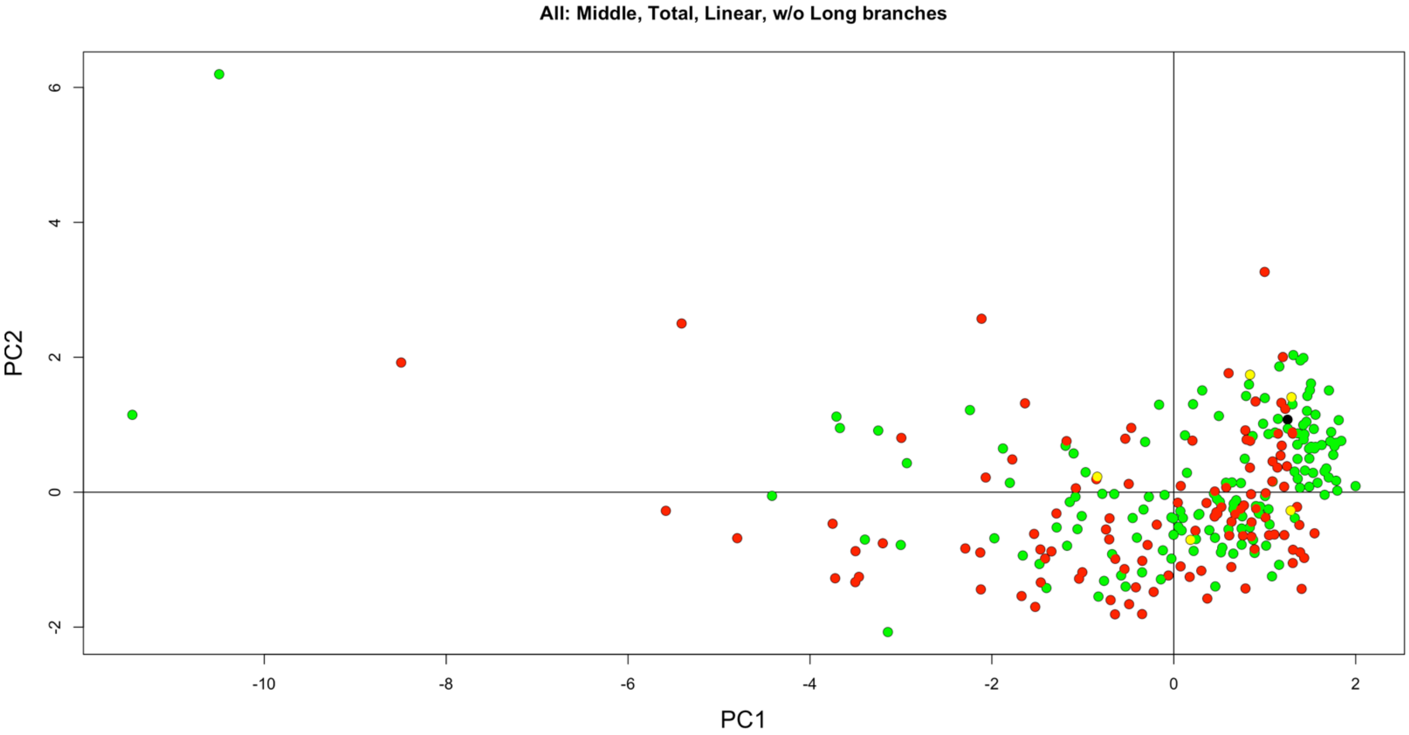


Figure S17. PCA biplot of feathers from the whole plumage of all clades, except penguins/tubenoses and ratites/tinamous. Green are volant taxa, red are flightless taxa, and yellow are poor flying/'incipiently flightless' taxa. N = 267 feathers, 75 taxa.

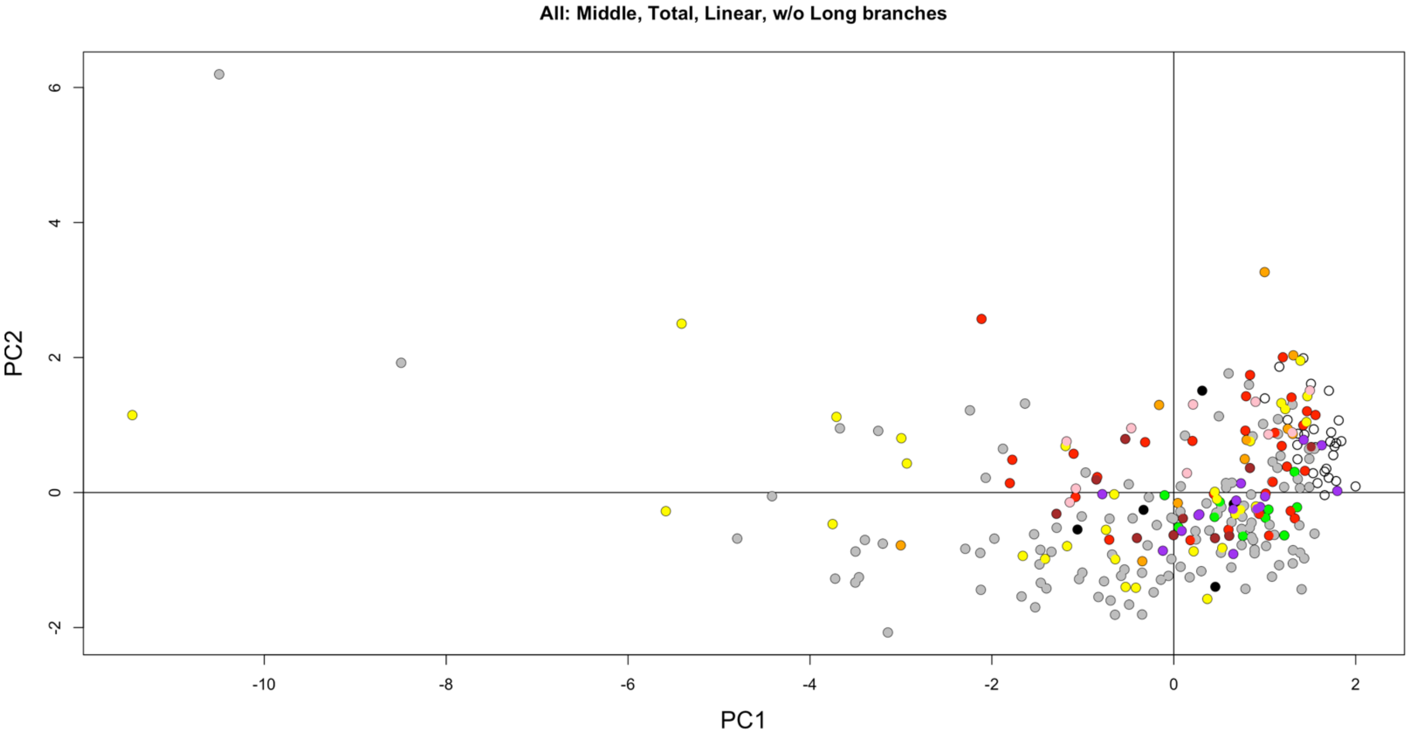


Figure S18. PCA biplot of feathers from the whole plumage of all clades, except penguins/tubenoses and ratites/tinamous. Color indicates clade as in Figure 2B. N = 267 feathers, 75 taxa.

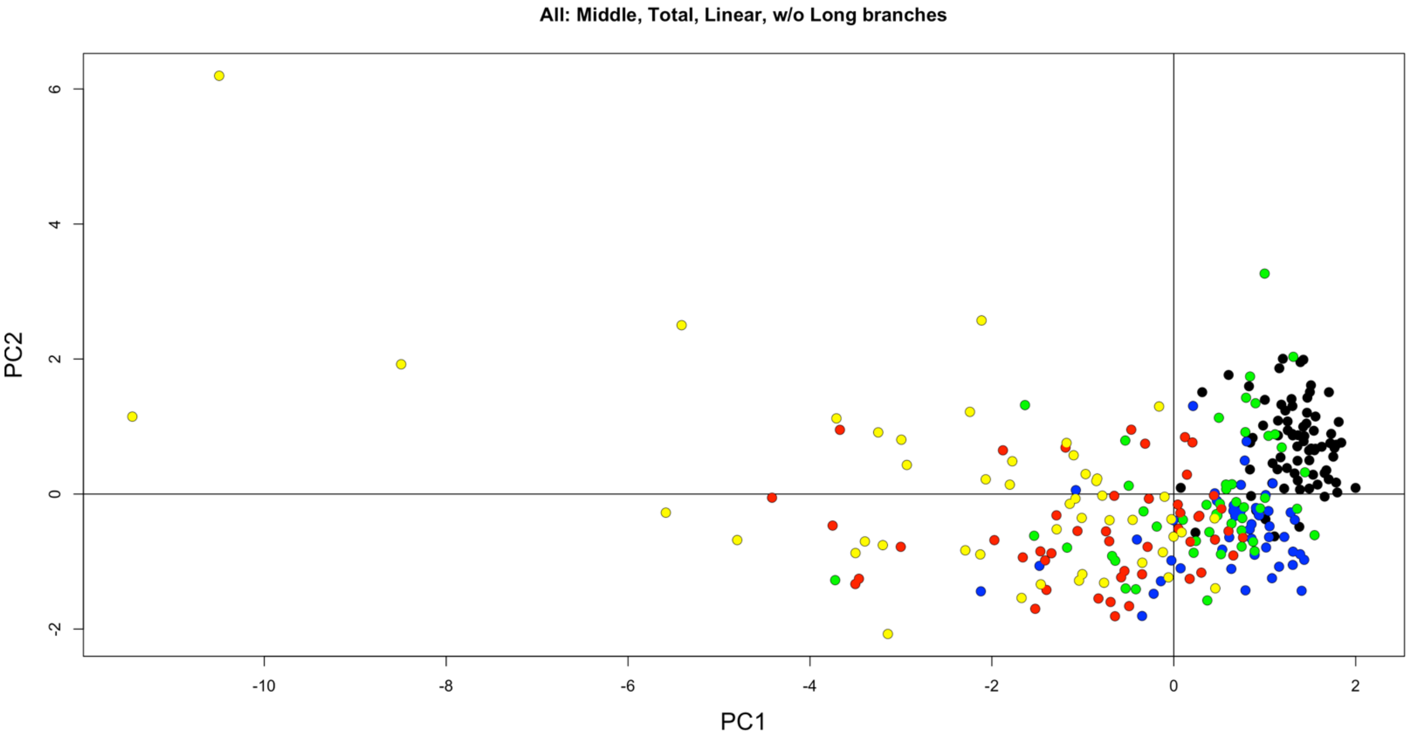

Figure S19. PCA biplot of feathers from the whole plumage of all clades, except penguins/tubenoses and ratites/tinamous. Black are primaries, blue are tertials, green are rectrices, red are dorsal contours, and yellow are ventral contours. N = 267 feathers, 75 taxa.

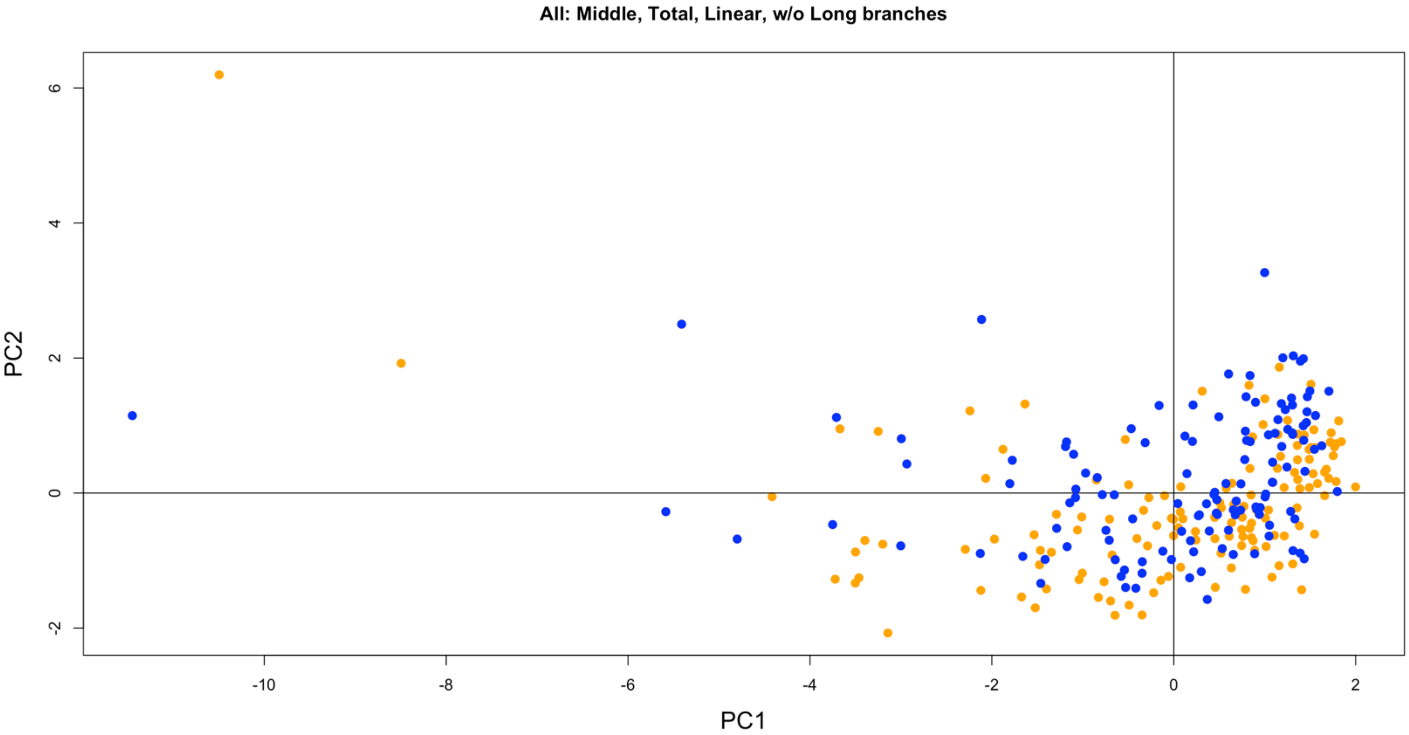

Figure S20. PCA biplot of feathers from the whole plumage of all clades, except penguins/tubenoses and ratites/tinamous. Blue are semiaquatic, and orange are terrestrial. N = 267 feathers, 75 taxa.

#### 4.2. Primaries

As a result of not log-transforming the data, bear in mind that for the shifts in metrics from a volant ancestor (i.e., the origin on the plots of ΔPC1 and ΔPC2), extreme values are not consistently scaled after dividing the metric by the rachis length at that middle barb pair. The same holds true for the feather types other than primaries (sections 3.3–3.7).

#### 4.2.1. Primary: long branches, PC space established by only clades with flightless taxa

PC1 and PC2 explain ~64.55% and ~19.71% of the variation, respectively.

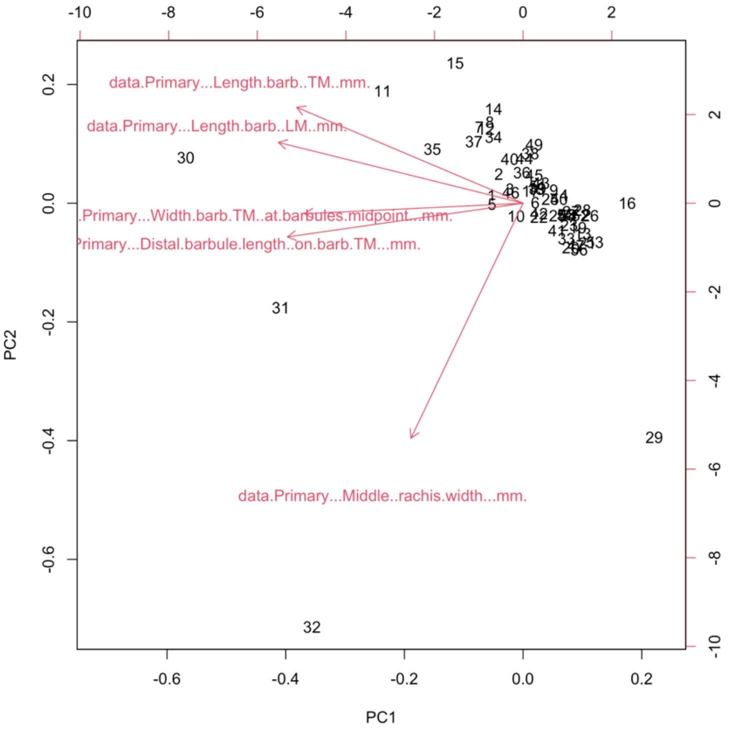


Figure S21. PCA biplot of primaries of only clades containing flightless taxa, with penguins/tubenoses and ratites/tinamous included. Red arrows indicate variable loadings. N = 56 taxa.

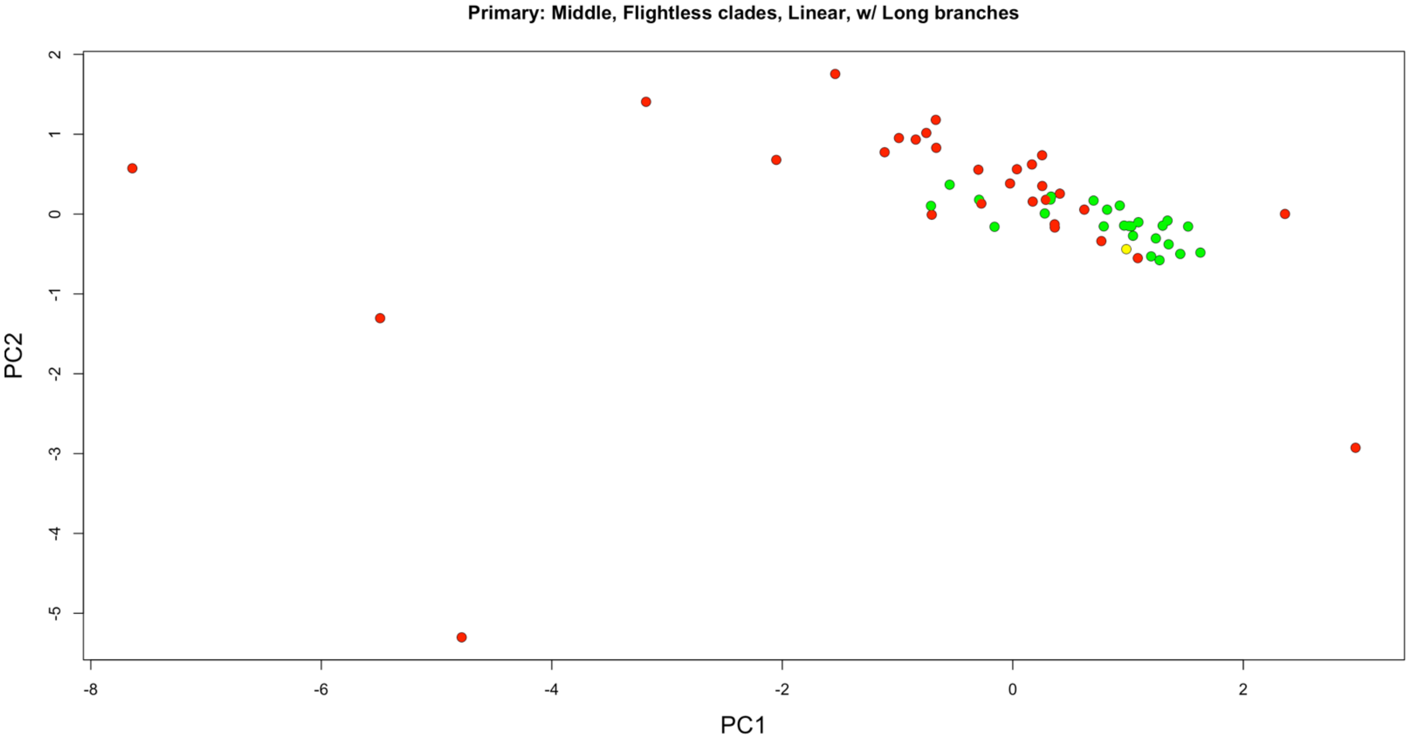

Figure S22. PCA of primaries of only clades containing flightless taxa, with penguins/tubenoses and ratites/tinamous included. Green are volant taxa, red are flightless taxa, and yellow are poor flying/'incipiently flightless' taxa. N = 56 taxa.

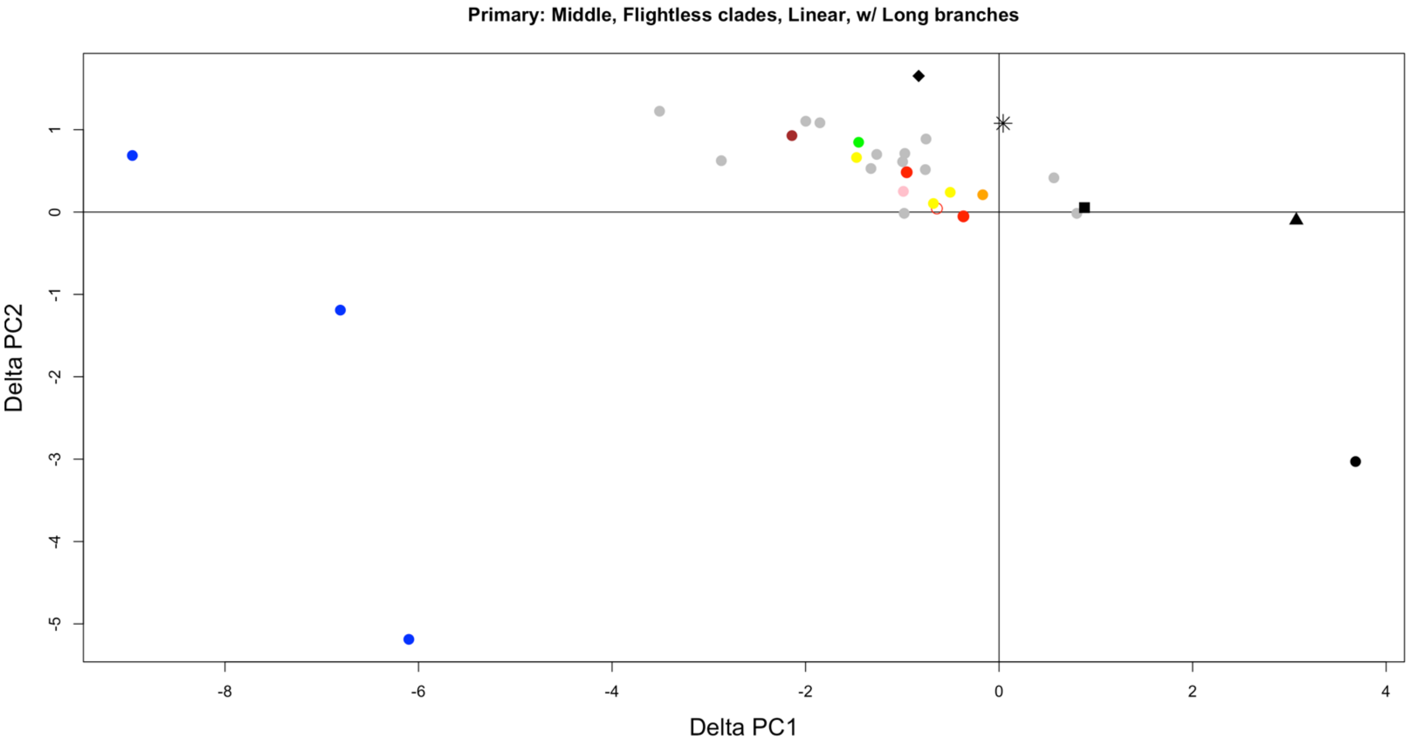


Figure S23. Simple pairwise phylogenetic PCA shift of primaries during flight loss based on an initial morphospace of only clades containing flightless taxa, with penguins/tubenoses and ratites/tinamous included. Color and shape indicate taxonomy as in Figure 3. Open circle is poor flyer/'incipiently flightless'. N = 31 taxa.

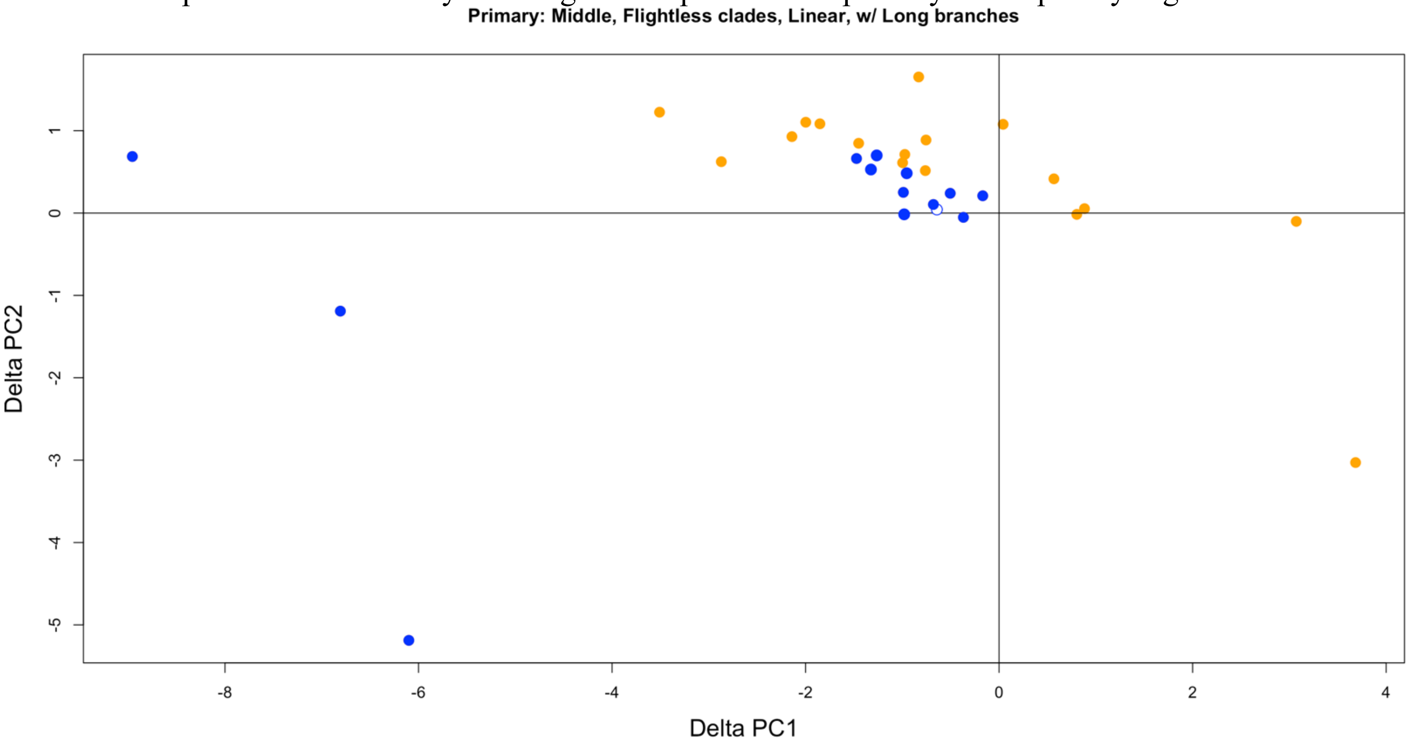


Figure S24. Simple pairwise phylogenetic PCA shift of primaries during flight loss based on an initial morphospace of only clades containing flightless taxa, with penguins/tubenoses and ratites/tinamous included. Blue are semiaquatic, and orange are terrestrial. Open circle is poor flyer/'incipiently flightless'. N = 31 taxa.

#### 4.2.2. Primary: long branches, PC space established by all clades

PC1 and PC2 explain ~65.30% and ~19.11% of the variation, respectively.

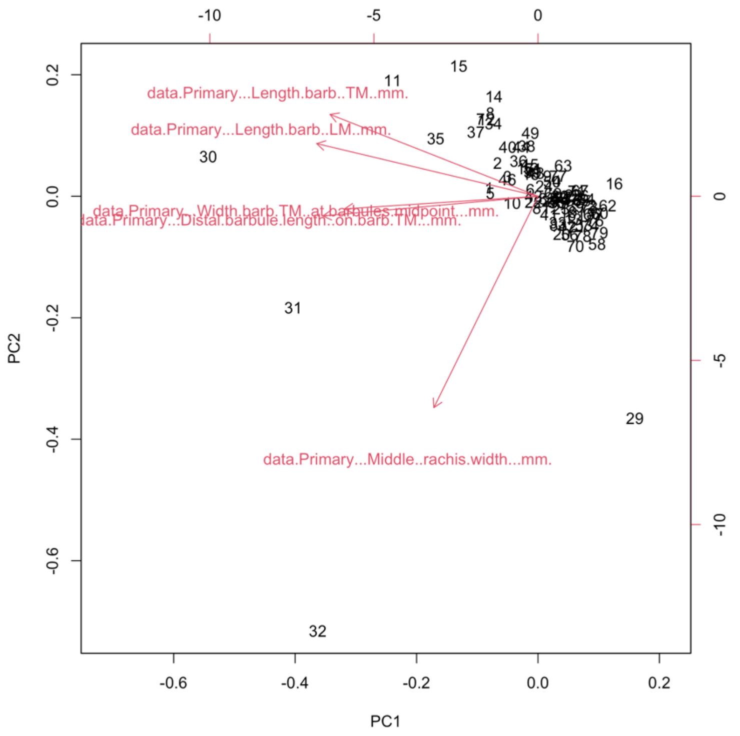


Figure S25. PCA biplot of primaries of all clades, with penguins/tubenoses and ratites/tinamous included. Red arrows indicate variable loadings. Barb width and barbule length show similar loadings. N = 83 taxa.

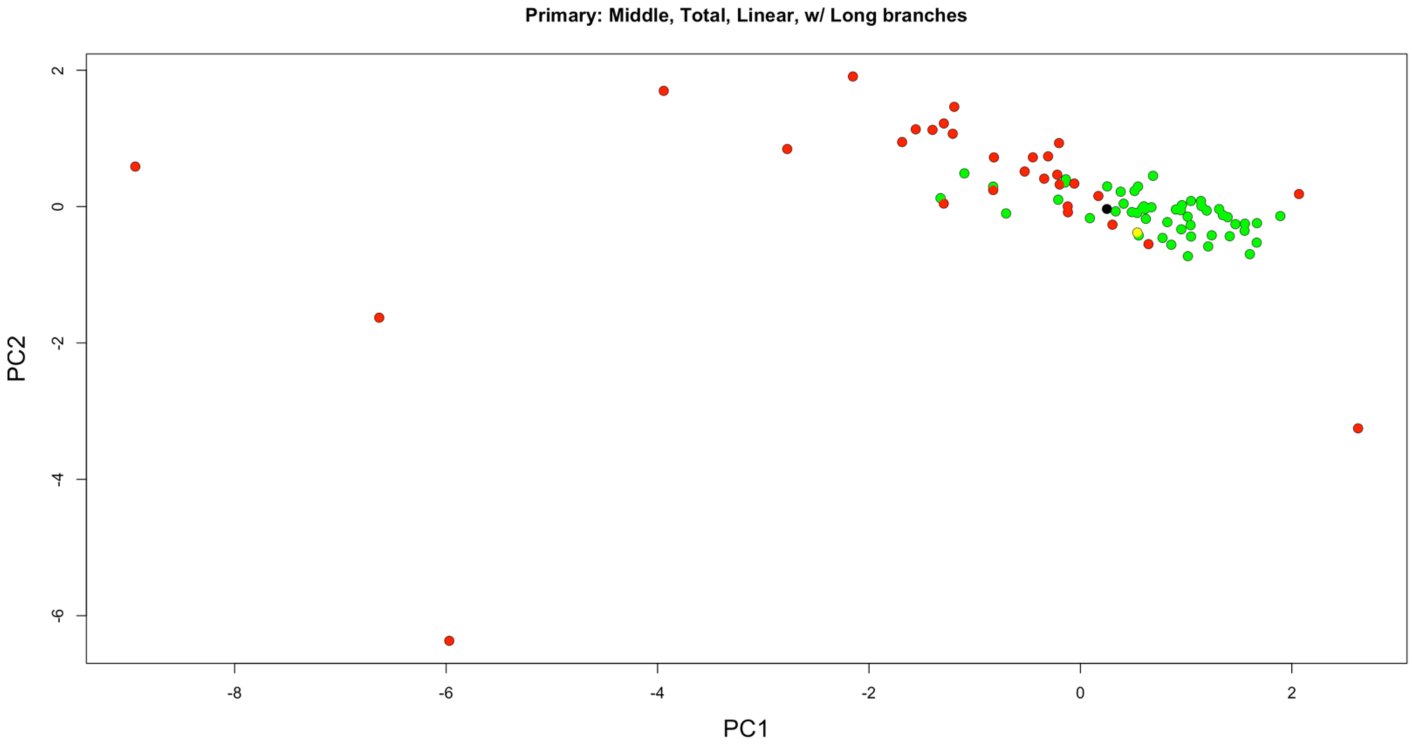

Figure S26. PCA of primaries of all clades, with penguins/tubenoses and ratites/tinamous included. Green are volant taxa, red are flightless taxa, yellow are poor flying/'incipiently flightless' taxa, and black are taxa of unknown flight capability. N = 83 taxa.

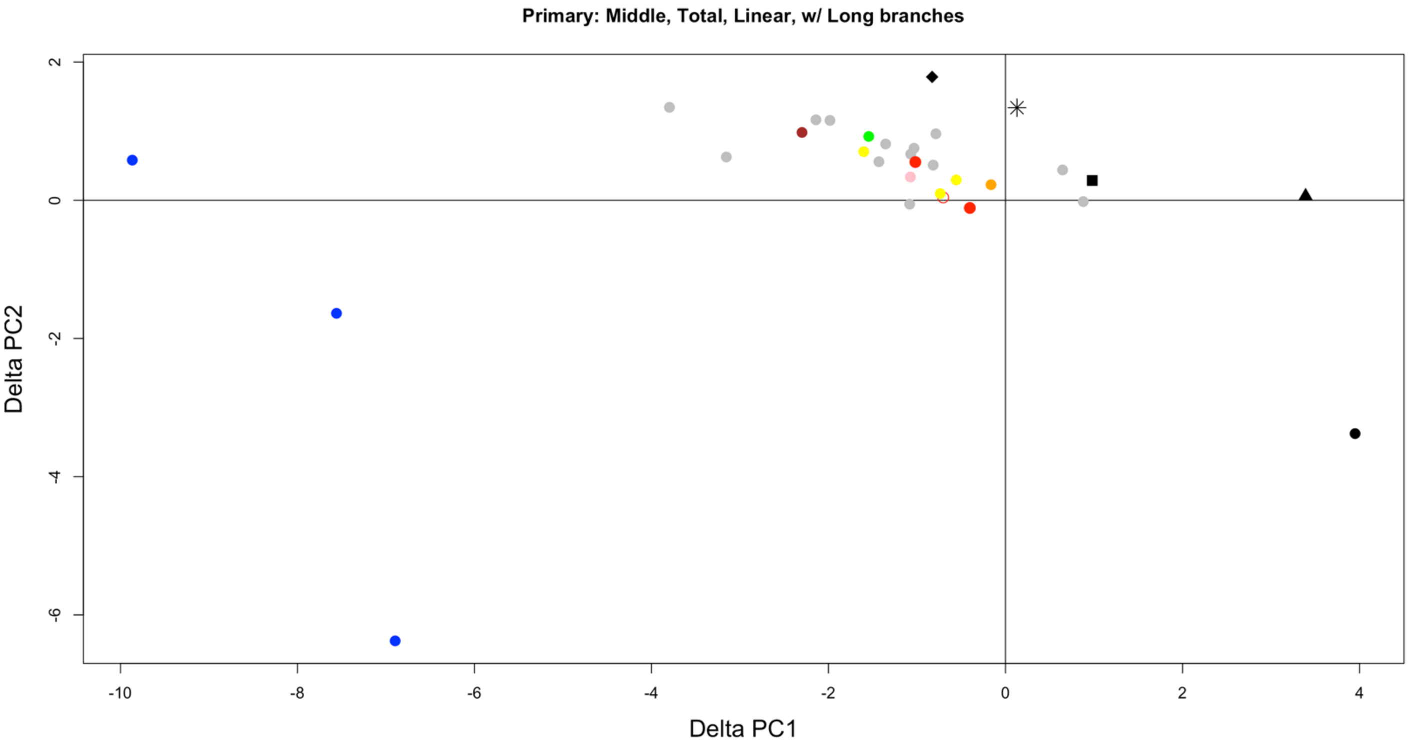


Figure S27. Simple pairwise phylogenetic PCA shift of primaries during flight loss based on an initial morphospace of all clades, with penguins/tubenoses and ratites/tinamous included. Color and shape indicate taxonomy as in Figure 3. Open circle is poor flyer/'incipiently flightless'. N = 31 taxa

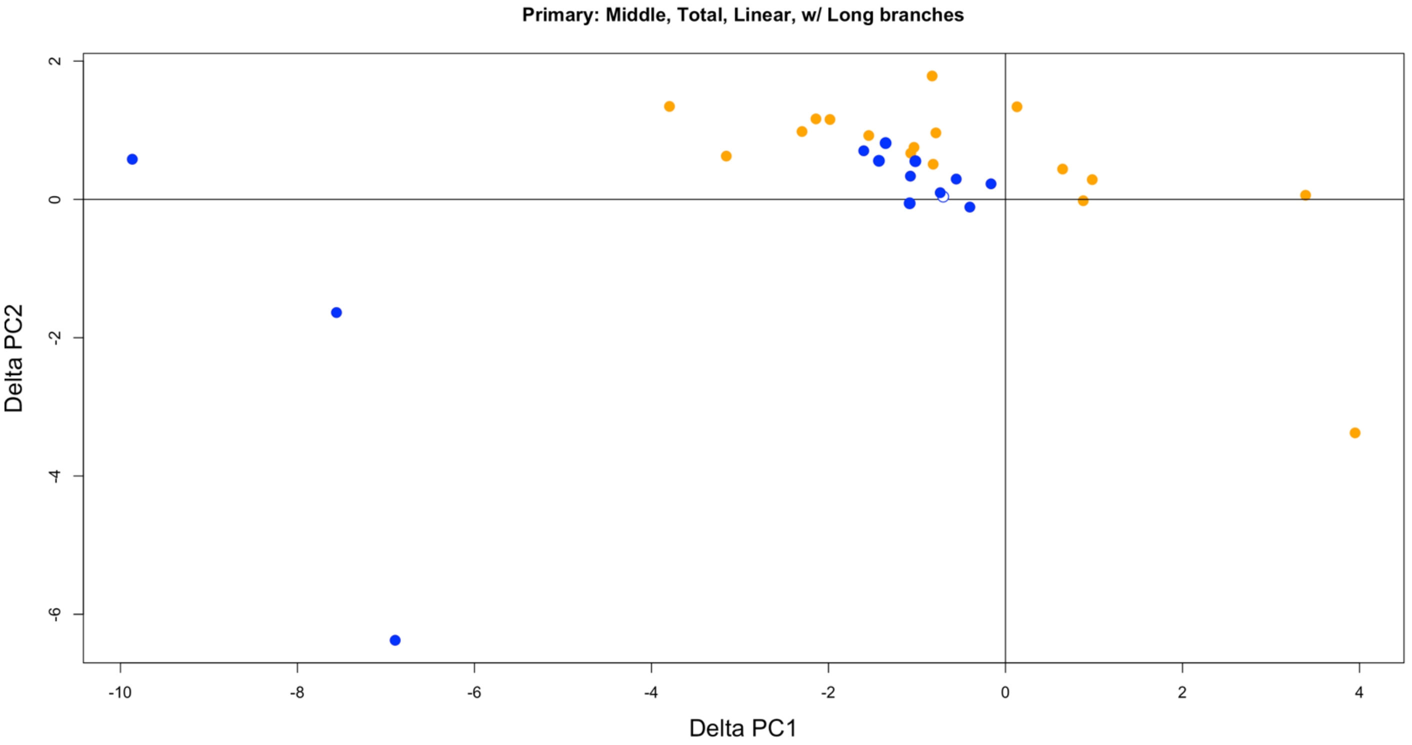


Figure S28. Simple pairwise phylogenetic PCA shift of primaries during flight loss based on an initial morphospace of all clades, with penguins/tubenoses and ratites/tinamous included. Blue are semiaquatic, and orange are terrestrial. Open circle is poor flyer/'incipiently flightless'. N = 31 taxa.

#### 4.2.3. Primary: no long branches (i.e., without Palaeognathae, Sphenisciformes, and Procellariiformes), PC space established by only clades with flightless taxa

PC1 and PC2 explain ~58.85% and ~19.06% of the variation, respectively.

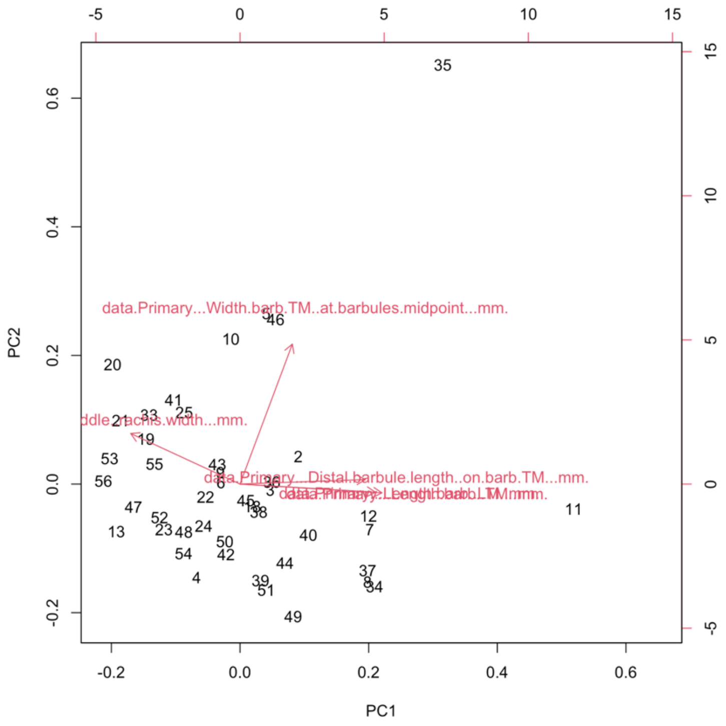


Figure S29. PCA biplot of primaries of only clades with flightless taxa, except penguins/tubenoses and ratites/tinamous. Red arrows indicate variable loadings. Leading and trailing barb lengths have similar loadings. N = 44 taxa.

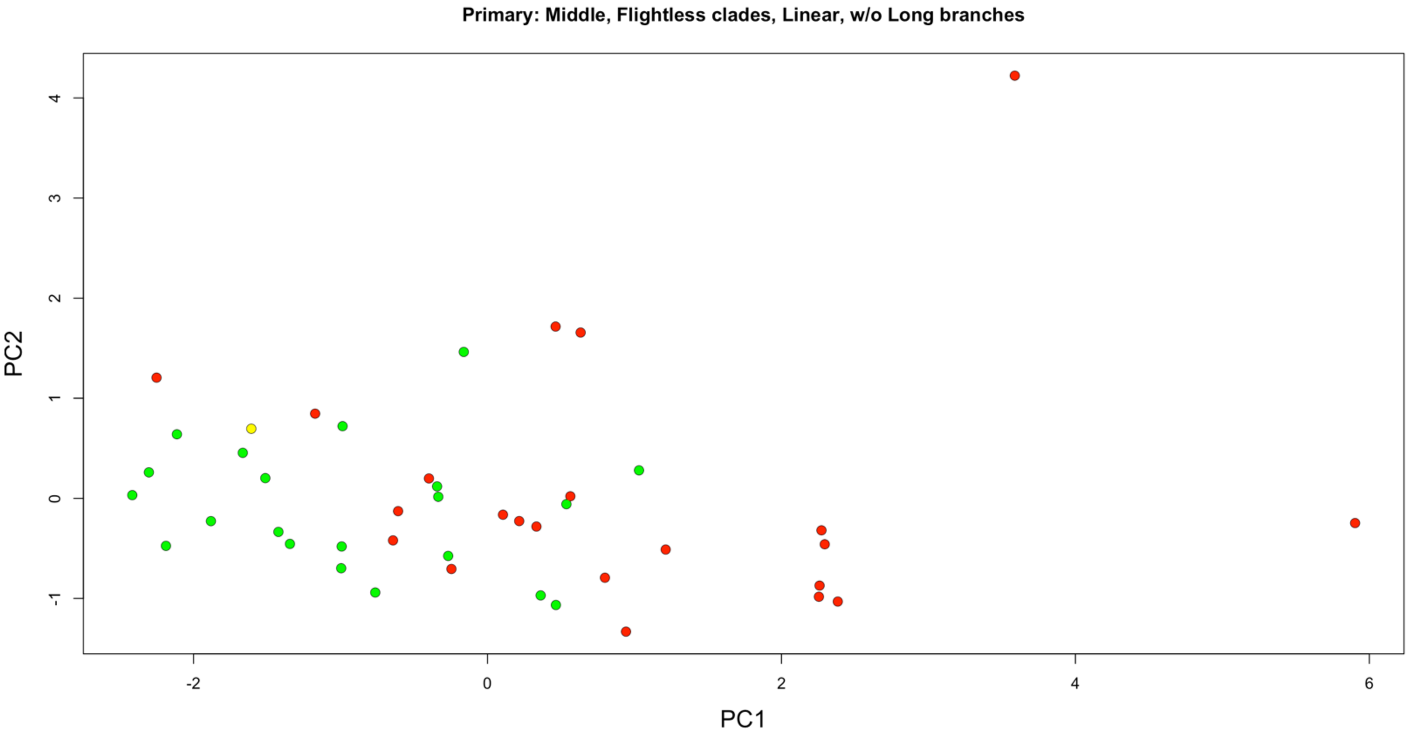

Figure S30. PCA of primaries of only clades with flightless taxa, except penguins/tubenoses and ratites/tinamous. Green are volant taxa, red are flightless taxa, and yellow are poor flying/'incipiently flightless' taxa. N = 44 taxa.

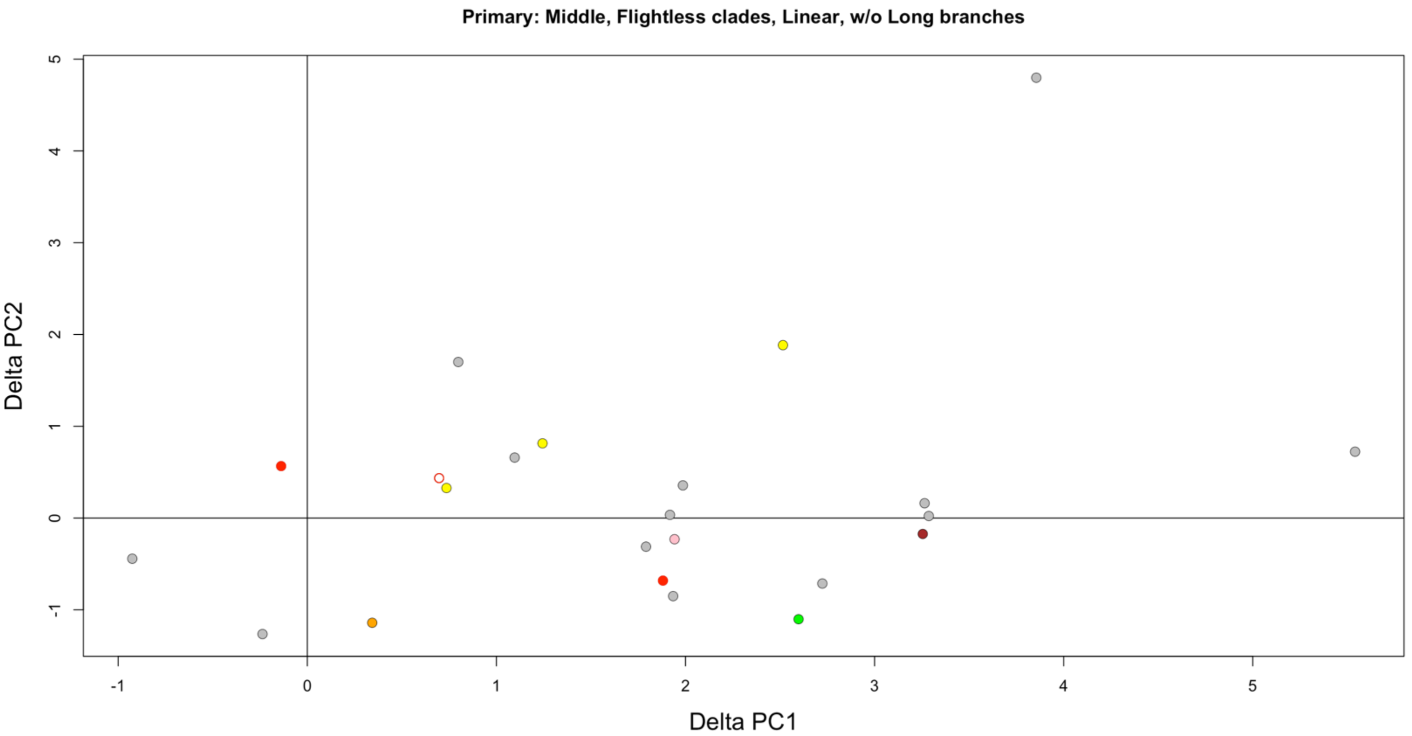


Figure S31. Simple pairwise phylogenetic PCA shift of primaries during flight loss based on an initial morphospace of only clades with flightless taxa, except penguins/tubenoses and ratites/tinamous. Color indicates clade as in Figure 3. Open circle is poor flyer/'incipiently flightless'. N = 23 taxa.

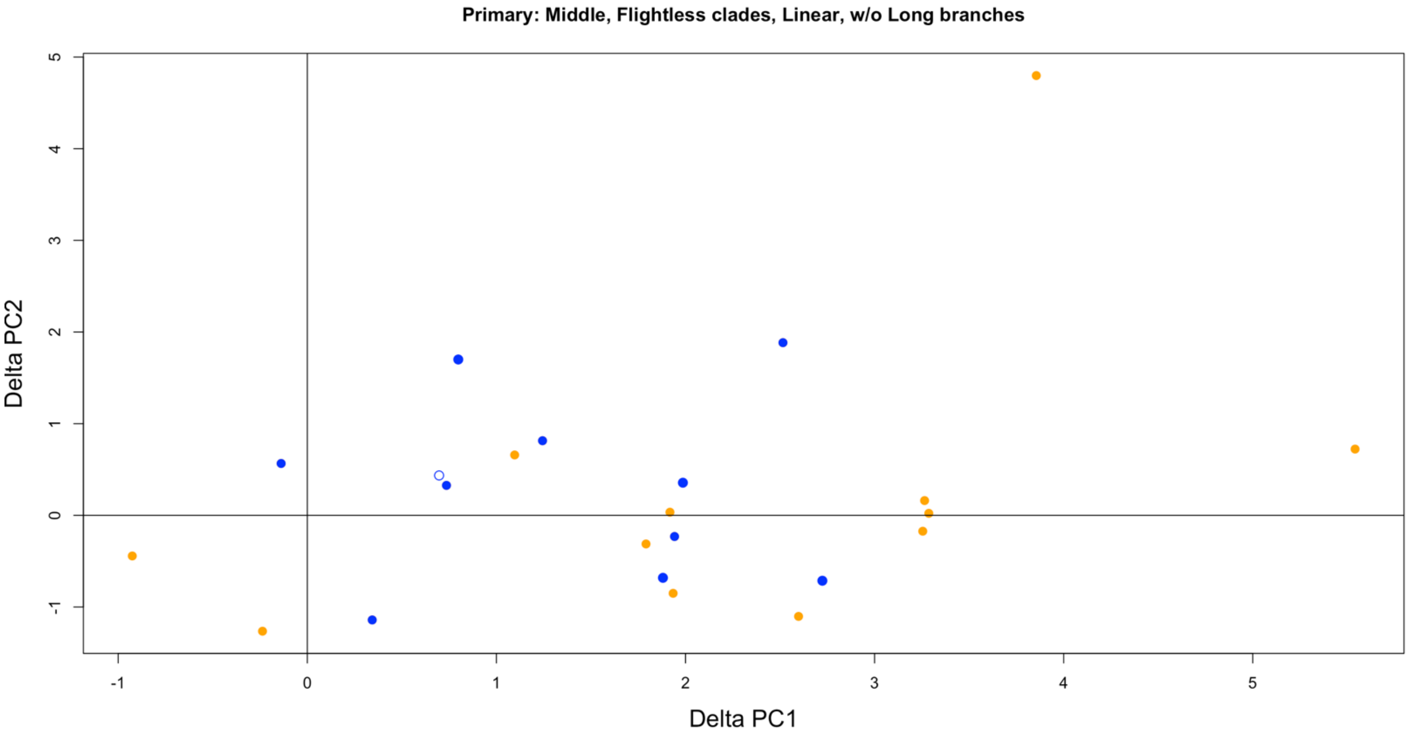


Figure S32. Simple pairwise phylogenetic PCA shift of primaries during flight loss based on an initial morphospace of only clades with flightless taxa, except penguins/tubenoses and ratites/tinamous. Blue are semiaquatic, and orange are terrestrial. Open circle is poor flyer/'incipiently flightless'. N = 23 taxa.

#### 4.2.4. Primary: no long branches (i.e., without Palaeognathae, Sphenisciformes, and Procellariiformes), PC space established by all clades

PC1 and PC2 explain ~55.51% and ~21.82% of the variation, respectively.

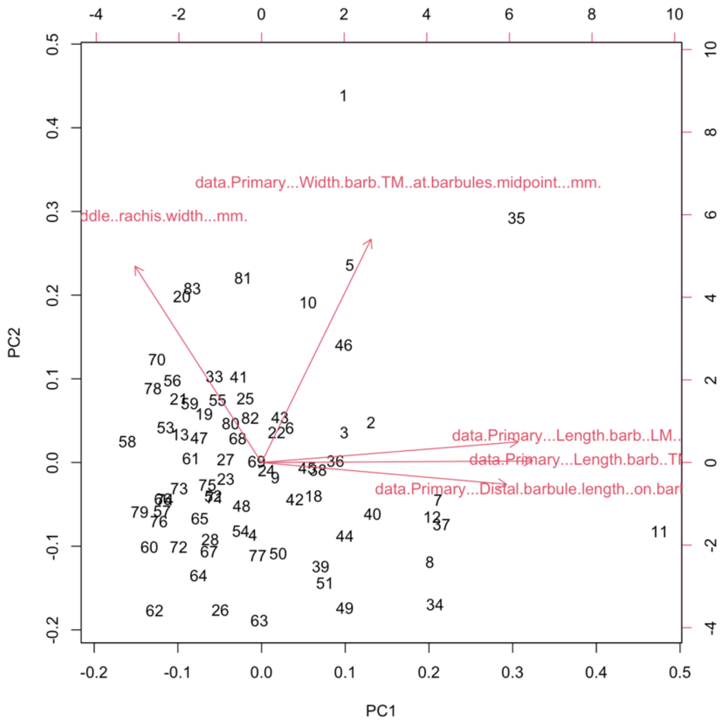


Figure S33. PCA biplot of primaries of all clades, except penguins/tubenoses and ratites/tinamous. Red arrows indicate variable loadings. N = 75 taxa.

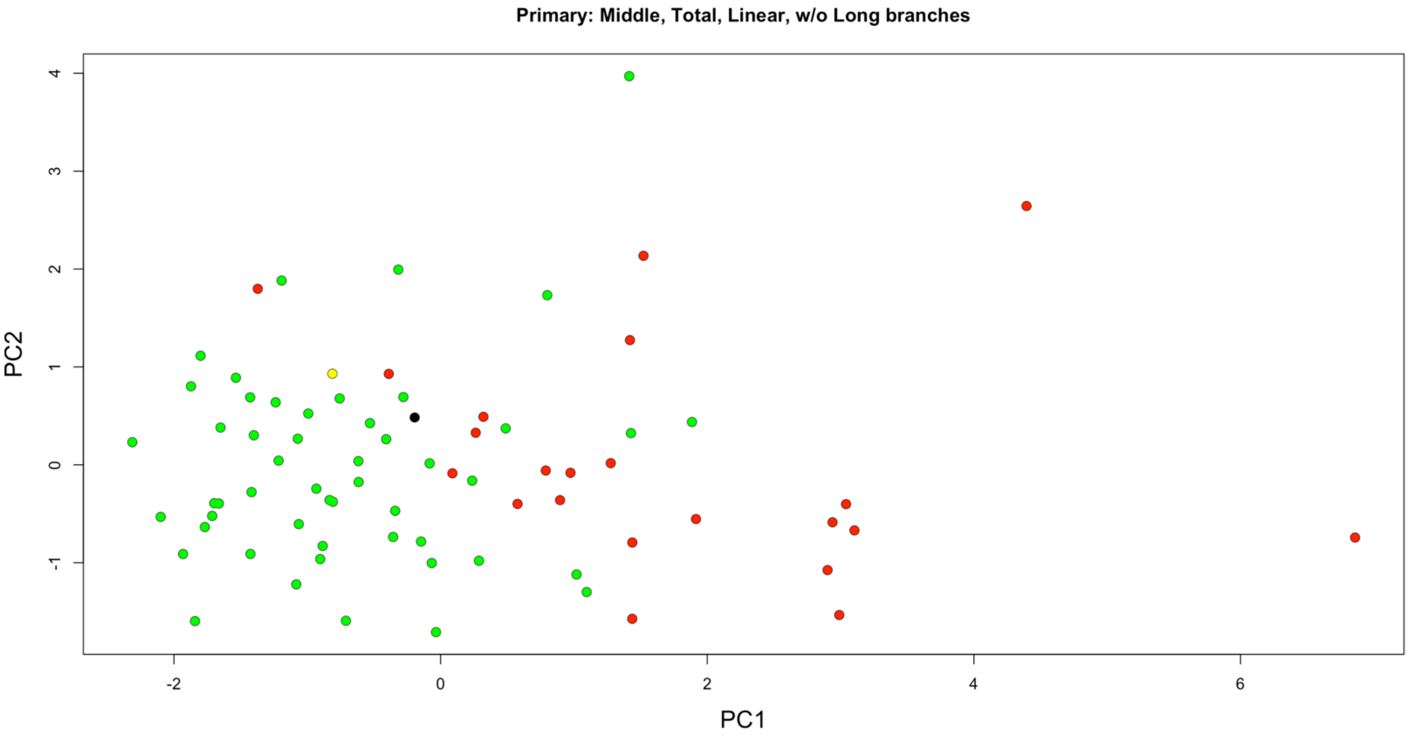

Figure S34. PCA of primaries of all clades, except penguins/tubenoses and ratites/tinamous. Green are volant taxa, red are flightless taxa, yellow are poor flying/‘incipiently flightless’ taxa, and black are taxa of unknown flight capability. N = 75 taxa.

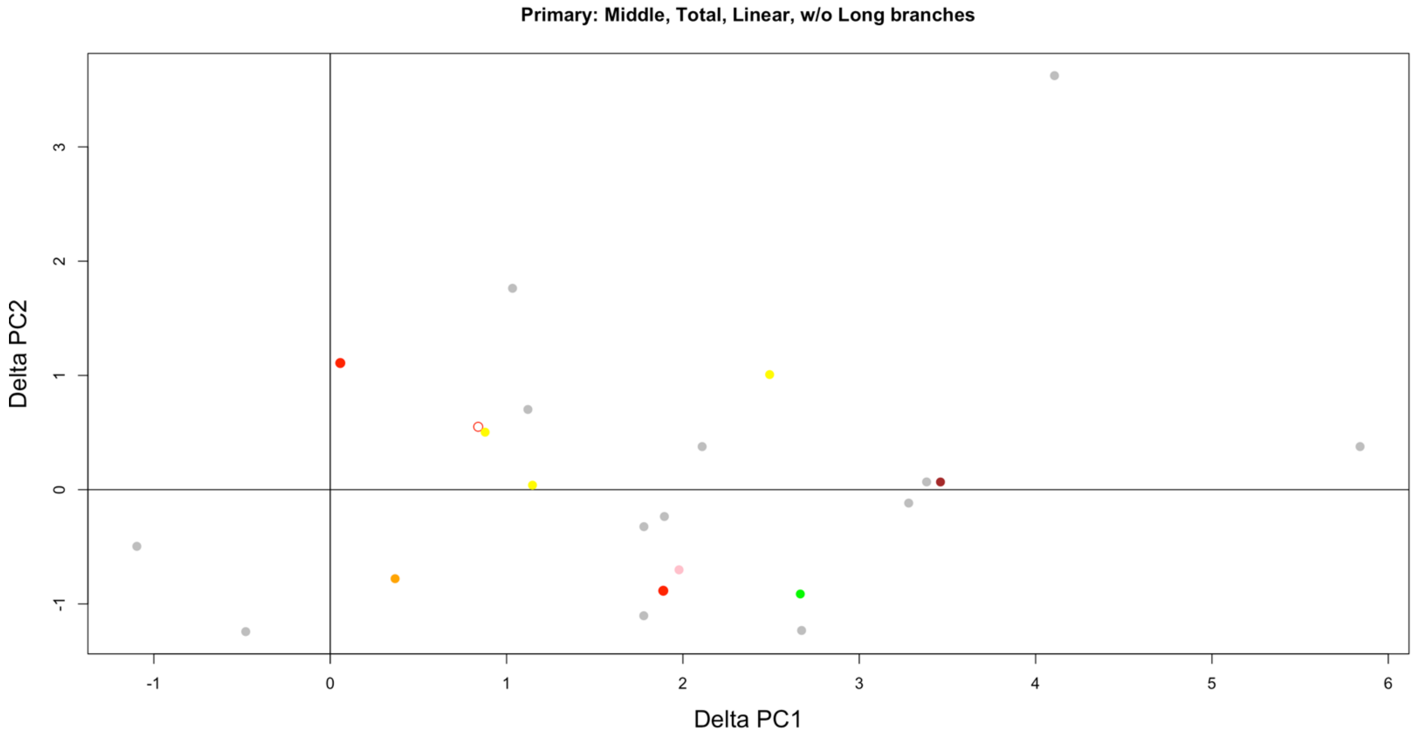


Figure S35. Simple pairwise phylogenetic PCA shift of primaries during flight loss based on an initial morphospace of all clades, except penguins/tubenoses and ratites/tinamous. Color indicates clade as in Figure 3. Open circle is poor flyer/‘incipiently flightless’. N = 23 taxa.

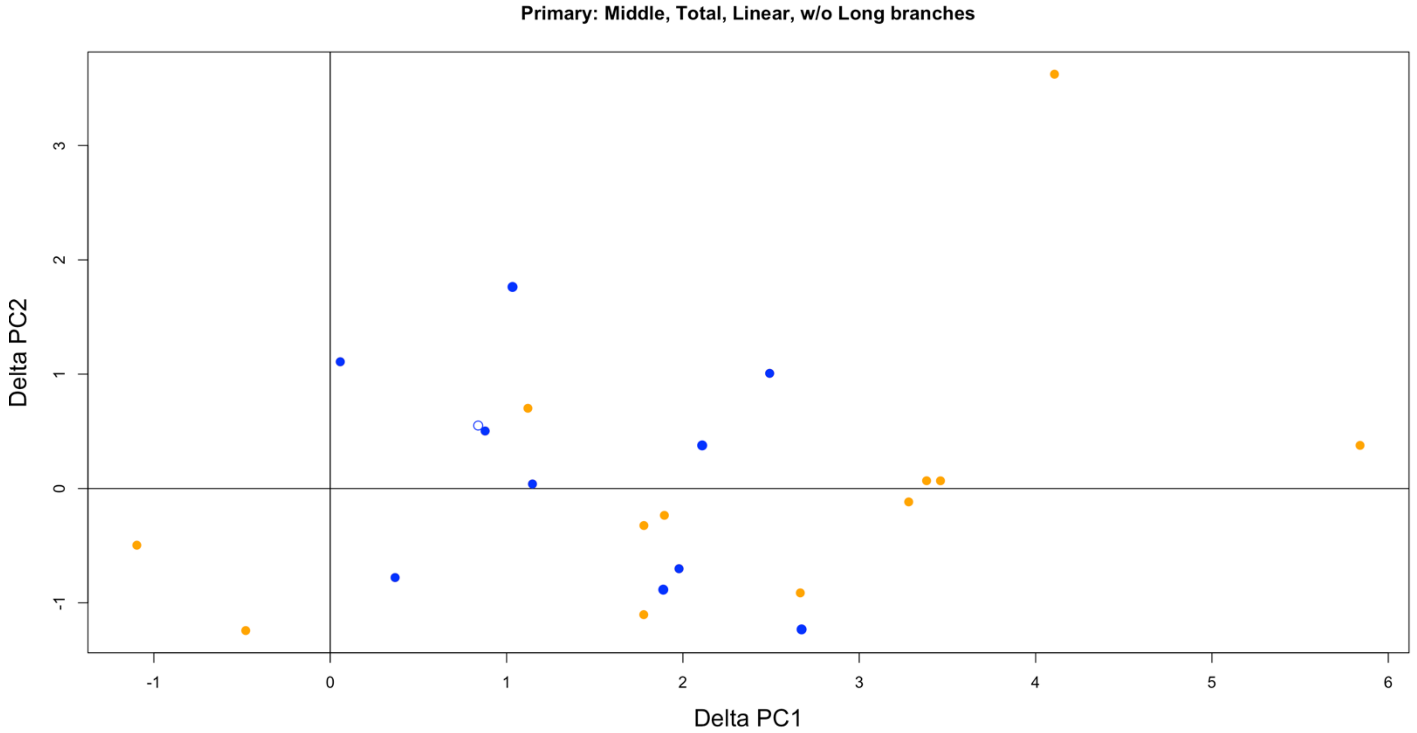


Figure S36. Simple pairwise phylogenetic PCA shift of primaries during flight loss based on an initial morphospace of all clades, except penguins/tubenoses and ratites/tinamous. Blue are semiaquatic, and orange are terrestrial. Open circle is poor flyer/‘incipiently flightless’. N = 23 taxa.

### 4.3. Tertials

#### 4.3.1. Tertial: long branches, PC space established by only clades with flightless taxa

PC1 and PC2 explain ~58.39% and ~28.05% of the variation, respectively.

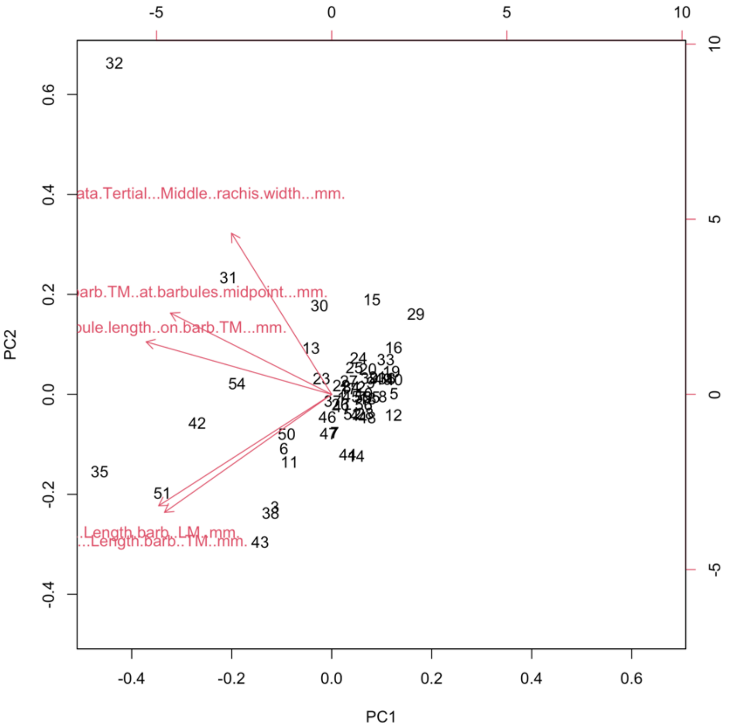


Figure S37. PCA biplot of tertials of only clades with flightless taxa, with penguins/tubenoses and ratites/tinamous included. Red arrows indicate variable loadings. N = 56 taxa.

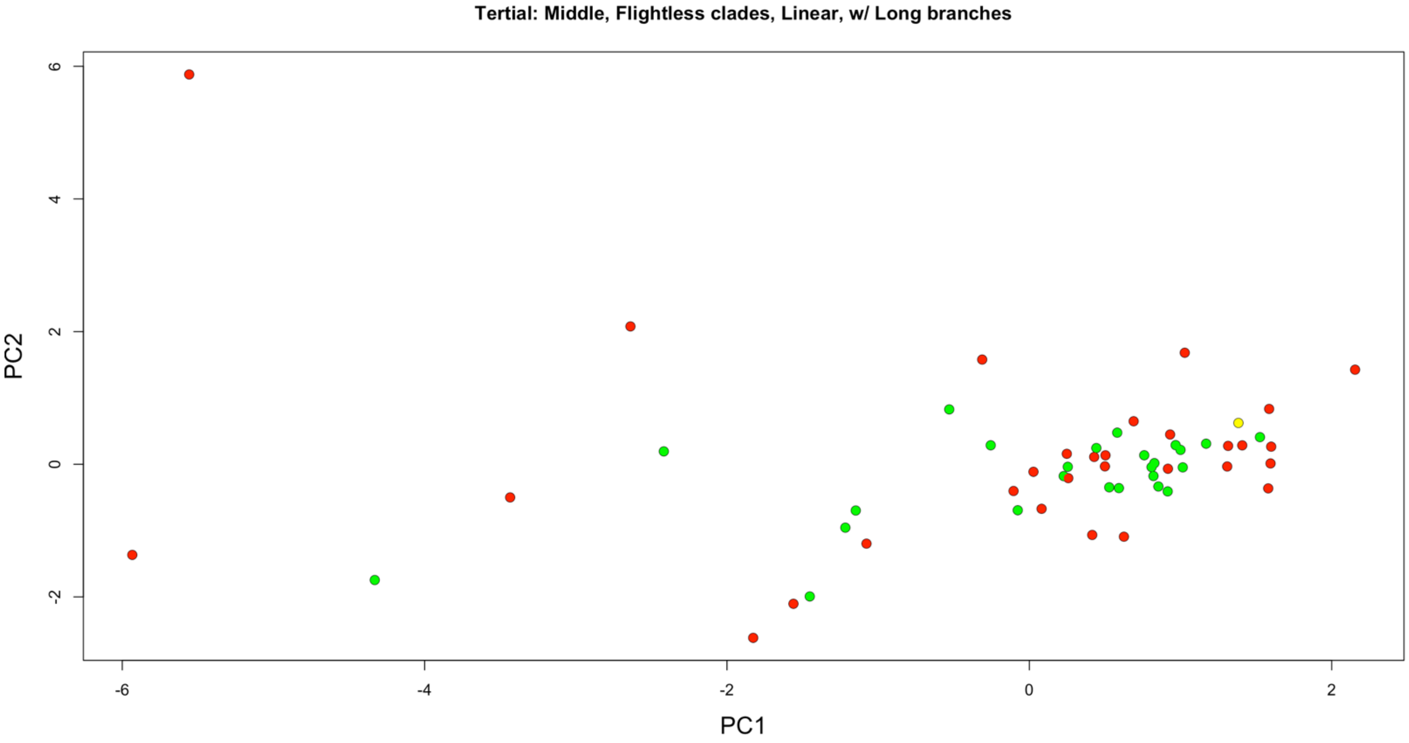

Figure S38. PCA of tertials of only clades with flightless taxa, with penguins/tubenoses and ratites/tinamous included. Green are volant taxa, red are flightless taxa, and yellow are poor flying/'incipiently flightless' taxa. N = 56 taxa.

#### 4.3.2. Tertial: no long branches (i.e., without Palaeognathae, Sphenisciformes, and Procellariiformes), PC space established by only clades with flightless taxa

PC1 and PC2 explain ~65.82% and ~20.80% of the variation, respectively.

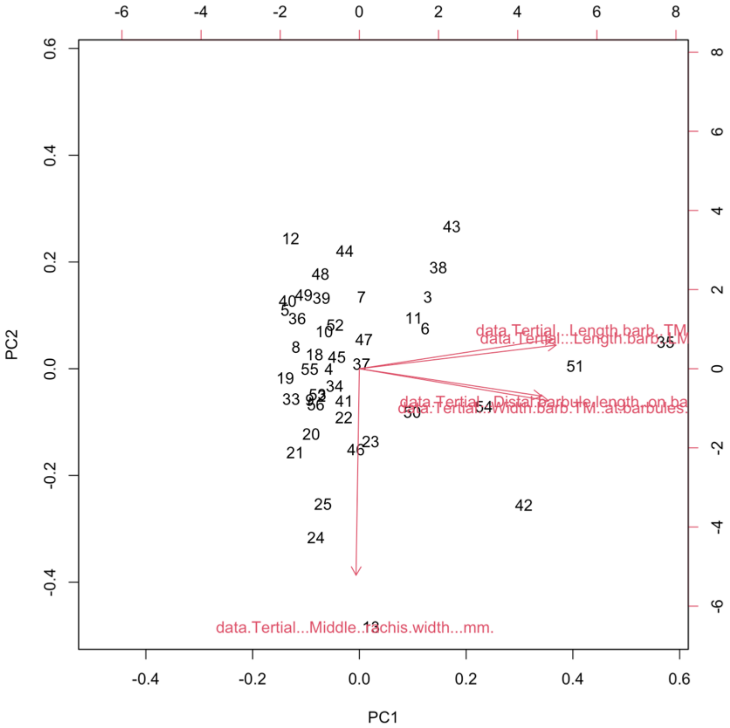


Figure S39. PCA biplot of tertials of only clades with flightless taxa, except penguins/tubenoses and ratites/tinamous. Red arrows indicate variable loadings. Leading and trailing barb lengths have similar loadings. Barbule length and barb width have similar loadings. N = 44 taxa.

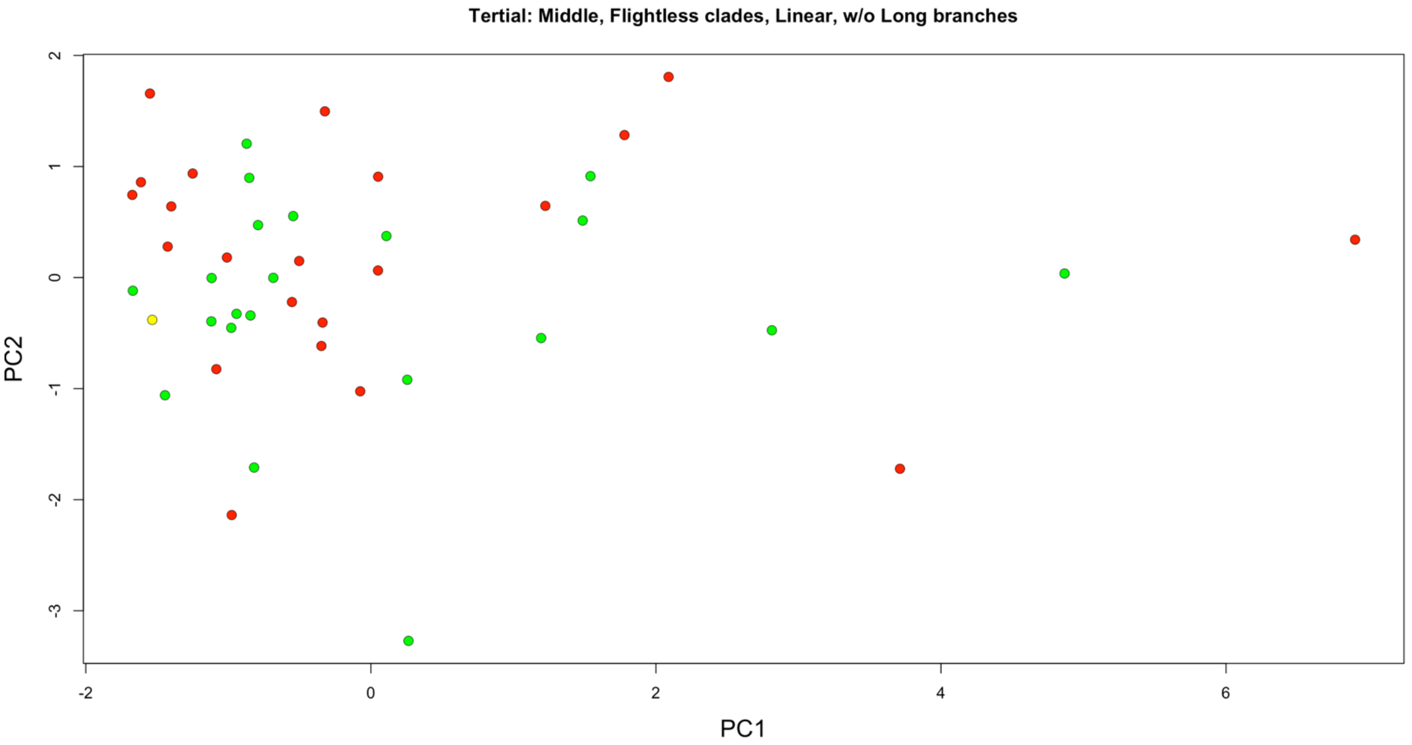


Figure S40. PCA of tertials of only clades with flightless taxa, except penguins/tubenoses and ratites/tinamous. Green are volant taxa, red are flightless taxa, and yellow are poor flying/'incipiently flightless' taxa. N = 44 taxa.

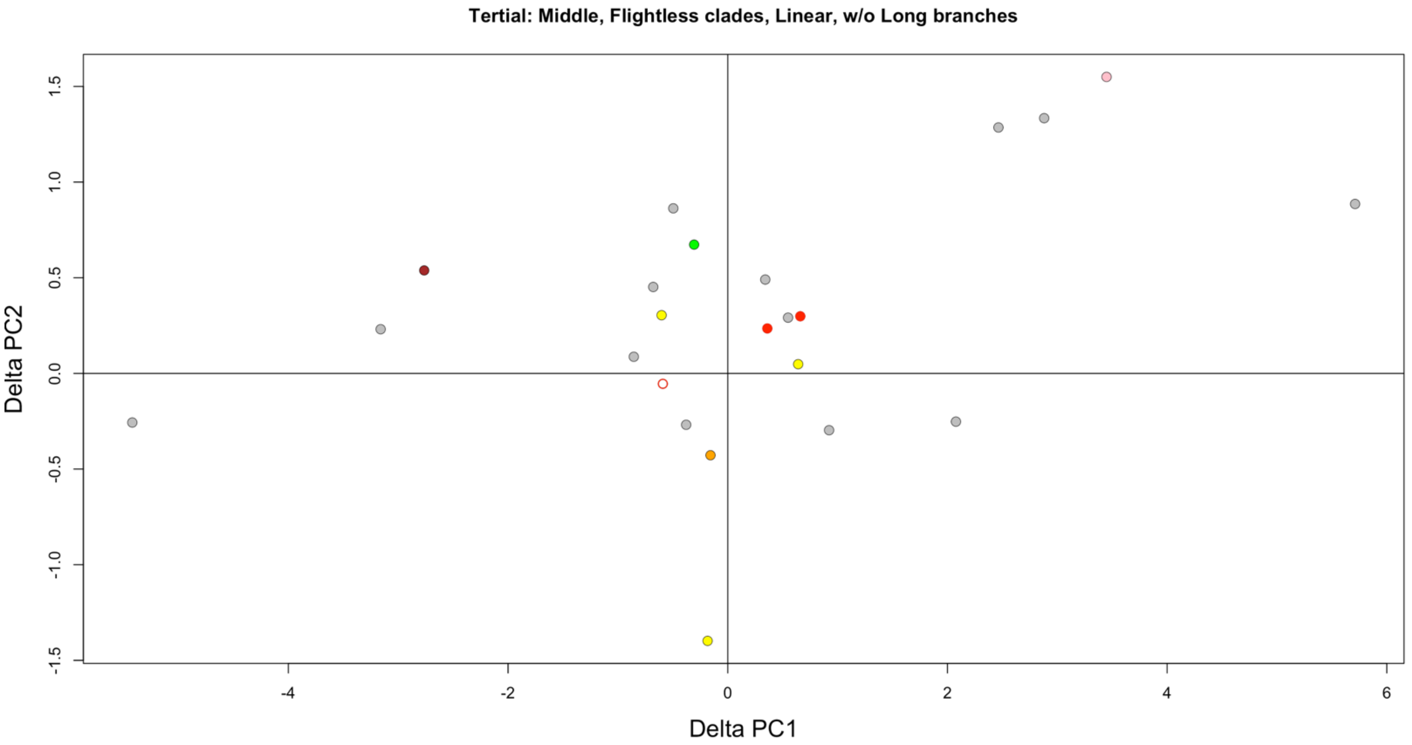


Figure S41. Simple pairwise phylogenetic PCA shift of tertials during flight loss based on an initial morphospace of only clades with flightless taxa, except penguins/tubenoses and ratites/tinamous. Color indicates clade as in Figure 3. Open circle is poor flyer/'incipiently flightless'. N = 23 taxa.

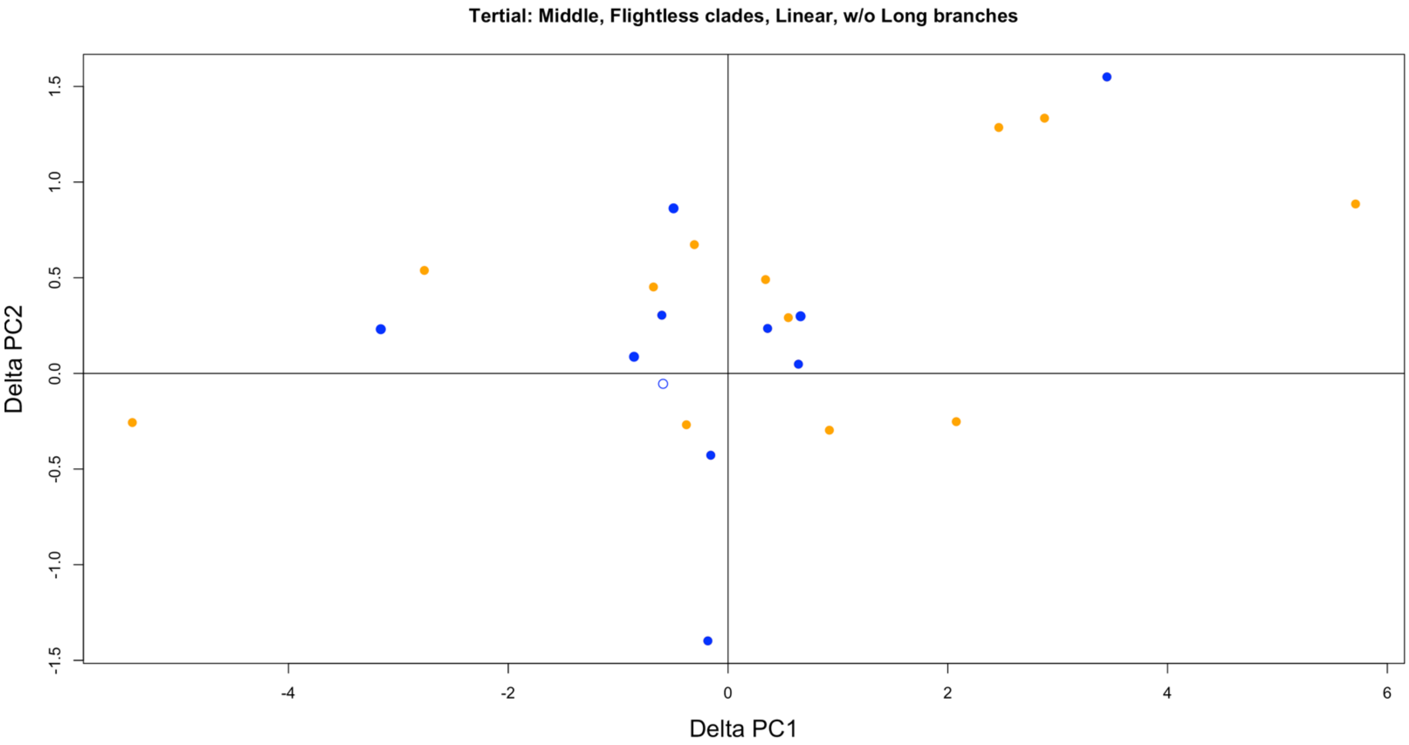


Figure S42. Simple pairwise phylogenetic PCA shift of tertials during flight loss based on an initial morphospace of only clades with flightless taxa, except penguins/tubenoses and ratites/tinamous. Blue are semiaquatic, and orange are terrestrial. Open circle is poor flyer/‘incipiently flightless’. N = 23 taxa.

### 4.4. Rectrices

#### 4.4.1. Rectrix: long branches, PC space established by only clades with flightless taxa

PC1 and PC2 explain ~58.66% and ~22.61% of the variation, respectively.

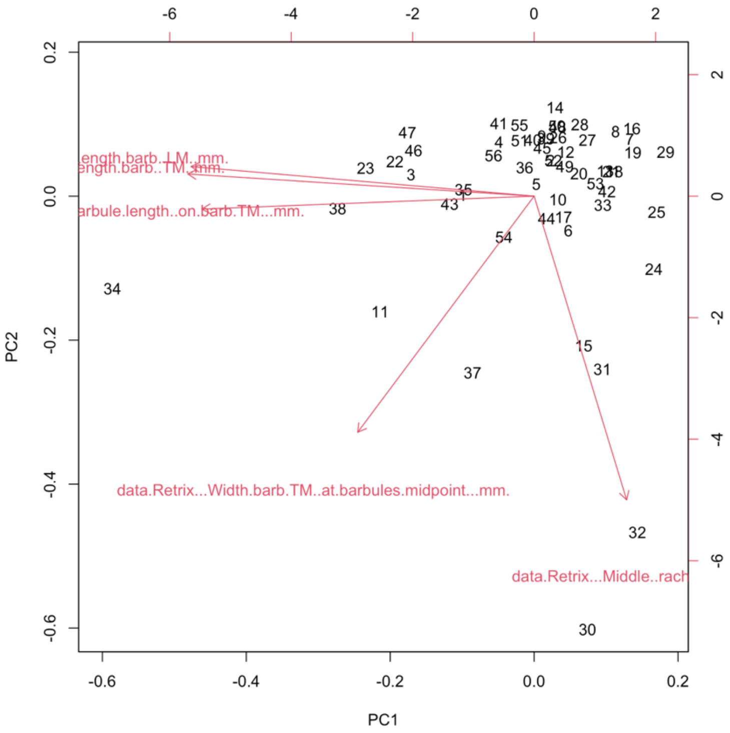

Figure S43. PCA biplot of rectrices of only clades with flightless taxa, with penguins/tubenoses and ratites/tinamous included. Red arrows indicate variable loadings. N = 56 taxa.

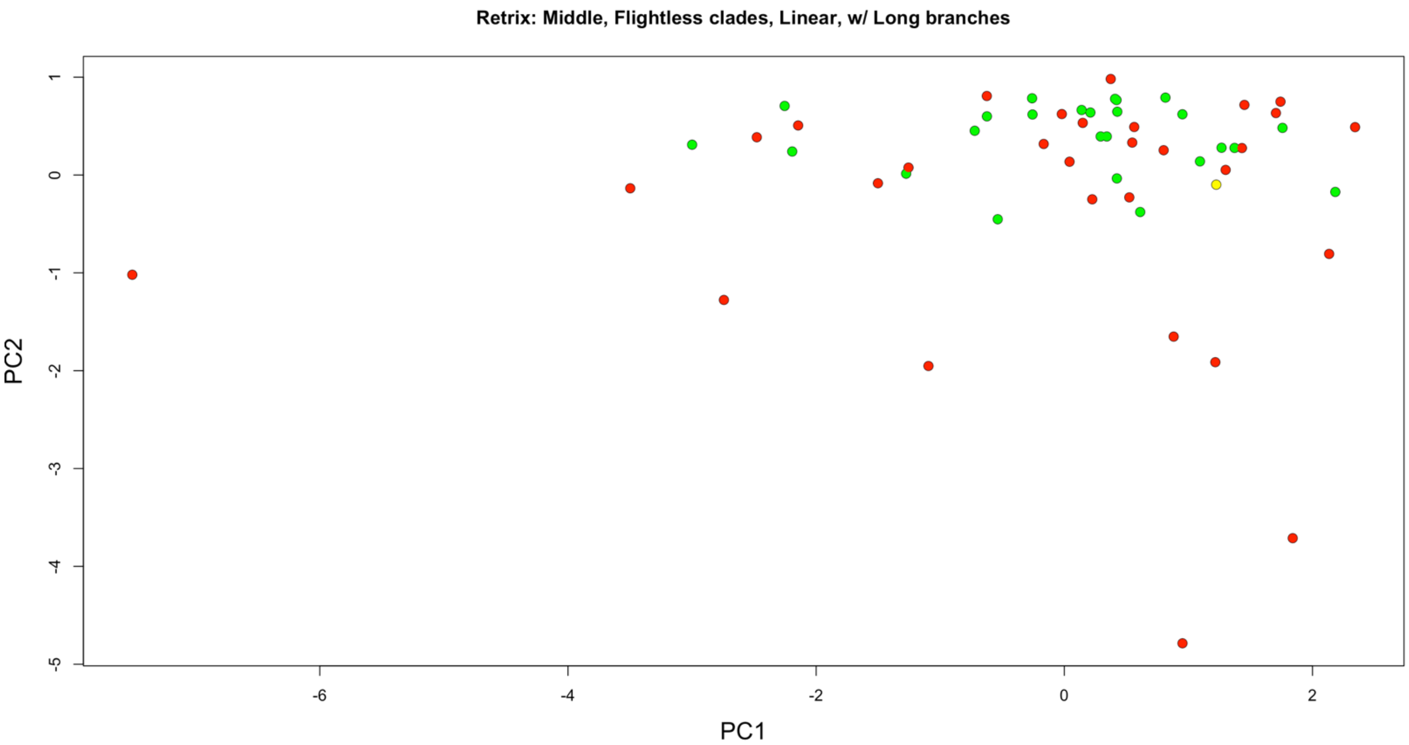


Figure S44. PCA of rectrices of only clades with flightless taxa, with penguins/tubenoses and ratites/tinamous included. Green are volant taxa, red are flightless taxa, and yellow are poor flying/'incipiently flightless' taxa. N = 56 taxa.

#### 4.4.2. Rectrix: no long branches (i.e., without Palaeognathae, Sphenisciformes, and Procellariiformes), PC space established by only clades with flightless taxa

PC1 and PC2 explain ~67.20% and ~18.00% of the variation, respectively.

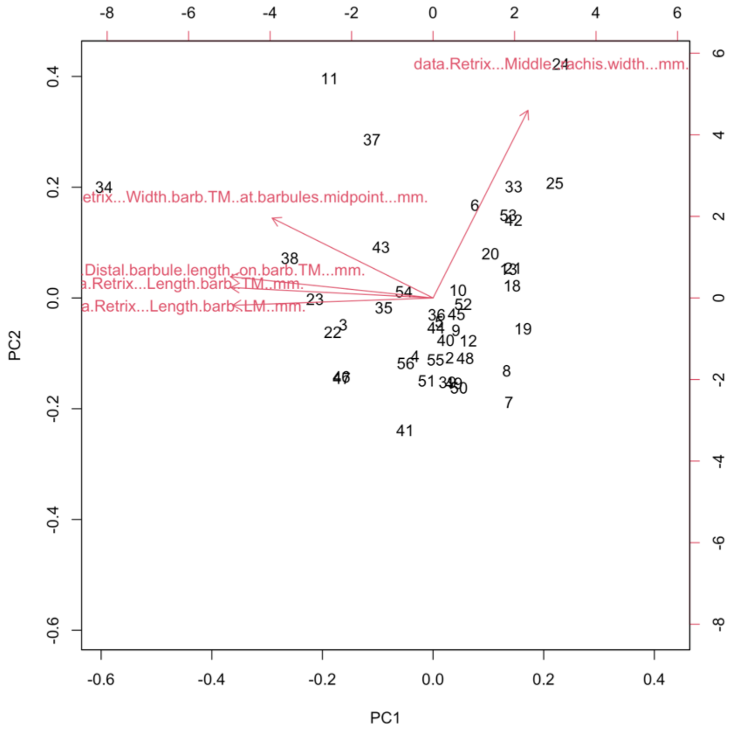


Figure S45. PCA biplot of rectrices of only clades with flightless taxa, except penguins/tubenoses and ratites/tinamous. Red arrows indicate variable loadings. N = 44 taxa.

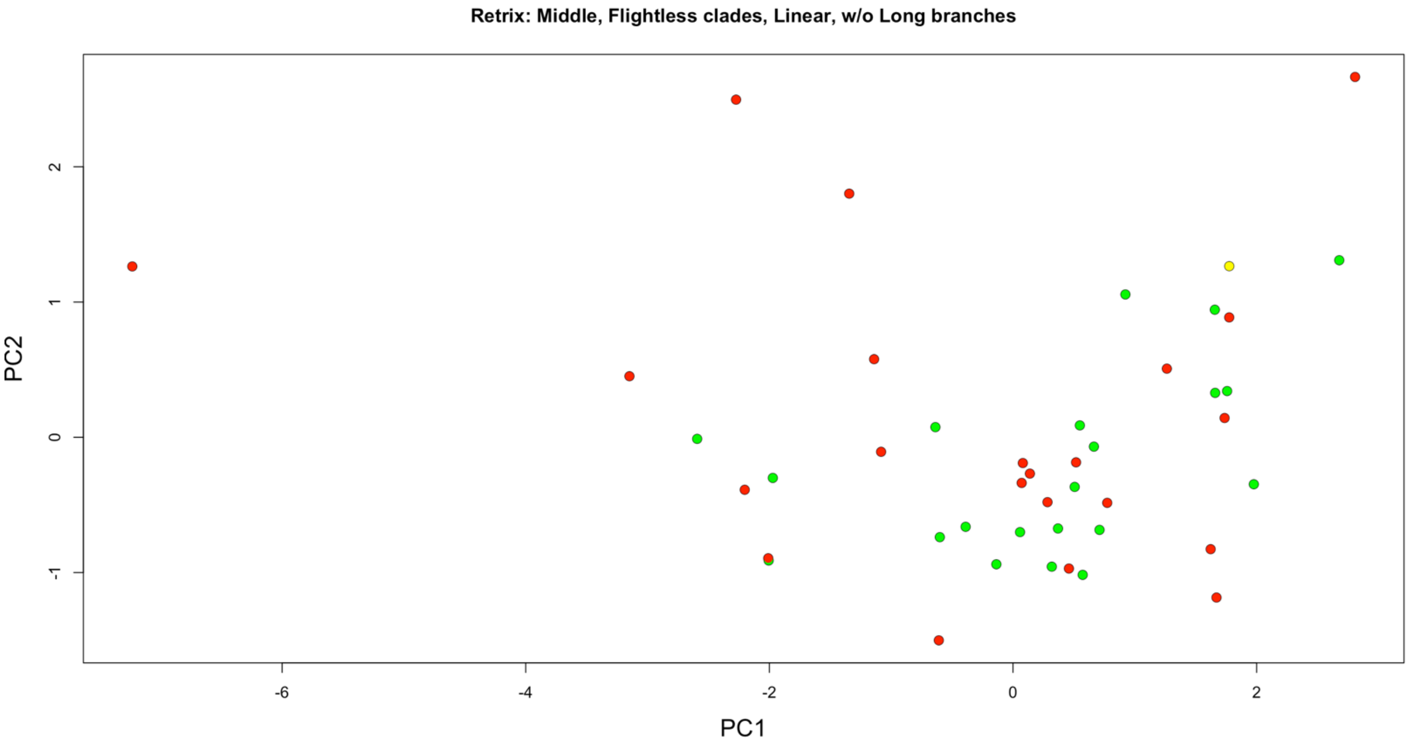


Figure S46. PCA of rectrices of only clades with flightless taxa, except penguins/tubenoses and ratites/tinamous. Green are volant taxa, red are flightless taxa, and yellow are poor flying/'incipiently flightless' taxa. N = 44 taxa.

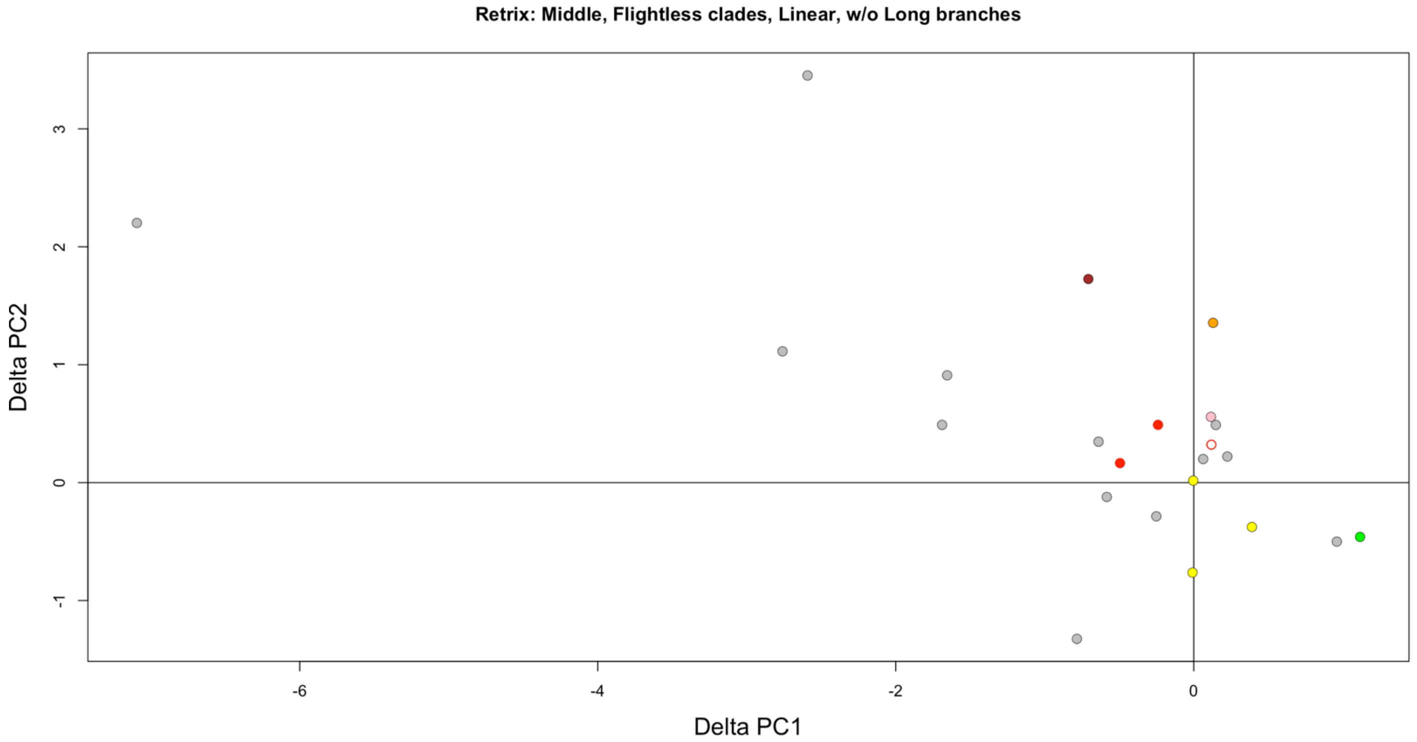

Figure S47. Simple pairwise phylogenetic PCA shift of rectrices during flight loss based on an initial morphospace of only clades with flightless taxa, except penguins/tubenoses and ratites/tinamous. Color indicates clade as in Figure 3. Open circle is poor flyer/'incipiently flightless'. N = 23 taxa.

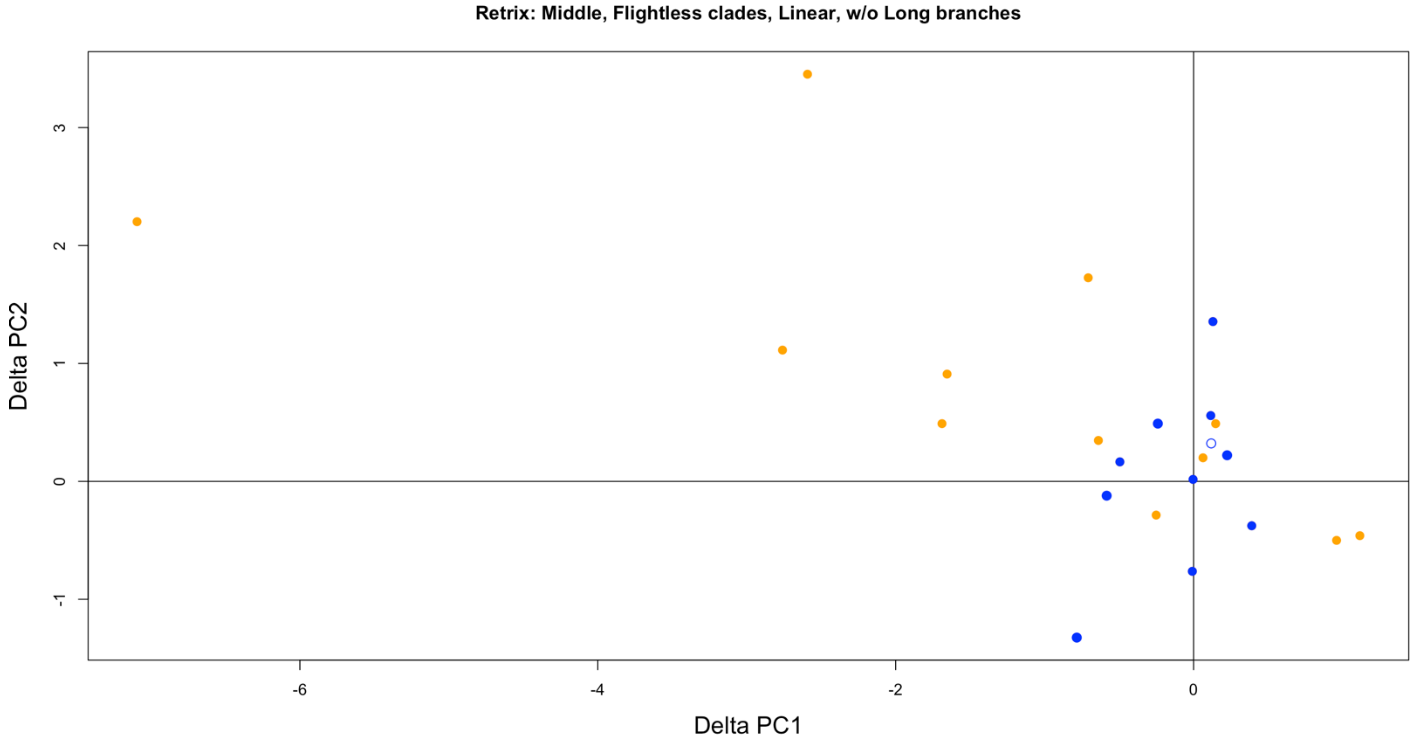

Figure S48. Simple pairwise phylogenetic PCA shift of rectrices during flight loss based on an initial morphospace of only clades with flightless taxa, except penguins/tubenoses and ratites/tinamous. Blue are semiaquatic, and orange are terrestrial. Open circle is poor flyer/'incipiently flightless'. N = 23 taxa.

#### 4.5. Dorsal contours

##### 4.5.1. Dorsal contour: long branches, PC space established by only clades with flightless taxa

PC1 and PC2 explain ~56.80% and ~21.95% of the variation, respectively.

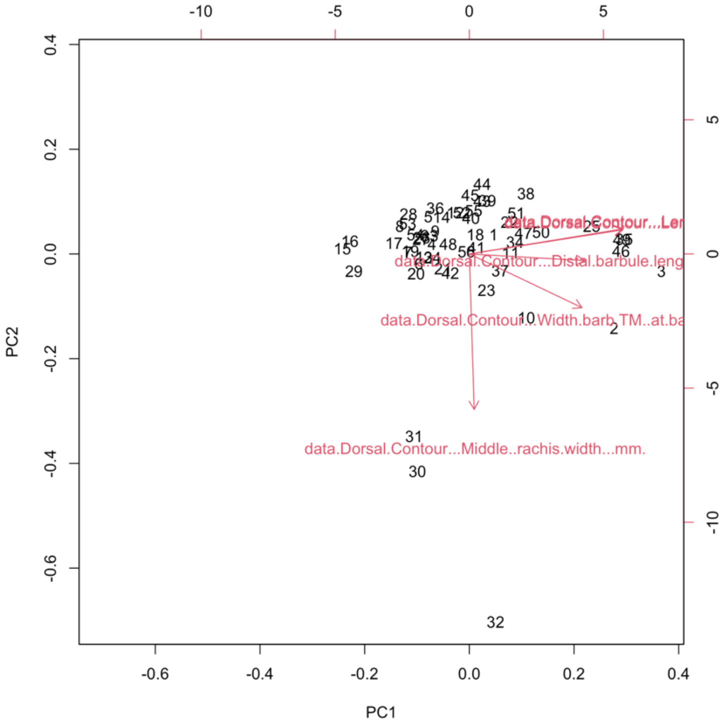


Figure S49. PCA biplot of dorsal contours of only clades with flightless taxa, with penguins/tubenoses and ratites/tinamous included. Red arrows indicate variable loadings. N = 56 taxa.

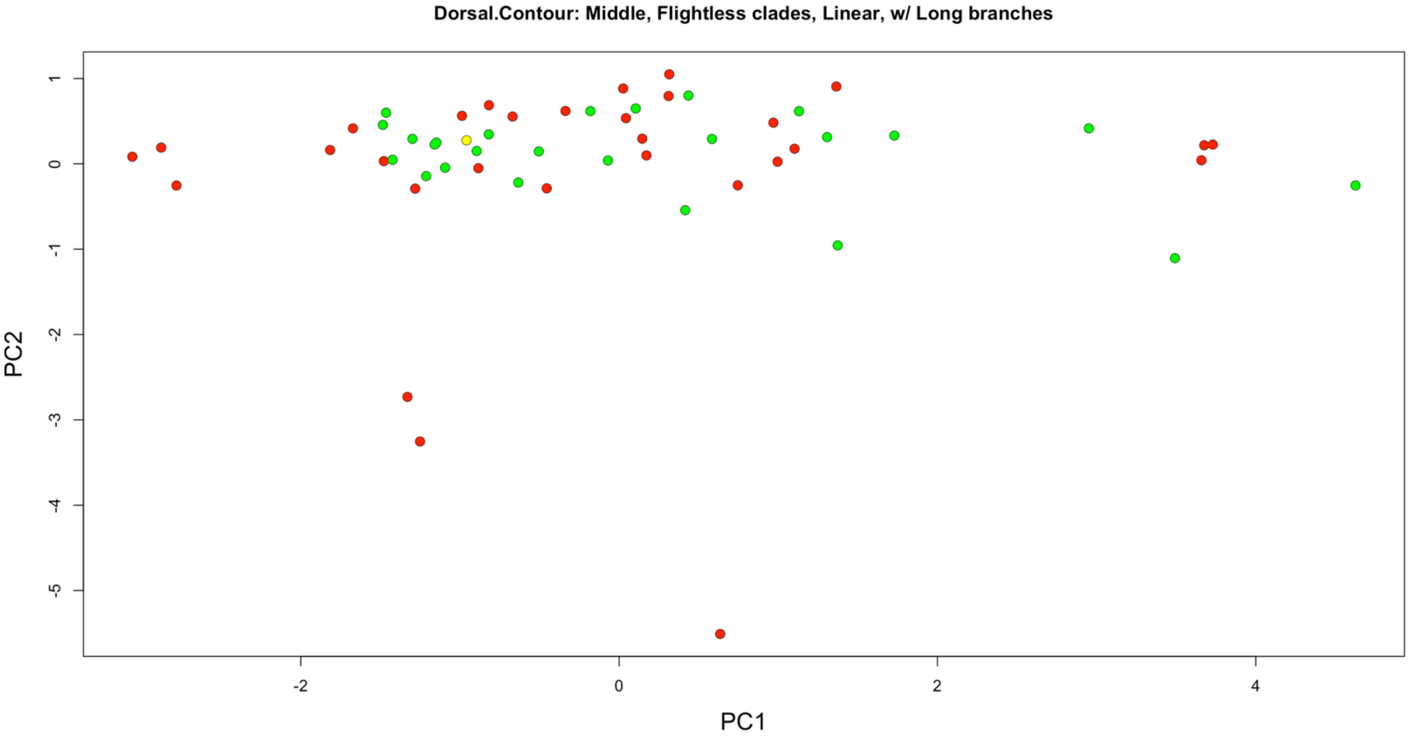

Figure S50. PCA of dorsal contours of only clades with flightless taxa, with penguins/tubenoses and ratites/tinamous included. Green are volant taxa, red are flightless taxa, and yellow are poor flying/‘incipiently flightless’ taxa. N = 56 taxa.

#### 4.5.2. Dorsal contour: no long branches (i.e., without Palaeognathae, Sphenisciformes, and Procellariiformes), PC space established by only clades with flightless taxa

PC1 and PC2 explain ~57.67% and ~21.80% of the variation, respectively.

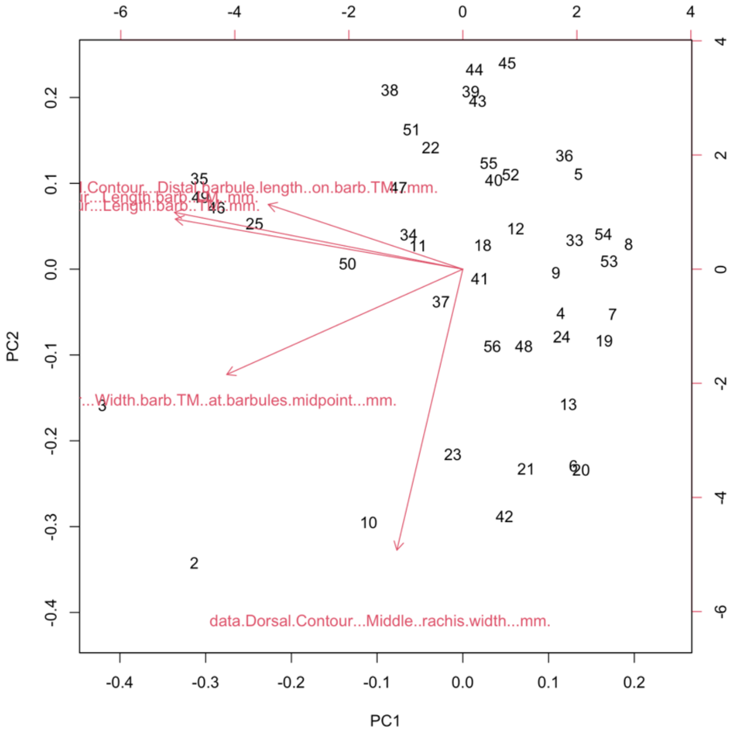


Figure S51. PCA biplot of dorsal contours of only clades with flightless taxa, except penguins/tubenoses and ratites/tinamous. Red arrows indicate variable loadings. N = 44 taxa.

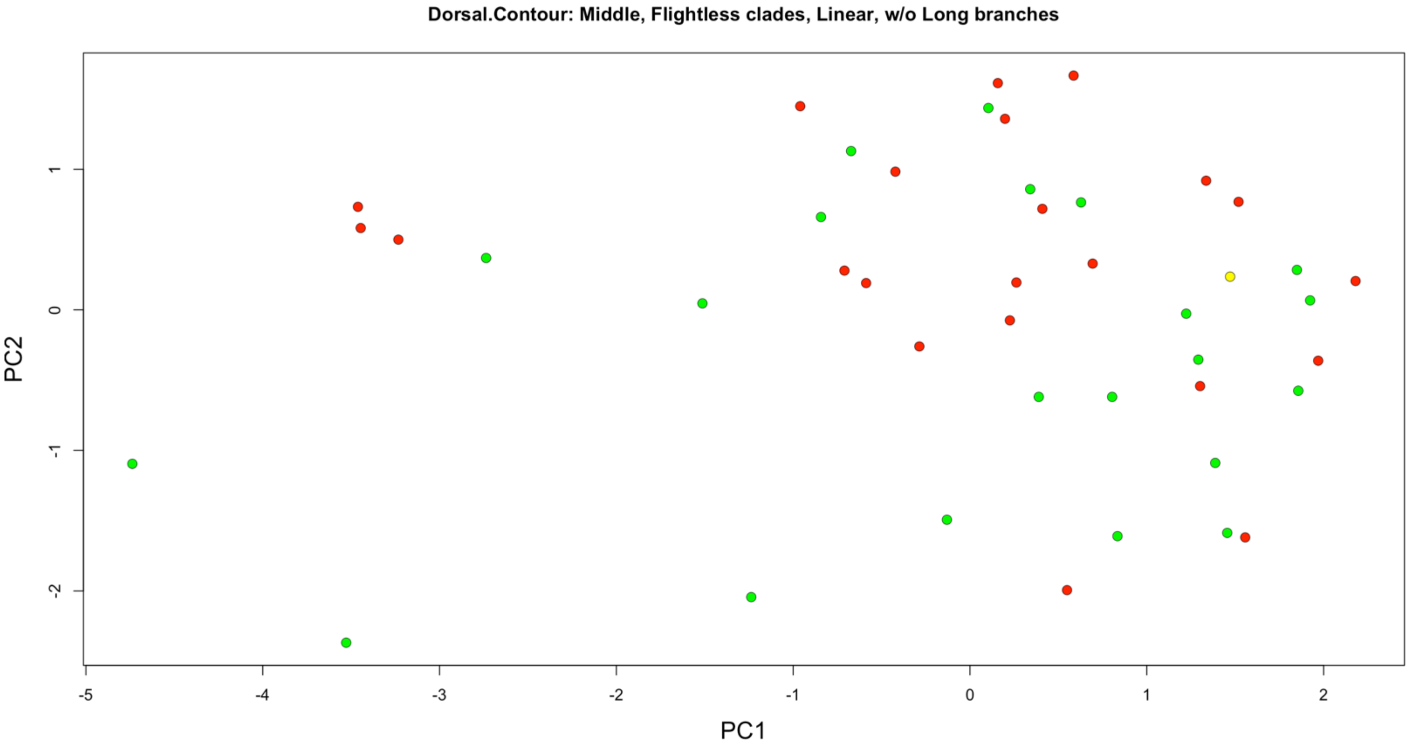


Figure S52. PCA of dorsal contours of only clades with flightless taxa, except penguins/tubenoses and ratites/tinamous. Green are volant taxa, red are flightless taxa, and yellow are poor flying/'incipiently flightless' taxa. N = 44 taxa.

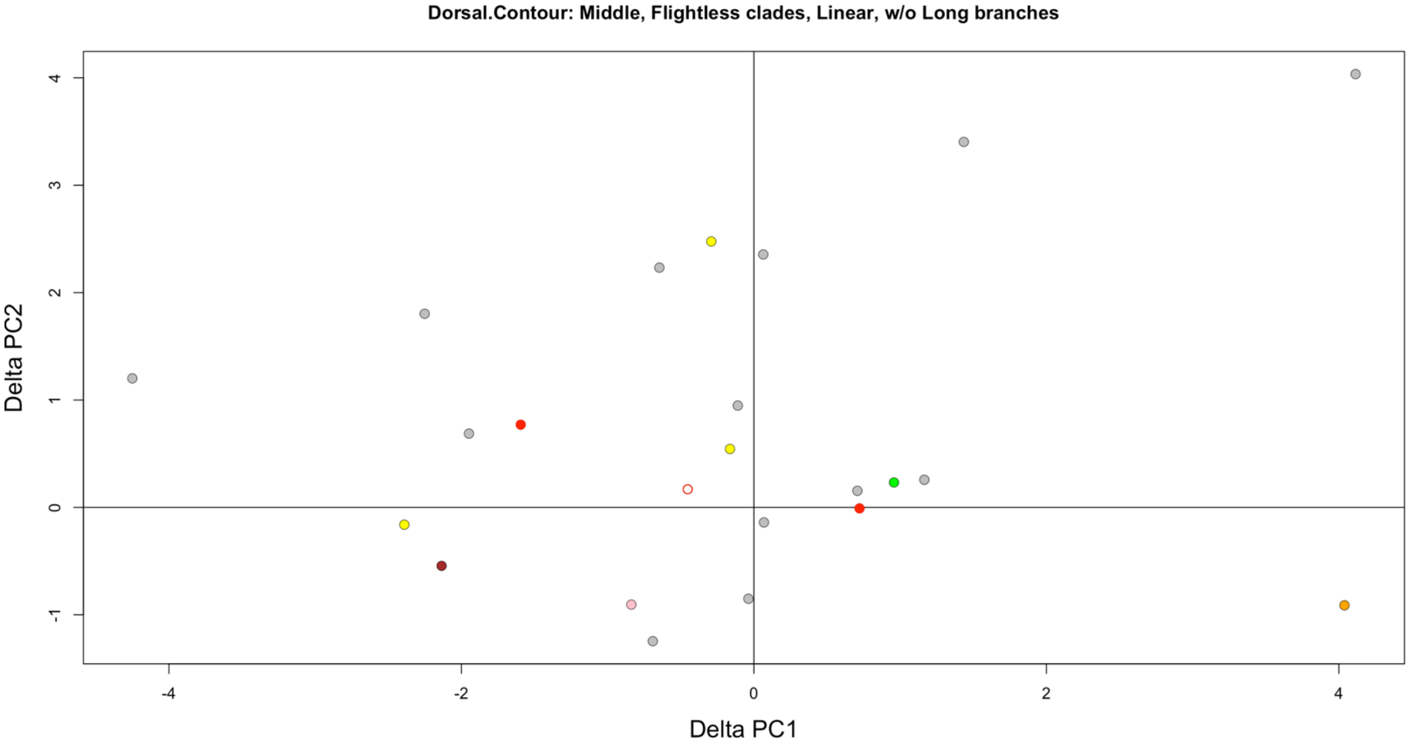


Figure S53. Simple pairwise phylogenetic PCA shift of dorsal contours during flight loss based on an initial morphospace of only clades with flightless taxa, except penguins/tubenoses and ratites/tinamous. Color indicates clade as in Figure 3. Open circle is poor flyer/'incipiently flightless'. N = 23 taxa

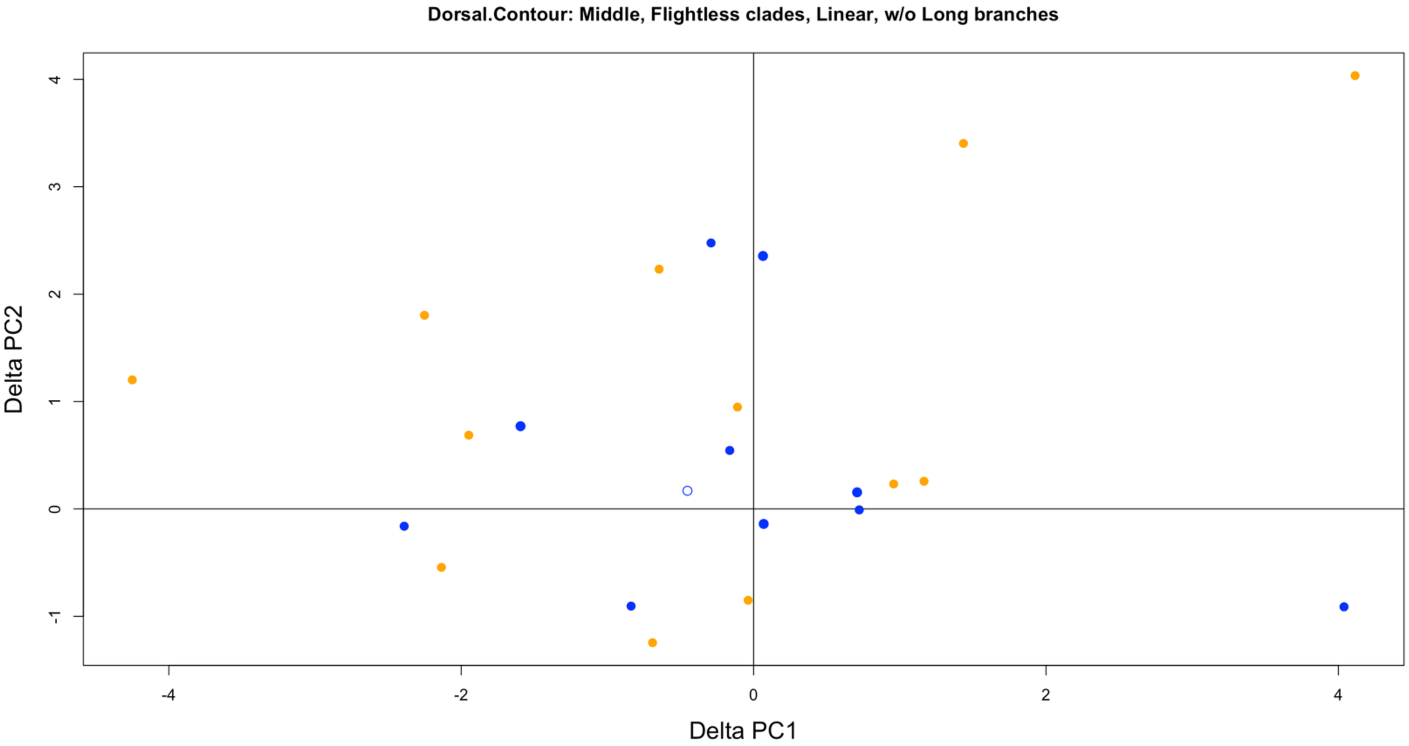


Figure S54. Simple pairwise phylogenetic PCA shift of dorsal contours during flight loss based on an initial morphospace of only clades with flightless taxa, except penguins/tubenoses and ratites/tinamous. Blue are semiaquatic, and orange are terrestrial. Open circle is poor flyer/'incipiently flightless'. N = 23 taxa.

### 4.6. Ventral contours

#### 4.6.1. Ventral contour: long branches, PC space established by only clades with flightless taxa

PC1 and PC2 explain ~66.12% and ~16.25% of the variation, respectively.

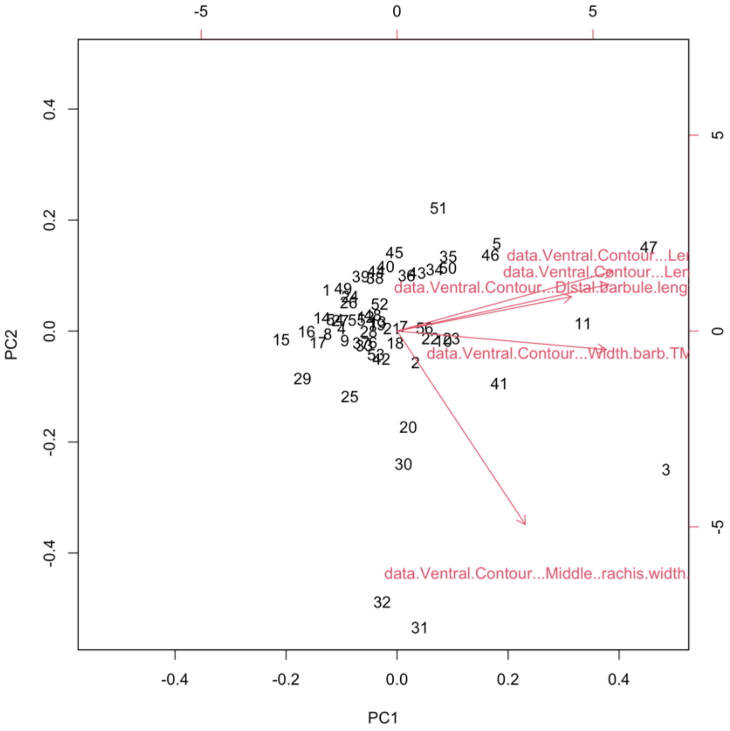

Figure S55. PCA biplot of ventral contours of only clades with flightless taxa, with penguins/tubenoses and ratites/tinamous included. Red arrows indicate variable loadings. N = 56 taxa.

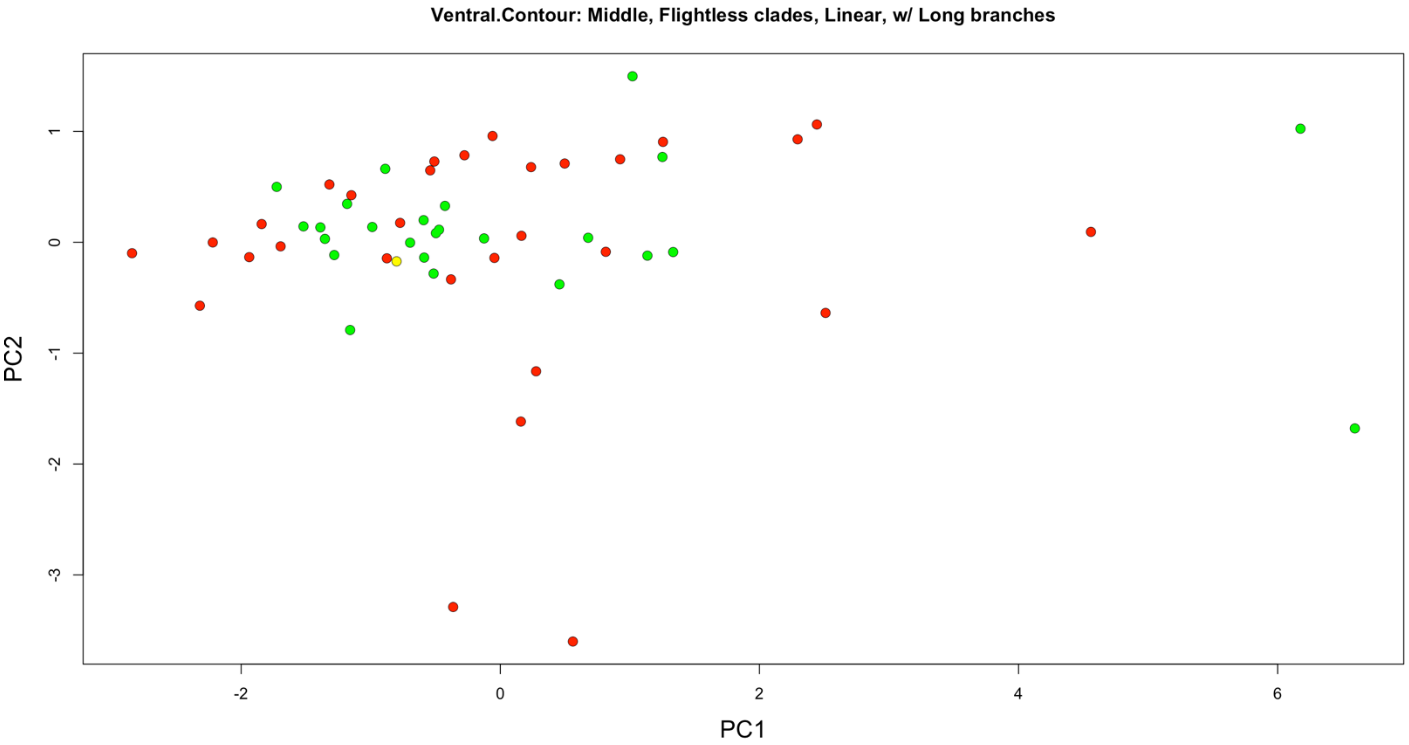


Figure S56. PCA of ventral contours of only clades with flightless taxa, with penguins/tubenoses and ratites/tinamous included. Green are volant taxa, red are flightless taxa, and yellow are poor flying/'incipiently flightless' taxa. N = 56 taxa.

**4.6.2. Ventral contour: no long branches (i.e., without Palaeognathae, Sphenisciformes, and Procellariiformes), PC space established by only clades with flightless taxa**

PC1 and PC2 explain ~70.77% and ~13.60% of the variation, respectively.

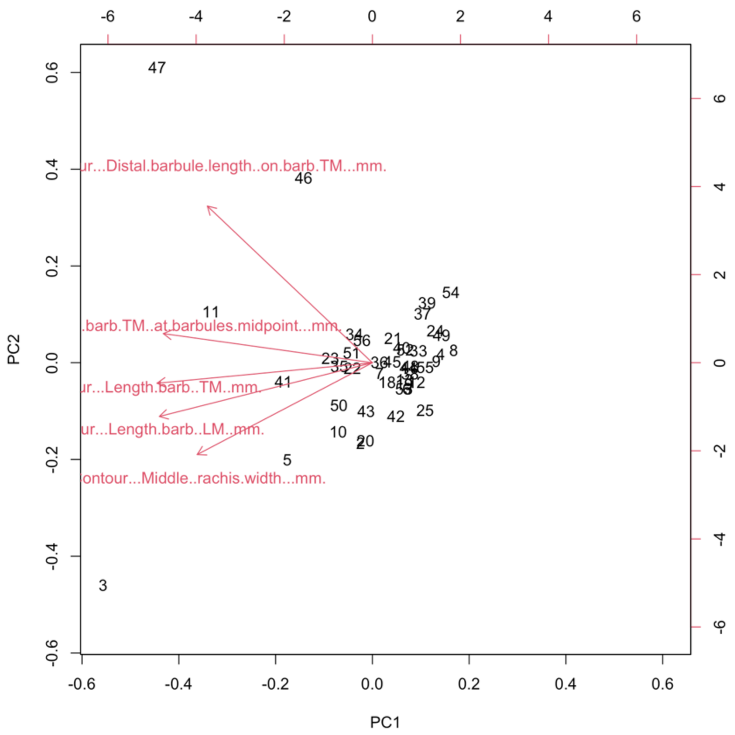


Figure S57. PCA biplot of ventral contours of only clades with flightless taxa, except penguins/tubenoses and ratites/tinamous. Red arrows indicate variable loadings. N = 44 taxa.

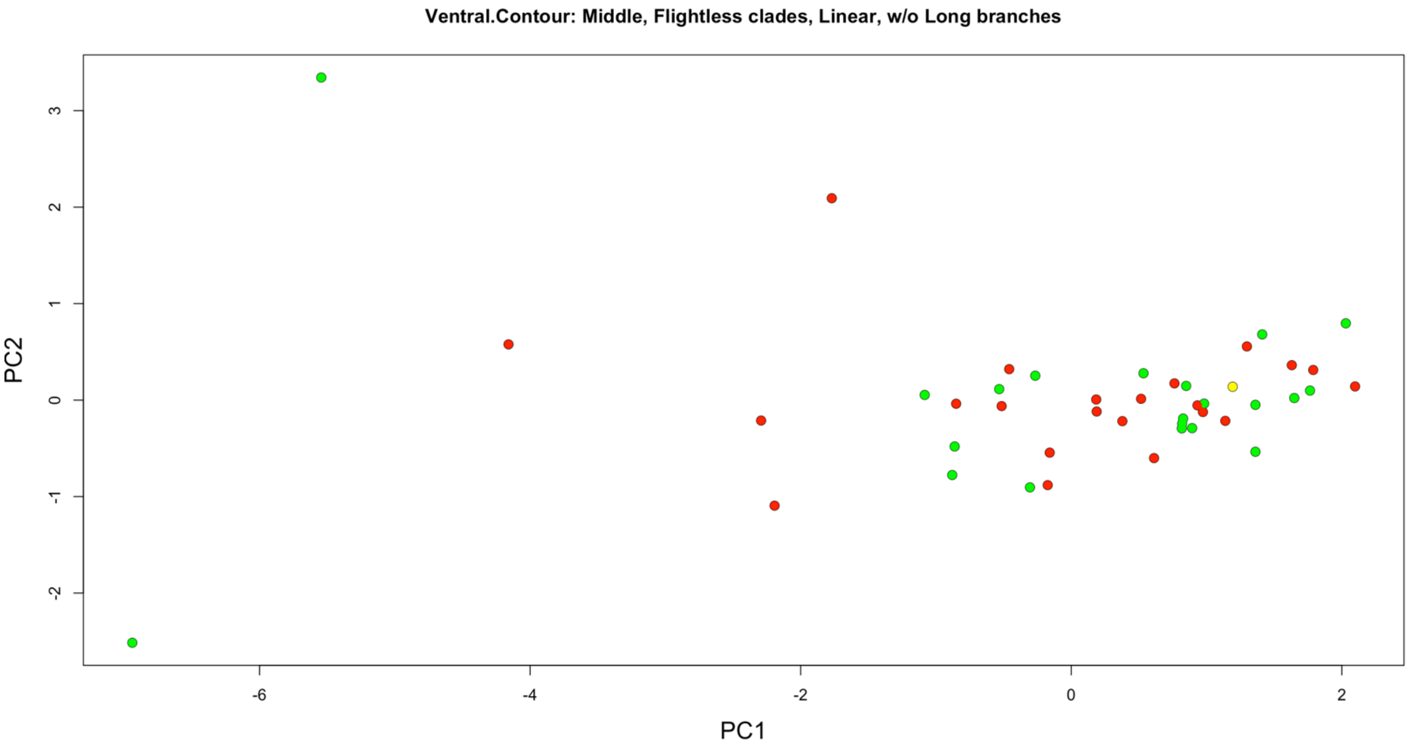


Figure S58. PCA of ventral contours of only clades with flightless taxa, except penguins/tubenoses and ratites/tinamous. Green are volant taxa, red are flightless taxa, and yellow are poor flying/‘incipiently flightless’ taxa. N = 44 taxa.

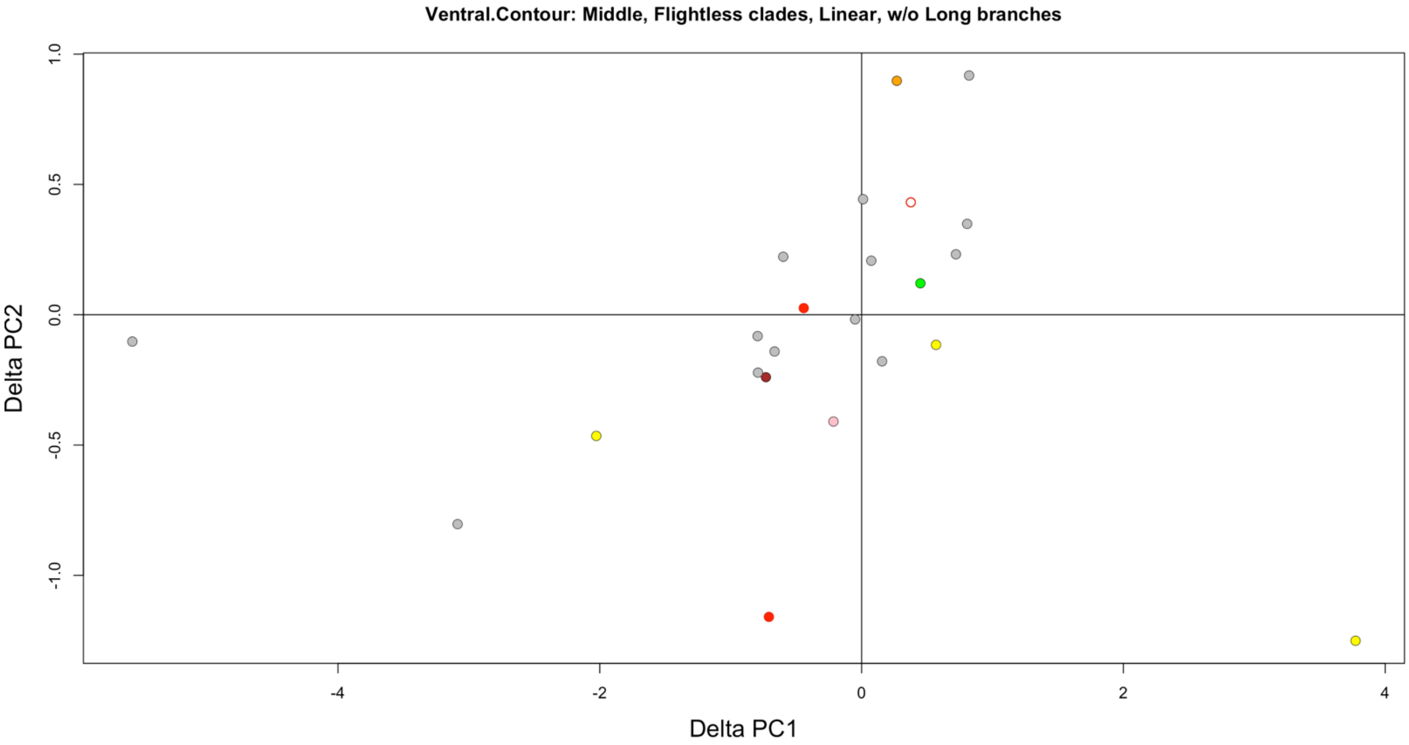


Figure S59. Simple pairwise phylogenetic PCA shift of ventral contours during flight loss based on an initial morphospace of only clades with flightless taxa, except penguins/tubenoses and ratites/tinamous. Color indicates clade as in Figure 3. Open circle is poor flyer/'incipiently flightless'. N = 23 taxa.

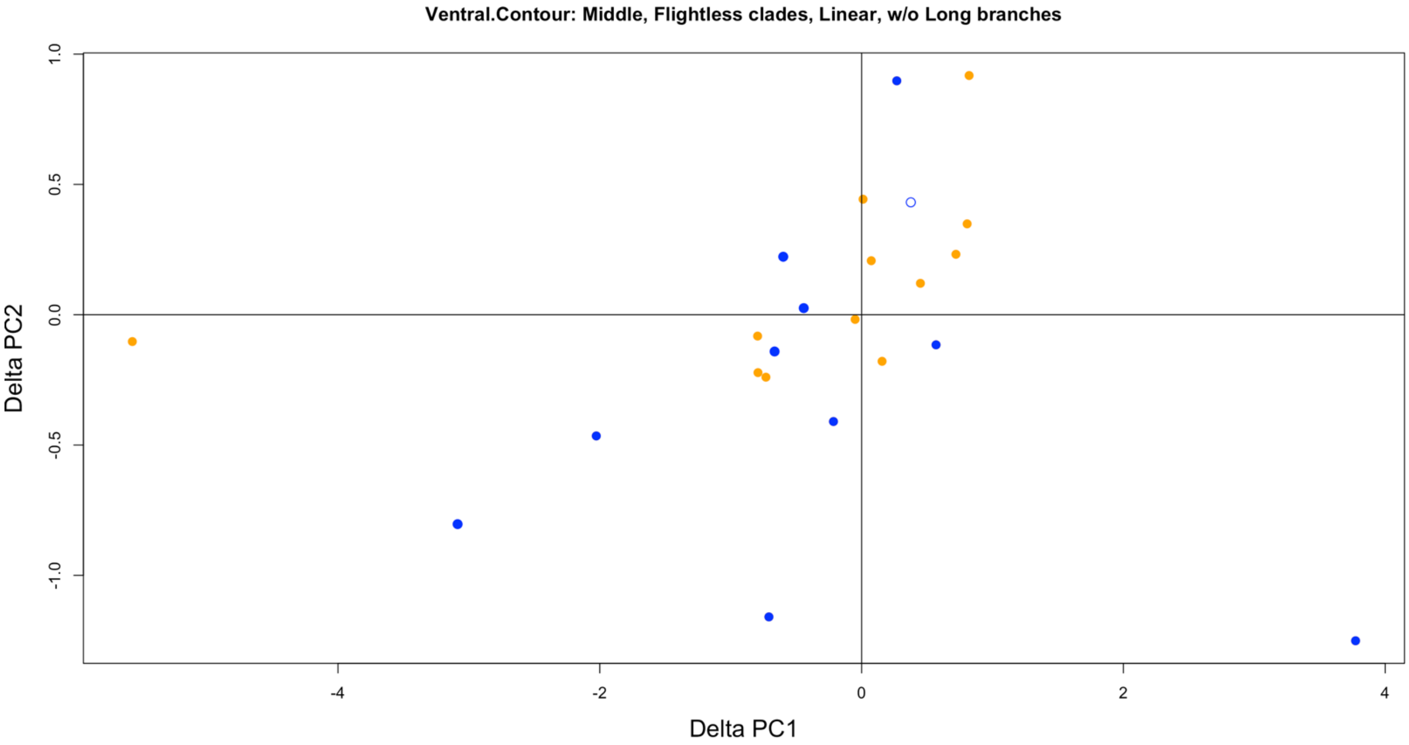


Figure S60. Simple pairwise phylogenetic PCA shift of ventral contours during flight loss based on an initial morphospace of only clades with flightless taxa, except penguins/tubenoses and ratites/tinamous. Blue are semiaquatic, and orange are terrestrial. Open circle is poor flyer/'incipiently flightless'. N = 23 taxa.

### 4.7. Summary plots of simple, pairwise phylogenetically controlled PCA

We ran PCA to establish a morphospace and then added phylogenetic control, such that the shift in PC values can be observed between flightless taxa and their closest volant relative(s), with the origin of the ΔPC plots representing the volant relative(s) (i.e., a hypothetical ancestral state). In combination with the above whole plumage PCA without phylogenetic control, we see that 1) flightless taxa tend to show greater feather morphological diversity than do volant taxa prior to phylogenetic control and 2) barb width tends to intermediately correlate between the filament length metrics and rachis width or that barb width correlates with barbule length (Fig. 2; Figs. S61–S62).

When examining the primaries, upon loss of flight (or incipient loss of flight), there appears to be an initial increase in filament lengths from their volant relatives. Terrestrial flightless taxa, especially some rails, tend to have greater shifts toward longer barbs from their volant relatives than do semiaquatic flightless taxa among more recently flightless clades. Feather types in ratites other than primaries tend to shift toward filament length reduction, rather than the more diverse changes seen in their primaries. Semiaquatic flightless taxa tend to shift their primaries more towards wider filaments and longer barbules from their volant relatives than do terrestrial flightless taxa among more recently flightless clades.

These patterns appear to be strongest in primaries, followed by the rectrix, tertial, dorsal contour, and ventral contour. In other words, clade and ecological categories begin to center around the ΔPC plot origin and the difference between flightless and volant taxa feather morphological diversity prior to phylogenetic control seems to decrease when simple phylogenetic control is added. Our results are generally supported when repeating the PCA under varying initial conditions (i.e., with and without 'long branch' clades where flightlessness occurred 10's of millions of years ago, as well as with and without clades that don't include flightless taxa in the case of the primaries).

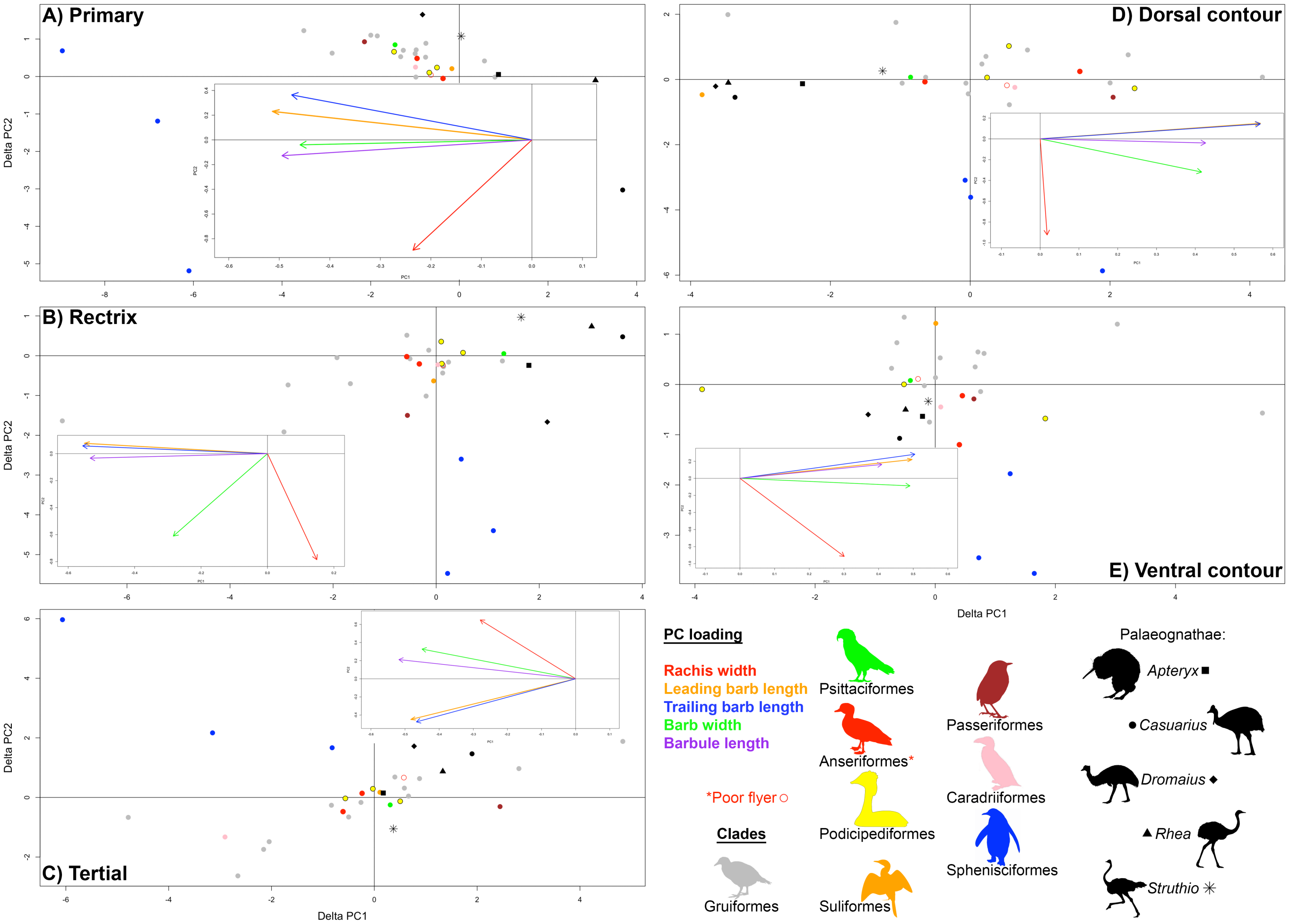


Figure S61. Phylogenetically controlled shifts in feather morphospace of flightless and 'incipiently flightless' taxa from their closest volant relative(s) in, A, primaries, B, rectrices, C, tertials, D, dorsal contours, and, E, ventral contours according to clade. Initial PCA morphospace prior to phylogenetic control established using only clades with flightless taxa for all panels. Insets show the loading plots for the five linear metrics on the PC1 and PC2. Each of the five metrics is divided by the length of the rachis from the apical-most node to the middle barb pair measured here. Ratite point shape indicates genus (bearing in mind that the semi-alternating barb development pattern in *Apteryx* could shift its position either towards longer filament ratites like *Struthio* and *Dromaius* or towards shorter filament ratites like *Rhea* and *Casuarius*, if using the example of the primary). *The red open circle of the Anseriformes represents the poor-flying/'incipiently flightless' *Mergus australis*. N = 30 flightless and 1 'incipiently flightless' taxa (initial morphospace established by 56 taxa).

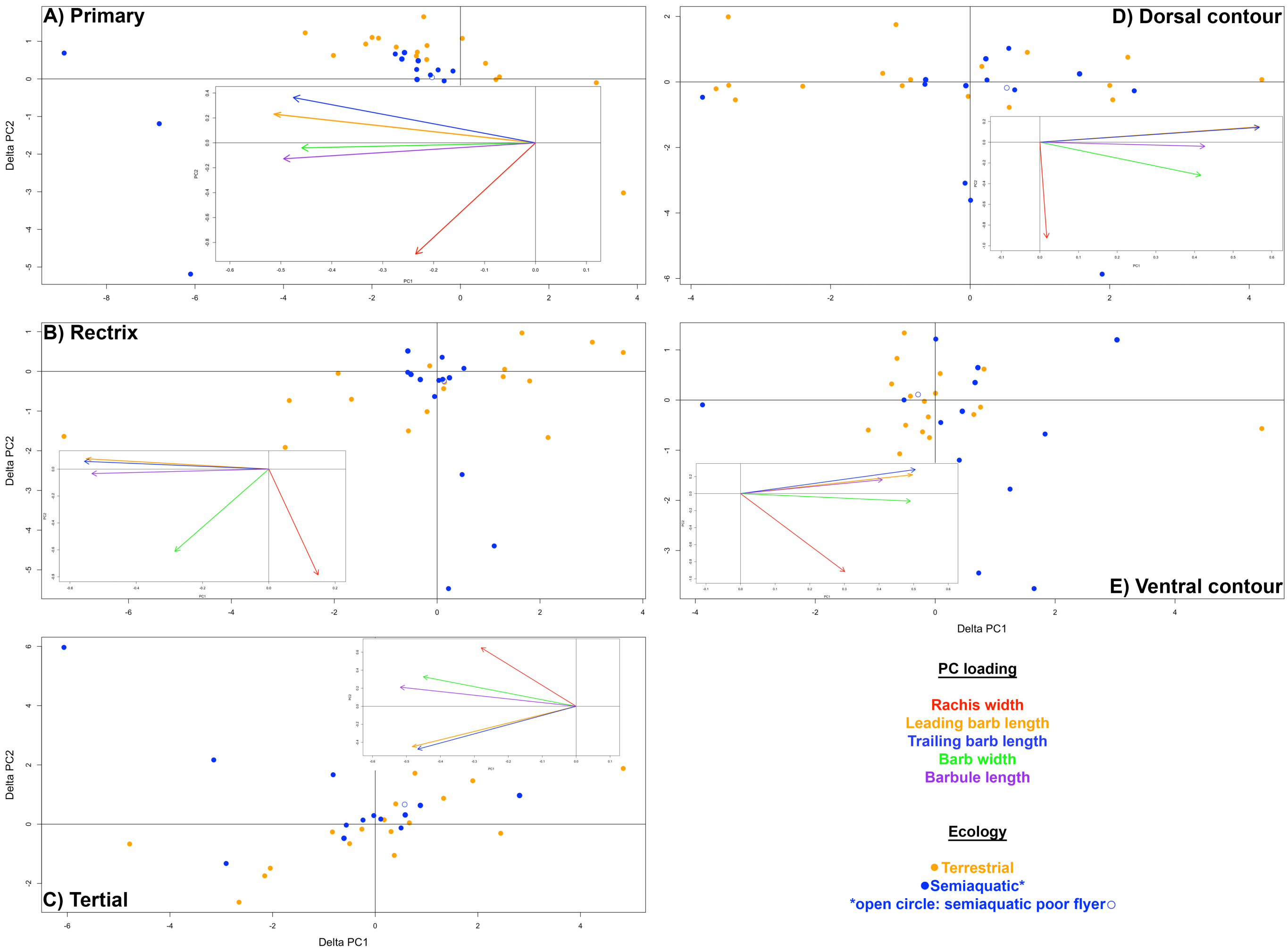


Figure S62. Phylogenetically controlled shifts in feather morphospace of flightless and 'incipiently flightless' taxa from their closest volant relative(s) in, A, primaries, B, rectrices, C, tertials, D, dorsal contours, and, E, ventral contours according to ecology. Initial PCA morphospace prior to phylogenetic control established using only clades with flightless taxa for all panels. Insets show the loading plots for the five linear metrics on the PC1 and PC2. Each of the five metrics is divided by the length of the rachis from the apical-most node to the middle barb pair measured here. *The blue open circle of the semiaquatic taxa represents the poor-flying/'incipiently flightless' *Mergus australis*. N = 30 flightless and 1 'incipiently flightless' taxa (initial morphospace established by 56 taxa).

When simple phylogenetic controls are introduced and the shift from volant relative(s) to flightless taxa in the morphospaces are observed, the most prevalent patterns according to clade and ecology seem to occur in the primaries. This is consistent with the primary's crucial role in flight, as well as its occasional use in swimming in some taxa. As observed in the primaries, there appears to possibly be an initial increase in filament lengths upon the loss of flight, possibly indicating less robust feathers that are under less selective pressure for use in flight and perhaps for increased thermoregulation.

Overall, semiaquatic flightless taxa tend to have shifts directed towards wider filaments and longer barbules compared to the terrestrial flightless taxa, hypothetically consistent with locomotion through a dense water medium.

Among taxa that lost flight relatively recently, terrestrial flightless taxa tend to shift towards longer filaments than semiaquatic flightless taxa.

### 5. Asymmetry: middle & basal regions (see Fig. 3)

#### 5.1. Tertial asymmetry, middle & basal regions

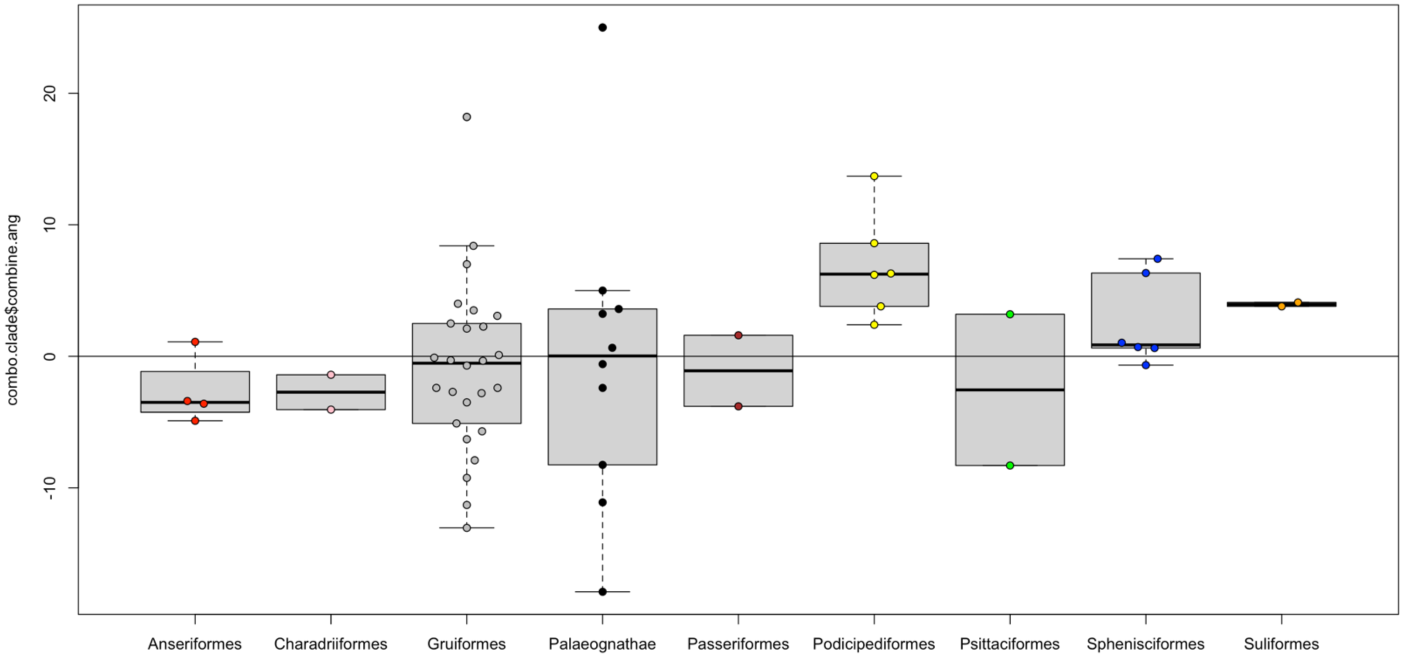


Figure S63. Pairwise phylogenetic shifts in barb asymmetry values due to flight loss of tertials (middle & basal region combined) according to clade. N = 60 barb pairs (middle & basal region), 30 taxa.

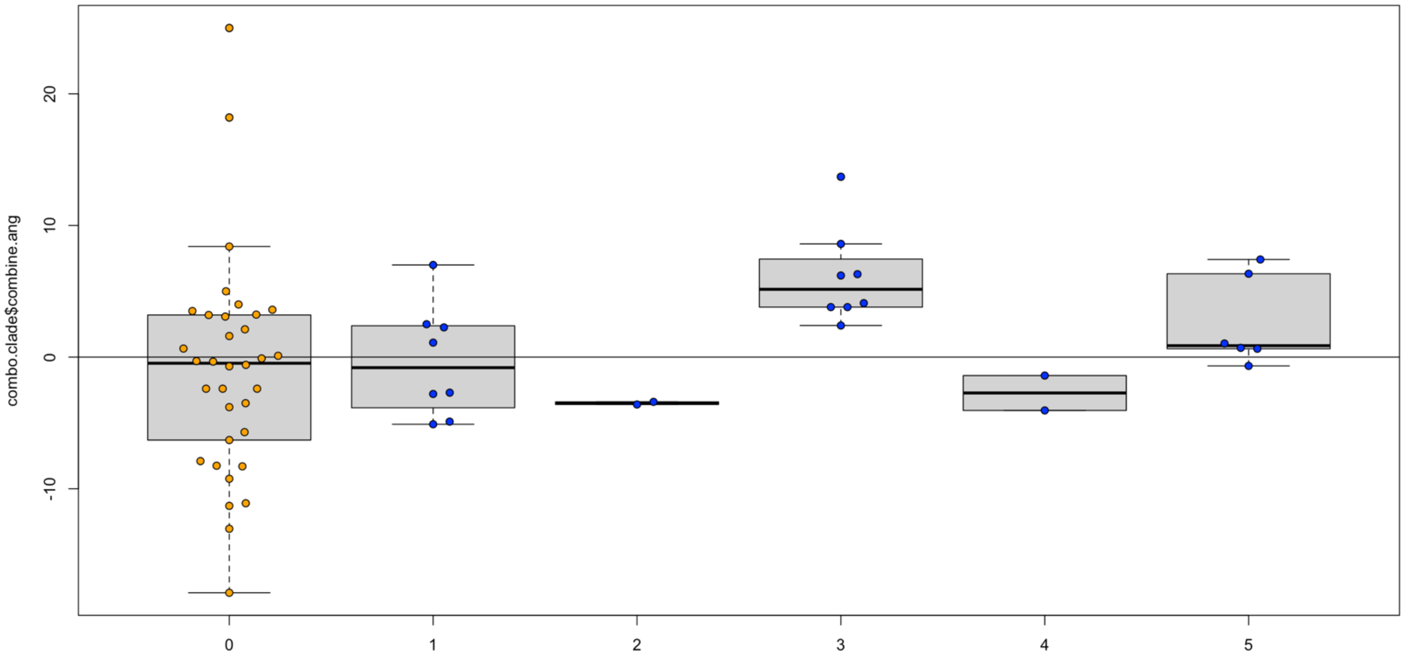

Figure S64. Pairwise phylogenetic shifts in barb asymmetry values due to flight loss of tertials (middle & basal region combined) according to swim score. Blue are semiaquatic, and orange are terrestrial. N = 60 barb pairs (middle & basal region), 30 taxa.

### 5.2. Rectrix asymmetry, middle & basal regions

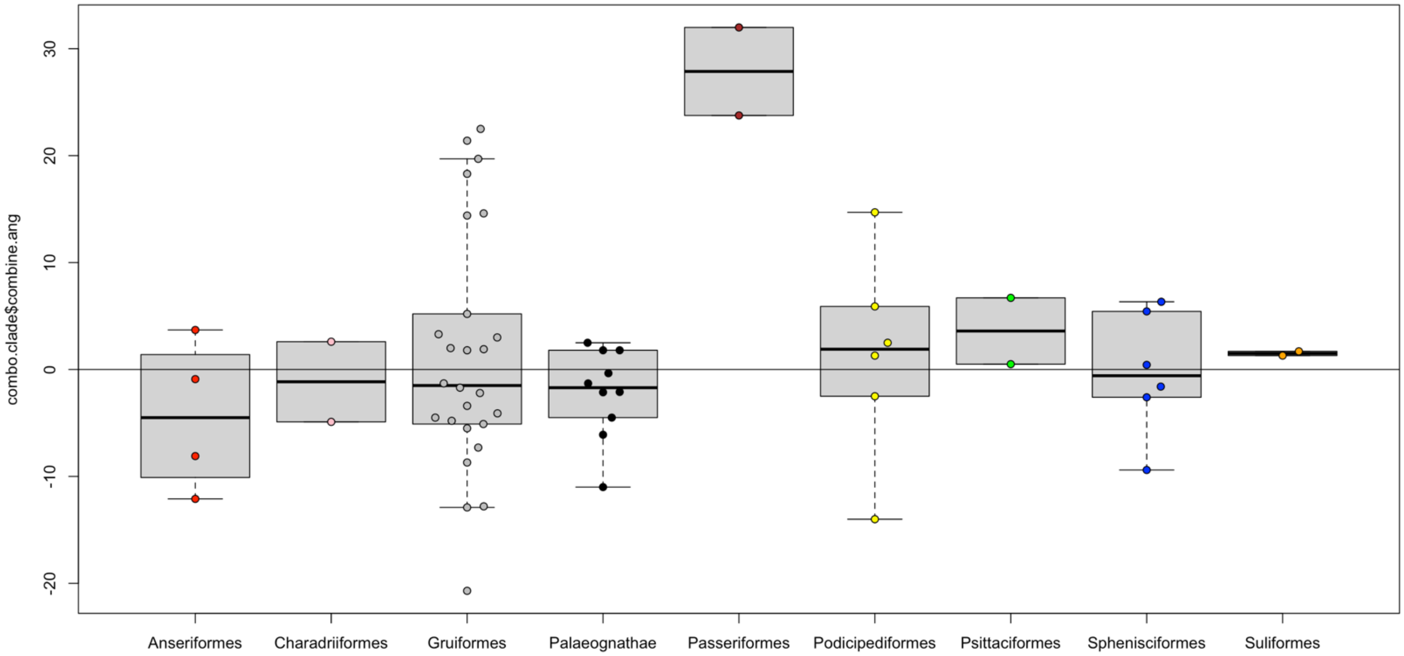


Figure S65. Pairwise phylogenetic shifts in barb asymmetry values due to flight loss of rectrices (middle & basal region combined) according to clade. N = 60 barb pairs (middle & basal region), 30 taxa.

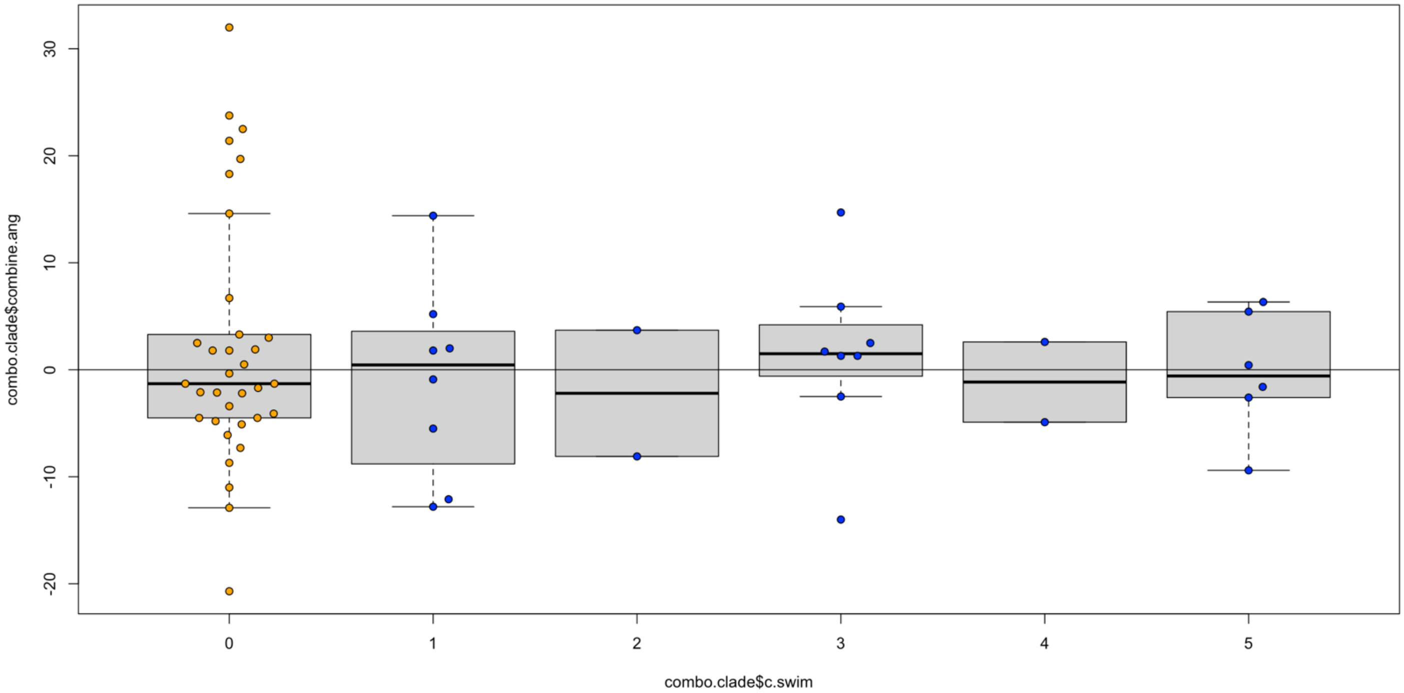


Figure S66. Pairwise phylogenetic shifts in barb asymmetry values due to flight loss of rectrices (middle & basal region combined) according to swim score. Blue are semiaquatic, and orange are terrestrial. N = 60 barb pairs (middle & basal region), 30 taxa.

#### 5.3. Dorsal contour asymmetry, middle & basal regions

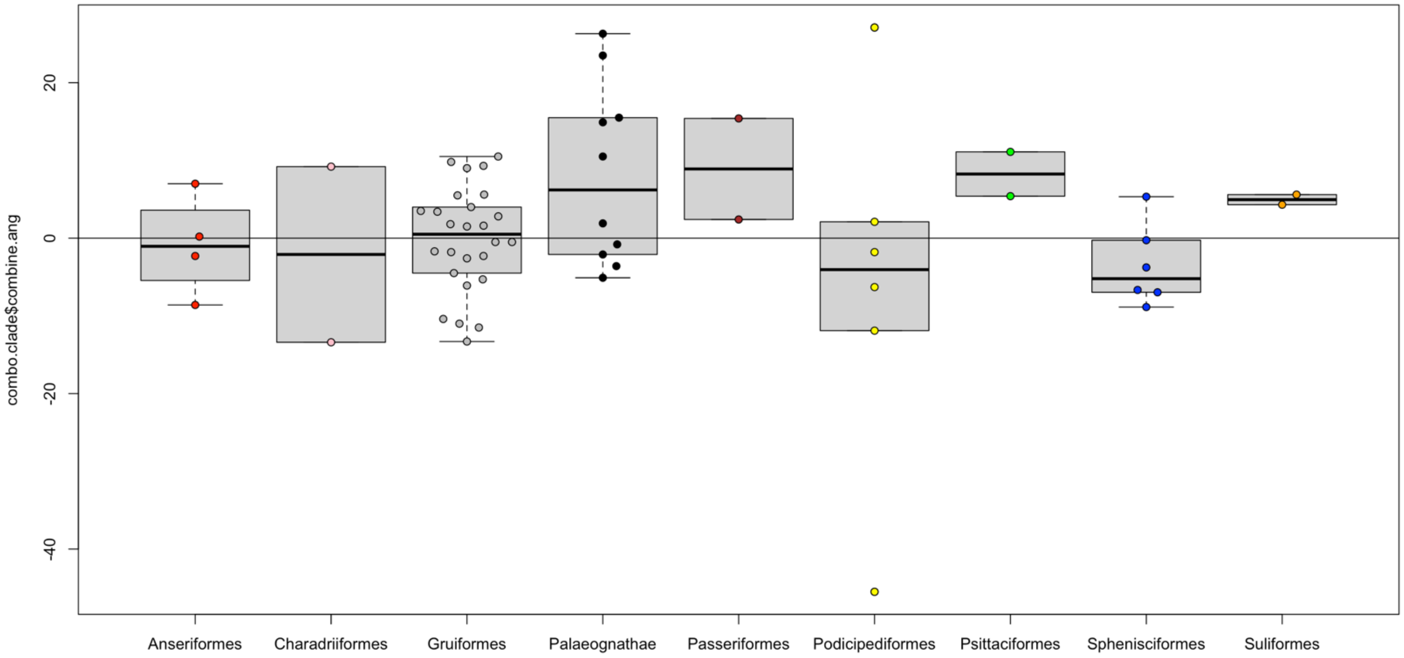


Figure S67. Pairwise phylogenetic shifts in barb asymmetry values due to flight loss of dorsal contours (middle & basal region combined) according to clade. N = 60 barb pairs (middle & basal region), 30 taxa.

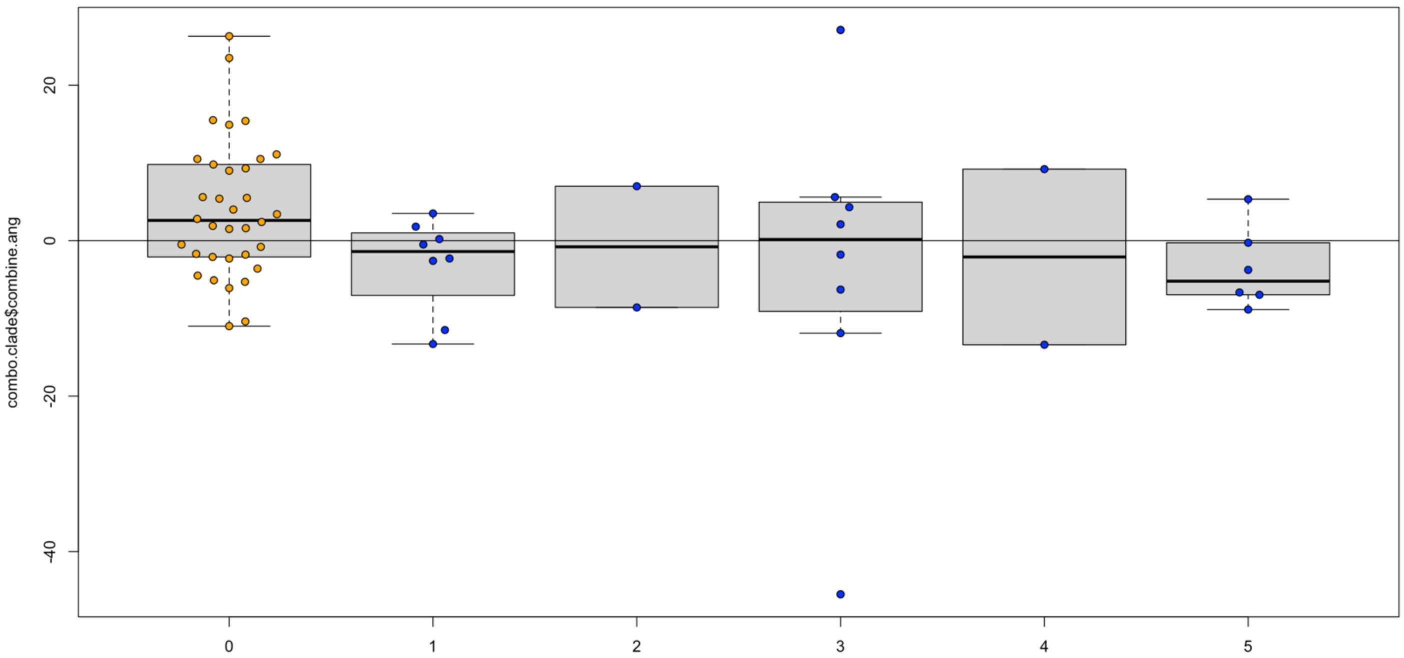


Figure S68. Pairwise phylogenetic shifts in barb asymmetry values due to flight loss of dorsal contours (middle & basal region combined) according to swim score. Blue are semiaquatic, and orange are terrestrial. N = 60 barb pairs (middle & basal region), 30 taxa.

### 5.4. Ventral contour asymmetry, middle & basal regions

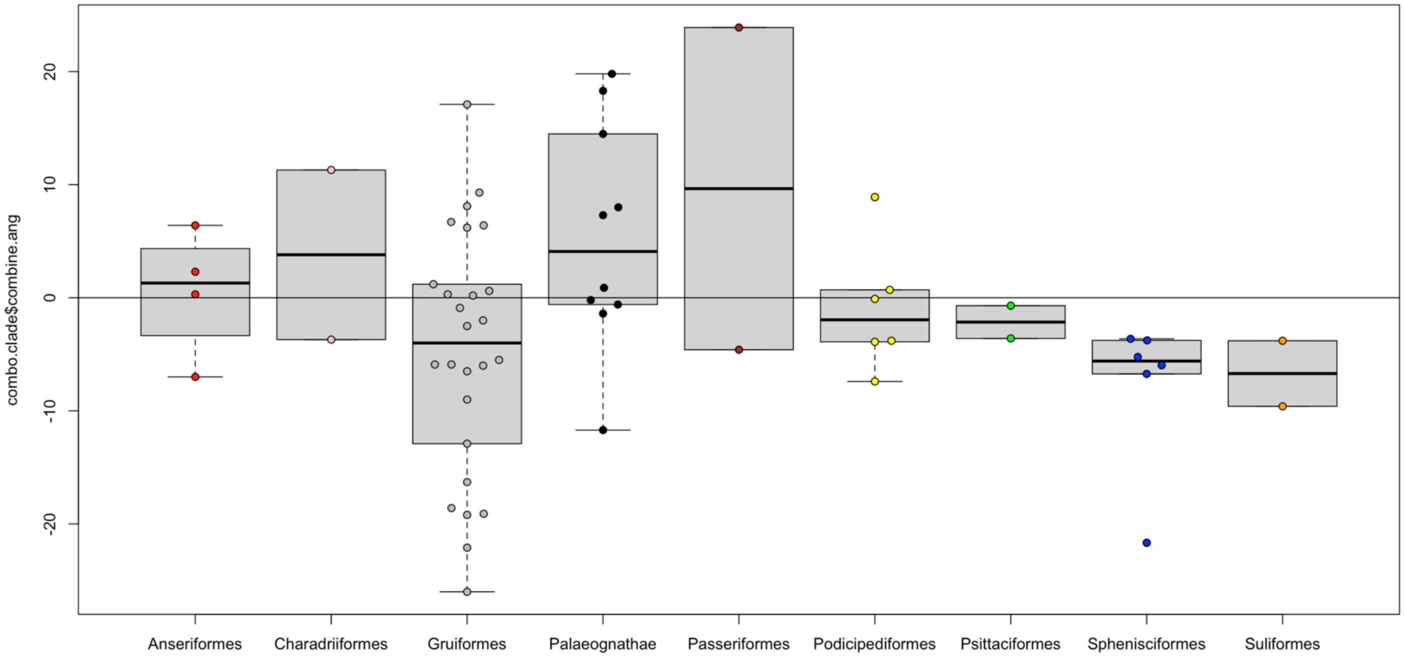


Figure S69. Pairwise phylogenetic shifts in barb asymmetry values due to flight loss of ventral contours (middle & basal region combined) according to clade. N = 60 barb pairs (middle & basal region), 30 taxa.

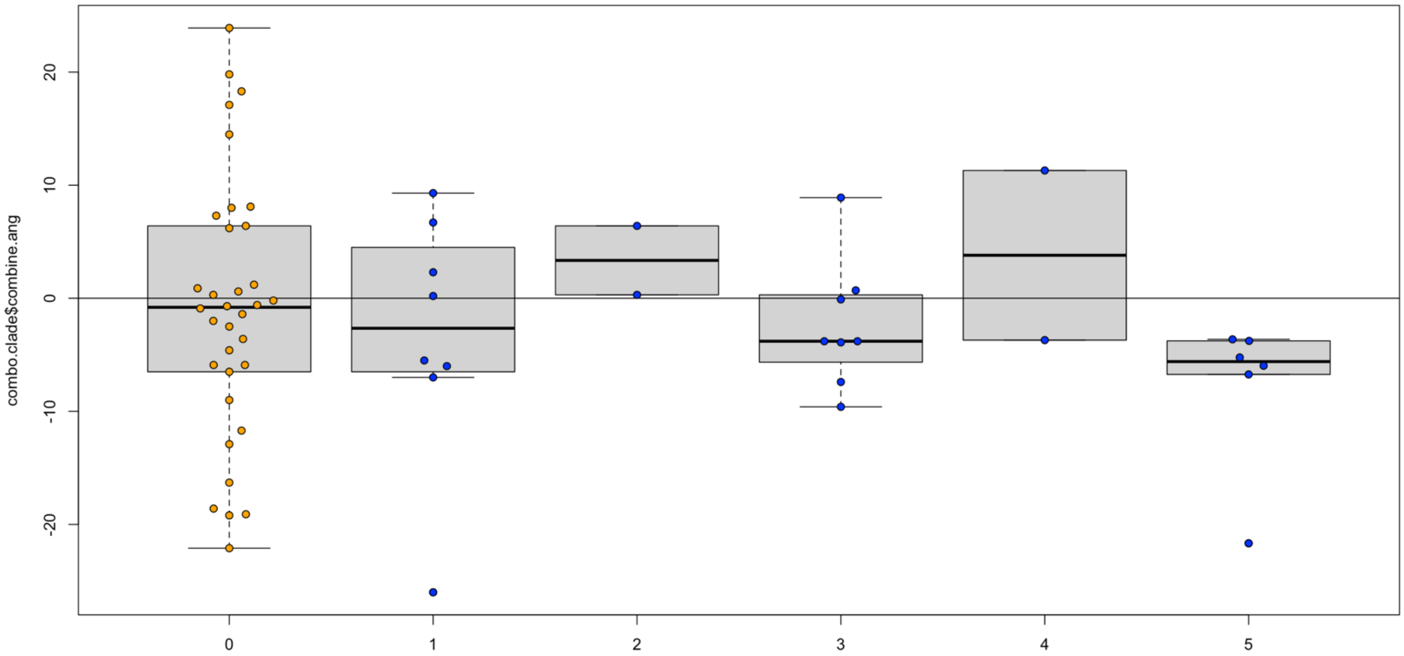


Figure S70. Pairwise phylogenetic shifts in barb asymmetry values due to flight loss of ventral contours (middle & basal region combined) according to swim score. Blue are semiaquatic, and orange are terrestrial. N = 60 barb pairs (middle & basal region), 30 taxa.

### 6. Filament densities: leading/trailing barbs, distal/proximal barbules, middle & basal regions (see Fig. 4)

#### 6.1. Tertial densities, leading/trailing barbs, distal/proximal barbules, middle & basal regions

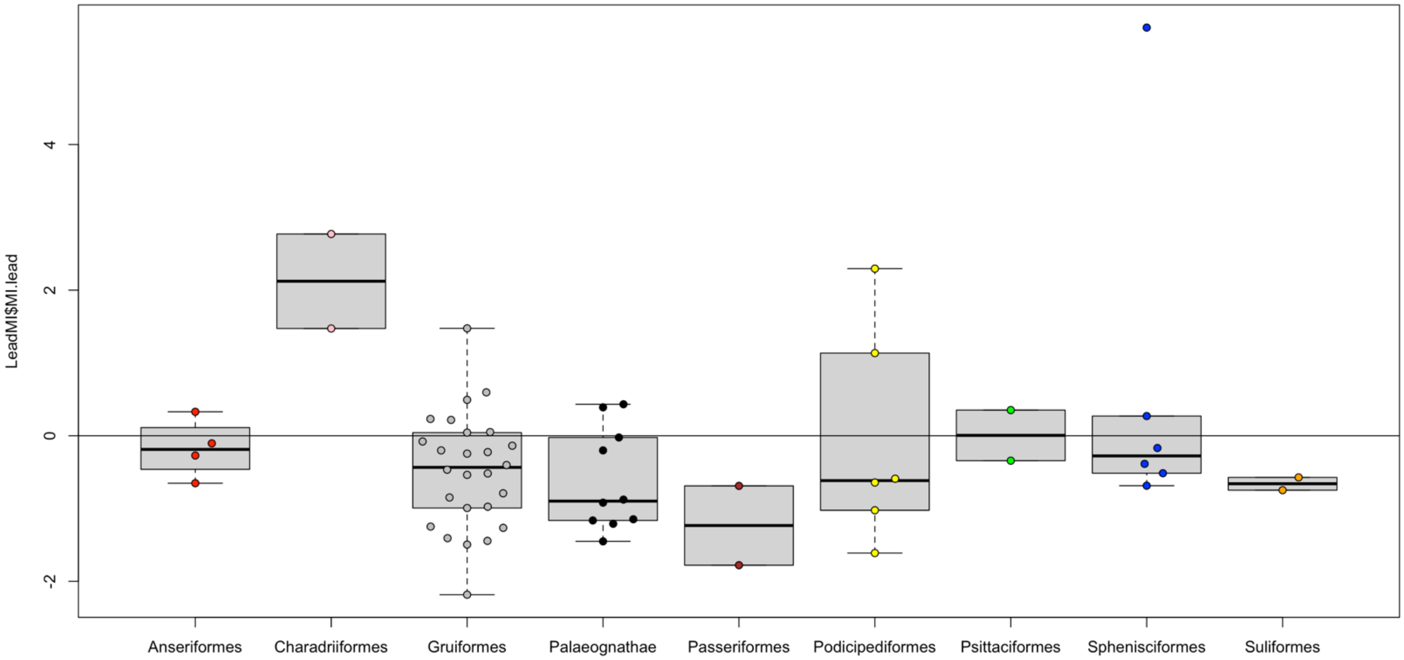


Figure S71. Pairwise phylogenetic shifts in leading barb densities due to flight loss of tertials (middle & basal region combined) according to clade. N = 60 leading barb density measurements (middle & basal region), 30 taxa.

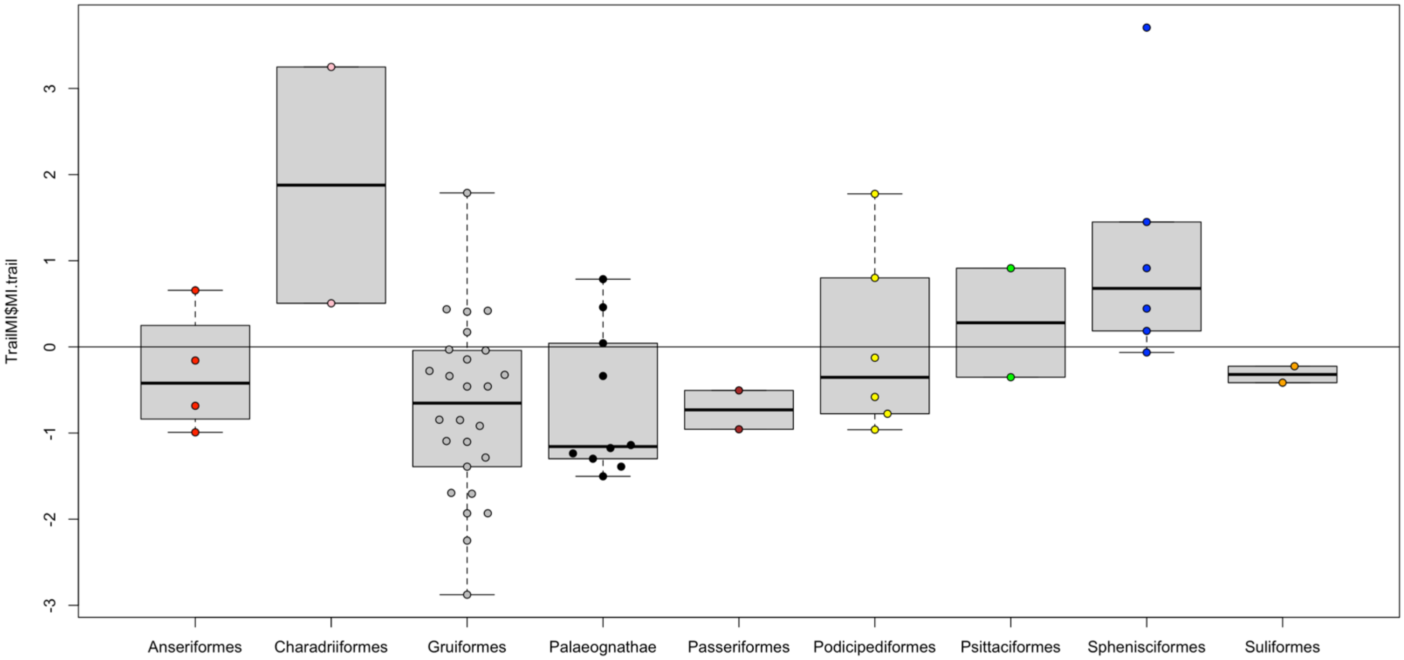


Figure S72. Pairwise phylogenetic shifts in trailing barb densities due to flight loss of tertials (middle & basal region combined) according to clade. N = 60 trailing barb density measurements (middle & basal region), 30 taxa.

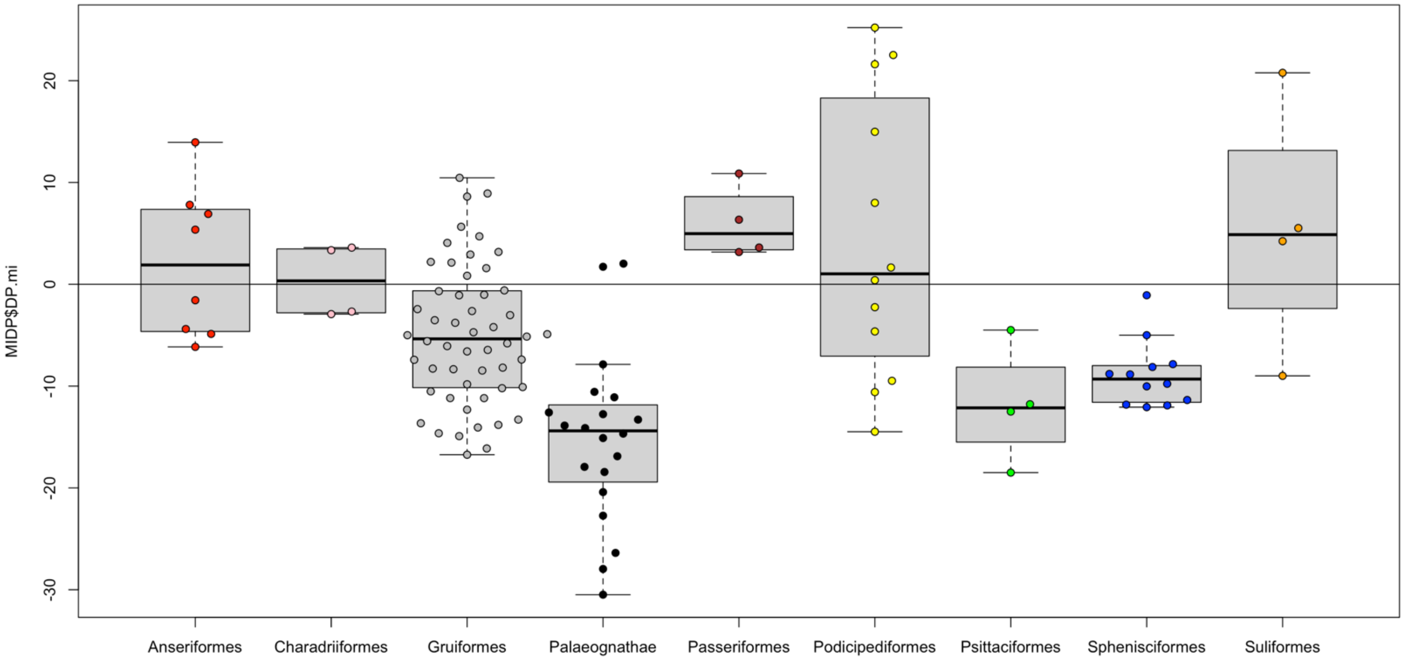


Figure S73. Pairwise phylogenetic shifts in trailing barbule densities (distal & proximal combined) due to flight loss of tertials (middle & basal region combined) according to clade. N = 120 barbule density measurements (distal & proximal), 60 trailing barbs (middle & basal region), 30 taxa.

### 6.2. Rectrix densities, leading/trailing barbs, distal/proximal barbules, middle & basal regions

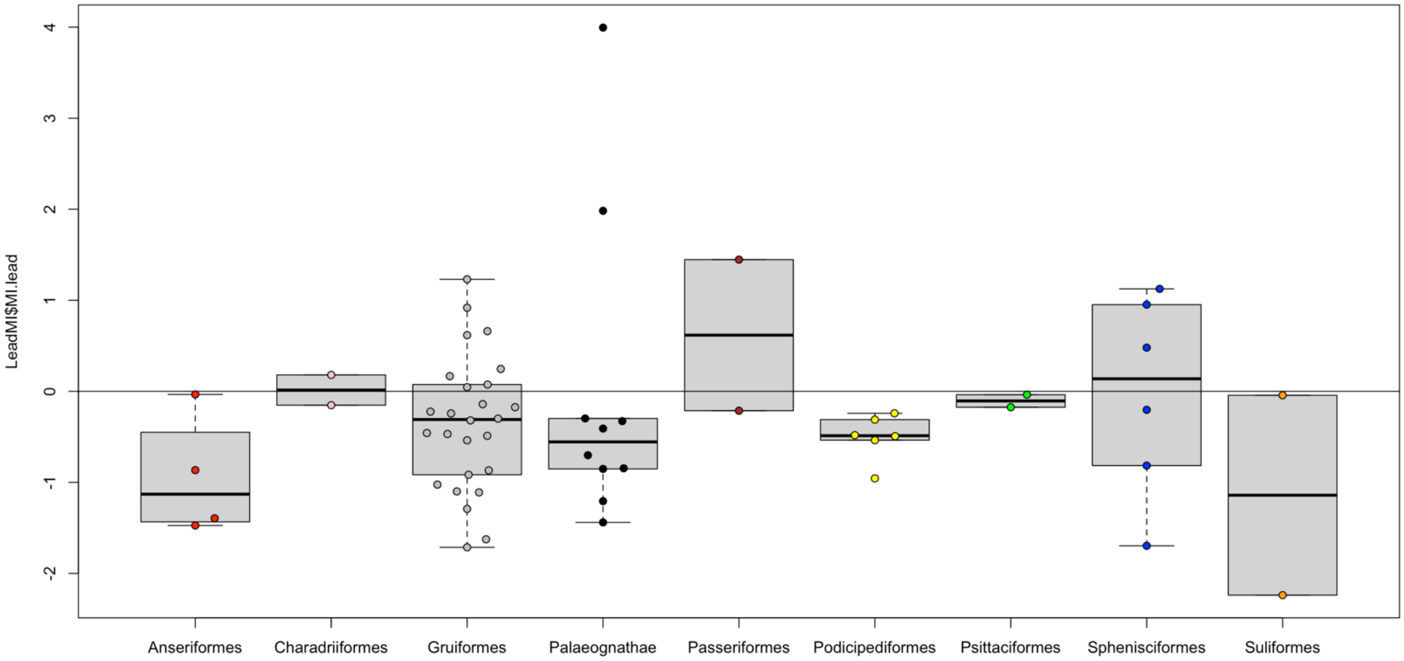


Figure S74. Pairwise phylogenetic shifts in leading barb densities due to flight loss of rectrices (middle & basal region combined) according to clade. N = 60 leading barb density measurements (middle & basal region), 30 taxa.

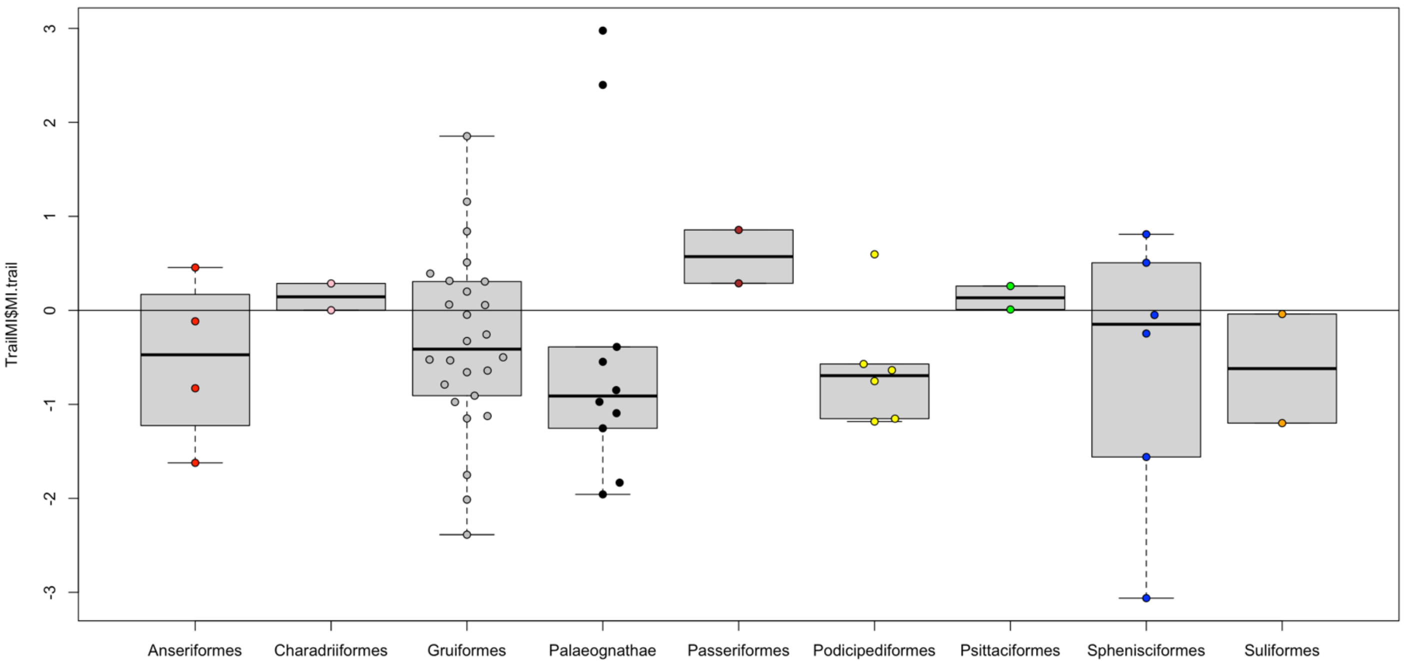

Figure S75. Pairwise phylogenetic shifts in trailing barb densities due to flight loss of rectrices (middle & basal region combined) according to clade. N = 60 trailing barb density measurements (middle & basal region), 30 taxa.

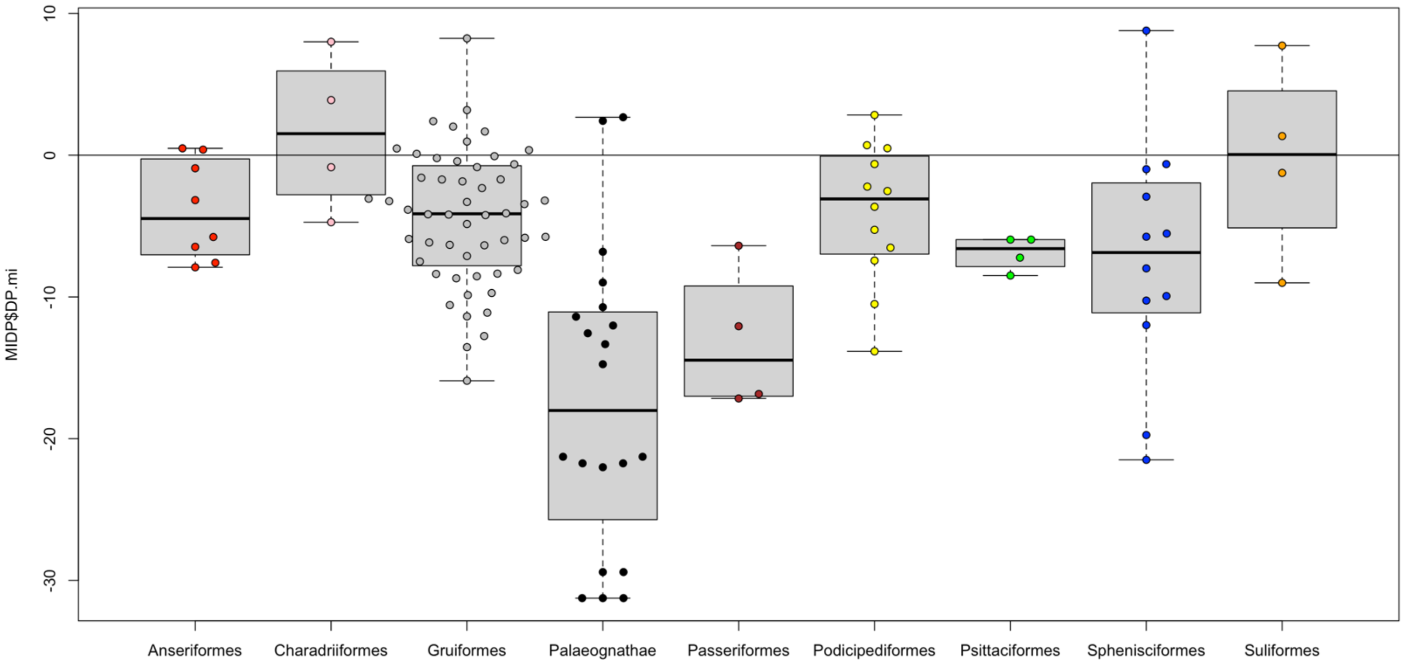

Figure S76. Pairwise phylogenetic shifts in trailing barbule densities (distal & proximal combined) due to flight loss of rectrices (middle & basal region combined) according to clade. N = 120 barbule density measurements (distal & proximal), 60 trailing barbs (middle & basal region), 30 taxa.

#### 6.3. Dorsal contour densities, leading/trailing barbs, distal/proximal barbules, middle & basal regions

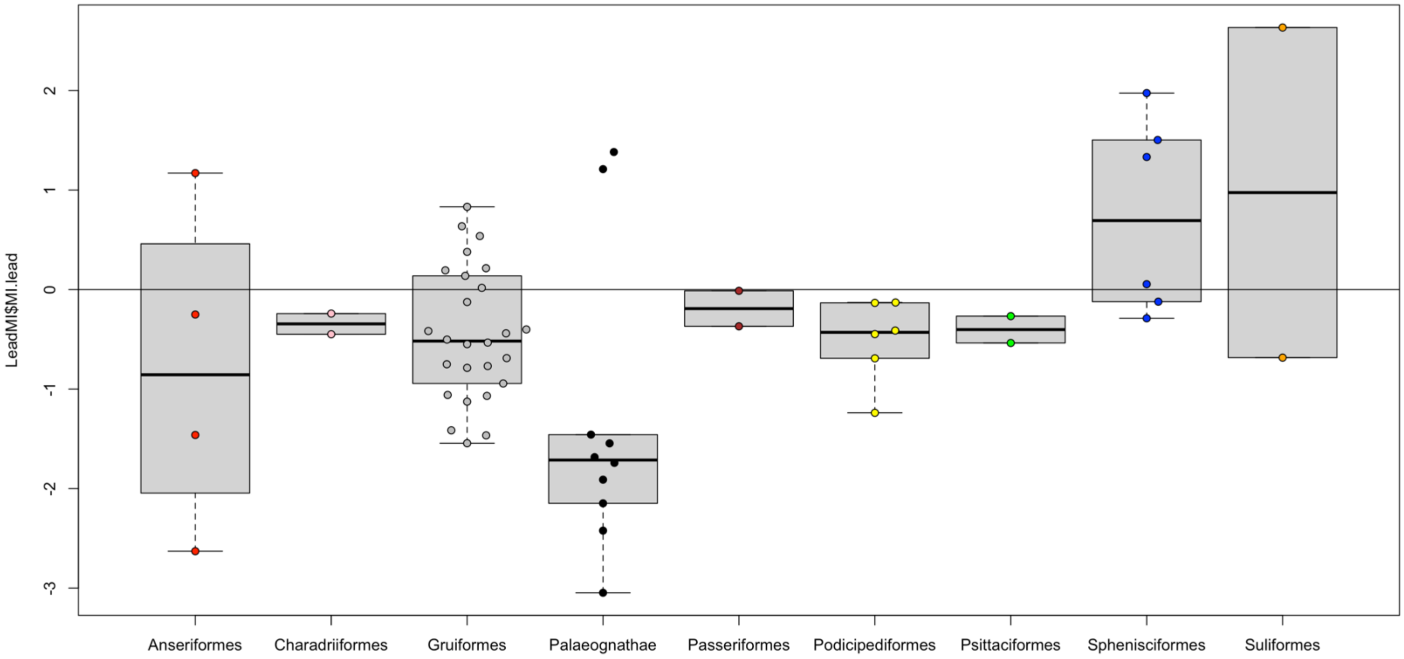


Figure S77. Pairwise phylogenetic shifts in leading barb densities due to flight loss of dorsal contours (middle & basal region combined) according to clade. N = 60 leading barb density measurements (middle & basal region), 30 taxa.

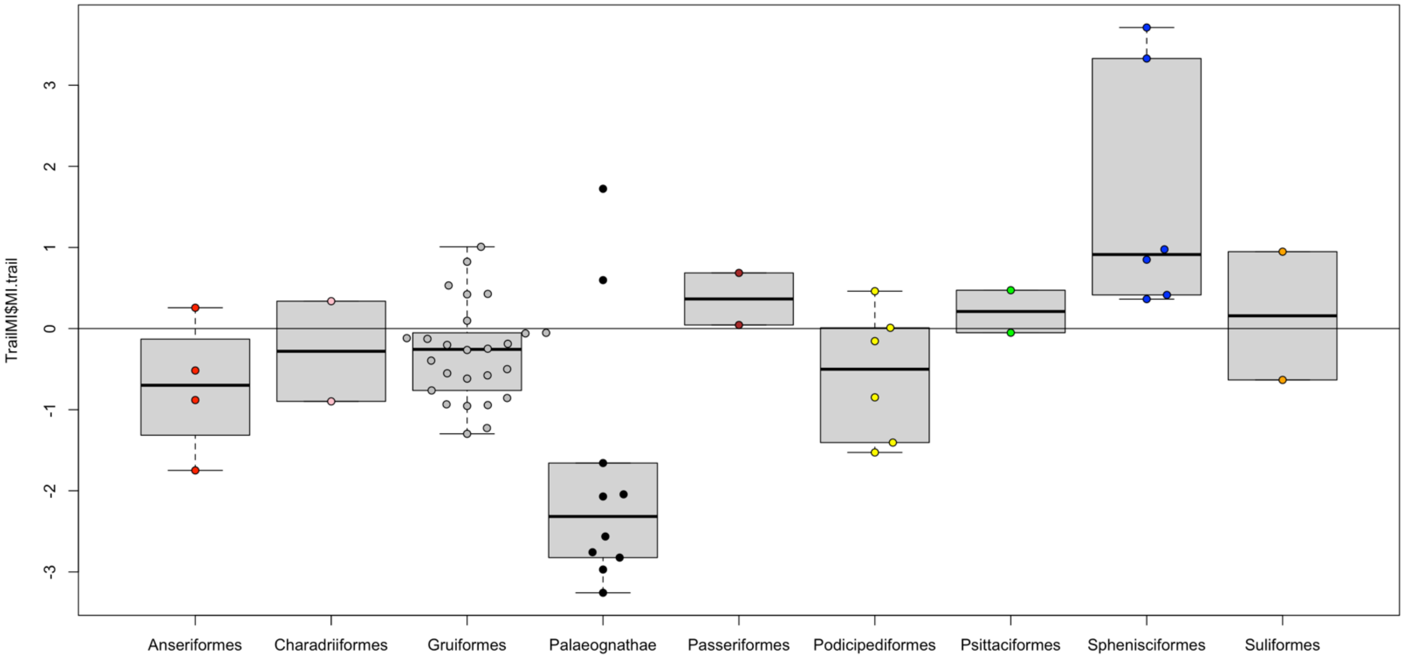


Figure S78. Pairwise phylogenetic shifts in trailing barb densities due to flight loss of dorsal contours (middle & basal region combined) according to clade. N = 60 trailing barb density measurements (middle & basal region), 30 taxa.

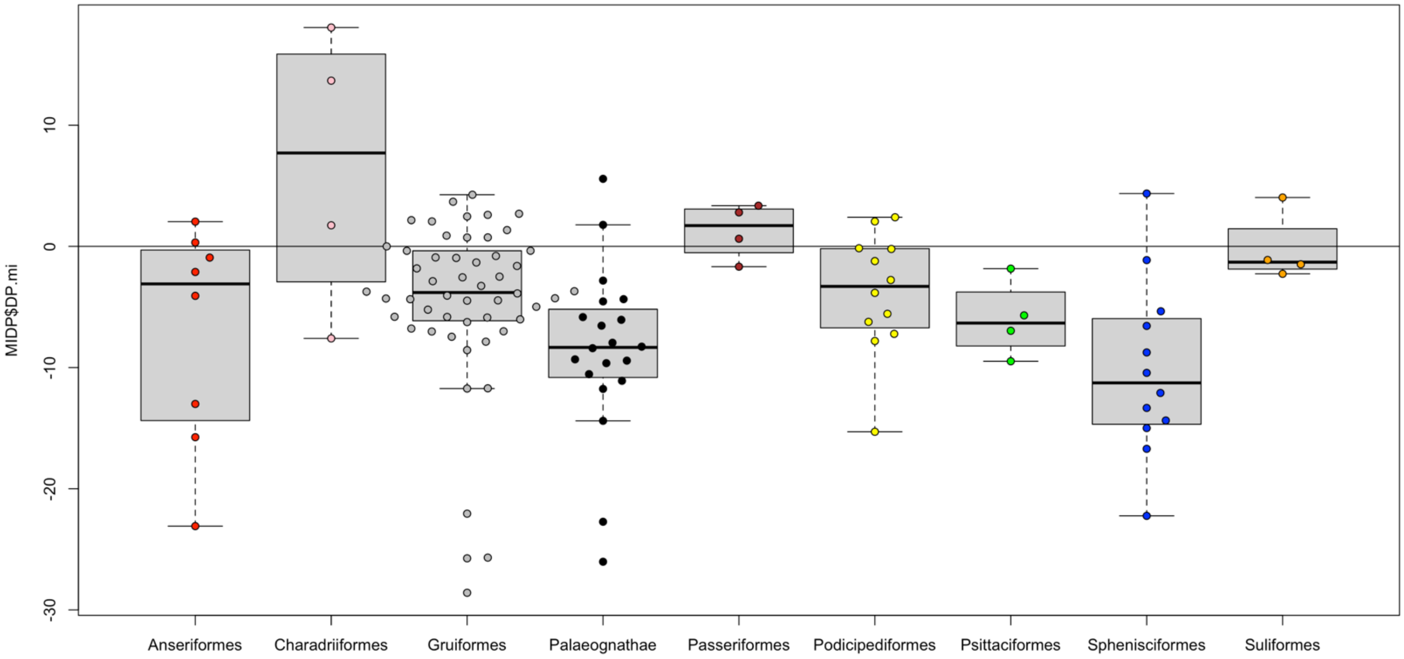


Figure S79. Pairwise phylogenetic shifts in trailing barbule densities (distal & proximal combined) due to flight loss of dorsal contours (middle & basal region combined) according to clade. N = 120 barbule density measurements (distal & proximal), 60 trailing barbs (middle & basal region), 30 taxa.

#### 6.4. Ventral contour densities, leading/trailing barbs, distal/proximal barbules, middle & basal regions

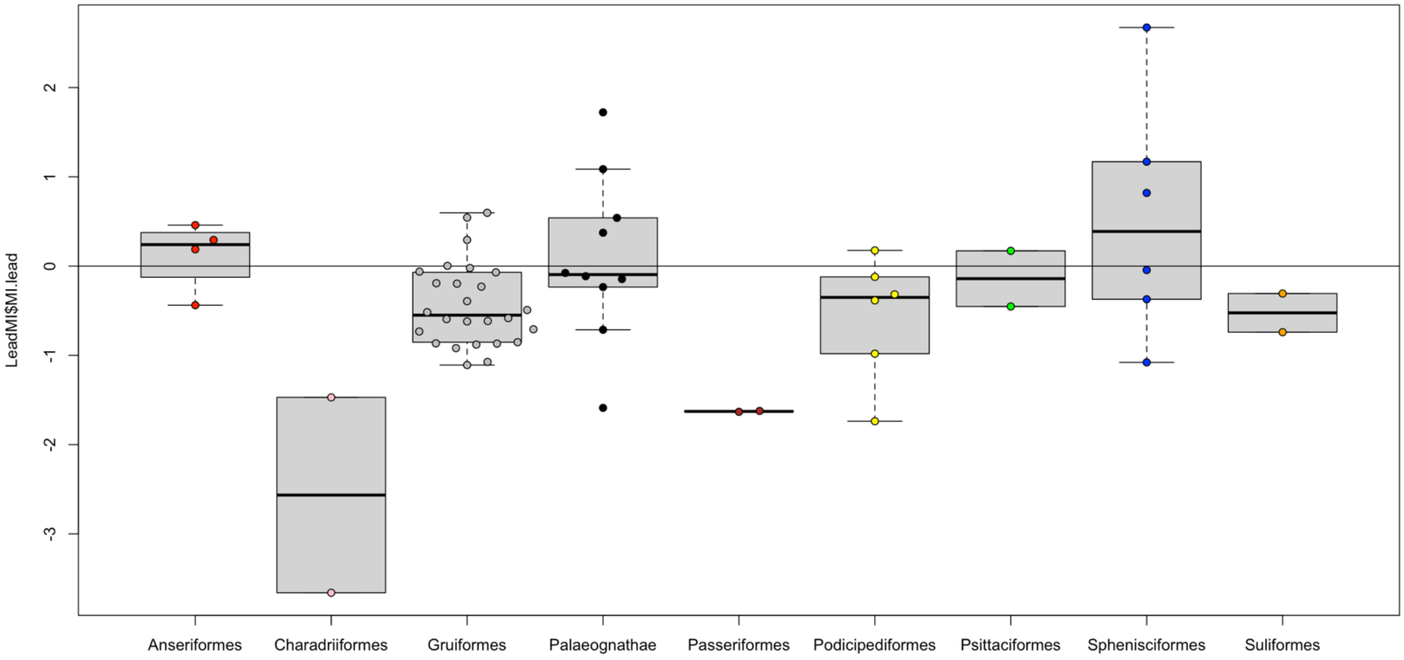


Figure S80. Pairwise phylogenetic shifts in leading barb densities due to flight loss of ventral contours (middle & basal region combined) according to clade. N = 60 leading barb density measurements (middle & basal region), 30 taxa.

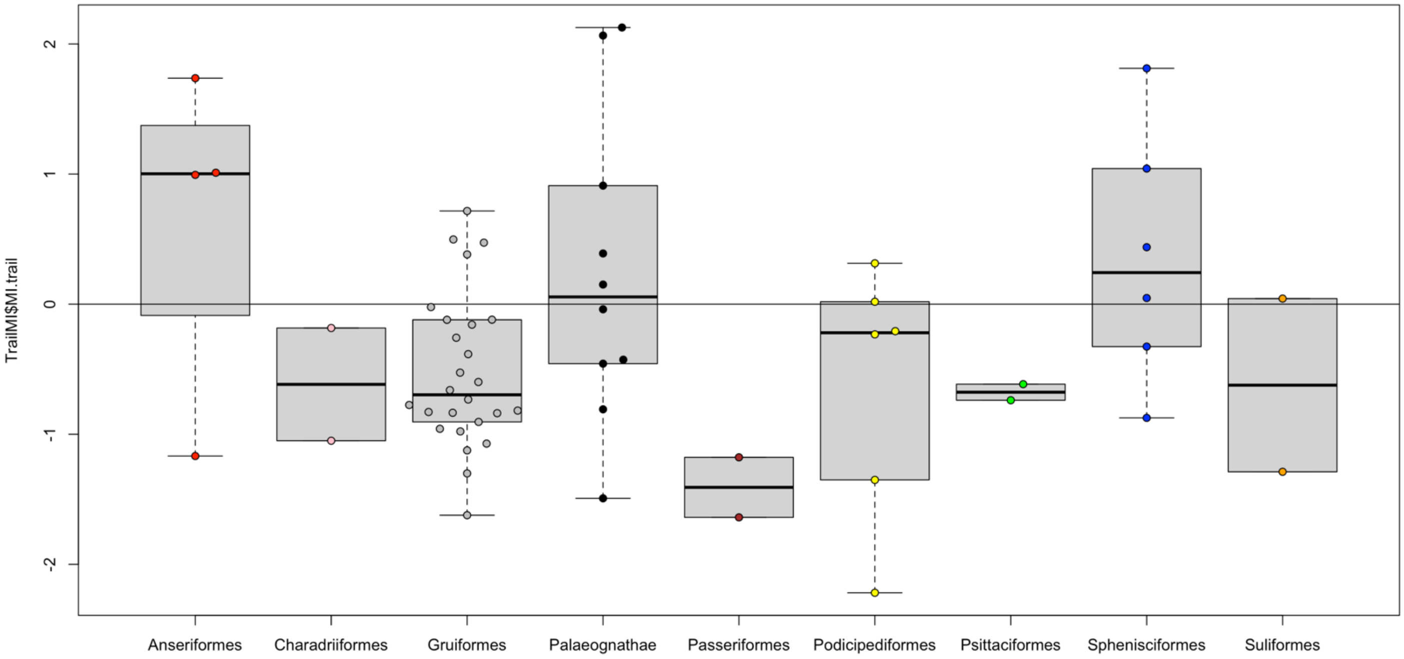

Figure S81. Pairwise phylogenetic shifts in trailing barb densities due to flight loss of ventral contours (middle & basal region combined) according to clade. N = 60 trailing barb density measurements (middle & basal region), 30 taxa.

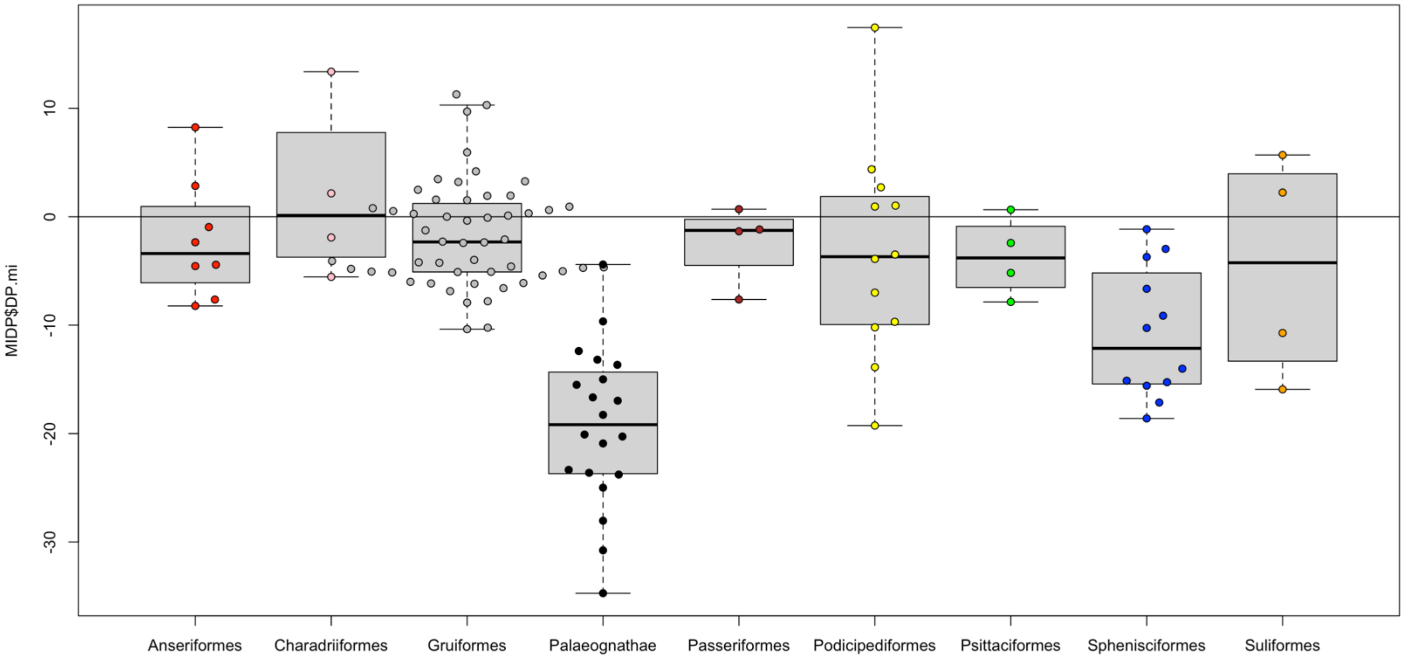

Figure S82. Pairwise phylogenetic shifts in trailing barbule densities (distal & proximal combined) due to flight loss of ventral contours (middle & basal region combined) according to clade. N = 120 barbule density measurements (distal & proximal), 60 trailing barbs (middle & basal region), 30 taxa.

### 7. Simple, phylogenetic pairwise regressions on macro- and microscopic traits

We report simplistic regression analyses to examine morphological divergence (Δ) versus maximum estimated time of flightlessness (based mostly on phylogenetic divergence ages from published molecular studies with dated nodes, but also informed by time of geographic isolation) between the phylogenetically controlled volant-flightless sister pairs. More specifically, the shift (Δ) is the morphometric variable from the phylogenetically closest volant taxa to the flightless taxon, and it is examined against divergence age. These helped us establish hypotheses for morphological changes potentially linked to flight loss that can then be more rigorously tested through more sophisticated phylogenetic comparative methods.

We lacked known age estimates for *Gallinula nesiotis* and *Podilymbus gigas*, the latter is simply estimated to be less than 960 Ka. We can $\log_{10}$-transform before the linear model is applied in order to examine the intercept, slope, and $R^2$, but some ratites return ‘inf. error’ in R when filaments are absent (i.e., you cannot take the log of zero). Finally, we compared macroscopic whole body changes to microscopic changes in the primaries.

### 7.1. Macroscopic traits: pairwise regressions

#### 7.1.1. Body mass vs. divergence age

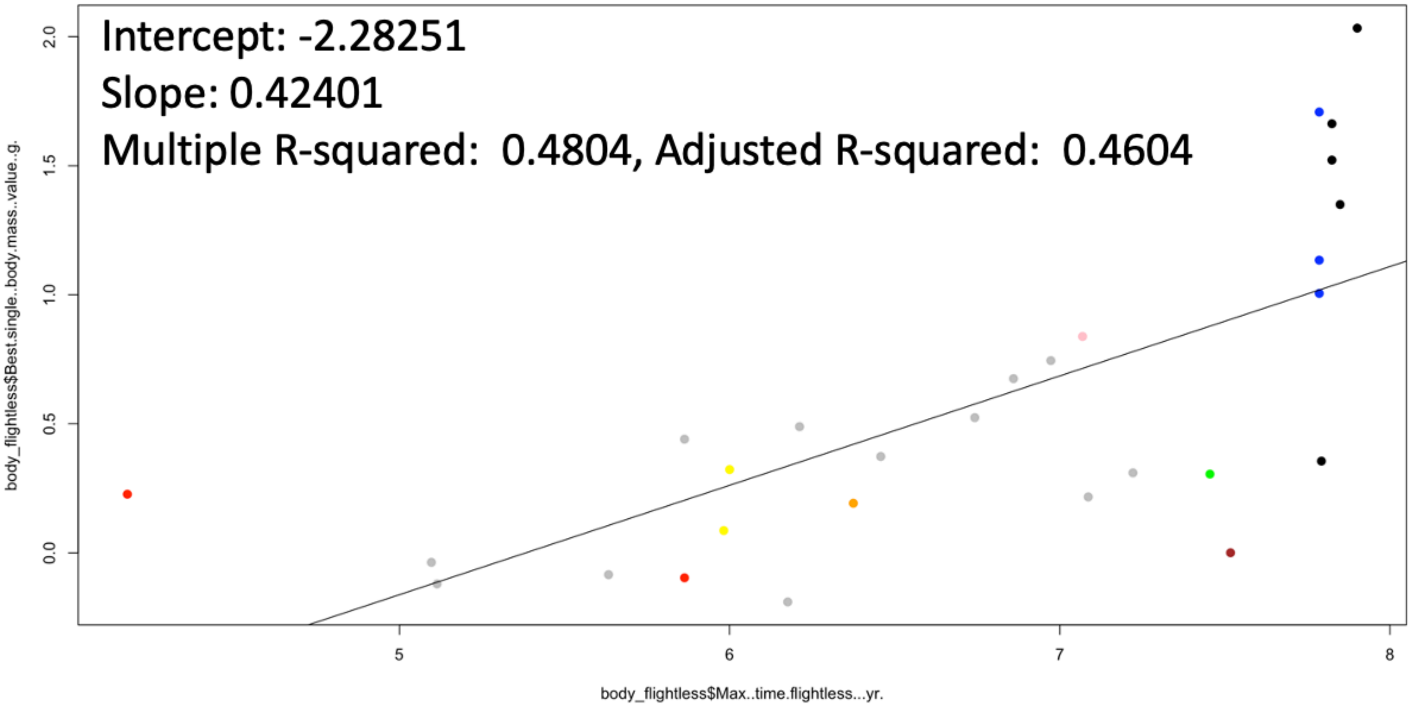


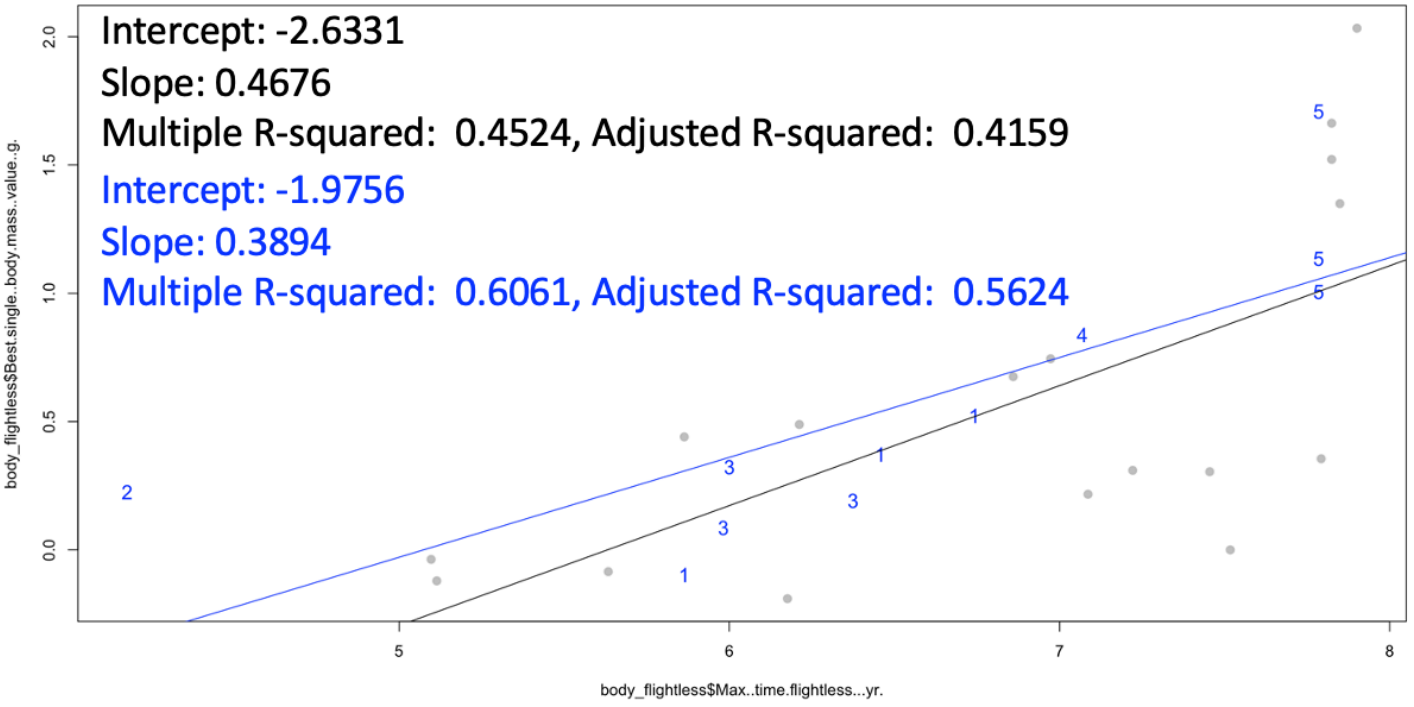

Figure S83. Shift in $\log_{10}$-tranformed body mass (flightless minus volant sister taxon comparisons) versus divergence age. $\Delta = \mathrm{Log}_{10}(\mathrm{Mass}_F) - \mathrm{Log}_{10}(\mathrm{Mass}_V)$. Top: clade colors as in plots above. Bottom: blue points are semiaquatic, while black are terrestrial. Linear regression results shown.

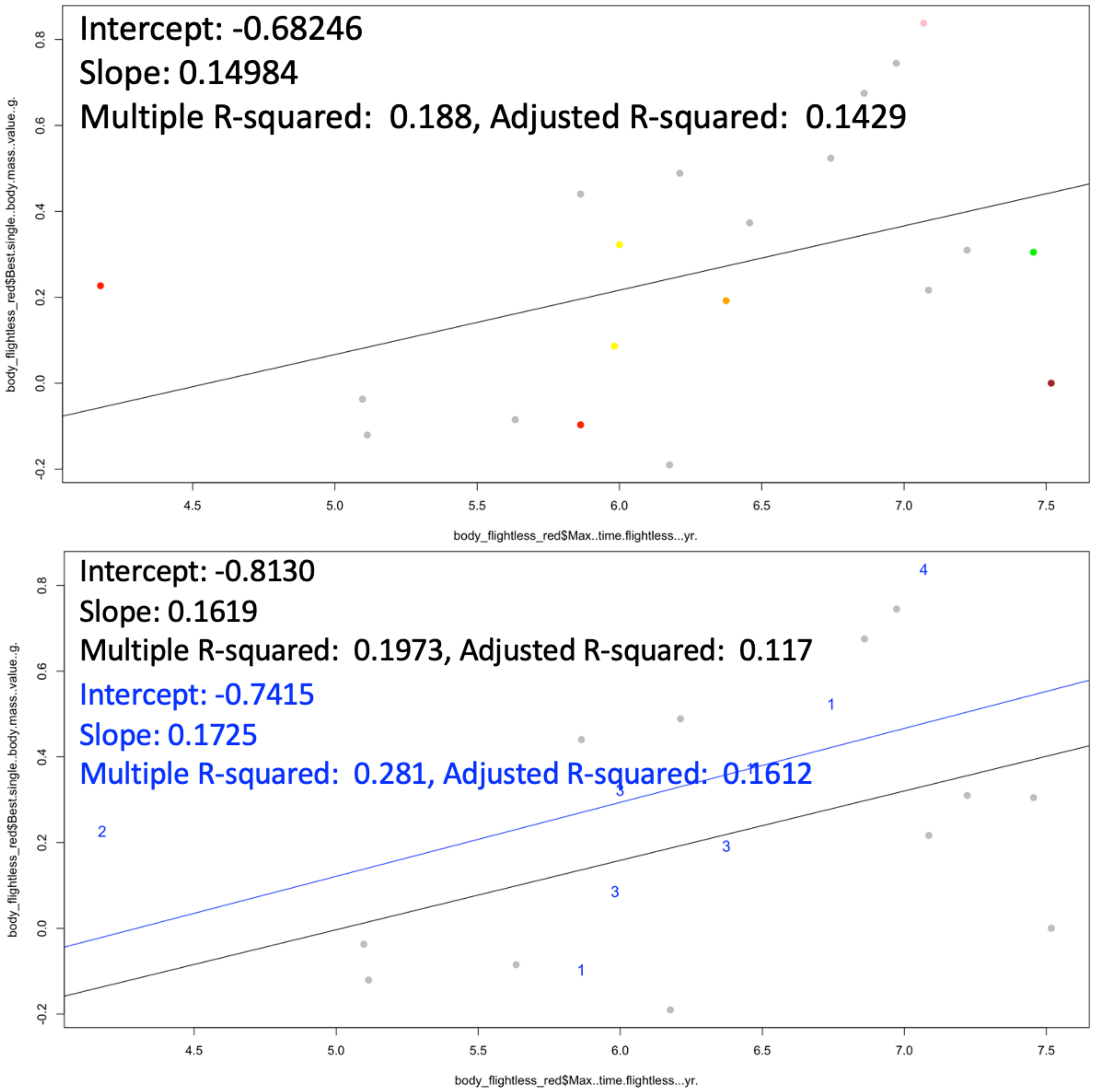


Figure S84. Shift in $\log_{10}$-tranformed body mass (flightless minus volant sister taxon comparisons) versus divergence age with penguins and ratites removed. Top: clade colors as in plots above. Bottom: blue points are semiaquatic, while black are terrestrial. Linear regression results shown.

#### 7.1.2. Wing length (scaled to body mass) vs. divergence age

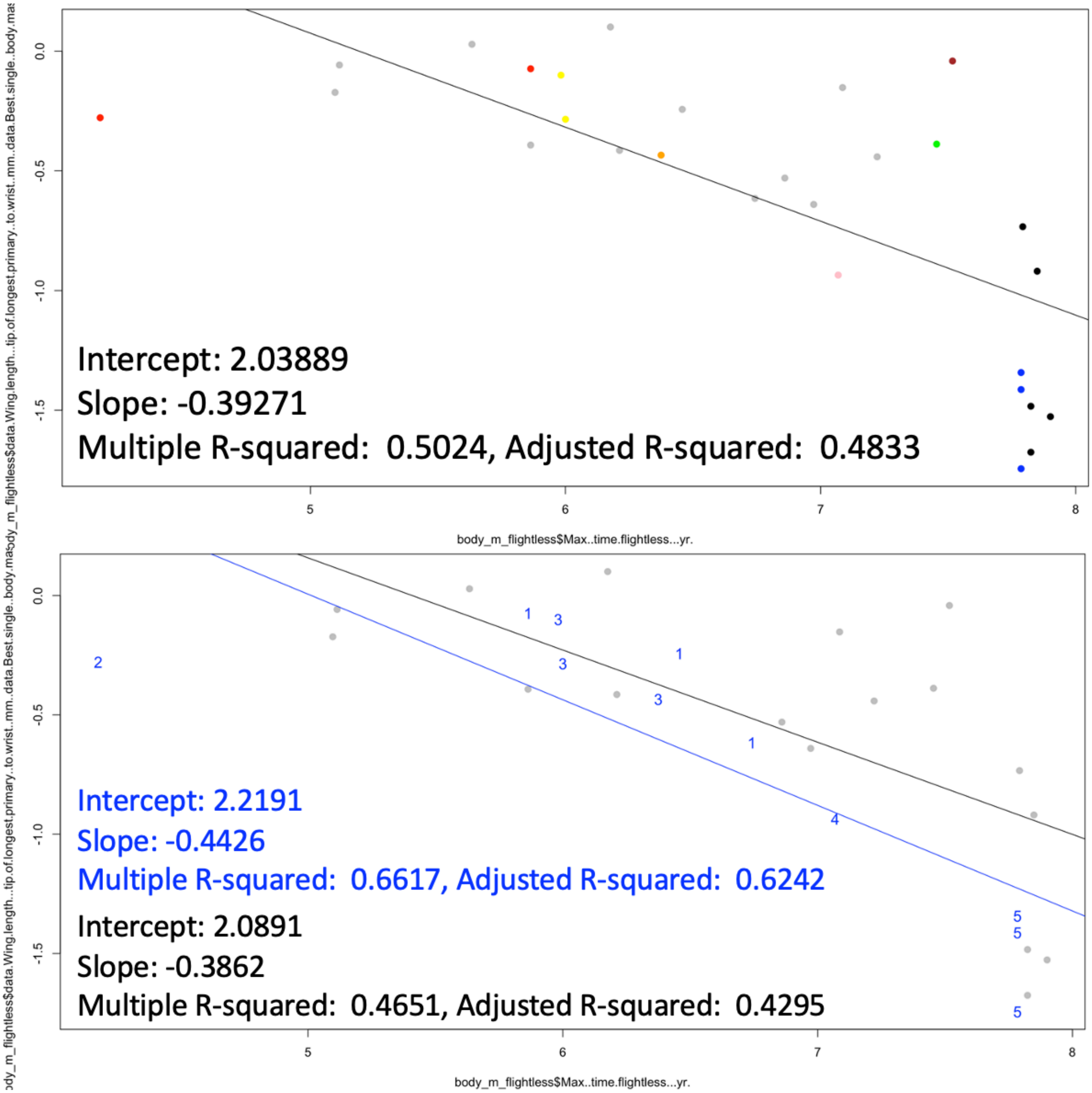


Figure S85. Shift in $log_{10}$-tranformed wing length/body mass (flightless minus volant sister taxon comparisons) versus divergence age. $\Delta = Log_{10}(Wing_F/Mass_F) - Log_{10}(Wing_V/Mass_V)$. Top: clade colors as in plots above. Bottom: blue points are semiaquatic, while black are terrestrial. Linear regression results shown.

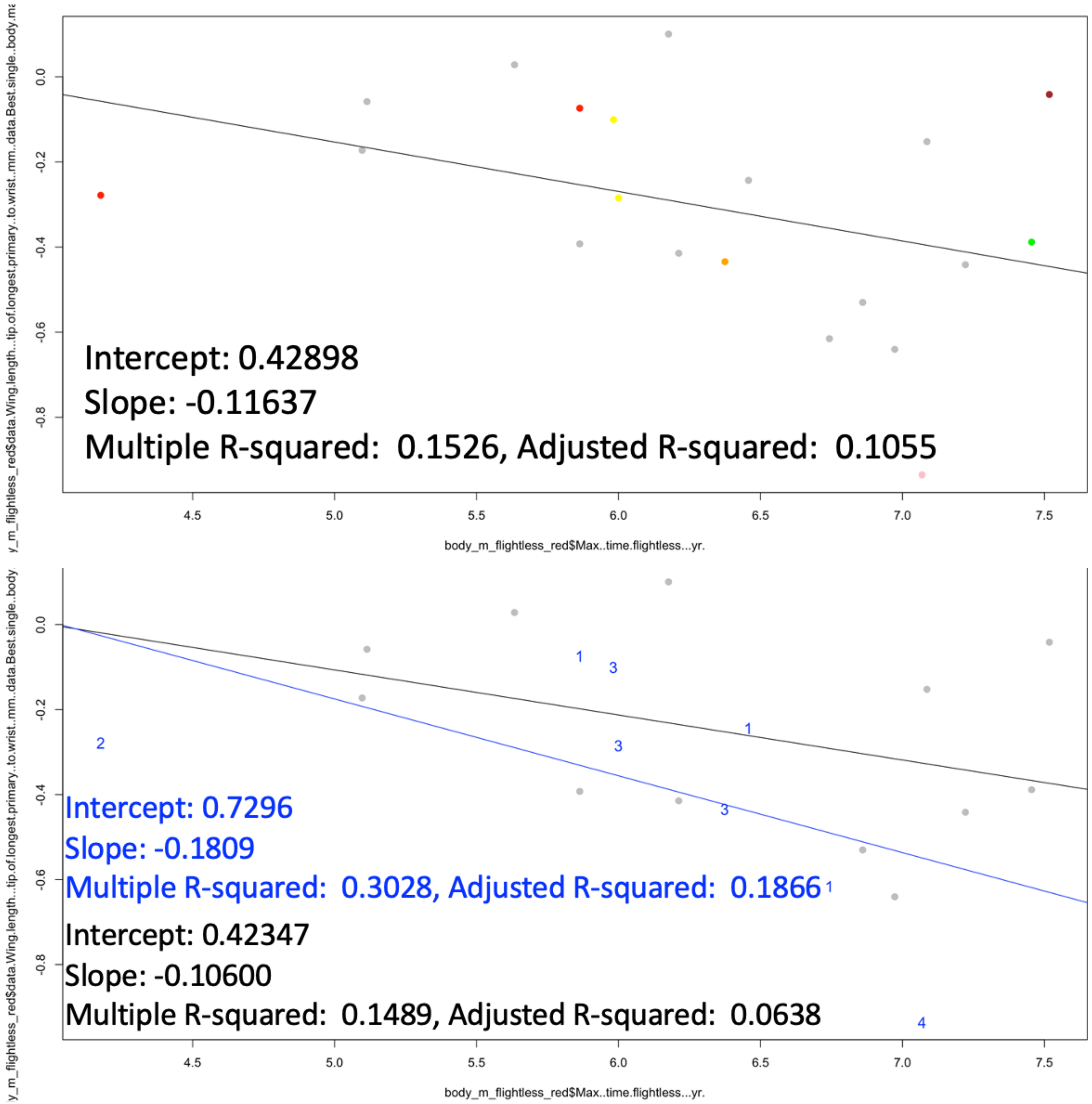


Figure S86. Shift in $\log_{10}$-tranformed wing length/body mass (flightless minus volant sister taxon comparisons) versus divergence age with penguins and ratites removed. Top: clade colors as in plots above. Bottom: blue points are semiaquatic, while black are terrestrial. Linear regression results shown.

#### 7.1.3. Tail length (scaled to body mass) vs. divergence age

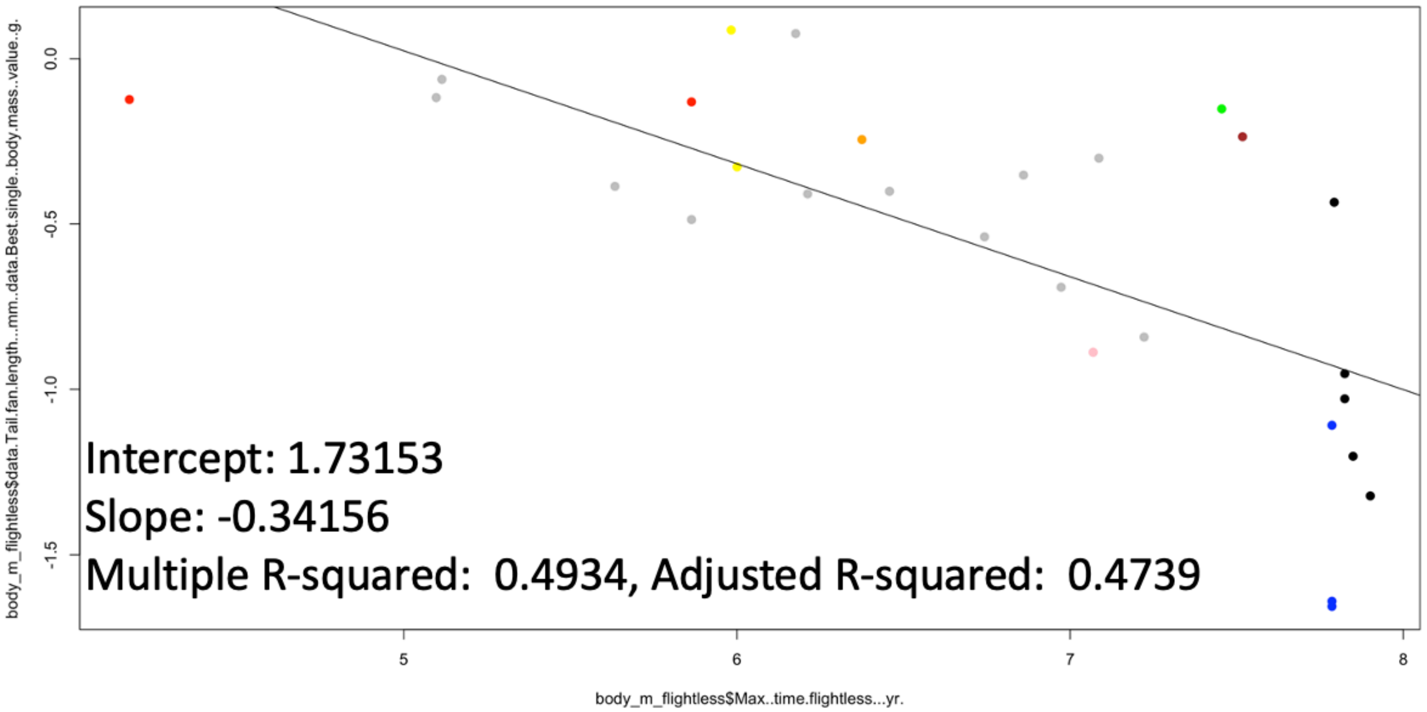


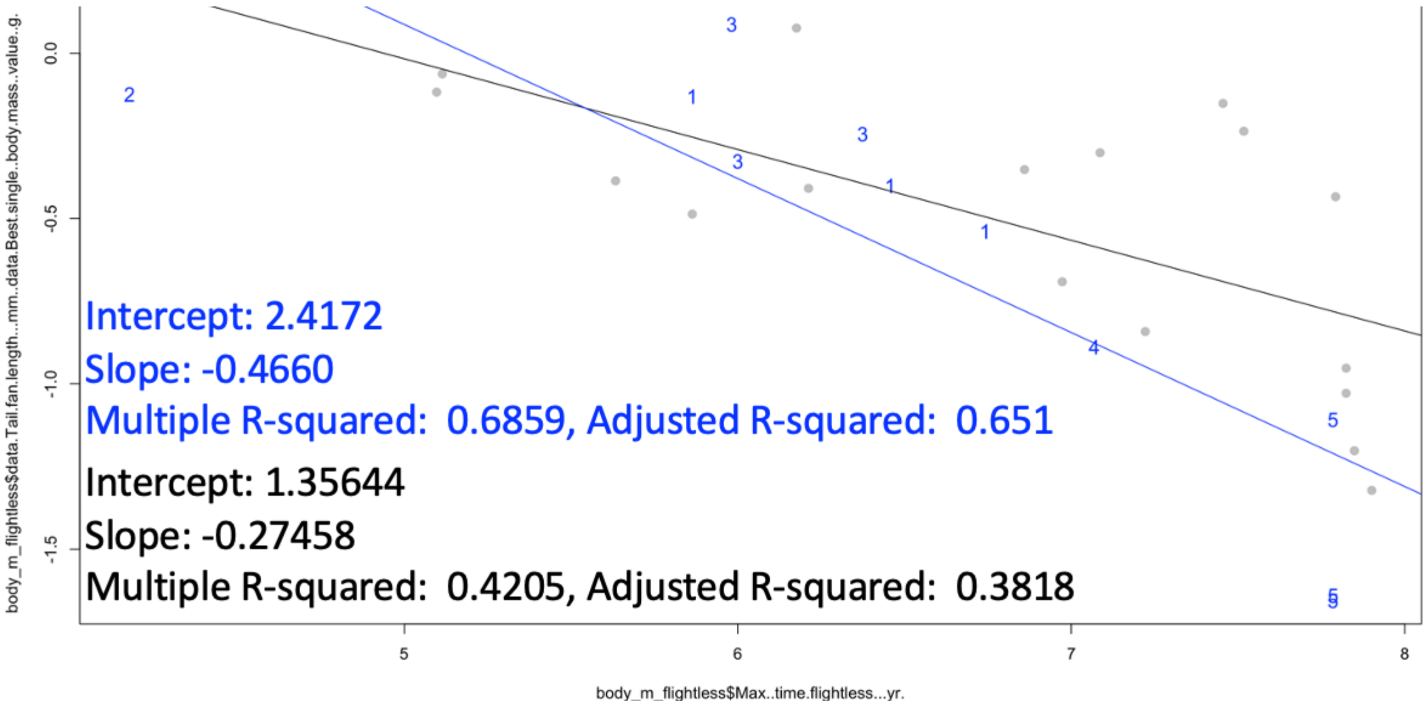


Figure S87. Shift in $\log_{10}$-tranformed tail fan length/body mass (flightless minus volant sister taxon comparisons) versus divergence age. $\Delta = \text{Log}_{10}(\text{Tail}_F/\text{Mass}_F) - \text{Log}_{10}(\text{Tail}_V/\text{Mass}_V)$. Top: clade colors as in plots above. Bottom: blue points are semiaquatic, while black are terrestrial. Linear regression results shown.

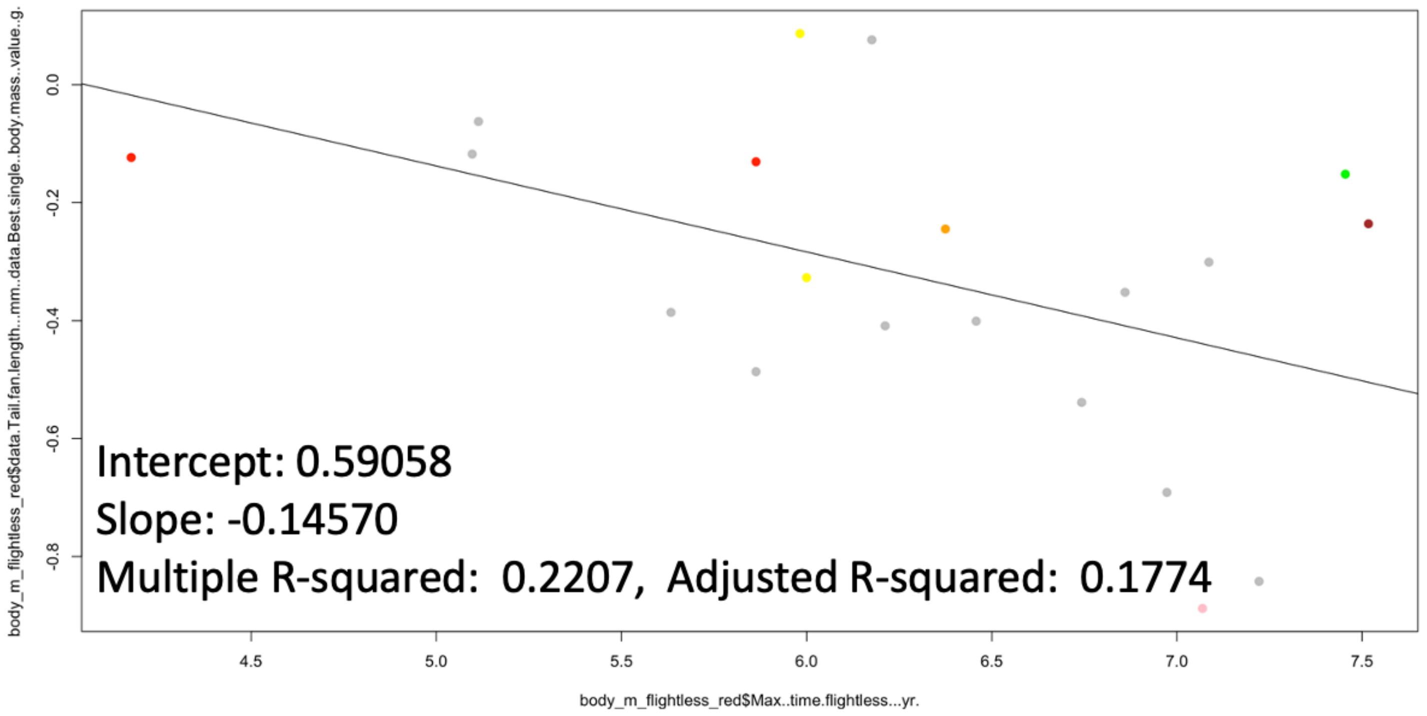


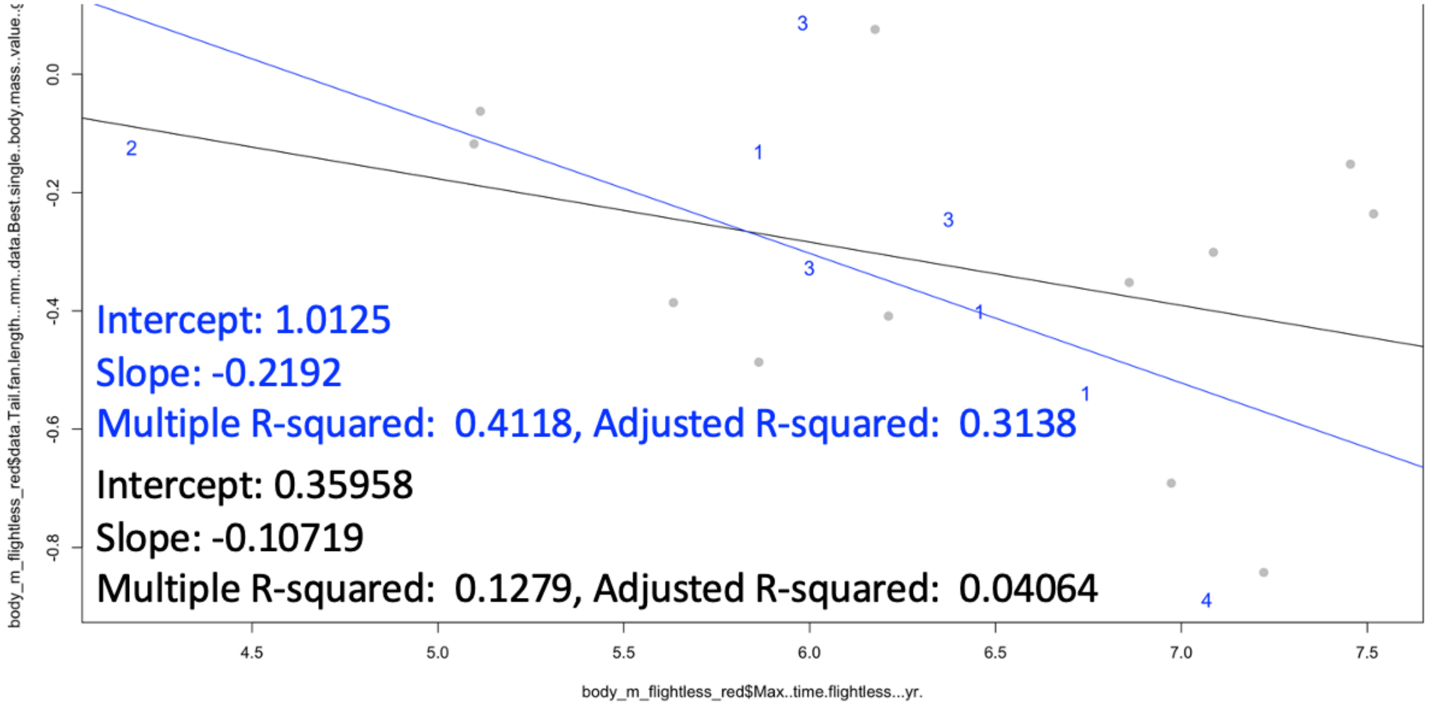


Figure S88. Shift in $\log_{10}$-tranformed tail fan length/body mass (flightless minus volant sister taxon comparisons) versus divergence age with penguins and ratites removed. Top: clade colors as in plots above. Bottom: blue points are semiaquatic, while black are terrestrial. Linear regression results shown.

#### 7.1.4. Difference in wing and tail length vs. divergence age

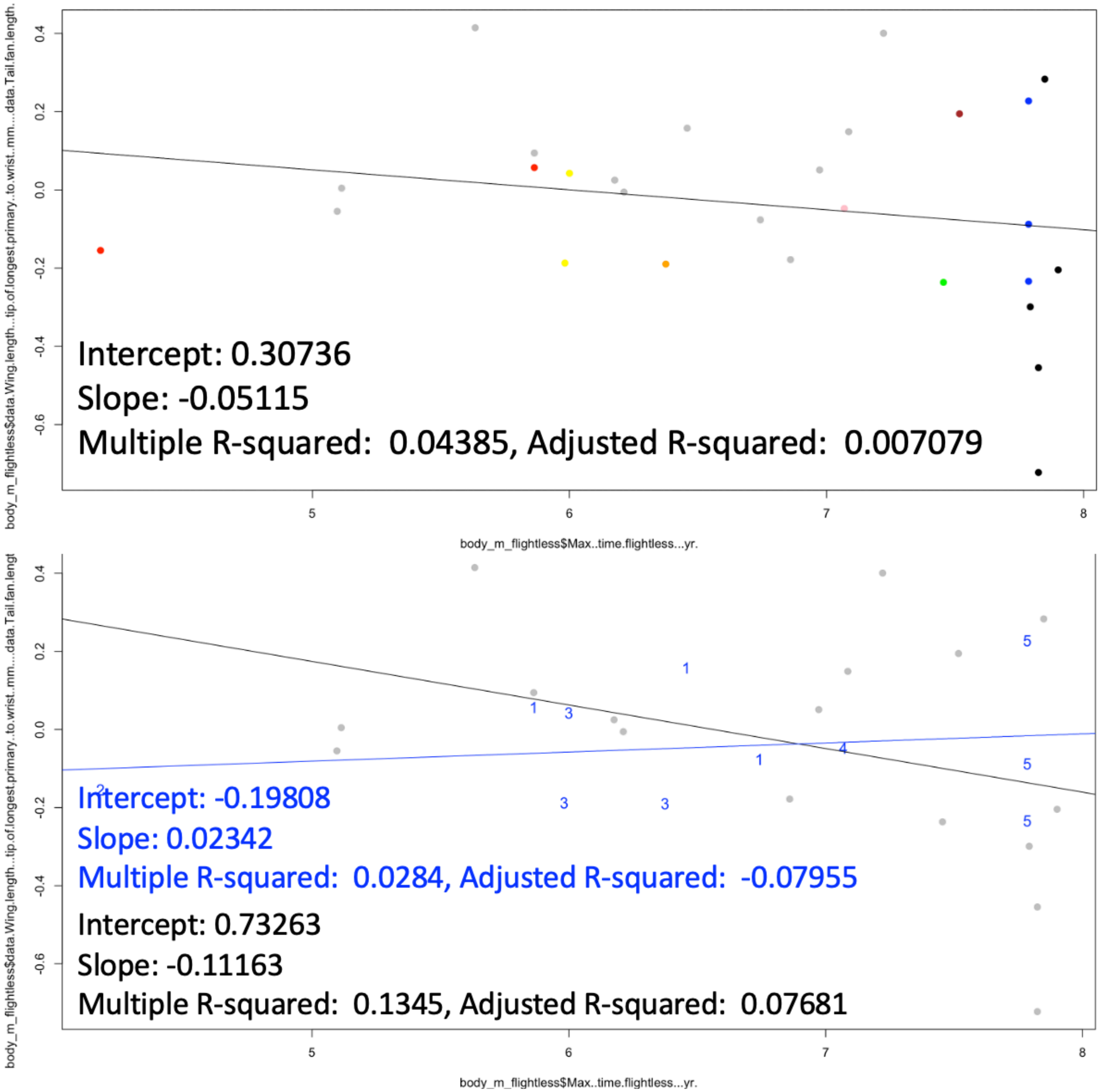


Figure S89. Shift in log$_{10}$-tranformed wing length minus tail fan length (flightless minus volant sister taxon comparisons) versus divergence age. $\Delta = (\text{Log}_{10}(\text{Wing}_F) - \text{Log}_{10}(\text{Tail}_F)) - (\text{Log}_{10}(\text{Wing}_V) - \text{Log}_{10}(\text{Tail}_V))$. Top: clade colors as in plots above. Bottom: blue points are semiaquatic, while black are terrestrial. Linear regression results shown.

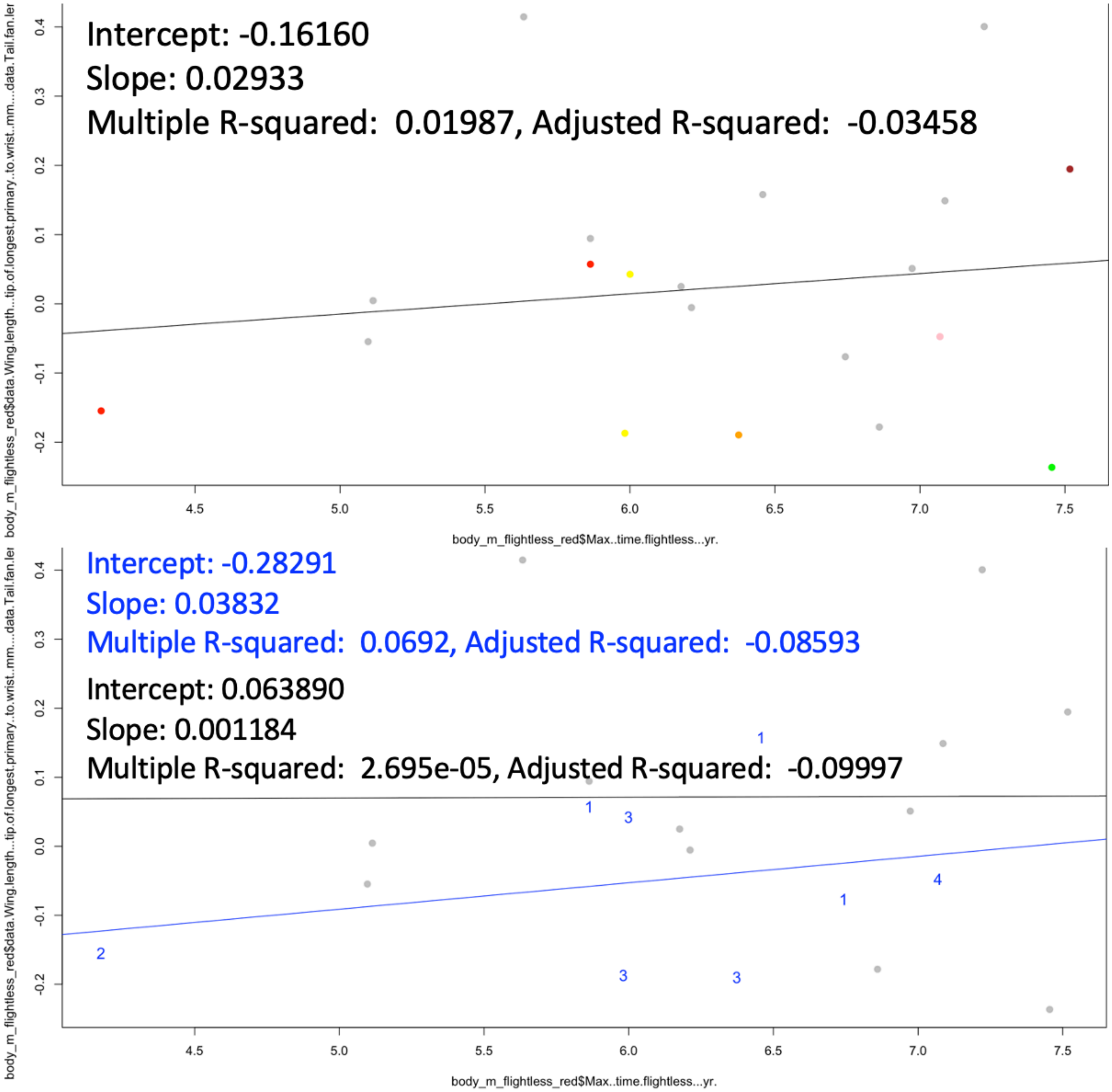


Figure S90. Shift in $\log_{10}$-tranformed wing length minus tail fan length (flightless minus volant sister taxon comparisons) versus divergence age with penguins and ratites removed. Top: clade colors as in plots above. Bottom: blue points are semiaquatic, while black are terrestrial. Linear regression results shown.

#### 7.1.5. Difference in wing and tarsus length vs. divergence age

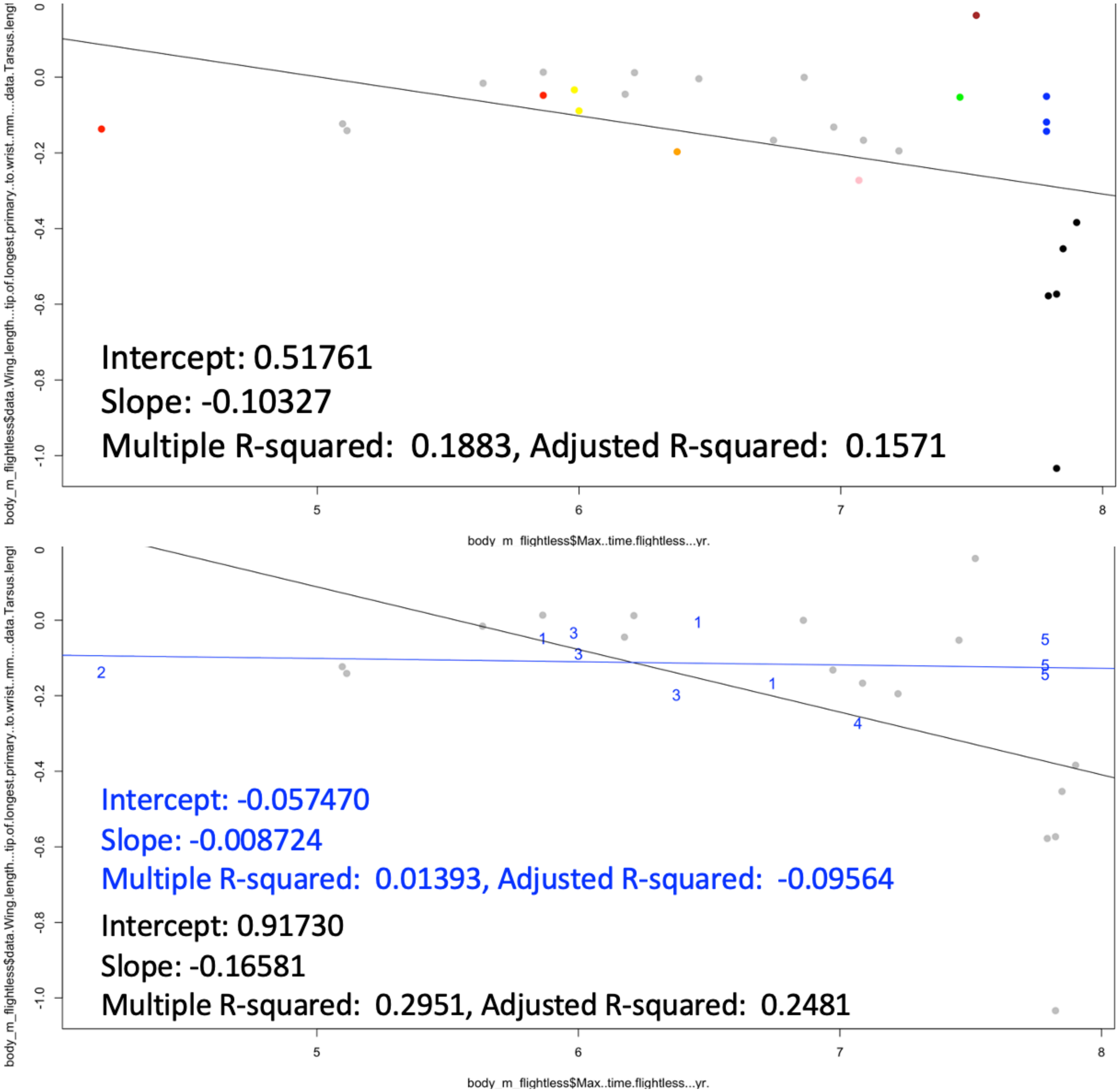


Figure S91. Shift in $\log_{10}$-tranformed wing length minus tarsus length (flightless minus volant sister taxon comparisons) versus divergence age. $\Delta = (\mathrm{Log}_{10}(\mathrm{Wing}_F) - \mathrm{Log}_{10}(\mathrm{Tarsus}_F)) - (\mathrm{Log}_{10}(\mathrm{Wing}_V) - \mathrm{Log}_{10}(\mathrm{Tarsus}_V))$. Top: clade colors as in plots above. Bottom: blue points are semiaquatic, while black are terrestrial. Linear regression results shown.

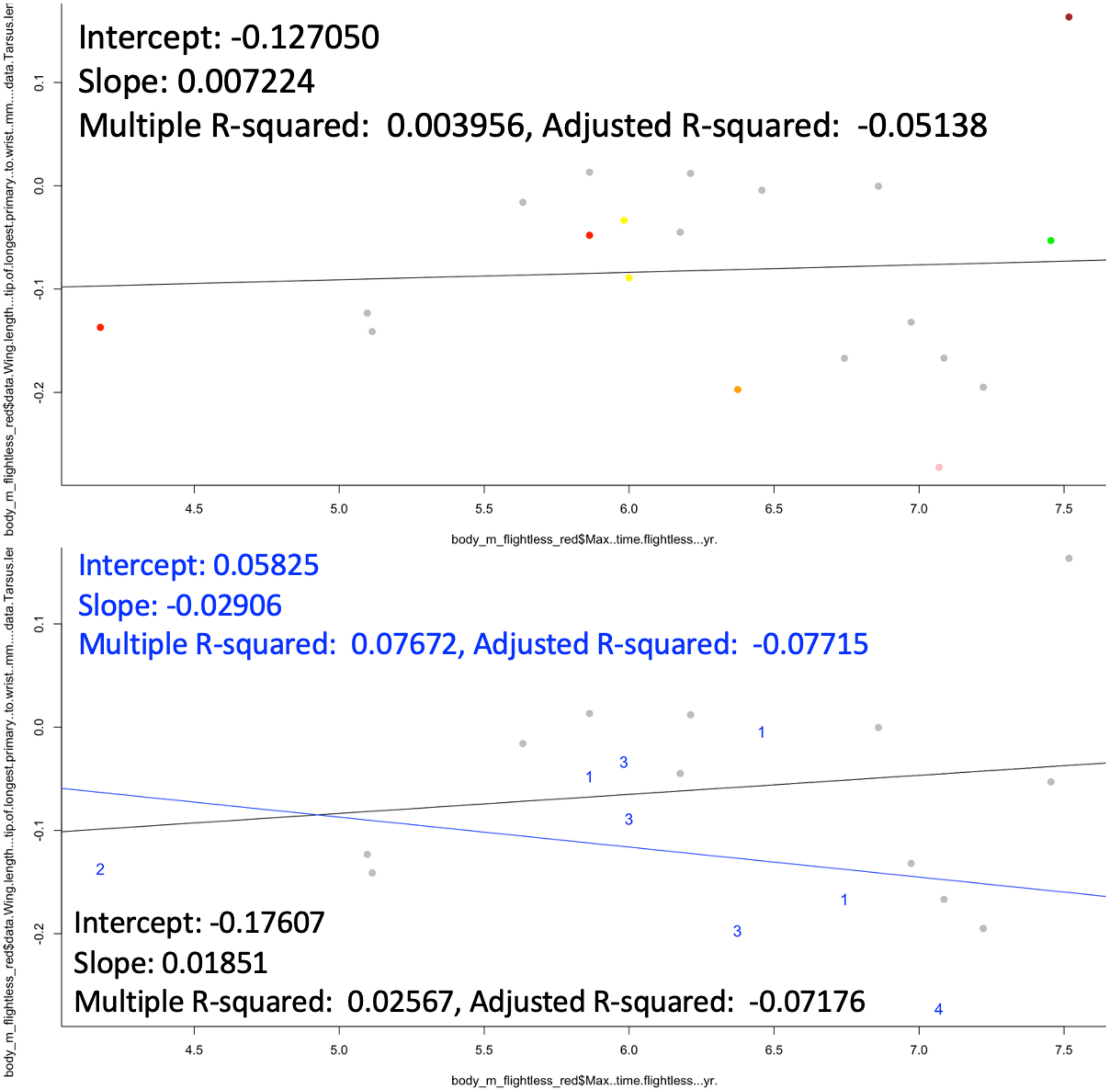


Figure S92. Shift in $log_{10}$-tranformed wing length minus tarsus length (flightless minus volant sister taxon comparisons) versus divergence age with penguins and ratites removed. Top: clade colors as in plots above. Bottom: blue points are semiaquatic, while black are terrestrial. Linear regression results shown.

#### 7.1.6. Macroscopic trait pairwise regression summary

The best signatures of flightlessness based on these regressions with simplistic phylogenetic control (above) are body mass increase, wing length/mass ratio decrease, and tail fan length/mass ratio decrease. This may be consistent even in early evolution, according to these results.

The shifts might appear to be more rapid in semiaquatic taxa in these results due to either water buoyancy that more readily allows for larger body size or novel selective pressures related to locomotion in water (rather than simply a relaxation of selection for flight).

### 7.2. Microscopic traits: pairwise regressions

Primary remiges were examined under the assumption that they are likely to show the most prominent signal to noise ratio, because they are most crucial for flight among our sampled feather positions. The middle region exposed on the rachis was examined for barb (leading and trailing) lengths, rachis width, and (distal) barbule length, all of which were scaled to rachis length. $Log_{10}$-transformation was used to determine the shift from volant to flightless phylogenetically related taxa as follows: $\Delta = Log_{10}(M_F/L_F)-Log_{10}(M_V/L_V)$, where $M_F$ and $M_V$ are the microscopic morphometric and $L_F$ and $L_V$ are the rachis length from the apical/distal node of the feather to that point of the middle region exposed along the feather, for flightless and volant sister taxa respectively. Barb width was not examined since it correlates with other variables (e.g., barbule length) and pushes the limits of the microscope's measurement capability.

#### 7.2.1. Rachis width vs. divergence age

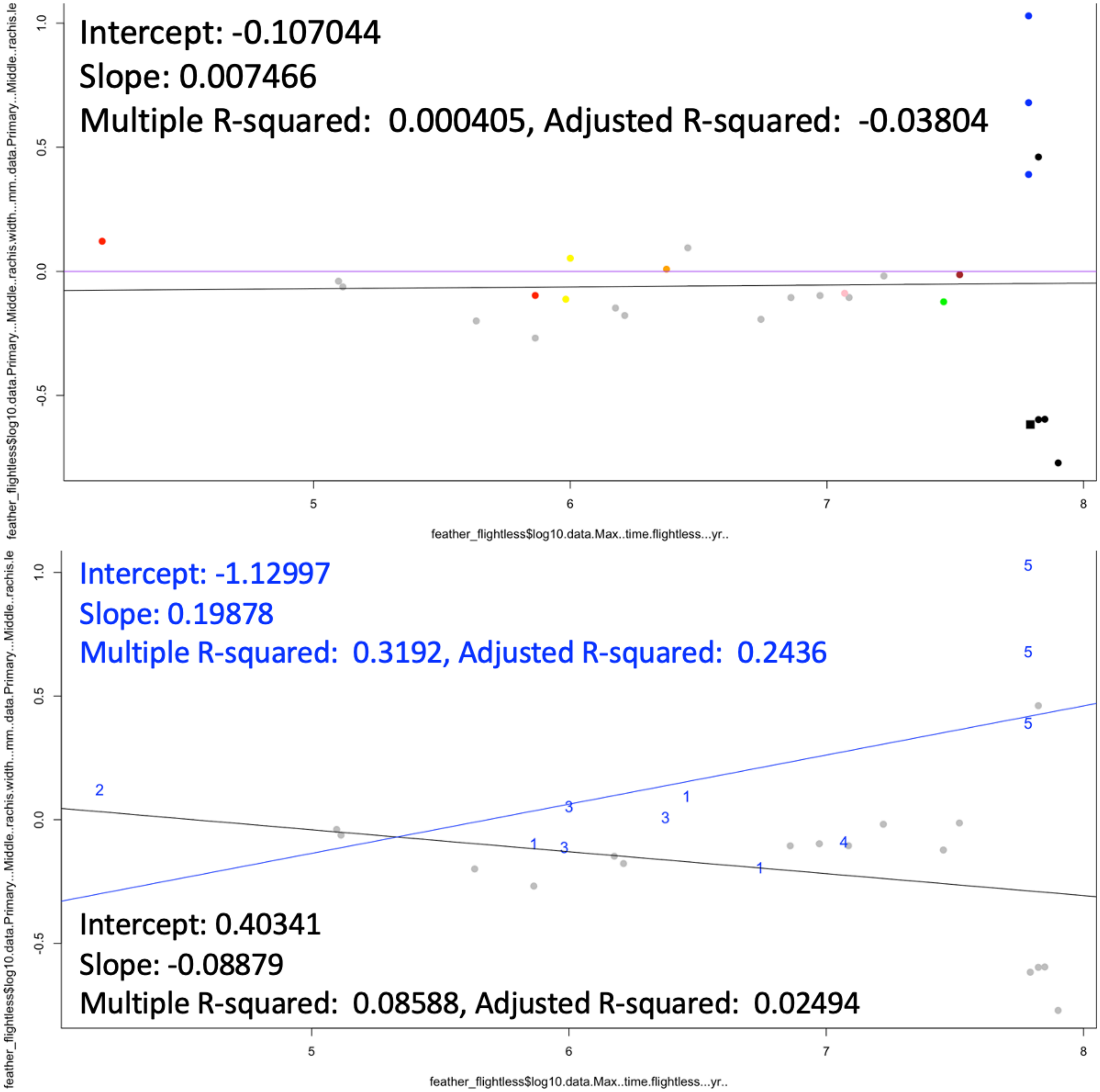

Figure S93. Shift in $log_{10}$-tranformed rachis width/rachis length (flightless minus volant sister taxon comparisons) versus divergence age. $\Delta = Log_{10}(Width_F/Length_F) - Log_{10}(Width_V/Length_V)$. Top: clade colors as in plots above. Bottom: blue points are semiaquatic, while black are terrestrial. Linear regression results shown.

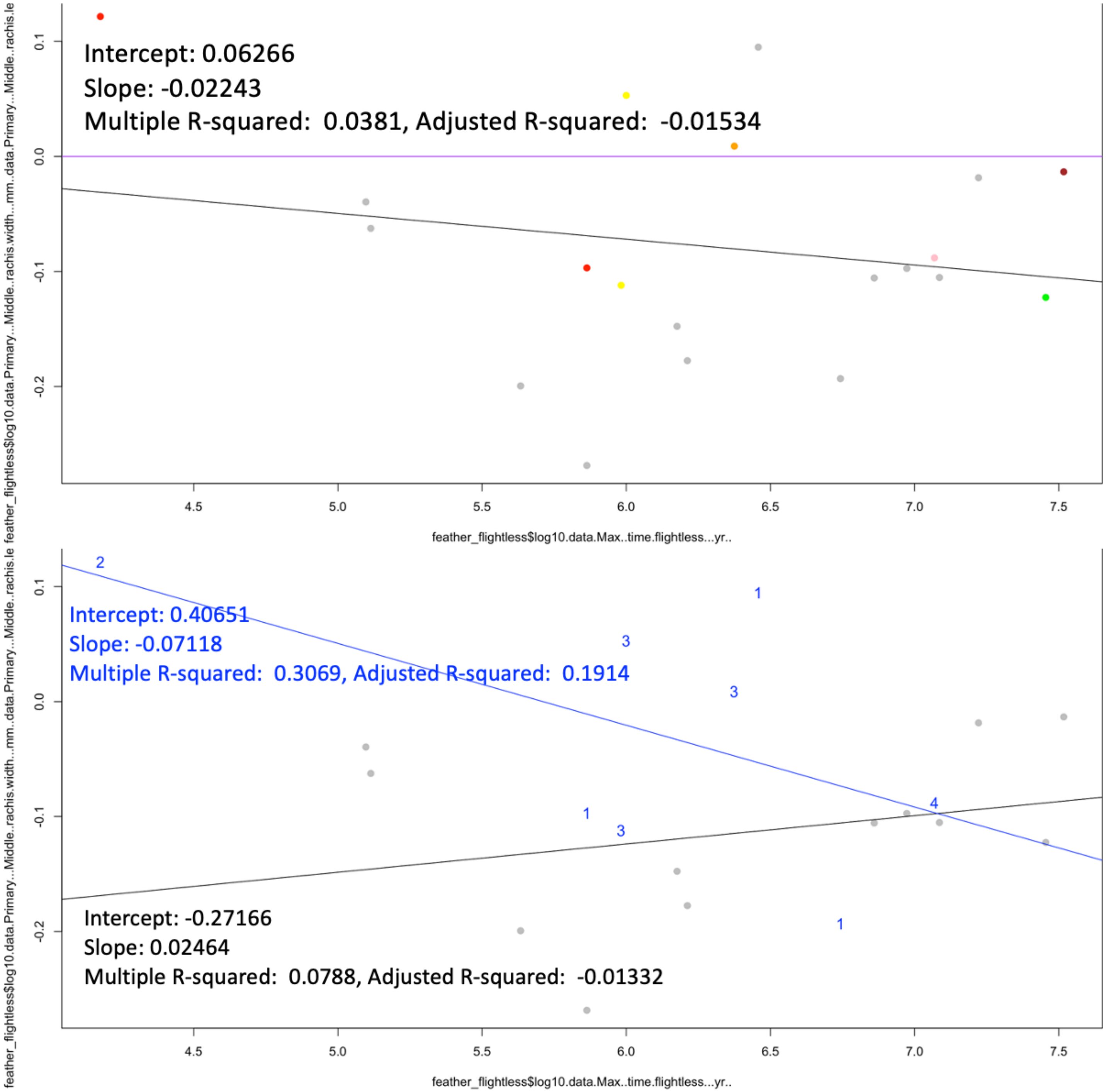


Figure S94. Shift in $\log_{10}$-tranformed rachis width/rachis length (flightless minus volant sister taxon comparisons) versus divergence age with penguins and ratites removed. Top: clade colors as in plots above. Bottom: blue points are semiaquatic, while black are terrestrial. Linear regression results shown.

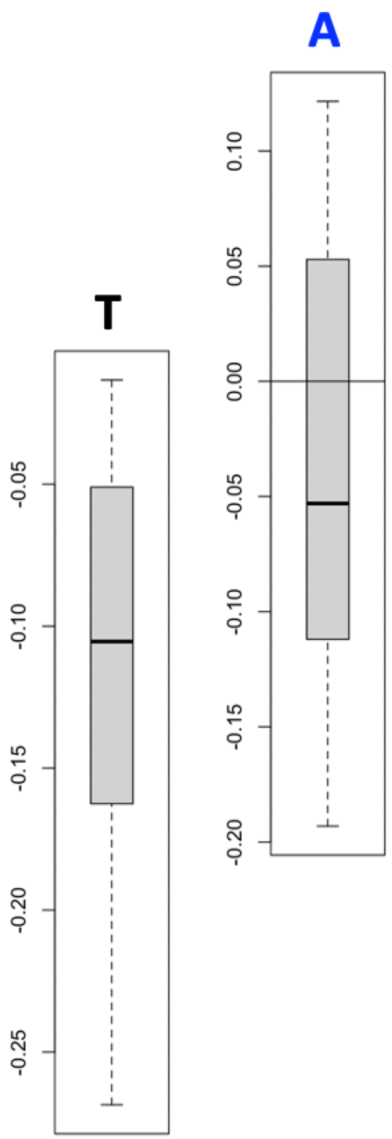


Figure S95. Boxplots of shift in $log_{10}$-tranformed rachis width/rachis length (flightless minus volant sister taxon comparisons) versus divergence age with penguins and ratites removed. T: Terrestrial taxa. A: Semiaquatic taxa. Not shown to scale.

#### 7.2.2. Leading barb length vs. divergence age

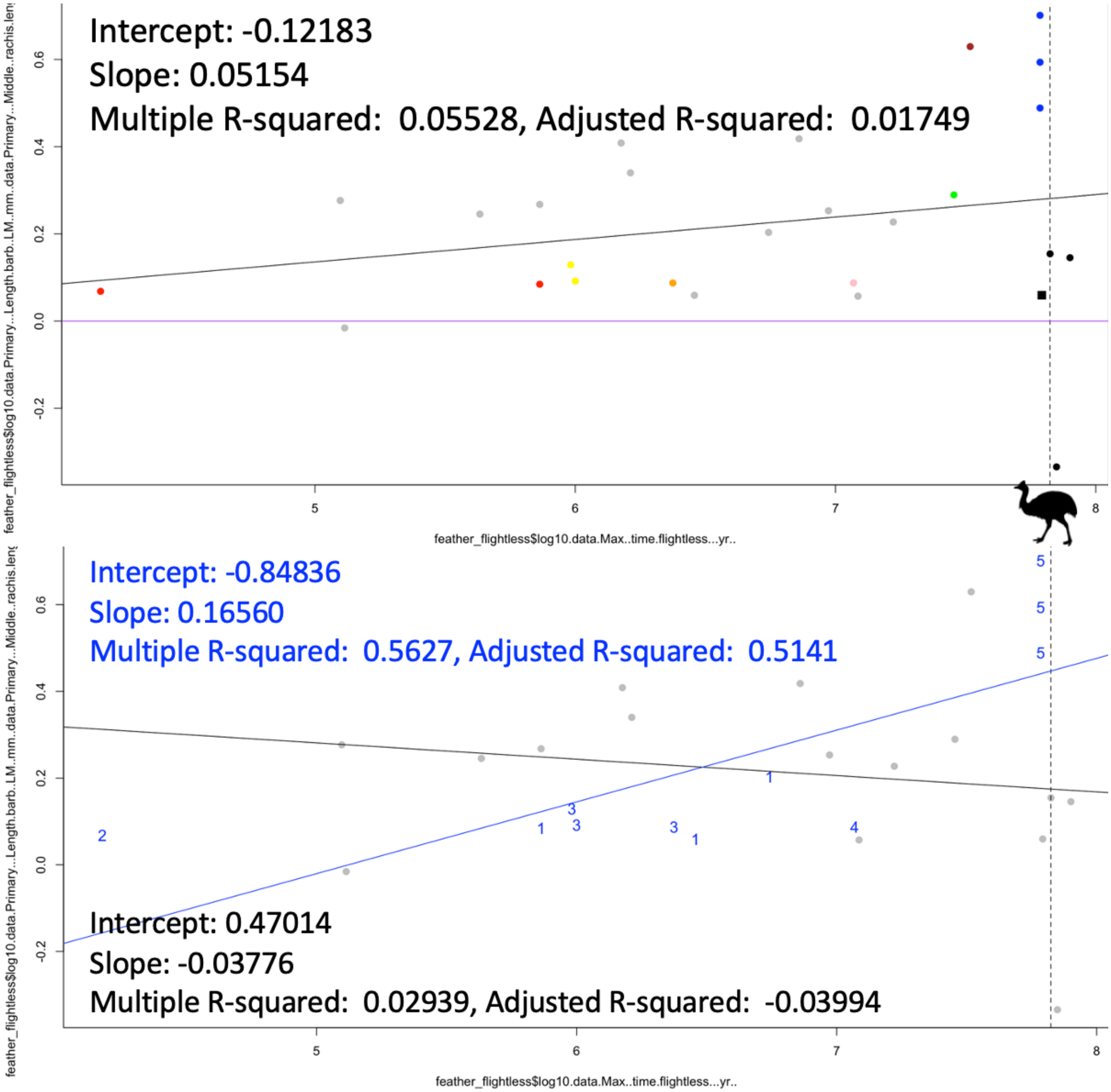


Figure S96. Shift in $log_{10}$-tranformed leading barb length/rachis length (flightless minus volant sister taxon comparisons) versus divergence age. $\Delta = Log_{10}(Barb_F/Rachis_F) - Log_{10}(Barb_V/Rachis_V)$. Top: clade colors as in plots above. Bottom: blue points are semiaquatic, while black are terrestrial. Linear regression results shown. Vertical dashed line indicates cassowary divergence age since barb loss precludes $log_{10}$-transformation and inclusion in the plot/regressions.

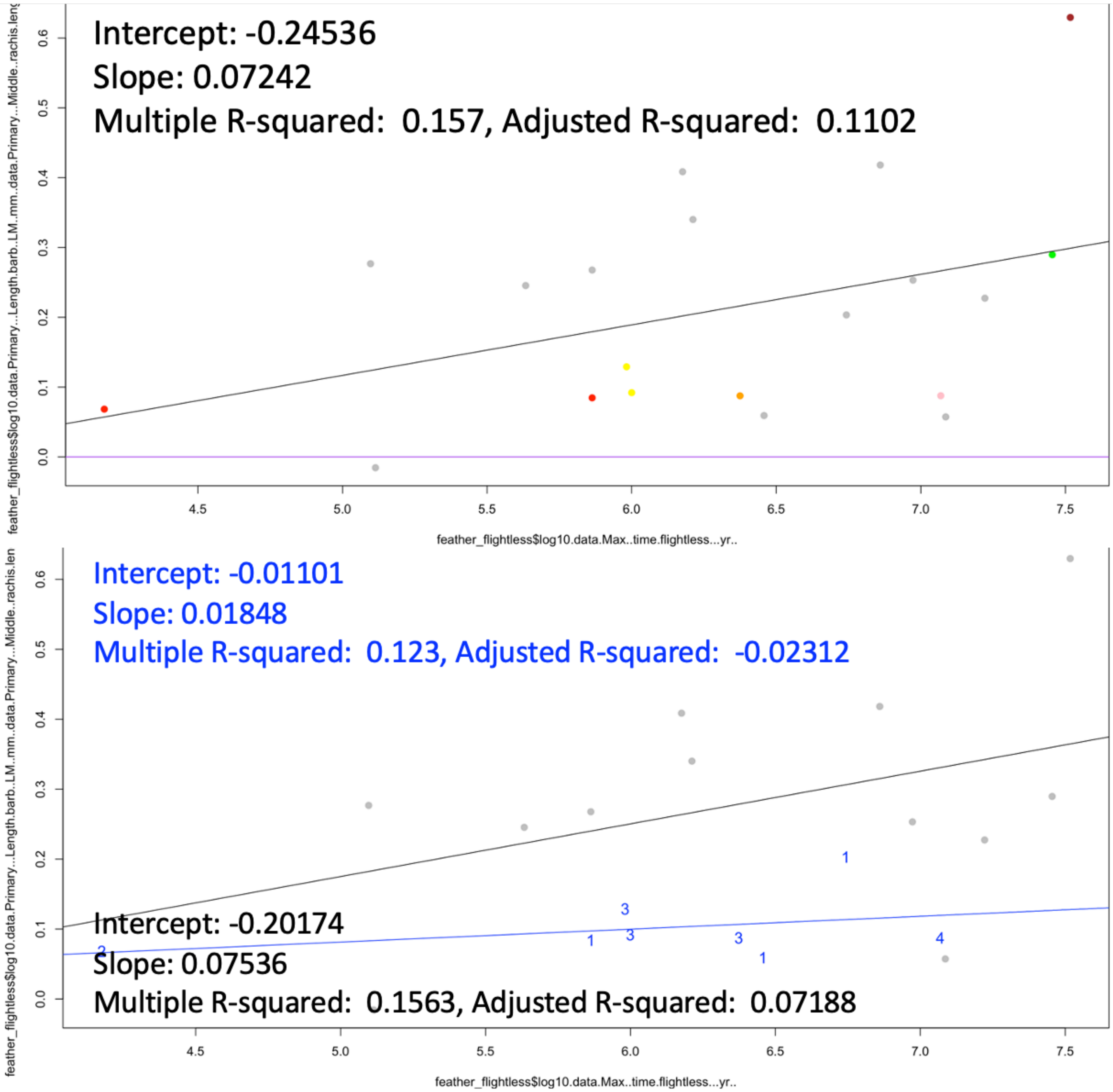


Figure S97. Shift in $log_{10}$-tranformed leading barb length/rachis length (flightless minus volant sister taxon comparisons) versus divergence age with penguins and ratites removed. Top: clade colors as in plots above. Bottom: blue points are semiaquatic, while black are terrestrial. Linear regression results shown.

#### 7.2.3. Trailing barb length vs. divergence age

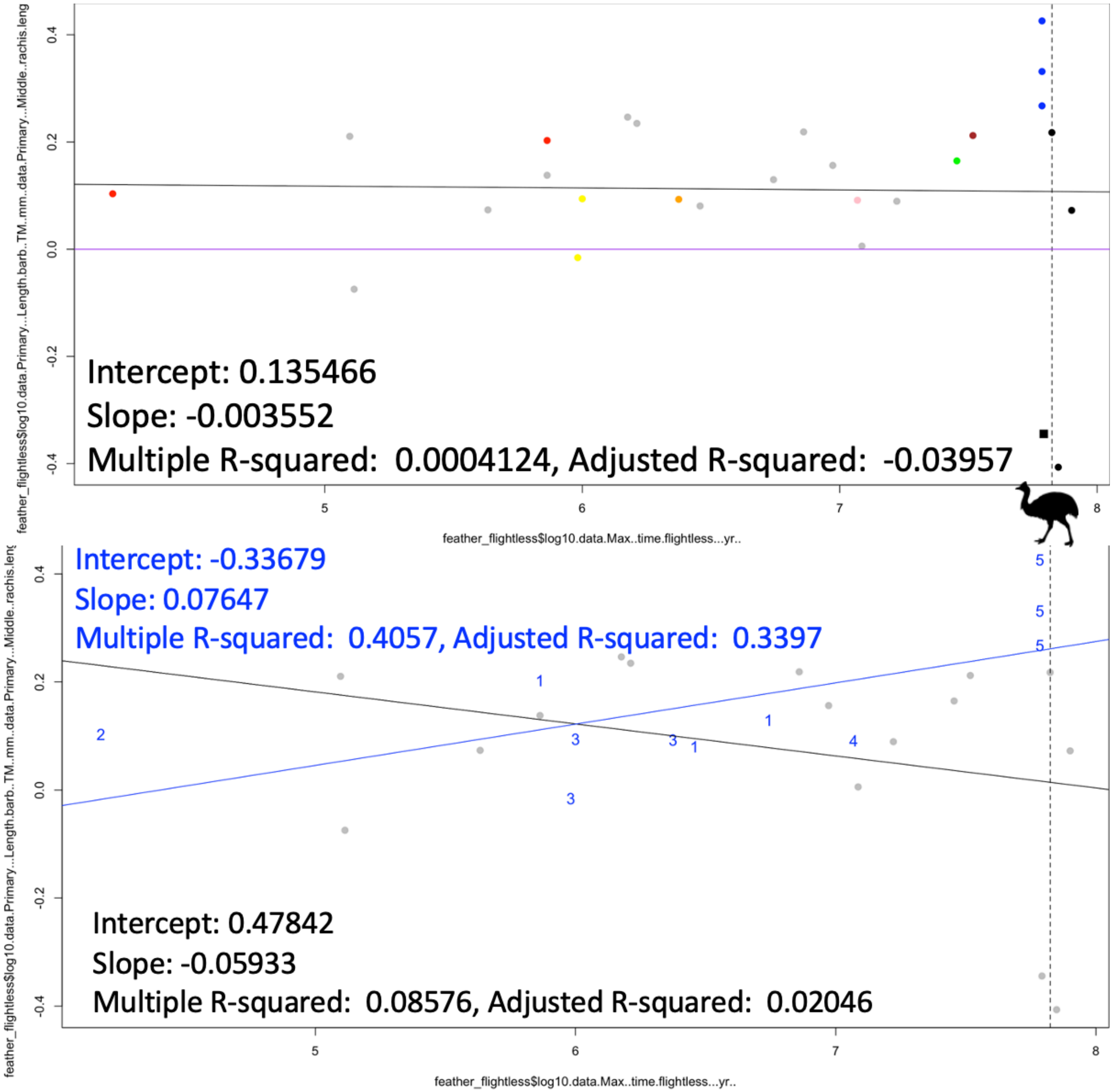


Figure S98. Shift in $\log_{10}$-tranformed trailing barb length/rachis length (flightless minus volant sister taxon comparisons) versus divergence age. $\Delta = \mathrm{Log}_{10}(\mathrm{Barb}_F/\mathrm{Rachis}_F)-\mathrm{Log}_{10}(\mathrm{Barb}_V/\mathrm{Rachis}_V)$. Top: clade colors as in plots above. Bottom: blue points are semiaquatic, while black are terrestrial. Linear regression results shown. Vertical dashed line indicates cassowary divergence age since barb loss precludes $\log_{10}$-transformation and inclusion in the plot/regressions.

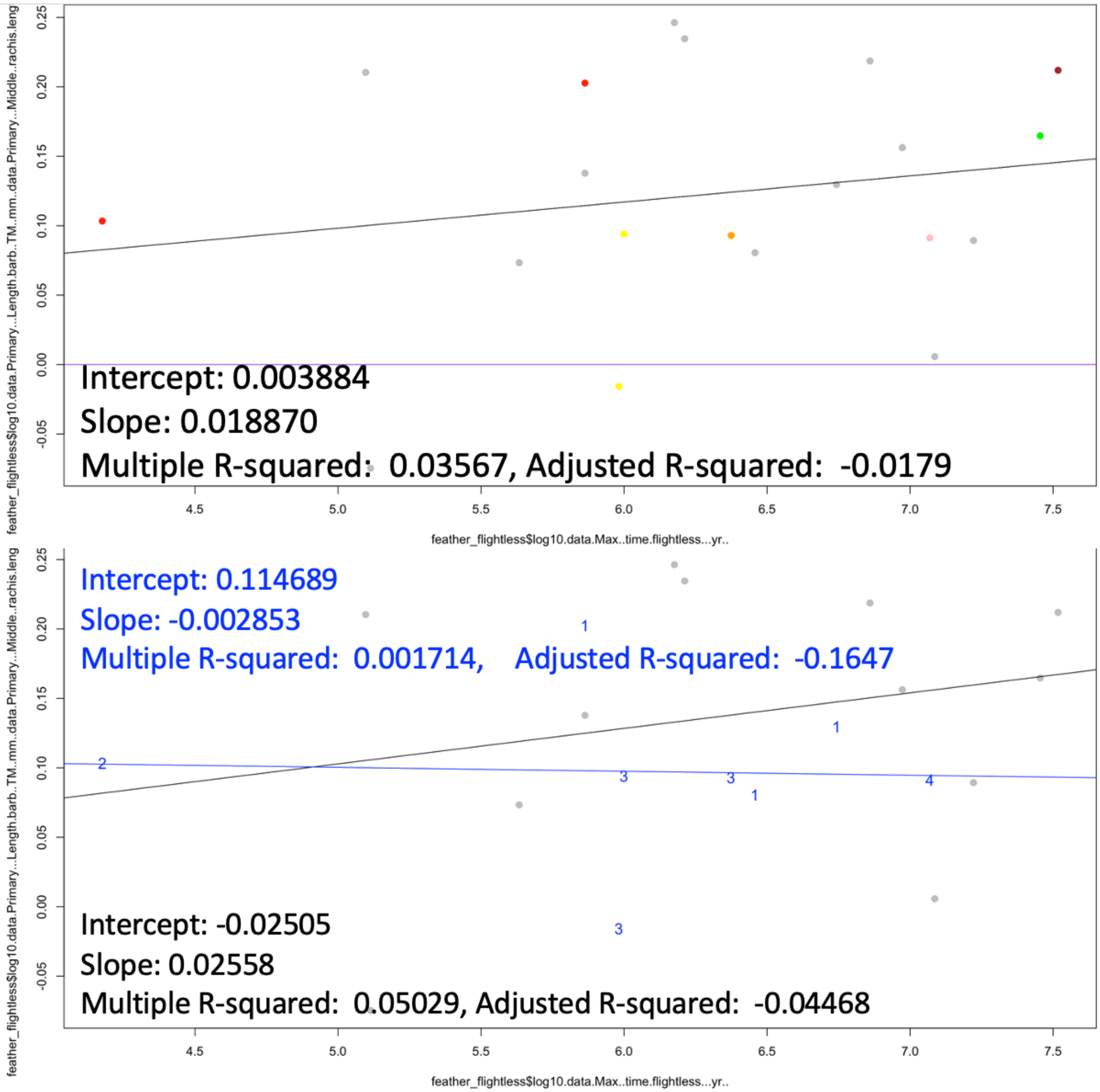


Figure S99. Shift in $log_{10}$-tranformed trailing barb length/rachis length (flightless minus volant sister taxon comparisons) versus divergence age with penguins and ratites removed. Top: clade colors as in plots above. Bottom: blue points are semiaquatic, while black are terrestrial. Linear regression results shown.

#### 7.2.4. Barbule length vs. divergence age

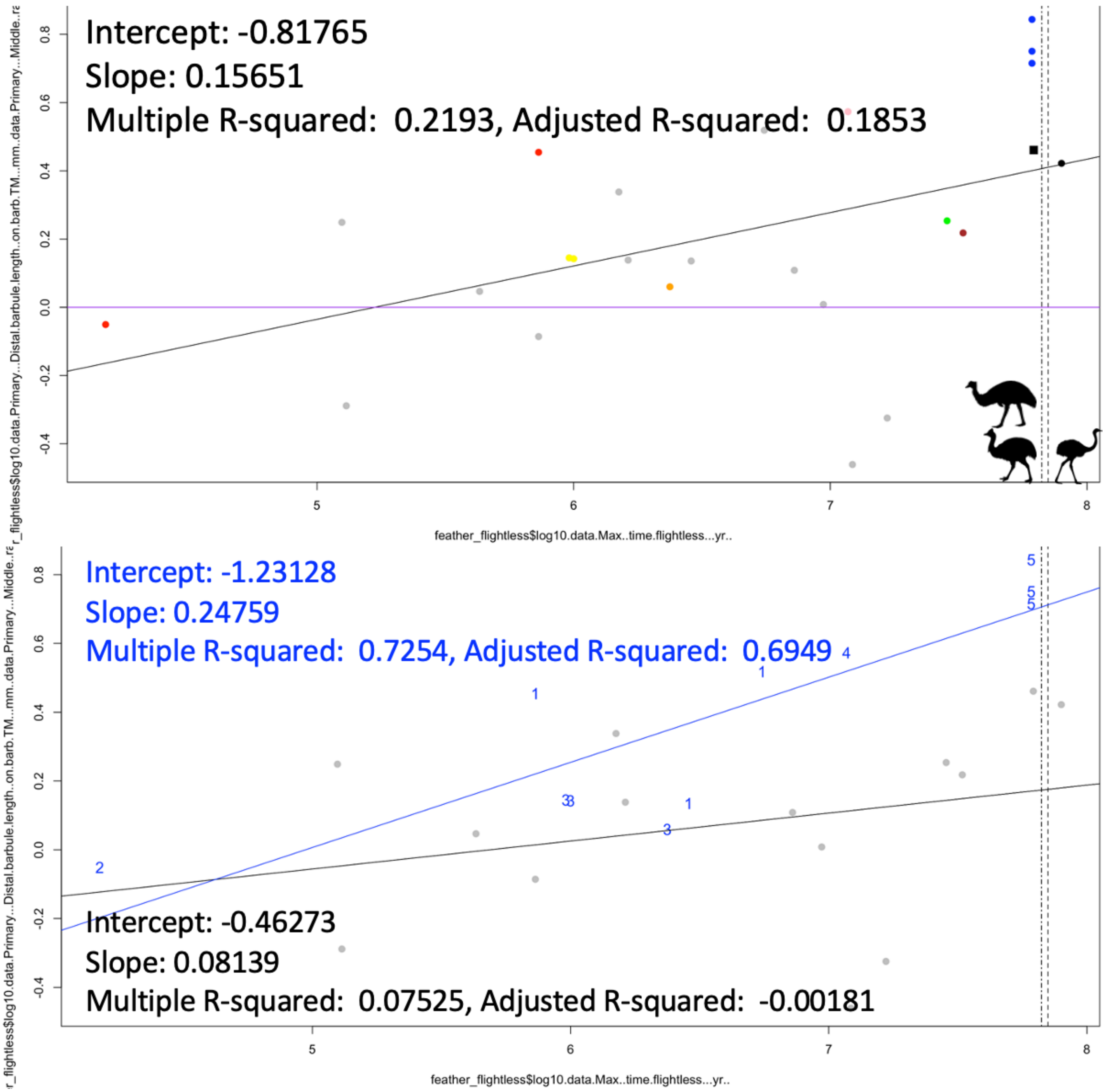


Figure S100. Shift in $\log_{10}$-tranformed distal barbule length/rachis length (flightless minus volant sister taxon comparisons) versus divergence age. $\Delta = \text{Log}_{10}(\text{Barbule}_F/\text{Rachis}_F) - \text{Log}_{10}(\text{Barbule}_V/\text{Rachis}_V)$. Top: clade colors as in plots above. Bottom: blue points are semiaquatic, while black are terrestrial. Linear regression results shown. Vertical dashed lines indicate emu and cassowary (left) and rhea (right) divergence ages since barbules loss precludes $\log_{10}$-transformation and inclusion in the plot/regressions.

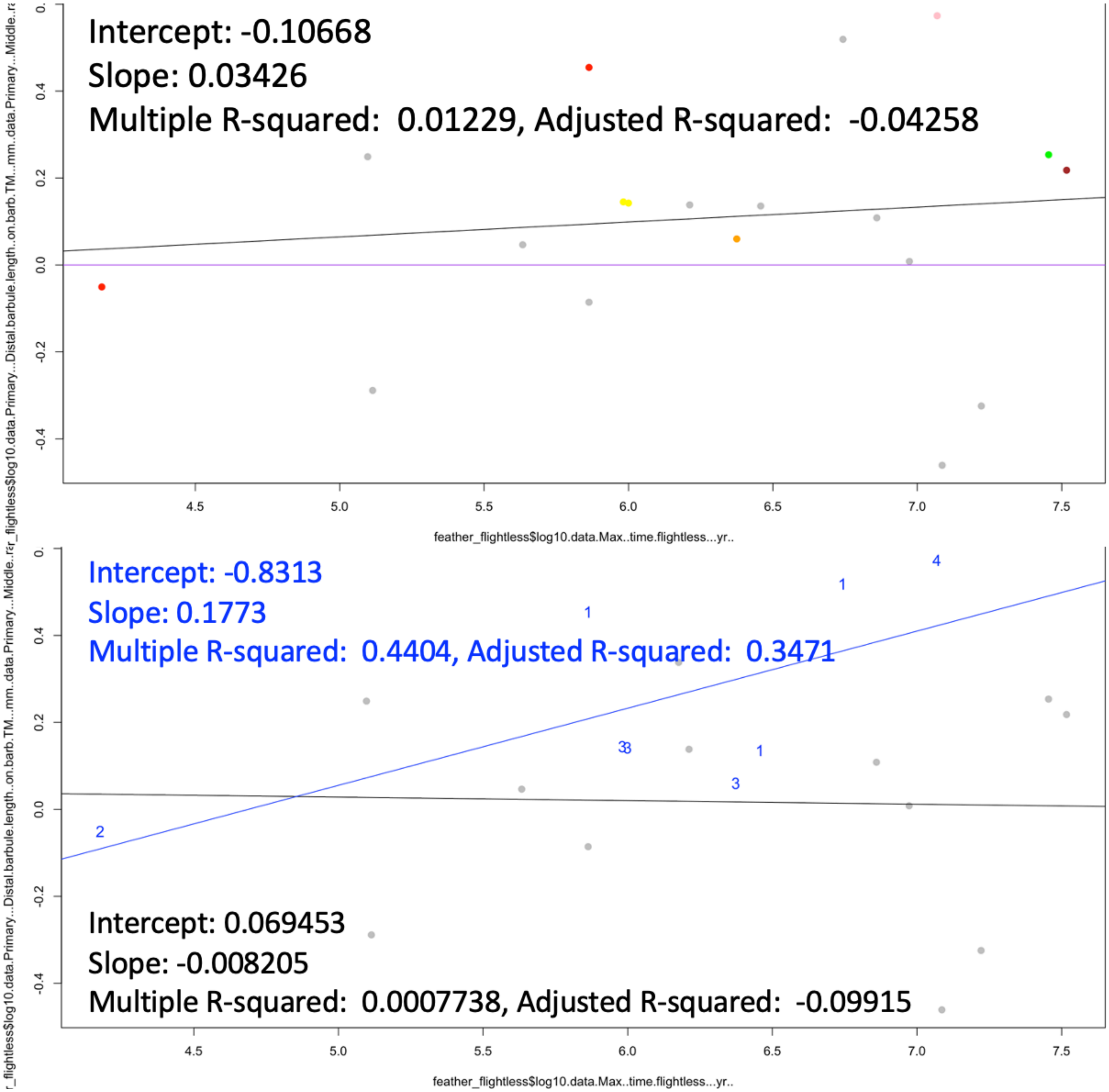

Figure S101. Shift in $\log_{10}$-tranformed distal barbule length/rachis length (flightless minus volant sister taxon comparisons) versus divergence age with penguins and ratites removed. Top: clade colors as in plots above. Bottom: blue points are semiaquatic, while black are terrestrial. Linear regression results shown.

#### 7.2.5. Microscopic trait pairwise regression summary

Rachis width generally shifts negatively from the volant ancestor – notably decreasing in terrestrial taxa, although there is a less negative shift in semiaquatic taxa. The reversed correlations seen without the 'long branches' of ratites and penguins are likely flukes of low sample size, since the data points representing recently diverging semiaquatic flightless taxa are more positive than terrestrial flightless taxa of similar age. This possibly shows that the pattern in rachis width upon flight loss is more apparent after long divergence times and specialized adaptation to non-flight.

Barb length increases with flightlessness generally, more so in the leading than the trailing vane because of high vane asymmetry in volant ancestors. Early evolution may show more rapid barb length increase in terrestrial flightless taxa than in semiaquatic flightless taxa. Late evolution may show a reversal of this trend – ratites eventually lose barbs, while penguins have long barbs relative to their rachis length.

Barbule length tends to increase with flightlessness, but this trend is likely not as prevalent as in barb lengths. There may be a strong shift in semiaquatic flightless taxa, even in early in their evolution. The ratite pattern is diverse – if they retain barbules, the barbules tend to be long; otherwise, barbules are lost entirely.

### 7.3. Comparison of pairwise regression slopes and $R^2$

#### 7.3.1. Absolute value of the regression slope

| Metric | Bin | Absolute value of slope, Long branches |
|---|---|---|
| Mass | Terr. | 0.4676 |
| Tail | Aqua. | 0.466 |
| Wing | Aqua. | 0.4426 |
| Mass | All | 0.42401 |
| Wing | All | 0.39271 |
| Mass | Aqua. | 0.3894 |
| Wing | Terr. | 0.3862 |
| Tail | All | 0.34156 |
| Tail | Terr. | 0.27458 |
| Barbule | Aqua. | 0.24759 |
| Barb L | Aqua. | 0.1656 |
| Barbule | All | 0.15651 |
| Rachis W | Terr. | 0.08879 |
| Barbule | Terr. | 0.08139 |
| Barb T | Aqua. | 0.07647 |
| Barb T | Terr. | 0.05933 |
| Barb L | All | 0.05154 |
| Barb L | Terr. | 0.03776 |
| Rachis W | Aqua. | 0.019878 |
| Rachis W | All | 0.007466 |
| Barb T | All | 0.003552 |

Table S3. Absolute value of regression slopes with penguins and ratites included for macroscopic (blue) and microscopic (red) traits.

| Metric | Bin | Absolute value of slope, No long branches |
|---|---|---|
| Tail | Aqua. | 0.2192 |
| Wing | Aqua. | 0.1809 |
| Barbule | Aqua. | 0.1773 |
| Mass | Aqua. | 0.1725 |

| | | |
|---|---|---|
| **Mass** | **Terr.** | **0.1619** |
| **Mass** | **All** | **0.14984** |
| **Tail** | **All** | **0.1457** |
| **Wing** | **All** | **0.11637** |
| **Tail** | **Terr.** | **0.10719** |
| **Barbule** | **All** | **0.10668** |
| **Wing** | **Terr.** | **0.106** |
| **Barb L** | **Terr.** | **0.07536** |
| **Barb L** | **All** | **0.07242** |
| **Rachis W** | **Terr.** | **0.07118** |
| **Barb T** | **Terr.** | **0.02558** |
| **Rachis W** | **Aqua.** | **0.02464** |
| **Rachis W** | **All** | **0.02243** |
| **Barb T** | **All** | **0.01887** |
| **Barb L** | **Aqua.** | **0.01848** |
| **Barbule** | **Terr.** | **0.008205** |
| **Barb T** | **Aqua.** | **0.002853** |

Table S4. Absolute value of regression slopes with penguins and ratites removed for macroscopic (blue) and microscopic (red) traits.

#### 7.3.2. $R^2$ value of the regression

| **Metric** | **Bin** | **$R^2$, Long branches** |
|---|---|---|
| **Barbule** | **Aqua.** | **0.7254** |
| **Tail** | **Aqua.** | **0.6859** |
| **Wing** | **Aqua.** | **0.6617** |
| **Mass** | **Aqua.** | **0.6061** |
| **Barb L** | **Aqua.** | **0.5627** |
| **Wing** | **All** | **0.5024** |
| **Tail** | **All** | **0.4934** |
| **Mass** | **All** | **0.4804** |
| **Wing** | **Terr.** | **0.4651** |
| **Mass** | **Terr.** | **0.4524** |
| **Tail** | **Terr.** | **0.4205** |
| **Barb T** | **Aqua.** | **0.4057** |
| **Rachis W** | **Aqua.** | **0.3192** |
| **Barbule** | **All** | **0.2193** |
| **Rachis W** | **Terr.** | **0.08588** |
| **Barb T** | **Terr.** | **0.08576** |
| **Barbule** | **Terr.** | **0.07525** |
| **Barb L** | **All** | **0.05528** |
| **Barb L** | **Terr.** | **0.02939** |
| **Barb T** | **All** | **0.0004124** |

| | | |
|---|---|---|
| Rachis W | All | 0.000405 |

Table S5. $R^2$ of regressions with penguins and ratites included for macroscopic (blue) and microscopic (red) traits.

| Metric | Bin | Adjusted $R^2$, Long branches |
|---|---|---|
| Barbule | Aqua. | 0.6949 |
| Tail | Aqua. | 0.651 |
| Wing | Aqua. | 0.6242 |
| Mass | Aqua. | 0.5624 |
| Barb L | Aqua. | 0.5141 |
| Wing | All | 0.4833 |
| Tail | All | 0.4739 |
| Mass | All | 0.4604 |
| Wing | Terr. | 0.4295 |
| Mass | Terr. | 0.4159 |
| Tail | Terr. | 0.3818 |
| Barb T | Aqua. | 0.3397 |
| Rachis W | Aqua. | 0.2436 |
| Barbule | All | 0.1853 |
| Rachis W | Terr. | 0.02494 |
| Barb T | Terr. | 0.02046 |
| Barb L | All | 0.01749 |
| Barbule | Terr. | -0.00181 |
| Rachis W | All | -0.03804 |
| Barb T | All | -0.03957 |
| Barb L | Terr. | -0.03994 |

Table S6. Adjusted $R^2$ of regressions with penguins and ratites included for macroscopic (blue) and microscopic (red) traits.

| Metric | Bin | $R^2$, No long branches |
|---|---|---|
| Barbule | Aqua. | 0.4404 |
| Tail | Aqua. | 0.4118 |
| Rachis W | Aqua. | 0.3069 |
| Wing | Aqua. | 0.3028 |
| Mass | Aqua. | 0.281 |
| Tail | All | 0.2207 |
| Mass | Terr. | 0.1973 |
| Mass | All | 0.188 |
| Barb L | All | 0.157 |
| Barb L | Terr. | 0.1563 |
| Wing | All | 0.1526 |
| Wing | Terr. | 0.1489 |
| Tail | Terr. | 0.1279 |
| Barb L | Aqua. | 0.123 |

| | | |
|---|---|---|
| **Rachis W** | **Terr.** | **0.0788** |
| **Barb T** | **Terr.** | **0.05029** |
| **Rachis W** | **All** | **0.0381** |
| **Barb T** | **All** | **0.03567** |
| **Barbule** | **All** | **0.01229** |
| **Barb T** | **Aqua.** | **0.001714** |
| **Barbule** | **Terr.** | **0.0007738** |

Table S7. $R^2$ of regressions with penguins and ratites removed for macroscopic (blue) and microscopic (red) traits.

| **Metric** | **Bin** | **Adjusted $R^2$, No long branches** |
|---|---|---|
| **Barbule** | **Aqua.** | **0.3471** |
| **Tail** | **Aqua.** | **0.3138** |
| **Rachis W** | **Aqua.** | **0.1914** |
| **Wing** | **Aqua.** | **0.1866** |
| **Tail** | **All** | **0.1774** |
| **Mass** | **Aqua.** | **0.1612** |
| **Mass** | **All** | **0.1429** |
| **Mass** | **Terr.** | **0.117** |
| **Barb L** | **All** | **0.1102** |
| **Wing** | **All** | **0.1055** |
| **Barb L** | **Terr.** | **0.07188** |
| **Wing** | **Terr.** | **0.0638** |
| **Tail** | **Terr.** | **0.04064** |
| **Rachis W** | **Terr.** | **-0.01332** |
| **Rachis W** | **All** | **-0.01534** |
| **Barb T** | **All** | **-0.0179** |
| **Barb L** | **Aqua.** | **-0.02312** |
| **Barbule** | **All** | **-0.04258** |
| **Barb T** | **Terr.** | **-0.04468** |
| **Barbule** | **Terr.** | **-0.09915** |
| **Barb T** | **Aqua.** | **-0.1647** |

Table S8. Adjusted $R^2$ of regressions with penguins and ratites removed for macroscopic (blue) and microscopic (red) traits.

#### 7.3.3. Regression parameters summary

Typically, macroscopic anatomical shifts seem to occur earlier and are stronger indicators of flightlessness than microscopic anatomical shifts. However, barbule length and rachis width in semiaquatic taxa, and to a lesser extent barb length in terrestrial taxa, might change soon after flight loss when examined with regression under simple phylogenetic control.

### 8. Literature cited in the supplemental material


Birkhead, T., Russell, D., Garbout, A., Attard, M., Thompson, J. and Jackson, D., 2020. New insights from old eggs–the shape and thickness of Great Auk Pinguinus impennis eggs. *Ibis*, *162*(4), pp.1345-1354.

Boyer, A.G., 2008. Extinction patterns in the avifauna of the Hawaiian islands. *Diversity and Distributions*, *14*(3), pp.509-517.

Burga, A., Wang, W., Ben-David, E., Wolf, P.C., Ramey, A.M., Verdugo, C., Lyons, K., Parker, P.G. and Kruglyak, L., 2017. A genetic signature of the evolution of loss of flight in the Galapagos cormorant. *Science*, *356*(6341), p.eaal3345.

Dunning Jr, J.B., 2008. *CRC Handbook of Avian Body Masses*. CRC press.

Fish, F.E., 2016. Secondary evolution of aquatic propulsion in higher vertebrates: validation and prospect. *Integrative and comparative biology*, *56*(6), pp.1285-1297.

Fulton, T.L., Letts, B. and Shapiro, B., 2012. Multiple losses of flight and recent speciation in steamer ducks. *Proceedings of the Royal Society B: Biological Sciences*, *279*(1737), pp.2339-2346.

Garcia-R, J.C., Gibb, G.C. and Trewick, S.A., 2014. Deep global evolutionary radiation in birds: Diversification and trait evolution in the cosmopolitan bird family Rallidae. *Molecular Phylogenetics and Evolution*, *81*, pp.96-108.

Gaspar, J., Gibb, G.C. and Trewick, S.A., 2020. Convergent morphological responses to loss of flight in rails (Aves: Rallidae). *Ecology and Evolution*, *10*(13), pp.6186-6207.

Harrison, P., Sallaberry, M., Gaskin, C.P., Baird, K.A., Jaramillo, A., Metz, S.M., Pearman, M., O'Keeffe, M., Dowdall, J., Enright, S. and Fahy, K., 2013. A new storm-petrel species from Chile. *The Auk*, *130*(1), pp.180-191.

Hume, J.P. and Martill, D., 2019. Repeated evolution of flightlessness in Dryolimnas rails (Aves: Rallidae) after extinction and recolonization on Aldabra. *Zoological Journal of the Linnean Society*, *186*(3), pp.666-672.

Jarvis, E.D., Mirarab, S., Aberer, A.J., Li, B., Houde, P., Li, C., Ho, S.Y., Faircloth, B.C., Nabholz, B., Howard, J.T. and Suh, A., 2014. Whole-genome analyses resolve early branches in the tree of life of modern birds. *Science*, *346*(6215), pp.1320-1331.

Johnson, A. and Cézilly, F., *The Greater Flamingo*. London: Poyser, 1975.

Lee, D. S. and Walsh-McGee, M., 2020. White-tailed Tropicbird (*Phaethon lepturus*), version 1.0. In Birds of the World (S. M. Billerman, Editor). Cornell Lab of Ornithology, Ithaca, NY, USA. https://doi-org.ezp2.lib.umn.edu/10.2173/bow.whttro.01

Lennard, C.J., 1997. *The causes of avian extinction and rarity* (Master's thesis, University of Cape Town).

Li, C., Zhang, Y., Li, J., Kong, L., Hu, H., Pan, H., Xu, L., Deng, Y., Li, Q., Jin, L. and Yu, H., 2014. Two Antarctic penguin genomes reveal insights into their evolutionary history and molecular changes related to the Antarctic environment. *GigaScience*, *3*(1), pp.2047-217X.

Livezey, B.C., 1988. Morphometrics of flightlessness in the Alcidae. *The Auk*, *105*(4), pp.681-698.

Meiri, S. and Dayan, T., 2003. On the validity of Bergmann's rule. *Journal of biogeography*, *30*(3), pp.331-351.

Mitchell, K.J., Wood, J.R., Scofield, R.P., Llamas, B. and Cooper, A., 2014. Ancient mitochondrial genome reveals unsuspected taxonomic affinity of the extinct Chatham duck (Pachyanas chathamica) and resolves divergence times for New Zealand and sub-Antarctic brown teals. *Molecular Phylogenetics and Evolution*, *70*, pp.420-428.

Mitchell, K.J., Wood, J.R., Llamas, B., McLenachan, P.A., Kardailsky, O., Scofield, R.P., Worthy, T.H. and Cooper, A., 2016. Ancient mitochondrial genomes clarify the evolutionary history of New Zealand's enigmatic acanthisittid wrens. *Molecular phylogenetics and evolution*, *102*, pp.295-304.

Ng, C.S. and Li, W.H., 2018. Genetic and molecular basis of feather diversity in birds. *Genome biology and evolution*, *10*(10), pp.2572-2586.

Ogawa, L.M., Pulgarin, P.C., Vance, D.A., Fjeldså, J. and van Tuinen, M., 2015. Opposing demographic histories reveal rapid evolution in grebes (Aves: Podicipedidae). *The Auk: Ornithological Advances*, *132*(4), pp.771-786.

Olson, S.L., 1999. Laysan Rail (Porzana palmeri) and Hawaiian Rail (Porzana sandwichensis). *The birds of North America*.

Poole, A. (Editor), 2005. *The Birds of North American Online*: http://bna.birds.cornell.edu/BNA/. Cornell Laboratory of Ornithology, Ithaca, NY.

Prince, P.A., Huin, N. and Weimerskirch, H., 1994. Diving depths of albatrosses. *Antarctic Science*, *6*(3), pp.353-354.

Rheindt, F.E., Christidis, L., Kuhn, S., de Kloet, S., Norman, J.A. and Fidler, A., 2014. The timing of diversification within the most divergent parrot clade. *Journal of Avian Biology*, *45*(2), pp.140-148.

Ripley, S.D., Lansdowne, J.F. and Olson, S.L., 1977. *Rails of the World: a Monograph of the Family Rallidae*. David R. Godine Publisher.

Sheard, C., Neate-Clegg, M.H., Alioravainen, N., Jones, S.E., Vincent, C., MacGregor, H.E., Bregman, T.P., Claramunt, S. and Tobias, J.A., 2020. Ecological drivers of global gradients in avian dispersal inferred from wing morphology. *Nature communications*, *11*(1), pp.1-9.

Slack, K.E., Jones, C.M., Ando, T., Harrison, G.L., Fordyce, R.E., Arnason, U. and Penny, D., 2006. Early penguin fossils, plus mitochondrial genomes, calibrate avian evolution. *Molecular biology and evolution*, *23*(6), pp.1144-1155.

Slikas, B., Olson, S.L. and Fleischer, R.C., 2002. Rapid, independent evolution of flightlessness in four species of Pacific Island rails (Rallidae): an analysis based on mitochondrial sequence data. *Journal of Avian Biology*, *33*(1), pp.5-14.

Smith, N.A. and Clarke, J.A., 2014. Osteological histology of the Pan-Alcidae (Aves, Charadriiformes): correlates of wing-propelled diving and flightlessness. *The Anatomical Record*, *297*(2), pp.188-199.

Stervander, M., Ryan, P.G., Melo, M. and Hansson, B., 2019. The origin of the world’s smallest flightless bird, the Inaccessible Island Rail Atlantisia rogersi (Aves: Rallidae). *Molecular phylogenetics and evolution*, *130*, pp.92-98.

van de Crommenacker, J., Bunbury, N., Jackson, H.A., Nupen, L.J., Wanless, R., Fleischer-Dogley, F., Groombridge, J.J. and Warren, B.H., 2019. Rapid loss of flight in the Aldabra white-throated rail. *PloS one*, *14*(12), p.e0226064.

Yonezawa, T., Segawa, T., Mori, H., Campos, P.F., Hongoh, Y., Endo, H., Akiyoshi, A., Kohno, N., Nishida, S., Wu, J. and Jin, H., 2017. Phylogenomics and morphology of extinct paleognaths reveal the origin and evolution of the ratites. *Current Biology*, *27*(1), pp.68-77.
